# Trade-off between foraging and vigilance in the Red-legged Seriema

**DOI:** 10.64898/2026.07.30.741728

**Authors:** Lotte Schroth, Leonardo Lopes, Filipe C.R. Cunha

## Abstract

As foraging increases exposure to predators, individuals must allocate time to vigilance in addition to food acquisition. Although those behaviors are assumed to be mutually exclusive, this is often hard to disentangle, given that multiple sensory modalities are often applied. However, when foraging, animals often have one sensory modality more impaired (e.g. vision), than others (e.g. auditory); thus, a level of compromise on vigilance capabilities is assumed while foraging. Here, we test the idea that when a food item requires manipulation, thus reducing vigilance capabilities, social foraging should promote shared costs of vigilance. We conducted video recordings to investigate whether there is a trade-off between foraging and vigilance behavior in red-legged seriemas (*Cariama cristata*) when foraging in pairs. We studied the social foraging behavioral patterns while seriemas were feeding on macauba palm (*Acrocomia aculeata*) fruit, which requires a series of throwing maneuvers to crack the hard outer shell. Our results showed that within a pair recording one individual was observed to spend more time vigilant than their partner, while the other individual spent more time throwing fruit than their partner. However, we found no evidence for synchronization or coordination of vigilance and foraging between individuals within a feeding bout. These results provide insight into the foraging ecology of red-legged seriemas and the trade-off between foraging and vigilance in social contexts.

## Introduction

Foraging is a costly activity that involves movement through the landscape, and occasionally, a level of manipulation of the food item. As searching for food exposes individuals to threats, being vigilant is a way for individuals to be aware of their surroundings and early detect potential predators (McNamara and Houston 1992, Linson et al. 2026), and/or kleptoparasites (Goss-Custard et al. 1999, Ridley et al. 2007). Although vigilance is often assumed to be mutually exclusive to foraging behavior (Barnard 1980, Beauchamp 1998), evidence suggests that these behaviors are not always incompatible, but that certain aspects of foraging, such as chewing in herbivores, can take place simultaneously with vigilance (Olson et al. 2015, Sirot et al. 2021). The compromise of certain sensory modalities (e.g. vision) while foraging influences the perception of the risk of being surprised (Tisdale et al. 2009, Pays et al. 2011). Thus, manipulating complex items should enhance vigilance and cooperation in social foraging. In social foraging events, individuals often share vigilance duties, and the presence of conspecifics should reduce the cost of vigilance per individual, thereby increasing the net energetic gain on foraging (Powell 1974, Cowlishaw et al. 2004, Ward et al. 2011).

This trade-off has often been suggested in socially living species, where vigilance is assumed to be shaped by the presence of conspecifics. The “many-eyes” hypothesis suggests that individuals living in groups can benefit from the alertness of their group members, thereby lowering an individual’s need to contribute to group vigilance (Lima & Dill 1990). Collective vigilance in groups has been proposed as a selection pressure for gregarious foraging (Olson et al. 2015). The trade-off between foraging and vigilance behavior is also seen in species with a sentinel system, where only one or a few individuals are vigilant at a given time, looking out for the whole group (McGowan & Woolfenden 1989). This has been observed in a handful of species, such as meerkats (*Suricata suricatta*) (Clutton-Brock et al. 1999) and Arabian babblers (*Argya squamiceps*) (Ostreiher et al. 2021).

Many bird species, however, forage in pairs; a study on 216 different Neotropical bird species found that 26.6% of them foraged in pairs (Thiollay & Julien 2008). Apart from pairs potentially promoting the reduction of individual vigilance, due to increased safety with an extra pair of eyes, they can also lead to behavioral differences between sexes. In pairs of white-tailed ptarmigans (*Lagopus leucurus*), for example, females were more likely to initiate foraging after their mate became vigilant (Artriss & Martin 1995). In this situation, male vigilance is believed to facilitate an increase in foraging opportunities for females. This pattern is observed more often in pairs, with males investing more time and energy in vigilance and females performing more foraging, potentially to acquire resources for egg production (Lou et al. 2017). Division of labor between males and females in foraging and vigilance again reinforces the idea of a trade-off between these two behaviors. Pairs represent the smallest social unit, with many bird species forming long-term pair bonds with their partner, and longer pair bonds can increase breeding success (Fowler 1995, Sánchez-Macouzet et al. 2014, Teitelbaum et al. 2017). Examining how foraging and vigilance behavior is distributed within pairs can help further understand the structure of these social units.

Pair-living organisms that share responsibility in survival and the rearing of offspring should rely on each other during recurrent behaviors (Roth et al. 2021), such as vigilance, which can be expressed either in the form of synchronization of coordination (Pays et al. 2008, Ge et al. 2011, Van Rooij and Griffith 2013, Brandl and Bellwood 2015). Synchronization of vigilance occurs when individuals overlap their vigilance more than expected by chance (Öst and Tierala 2011, Brügger et al. 2022). This way, individuals can minimize instances in which they are the least vigilant in a group, making them less of a target for predators (Sirot and Touzalin 2008). Synchronization also allows individuals to assess threats together and make faster decisions concerning the risks (Podgórski et al. 2016). However, synchronization increases the time in which no individual is vigilant and raises the risk that predator detection is missing at these moments (Ge et al. 2011). Coordination, on the other hand, occurs when individuals minimize the overlapping of their vigilance, thus maximizing the time at least one individual is alert and increasing predator detection (Rodríguez-Gironés and Vásquez 2002). If individuals coordinate their vigilance, the observed amount of time a group spends on this behavior should be higher than what is expected under the assumption that the behavior of individuals is independent (Brügger et al. 2022). For behavior to be synchronized, this observed amount of time vigilant should be lower than that expected under independence. Coordination is thought to be favored in smaller groups, as fewer group members make information sharing more efficient (Rodríguez-Gironés and Vásques 2002, Fernández-Juricic et al. 2004, Ge et al. 2011). Coordination could, therefore, also be found in some species that forage in pairs (Brandl and Bellwood 2015).

To investigate the trade-off between vigilance and foraging activities in a pair-living species, we conducted a field study by recording pairs of the red-legged seriema (*Cariama cristata*) foraging in the wild. The red-legged seriema is a large-bodied (∼2.3 Kg), terrestrial bird, widespread in South America, including most of Argentina, Paraguay, Bolivia and Brazil excluding the Amazonia (Jones et al. 2024). It is one of the two members of the Neotropical family Cariamidae. Red-legged seriemas have been observed to both forage with their partner as well as on their own (Jones et al. 2024). Although this is considered a common species, little is known about their behavior and ecology (Silva et al. 2016). Red-legged seriemas are opportunistic feeders, relying on the varying availability of arthropods, small vertebrates, seeds, and fruits (Schubart et al. 1965, Redford and Peters 1986, Silva et al. 2016). A peculiar behavior of red-legged seriemas is that they kill their prey by throwing them onto hard surfaces (Jones et al. 2024). In red-legged seriemas, this throwing behavior is also performed to crack open macauba palm (*Acrocomia aculeata*) fruits. Macauba fruit is a seasonal food source, with the peak of production happening in November (Scariot and Lleras 1991). The high fat content of the fruit is what makes it a valuable food resource, but due to the hard epicarp layer, the fruit requires specific manipulation to be open (Sorita et al. 2015, Lescano et al. 2021). The throwing technique of the red-legged seriema allows them to crack this hard outer layer, exposing the nutritious mesocarp of the fruit. By feeding on macauba fruit, red-legged seriemas play a potentially important role in the dispersal and therefore life cycle of the macauba palm (Dario 2020).

Here we tested whether individuals of red-legged seriemas differ from each other in foraging and vigilance behavior while foraging in pairs. If a cross-pair trade-off between foraging and vigilance is present, we expected one individual of the pair to spend more time foraging, while the other would spend more time vigilant (Artriss and Martin 1995, Lou et al. 2017) within a feeding event. Furthermore, we expected individuals to coordinate their vigilance behavior when foraging together as a way to maximize predator detection (Rodríguez-Gironés and Vásquez 2002, Wang et al. 2020). To investigate these ideas, we conducted field-based observational research at the University of Viçosa (UFV) in Florestal, Minas Gerais, Brazil.

## Methods

With the use of wildlife video cameras, we collected field-based observational data at the campus of the University of Viçosa (UFV) in Florestal, Minas Gerais, Brazil (19°52’29’’S, 44°25’12’’W). The study area has a size of about 1,500 ha, half of it covered by open habitats, including man-made pastures, crops, and gardens, which are used by the species. At least seven pairs of red-legged seriemas were present in our study area and their territories had been estimated in previous field observations. In four of these territories, a wildlife camera was positioned. Observations were made from mid-November until the end of December, coinciding with the fruiting season of macauba palms. A total of 48 days of non-stop footage was recorded, resulting in approximately 2700 hours of daylight footage. Of this, 48 hours contained seriemas interacting with macauba fruit.

The seriemas were not ringed, hence individual identification was not possible. The aforementioned territories were established by following seriemas across the study area and incorporating knowledge of the terrain, as well as boundaries being determined with the use of Minimum Complex Polygon (MCP) and GPS coordinates of individuals observed on a given day. Without the possibility of individual identification, these alleged territories provided a guide to the distinction between pairs’ home ranges. Although the territory delimitation has no major implications on the interpretation of the results in this study, any further speculation regarding landscape use should be refrained from until more data is available. Because of this, we will from here on out be referring to camera locations in different areas, rather than territories. Since individuals could not be recognized nor sexed, as this a sexually monomorphic species, it was not possible to track individual behaviors across recordings. Analyses were restricted to behavioral differences within a given recording, rather than individual differences across contexts. Patterns within actual long-term pairs could thus not be investigated, only patterns within specific foraging instances of two individuals.

We recorded behavior at four of seven areas in which we had evidence of pairs nesting previously. We used Green Feathers solar-powered Wi-Fi cameras (Green Feathers, 2025) for the observational research. Camera locations were based on nest sites for three of the pairs (Area 2, Area 3, and Area 4). Moreover, these were locations with flat ground areas and little vegetation that obstructed the camera view. At these locations, we placed multiple rocks, which functioned as cracking anvils. For the fourth area (i.e. Area 1), the camera’s location was based on cracking behavior recorded in earlier seasons (E. Angelli, personal communication, November 12, 2025). At this location, no additional rocks were placed as there were already rocks present. For all locations, macauba fruit was hand-placed every other day, ensuring there were between 15 and 20 pieces of fruit present every time.

The Green Feathers cameras, in combination with corresponding solar panels, were able to record up to three days continuously. At the same moment as placing new macauba fruit at locations, SD cards were switched out, and cameras and solar panels were replaced with fully charged ones. To extract media files from the cameras as MP4 files, we used the code critvid (Zukas 2026), which allowed us to split videos into approximately 50-minute-long clips. We only extracted recordings that took place between 5:00 and 19:00, as foraging behavior was only assumed during daytime, given that seriemas sleep during the night period.

To check videos for the presence of red-legged seriemas, we also developed a “Seriema Detection Model”. Frames were extracted from previous videos in the area (S. Trijp, personal communication, April 4, 2025) with the ‘animl’ package (CRAN 2026) in RStudio (Posit team, 2025). These frames contained seriemas as well as other species and empty images and were labelled with the LabelMe program (Russel et al. 2007) in Python (https://www.python.org/). These labels were divided into a training (184 images), validation (52 images), and test (28 images) set. To build the model, Ultralytics YOLOv8 (Ultralytics, 2023) was used (see Appendix 1 for detailed information on model construction). The model had a strong performance, with a precision of 93.7% and a recall of 89.5%. However, due to the model’s lack of speed and hardware limitations, a large part of the video selection for this study was still performed manually. All videos with at least one red-legged seriema present were used for behavioral scoring.

We used BORIS as a behavior scoring tool (Friard and Gamba 2016). As no ethogram concerned with macauba foraging existed yet, we built one during the early stage of the video analyses.

For this research, six different behaviors were classified: pecking fruit, picking up fruit, eating fruit, throwing fruit, dropping fruit and vigilance (Supplementary material Table S1). To study seriemas in a foraging context, only videos with at least one interaction with macauba fruit (picking up fruit, eating fruit, pecking fruit, throwing fruit, or dropping fruit) were selected. For each video, we also noted the presence of individuals from the moment they walked into the camera frame until they left. If individuals entered the frame but did not interact with the fruit within a minute, presence was recorded to begin one minute prior to the first fruit interaction. Similarly, presence was ended if all individuals spent more than a minute without a fruit interaction. This was done to standardize different locations, as not all cameras had equal viewing ranges, and to limit analyses to seriemas in a foraging context. The one-minute cut-off was chosen on the premise that seriemas were often observed not to interact with the fruit anymore after a minute since the last interaction had passed.

### Data preparation

Social context was added as an additional variable to structure the data. Behaviors observed in videos with only one individual were labelled “alone”, whereas behaviors in videos where there were two individuals present at any given moment were labelled “pair”. There were no observations with more than two individuals. For the analysis, we integrated all behaviors classified as ‘alone’ and for the behaviors classified as ‘pair’, we only kept the observations that took place when both individuals were present in the video frame. Furthermore, all observations where individuals displayed neither vigilance, eating, nor throwing behavior were removed, to control for false zeros associated with seriemas not being interested in the macauba fruit (Blasco-Moreno et al. 2019).

We further adjusted the variables vigilance, eating, and throwing, taking into account the varying duration of the observations. The amount of time an individual spent vigilant within a recording was summed and divided by the recording’s duration. The same was done for the amount of time individuals spent eating within a recording. As both vigilance and eating are continuous behaviors, this gave us the proportion of time individuals spent vigilant (VP) and eating (EP) per recording. For throwing, we calculated the throwing rate (TR) by dividing the total throws of an individual per recording by the duration of that recording in minutes.

Additionally, we calculated the difference in behavior between the two individuals in a recording. This was done for the difference in vigilance proportions (VP_Adult 1_ – VP_Adult 2_), the difference in eating proportion (EP_Adult 1_ – EP_Adult 2_), and the difference in absolute sum of throws (T_Adult 1_ – T_Adult 2_). Within each pair recording, individuals were labelled according to their behavioral contribution, with the individual that exhibited the higher value of a giving behavior labelled as “More” and the individual that exhibited the lower value of this behavior labelled as “Less”. These variables made it possible to compare differences in behavior between individuals within a specific recording.

Furthermore, we calculated both the predicted and observed probabilities of vigilance and eating behavior within a recording of a pair. For vigilance, the observed probability was quantified as the total amount of time at least one of the individuals was vigilant divided by the total time both individuals were present. For eating behavior, observed probabilities were calculated in the same way. We used the following formula to calculate predicted probabilities for vigilance and eating behavior per recording of a pair:

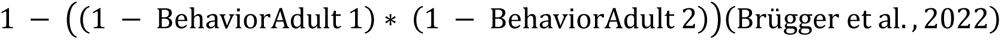

Predicted probability is described as the chance that at least one of the individuals is performing the behavior of interest, under the assumption that individuals execute their behavior independently from each other. If the observed probability is lower than the predicted probability, individual behavior is not independent, and pairs synchronize their behavior. However, if the observed probability is higher than the predicted probability, individuals coordinate their behavior. If there is no difference between observed and predicted probabilities, the behavior of individuals is considered independent.

### Statistical analysis

To test whether both individuals in a recording differ in the time spent vigilant. We tested if the difference in vigilance between two individuals within a recording (VP_Adult 1_ – VP_Adult 2_) was significantly different from zero using a Generalized linear mixed model (GLMM) from the ‘glmmTMB’ package (Brooks et al., 2017). A Beta distribution was used as vigilance proportion is based on continuous data. Furthermore, location was added as a random factor. A similar model was used to test if the difference in eating behavior between two individuals in a recording (EP_Adult 1_ – EP_Adult 2_) deviated from zero, this time we used a zero-inflated Beta distribution in order to account for the many zeros in this data. To test if two individuals within a recording differed in their throwing rate, we constructed a GLMM with a Negative Binomial distribution. Here we used the difference in absolute throwing (T_Adult 1_ – T_Adult 2_) as the dependent variable and added observation time as an offset to account for different recording times. Again, location was added as a random factor, and the model was again zero-inflated.

Additionally, we tested whether the differences in behavior between individuals within a recording were related to each other. For this, we used another GLMM with the difference in eating proportion as a dependent variable and the difference in vigilance proportion as an independent variable. This model used a Gaussian distribution and had location as a random factor. A similar model was made with the difference in absolute throwing as the dependent variable and the difference in vigilance proportion as the independent variable but added observation time as an offset. To test if differences in throwing rate were related to differences in eating proportion within recordings of pairs, we made a GLMM with a Gaussian distribution and the difference in absolute throwing as the dependent variable, the difference in eating proportion as the independent variable and again an offset for observation time.

To test if individuals within a recording coordinated or synchronized their vigilance, we compared predicted and observed probabilities of these recordings. For this we used a GLMM with the probability of vigilance as a response variable and the context of this vigilance, either predicted or observed, as the predictor variable. A Beta distribution was used, and location was added as a random factor. A similar model was used to test the difference in predicted and observed values of eating behavior per recording.

For all models, we used the ‘DHARMa’ package (Hartig et al. 2024) to check model assumptions. Data and code are available in the supplementary material.

## Results

We registered a total of 20.25 hours of foraging events, with 13.70 hours of seriemas foraging in pairs. For one of the areas that was recorded, there was only one observation with the two individuals within frame, shorter than one minute; thus, we decided to exclude this area from the analysis. It is noteworthy that the amount of data collected per location varied between the three other areas (Supplementary material Table S2).

When in a pair, there was a division of task allocation between the individuals in a given recording, with one individual spending a smaller proportion of time vigilant (mean ± SD) (22% ± 1.2%) than the other (38% ± 2.1%) (Figure 1, Table 1). Individuals devoting less time to vigilance than their social partner spent a mean of 2.33 minutes vigilant in a given recording (min: 0 – max: 9.75 minutes), while individuals devoting more time to vigilance spent a mean of 3.69 minutes vigilant in that same recording (min: 0.02 – max: 19.1 minutes). A similar pattern was observed for feeding behavior, with one individual allocating approximately twice as much proportion of the observed time eating (29% ± 2.1%) as its partner (13% ± 1.6%) in a given recording (Figure 1; Table 1). Individuals eating more than their partner spent a mean of 3.31 minutes eating (min: 0 – max: 13.8 minutes), while individuals eating less spent a mean of 1.82 minutes eating in a given recording (min: 0 – max: 12.4 minutes). For throwing behavior, we also recorded an asymmetric distribution of time within the pairs in a given recording, with one individual throwing about six times more fruit (0.74 ± 0.064 throws per minute) than their partner (0.13 ± 0.029 throws per minute) in a given recording (Figure 1; Table 1). The individual throwing less in a recording had a mean of 1.15 throws per recording (min: 0 – max: 12 throws), while the individual devoting more time to throwing than their partner had a mean of 6.91 throws in that same recording (min: 0 – max: 41 throws).

**Figure 1:**
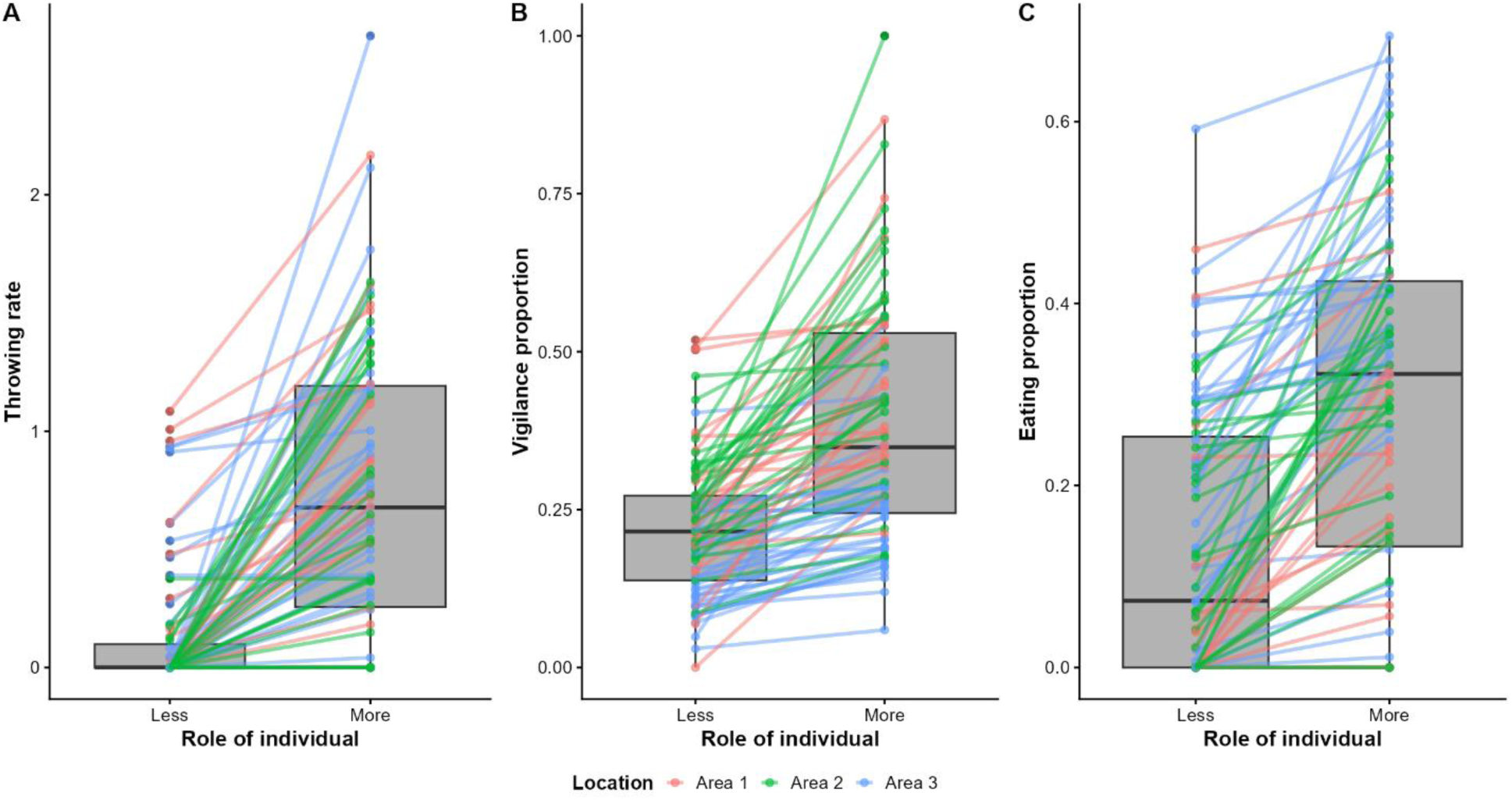
Boxplots displaying the differences in behavior between two individuals within a recording. A) the difference in throwing rate. B) the difference in proportion of time spent vigilant. C) the difference in proportion of time spent eating. Individuals were labelled Less or More, based on the amount of behavior they displayed compared to their partner in a recording. The colors indicate the different locations where recordings were made.

**Table 1:** Summary of Generalized linear mixed null models performed within recordings of pairs, testing if individuals within a pair differed from each other within a given recording. N= 87.

| Parameter | Estimate | SE | 95% CI | z-value | p-value |
| --- | --- | --- | --- | --- | --- |
| **Model 1: difference in proportion of time vigilant** |  |  |  |  |  |
| *(Beta distribution, Random factor: Location)* |  |  |  |  |  |
| Random factor SD | 0.32 |  |  |  |  |
| Intercept | -1.60 | 0.21 | [-2.01, -1.19] | -7.46 | < 0.0001 |
| **Model 2: difference in proportion of time eating** |  |  |  |  |  |
| *(Beta distribution, Zero-inflated, Random factor: Location)* |  |  |  |  |  |
| Random factor SD | < 0.0001 |  |  |  |  |
| Intercept | -1.50 | 0.11 | [-1.72, -1.28] | -13.52 | < 0.0001 |
| Intercept Zi | -1.65 | 0.29 | [-2.22, -1.08] | -5.66 | < 0.0001 |
| **Model 3: difference in throwing rate** |  |  |  |  |  |
| *(Negative binominal distribution, Zero-inflated, Random factor: Location)* |  |  |  |  |  |
| Random factor SD | < 0.0001 |  |  |  |  |
| Intercept | -4.5 | 0.1 | [-4.70, -4.30] | -47.35 | < 0.0001 |
| Intercept Zi | -3.34 | 1.34 | [-5.97, -0.71] | -2.50 | < 0.01 |

Moreover, we found a relationship between the difference in vigilance and the difference in throwing between individuals within a recording (Table 2). Individuals that spent more time vigilant (mean: 3.69, min: 0.02 – max: 19.1 minutes) than their partner (mean: 2.33, min: 0 – max: 9.75 minutes) in a given recording, spent less time throwing fruit (mean: 2.22, min: 0 – max: 19 throws) than their partner (mean: 5.84, min: 0 – max: 41 throws) in that same recording. If an individual was approximately ten percentage points (approximately 0.94 minutes) more vigilant than their partner, they were expected to throw 1.2 throws less than their partner in a given recording (Figure 2). We also found a relationship between the difference in vigilance and the difference in eating between individuals within a recording (Table 2). Individuals that spent more time vigilant (mean: 3.69, min: 0.02 – max: 19.1 minutes) than their partner (mean: 2.33, min: 0 – max: 9.75 minutes) in a given recording, spent less time eating fruit (mean: 2.01 min: 0 – max: 13.8 minutes) than their partner (mean: 3.11, min: 0 – max: 12.4 minutes) in that recording (Table 2). If an individual spent ten percentage points (approximately 0.94 minutes) more on vigilance behavior than their partner, they were expected to spend 4.2 percentage points less time on eating (approximately 0.39 minutes) than their partner in that recording (Figure 2). We found no relationship between the difference in eating and the difference in throwing between individuals within a recording (Table 2). Individuals that spent more time eating (mean: 3.31, min: 0 – max: 13.8 minutes) than their partner (mean: 1.82, min: 0 – max: 12.4 minutes), did not significantly differ from their partner in the amount of fruit they threw in a recording (mean: 4.36, min: 0 – max: 29 throws) (mean: 3.70, min: 0 – max: 41 throws) (Figure 2).

**Figure 2:**
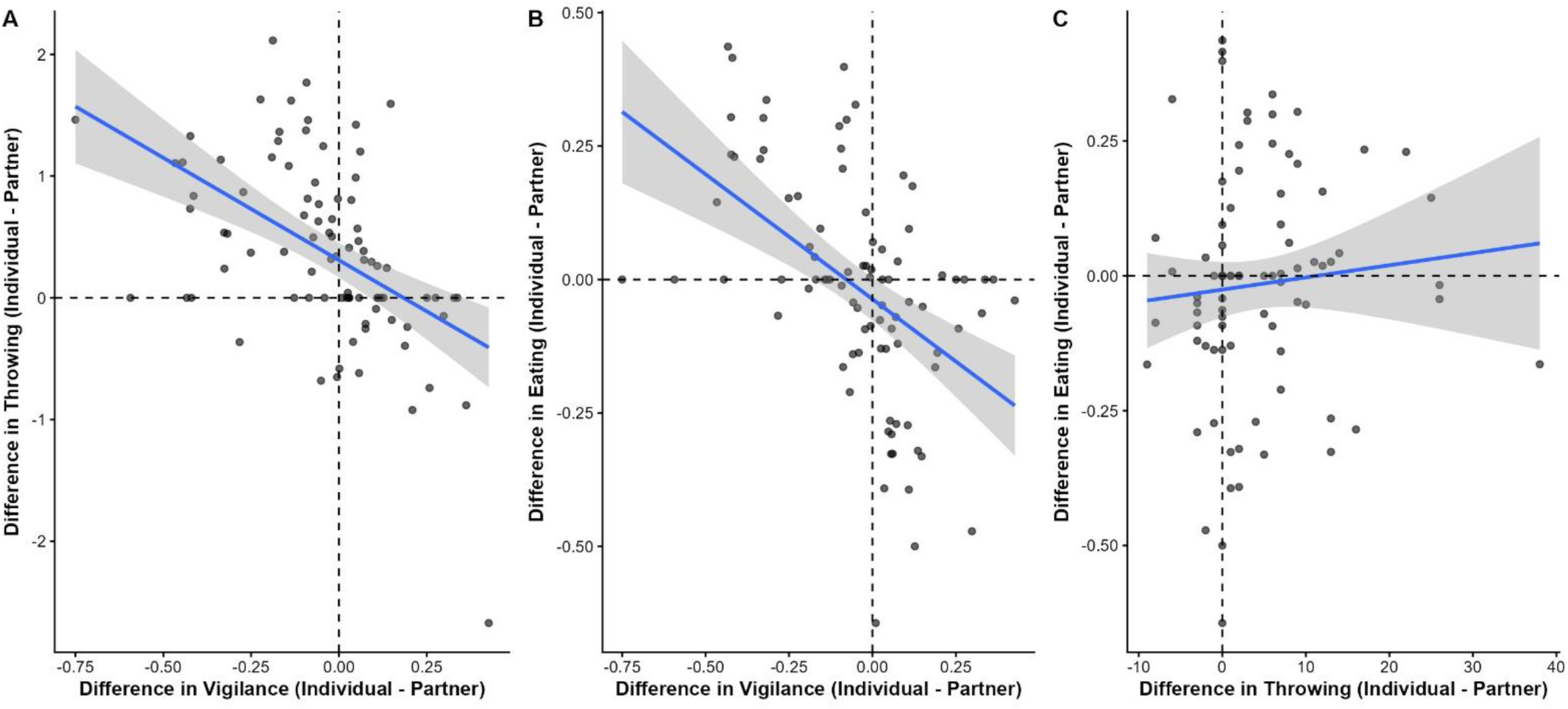
Scatterplot depicting the relation between the difference in behavior between individuals within a recording. A) The difference in vigilance proportion (x-axis) and the difference in total throwing amount (y-axis). There is a negative relation between these two factors, individuals that are more vigilant than their partner are throwing less than their partner in a given recording. B) The difference in vigilance proportion (x-axis) and the difference in eating proportion (y-axis). There is a negative relation between these two factors, individuals that are more vigilant than their partner are eating less than their partner in a given recording. C) The difference in eating proportion (x-axis) and the difference in total throwing amount (y-axis). There is no relation between these two factors. All plots are based on model predictions.

**Table 2:** Summary of Generalized linear mixed models performed within recordings of pairs, testing if differences in behavior are related to each other within a given recording. N= 83.

| Parameter | Estimate | SE | 95% CI | z-value | p-value |
| --- | --- | --- | --- | --- | --- |
| **Model 4: relation between the difference in throwing rate and the difference in vigilance proportion**<br>(Gaussian distribution, Random factor: Location) |  |  |  |  |  |
| Random factor SD | 2.39 |  |  |  |  |
| Intercept | -2.31 | 1.57 | [-5.39, 0.77] | -1.48 | 0.14 |
| difference vigilance | -11.92 | 3.52 | [-18.82, -5.02] | -3.39 | < 0.001 |
| **Model 5: relation between the difference in eating proportion and the difference in vigilance proportion**<br>(Gaussian distribution, Random factor: Location) |  |  |  |  |  |
| Random factor SD | 0.034 |  |  |  |  |
| Intercept | -0.030 | 0.029 | [-0.087, 0.027] | -1.04 | 0.30 |
| difference vigilance | -0.42 | 0.1 | [-0.62, -0.22] | -4.19 | < 0.0001 |
| **Model 6: relation between the difference in eating proportion and the difference in throwing rate**<br>(Gaussian distribution, Random factor: Location) |  |  |  |  |  |
| Random factor SD | 2.77 |  |  |  |  |
| Intercept | -1.64 | 1.77 | [-5.11, 1.83] | -0.93 | 0.35 |
| difference eating | 2.35 | 3.84 | [-5.18, 9.88] | 0.61 | 0.54 |

Furthermore, we found no difference between the observed probability of being vigilant (mean ± SD) (52% ± 6.7%) and the predicted probability (53% ± 6.7%), indicating there was no turn-taking in this behavior in a given recording (Supplementary material Table S1, Figure S2). There was also no turn-taking for eating behavior within a recording (Supplementary material Table S3, Figure S1), with observed probabilities of eating (42% ± 3.7%) being not statistically different from predicted probabilities of eating (42% ± 3.7%).

## Discussion

Within pairs of red-legged seriemas there seems to be an asymmetric effort allocation, with one individual being the most vigilant and spending approximately 60% more time vigilant than its partner within a recording. The other individual, in its turn, spent twice as much time eating and threw about six times more fruit within that same recording. Furthermore, within pair recordings, there was a negative relationship between vigilance and both eating and throwing behavior. If an individual spent approximately 0.94 minutes more on vigilance behavior than their partner within a recording, then that partner would throw about 1.2 fruits fewer and spent about 0.39 minutes less time eating than them. There was no relationship found between the difference in eating and the difference in throwing between two individuals within a recording. Moreover, no difference was found between predicted and observed probabilities for both vigilance and eating behavior in recordings of pairs, suggesting that seriemas in pairs are not coordinating nor synchronizing their behavior within a foraging session. It is noteworthy that individuals were not marked, limiting extrapolation of these results; therefore, interpretations must be taken with care. All analyses within pairs solely provide results for differences per feeding-recorded-bound.

The reduction of individual vigilance in larger groups is often linked to an increase in individual food intake, reflecting the often-assumed trade-off between vigilance and foraging (Beauchamp 1998), which has been demonstrated in multiple different species (Ruckstuhl et al. 2003, Cowlishaw et al. 2004, Favreau et al. 2014). Our research, however, focused on a pair setting, where the ratio of individuals vigilant and foraging is of a larger magnitude. Multiple studies have reported a division of labor between males and females in pairs when it comes to vigilance and foraging (Artiss and Martin 1995, Squires et al. 2007, Lou et al 2017). However, a study in zenaida doves (*Zenaida aurita*) showed that males and females do not differ from each other in their behavior when foraging in pairs (Cezilly and Keddar 2011). The main explanation for this finding was associated with the mutual benefits that individuals experience because of the long-term pair bonds zenaida doves have. Additionally, the lack of division of labor was linked to potentially lower levels of androgen in males, given that vigilance behavior in males might be related to higher levels of androgen (Dahlgren 1990) and that levels of androgen in male birds are generally lower in tropical species (Goymann et al. 2004, Hau et al. 2008). Hormonal variation could trigger certain levels of asymmetry in the trade-off between foraging and vigilance, suggesting that the breeding season progression might also influence this vigilance-foraging effort trade-off in social pairs.

Even though our findings showed a significant relationship between the difference in vigilance and the difference in both throwing and eating in pairs, there was no relationship between the difference in eating and the difference in throwing behavior between pair members in a recording. It is noteworthy that individuals might benefit from their partner opening macauba fruit, by consuming fruit without having to manipulate and crack the epicarp. Feeding on macauba fruit takes a lot of effort and time, because of this one fruit can offer multiple feeding opportunities for different individuals. We have observed individuals picking up and eating fruit that was opened by their partner, suggesting that not all individuals have to open fruit themselves in order to eat. If individuals can benefit from the throwing of their partner, this could further reinforce the idea of some level of division of labor between pair members, with one individual being more vigilant while the other individual opens fruit for both individuals to eat. Such coordination may rely on the ability to form partner-specific expectations based on repeated social interactions, allowing individuals to anticipate a partner’s behavior during foraging (Melis et al. 2006, Seyfarth & Cheney 2015). Whether this reflects prospective cognition or can be explained by learned expectations remains unresolved (Roberts 2002). However, the lack of relationship between eating and throwing could also be explained by other mechanisms, such as individuals opening more fruit than they can actually consume, and/or the fruit they open might not be of good quality. Similar behavior has been observed in blue jays (*Cyanocitta cristata L.*), which are more likely to not eat nuts they had opened if these were infested with weevils (Dixon et al. 1997).

Moreover, we found that there was no synchronization or coordination between individuals when it comes to vigilance and eating within a foraging session. Coordination is suggested to be the most effective collective way of detecting predators, as it minimizes the time during which no individuals are vigilant (Rodríguez-Gironés and Vásquez 2002). In an environment where there is low predation risk, it is costly to coordinate vigilance behavior, as this decreases foraging time without the actual need for it (Ge et al. 2011). There have been reports of seriemas being preyed on by Geoffroy’s cat (*Felis geoffroyi*) (Yanosky and Mercolli 1994), and other large felids and canids are considered potential predators. There have been at least two reports of domestic dogs killing juvenile seriemas in the study area. Overall, red-legged seriemas might experience low predation risks given the density of the predators in the area, therefore there is limited pressure to coordinate their vigilance. However, research on wild boars (*Sus scrofa*) found that groups can adjust their coordination or synchronization to ecological context and perceived risk (Podgórski et al. 2026). Red-legged seriemas might have a more dynamic level of coordination that fluctuates according to the immediate perceived risk.

Seriemas might still alternate vigilance across feeding sessions. If the individual that is more vigilant consistently eats less, seriemas could benefit from alternating behavioral roles between foraging sessions. There are multiple studies where turn-taking of vigilance has been proposed as a form of cooperation and direct reciprocity, with individuals sharing the cost of vigilance (Ridley et al. 2012, Brandl and Bellwood 2015, Brügger et al. 2023). In rabbitfishes (family Siganidae), for example, individuals that had repeated interactions with the same partner took turns in performing costly vigilance behavior (Brandl and Bellwood 2015). Direct reciprocity is defined as an individual helping a non-related individual and, in return, receiving delayed assistance from that same individual (Trivers 1971). In order for direct reciprocity to be possible, there should thus be repeated interactions between individuals (Trivers 1971). Also, long-term memory plays an important role in direct reciprocity, as it is dependent on the outcome of previous interactions with the same individual (Müller et al. 2017, Kettler et al. 2021). As seriemas are a long-lived, monogamous species, repeated interactions are thought to be common between individuals within a pair (Jones et al. 2024), but this idea of turn-taking and reciprocity remains untested in this system.

One of the main caveats of our research is the lack of identification of individuals. Therefore, individuals could be kept apart from each other within one observation but not across observations. The results suggest that there is an asymmetric effort allocation between individuals that are foraging together, but to fully understand if there is a persistent division of tasks between partners, or if behavioral roles within pairs are fixed or whether individuals switch roles between feeding sessions, multiple observations per pair over a longer period of time are needed. Additionally, sex differences might also play a role in the asymmetric allocation of time between foraging and vigilance in a pair setting. Evidence suggests that in multiple monogamous bird species, males tend to spend more time on vigilance behavior while females allocate more time to foraging (Powolny et al. 2014, Lou et al. 2017). As red-legged seriemas are also considered long-term monogamous (Jones et al. 2024), similar patterns might be present and fluctuate according to the breeding status.

There is still plenty unknown about the behavioral ecology of red-legged seriemas, but with this research we fill in some of the previous knowledge gaps. To our understanding this is the first research collecting large behavioral data on red-legged seriemas in the wild.

## AI Statement

AI was used as a supporting tool during this research. It aided in writing certain code for the statistical analyses and overcoming certain errors in RStudio. Furthermore, it played a supportive role in the construction of our Seriema Detection model. We used both ChatGPT (OpenAI 2026) and Claude (Anthropic 2025) as AI tools. It is important to state that the work of AI merely served as an assisting tool.

## Supporting information

Appendices containing all supplementary material, consisting of statistics, model description and supplementary tables/figures

