## Appendices containing all supplementary material, consisting of statistics, model description and supplementary tables/figures for "Trade-off between foraging and vigilance in the Red-legged Seriema": dataset.yaml

```
train: C:/Users/lotje/OneDrive/WUR/Thesis/Training data/seriemas/output model/train/images
val: C:/Users/lotje/OneDrive/WUR/Thesis/Training data/seriemas/output model/val/images
test: C:/Users/lotje/OneDrive/WUR/Thesis/Training data/seriemas/output model/test/images

nc: 1
names: ['seriema']
```
