## Supplementary figures and images for "Trade-off between foraging and vigilance in the Red-legged Seriema"

### 11080003-20734-302.jpg

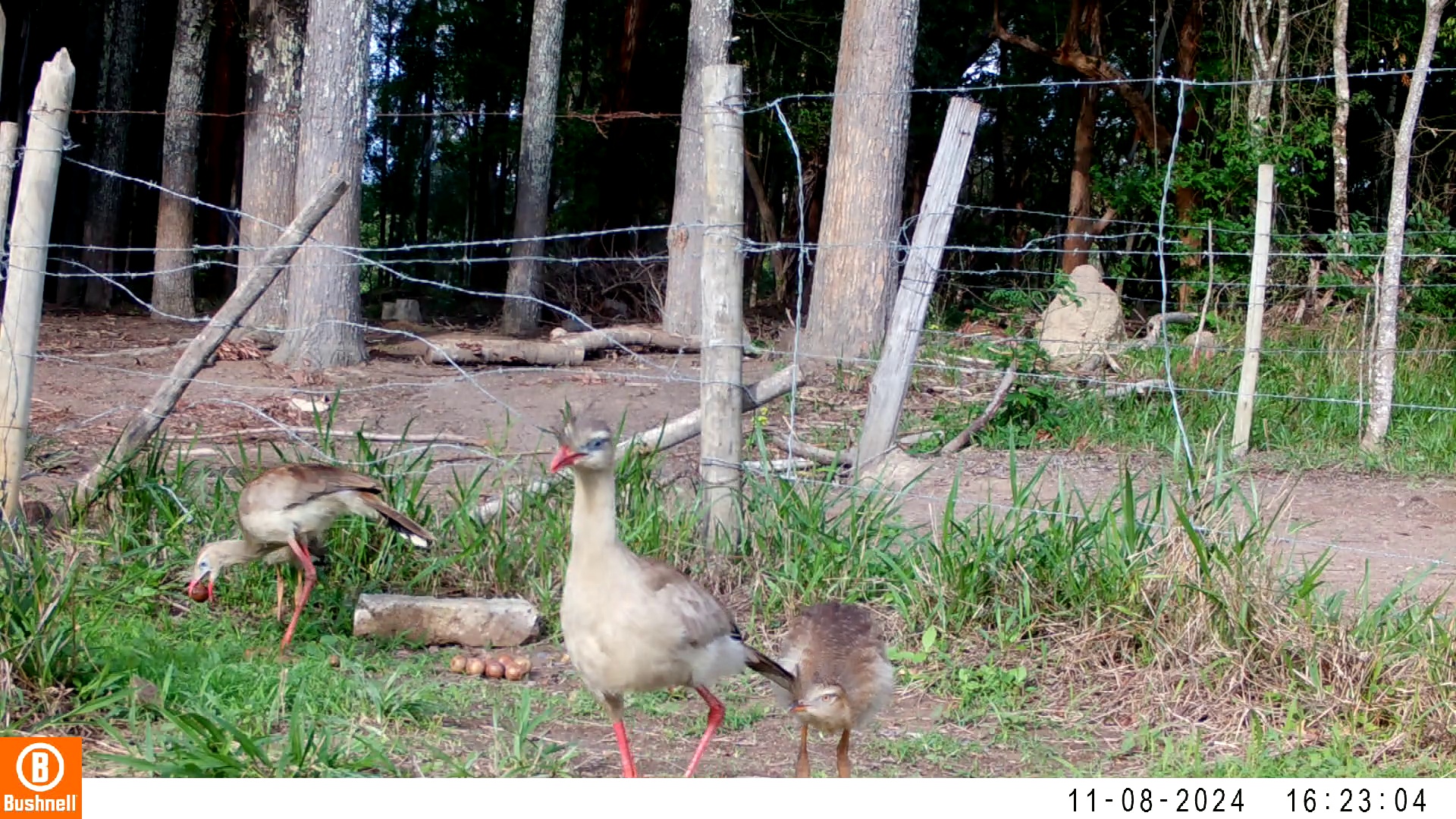

### 11080003-20734-604.jpg

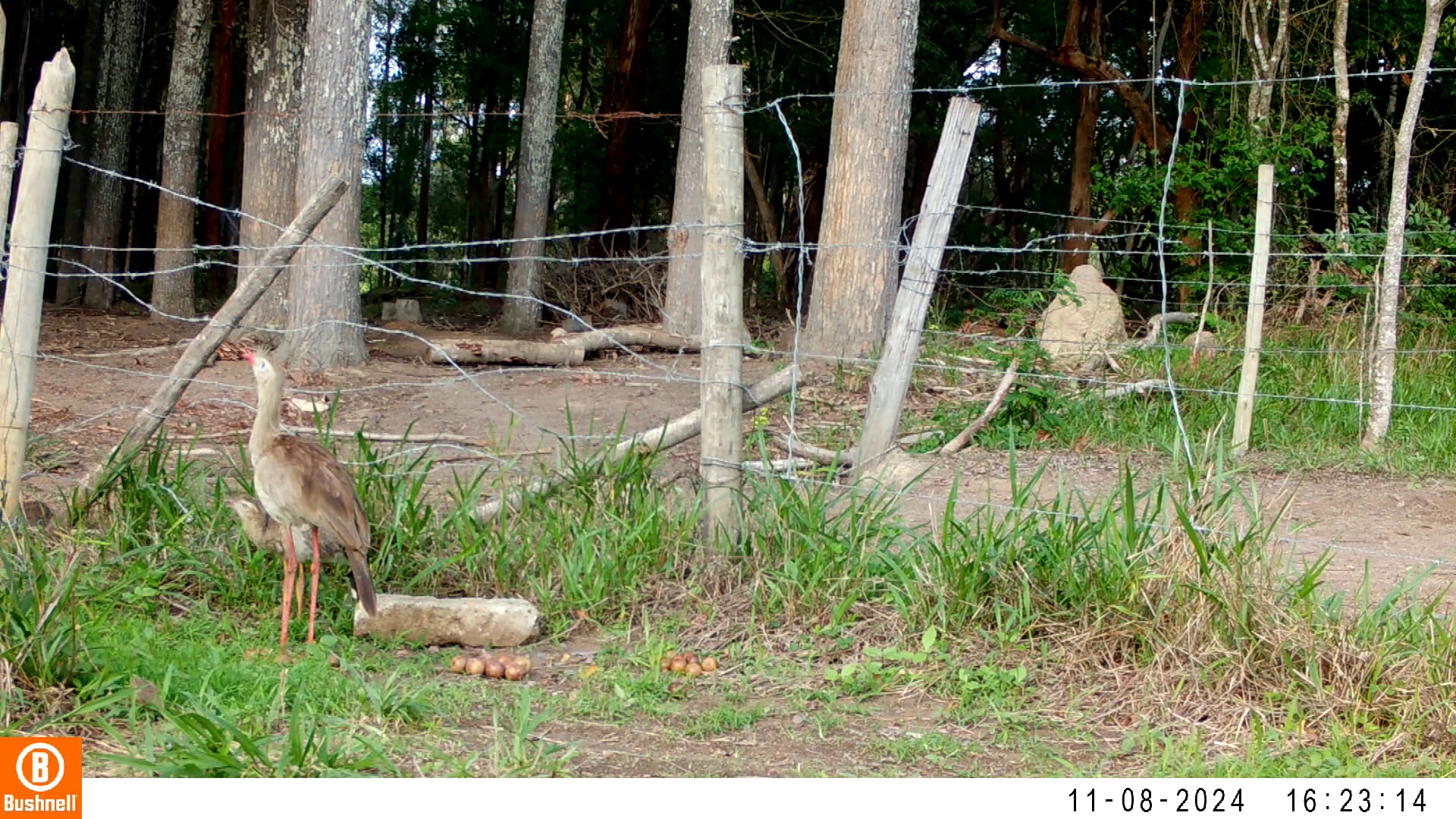

### 11130250-82886-0.jpg

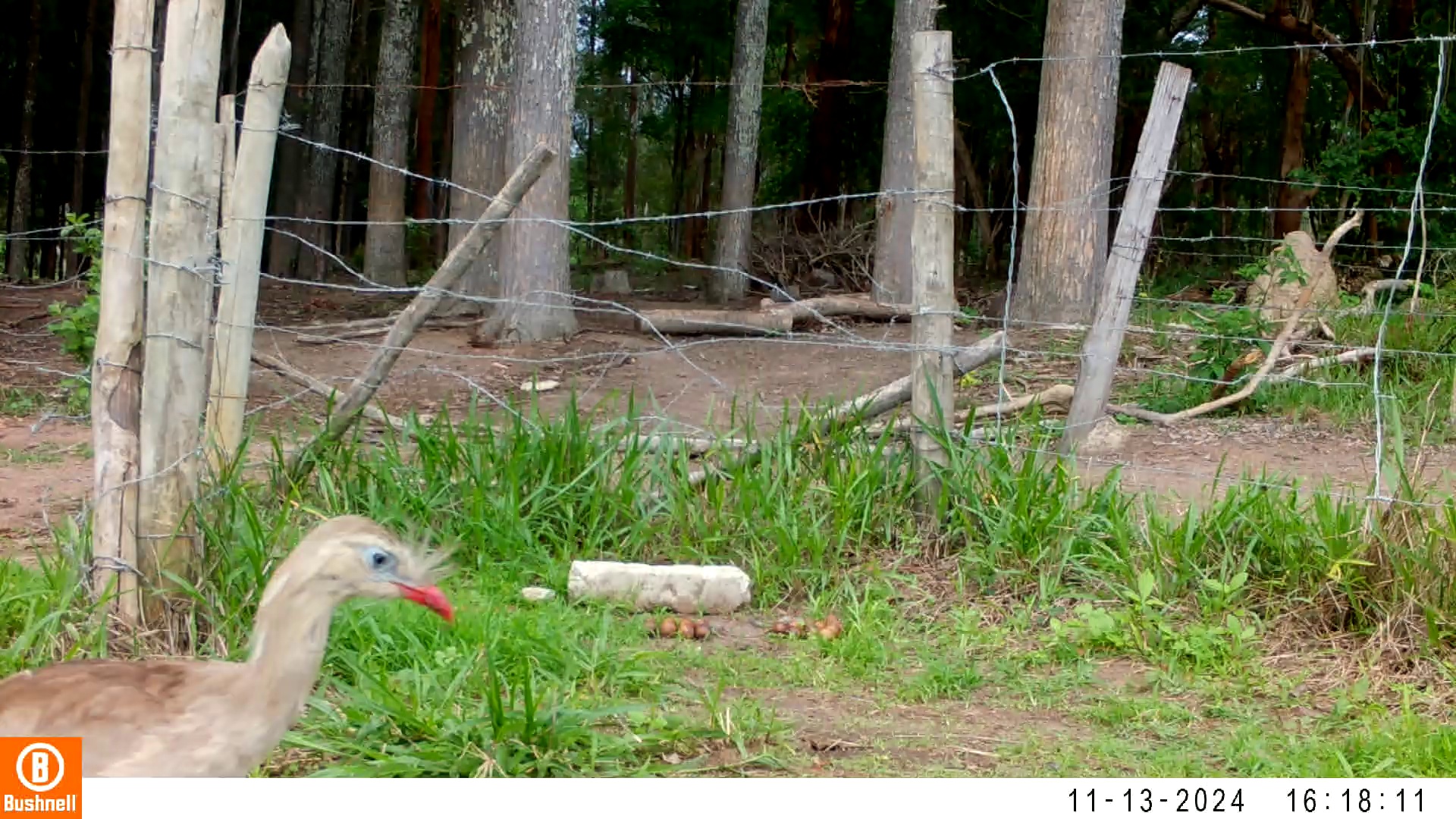

### 11130250-82886-604.jpg

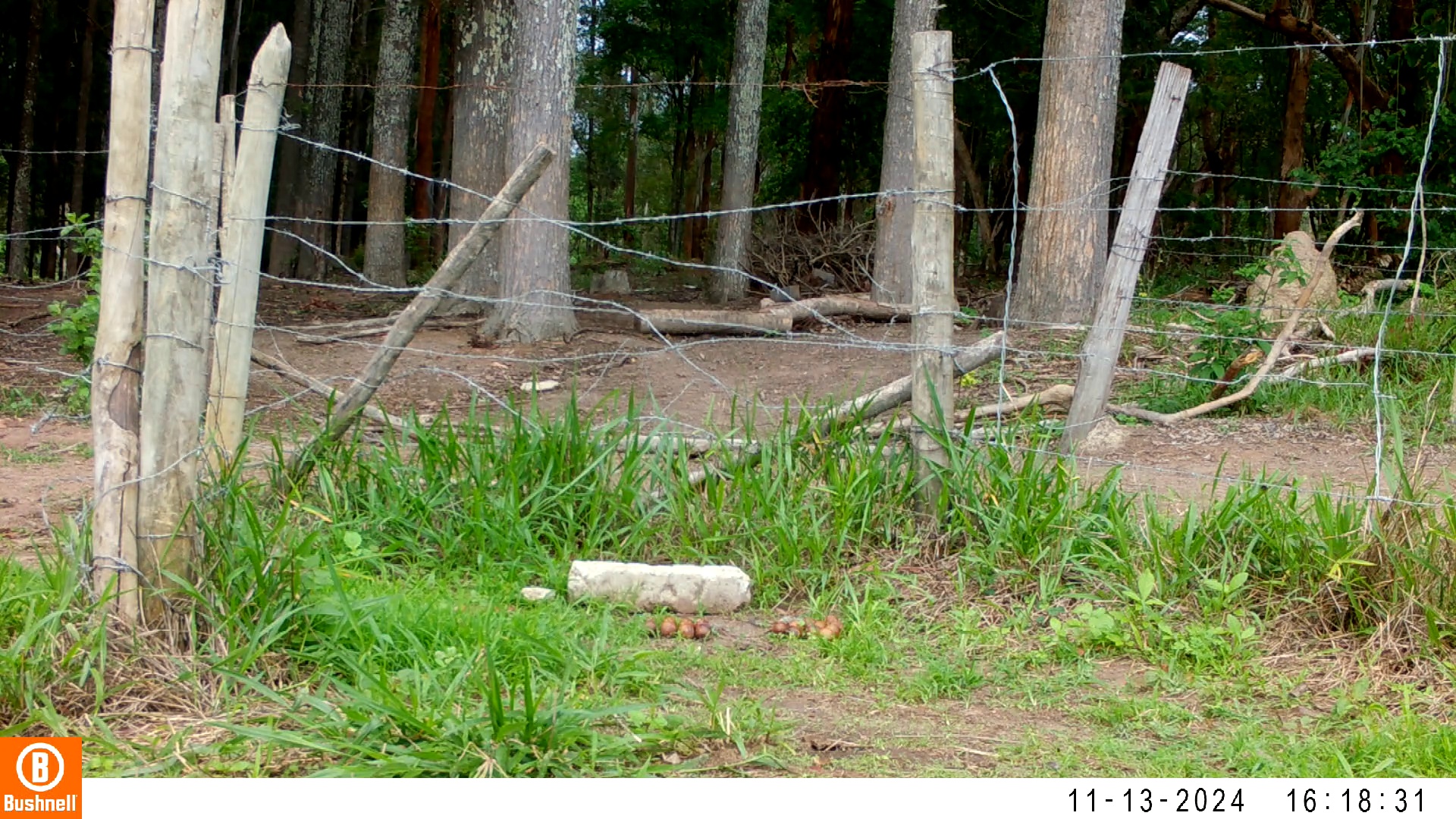

### 11130250-82886-906.jpg

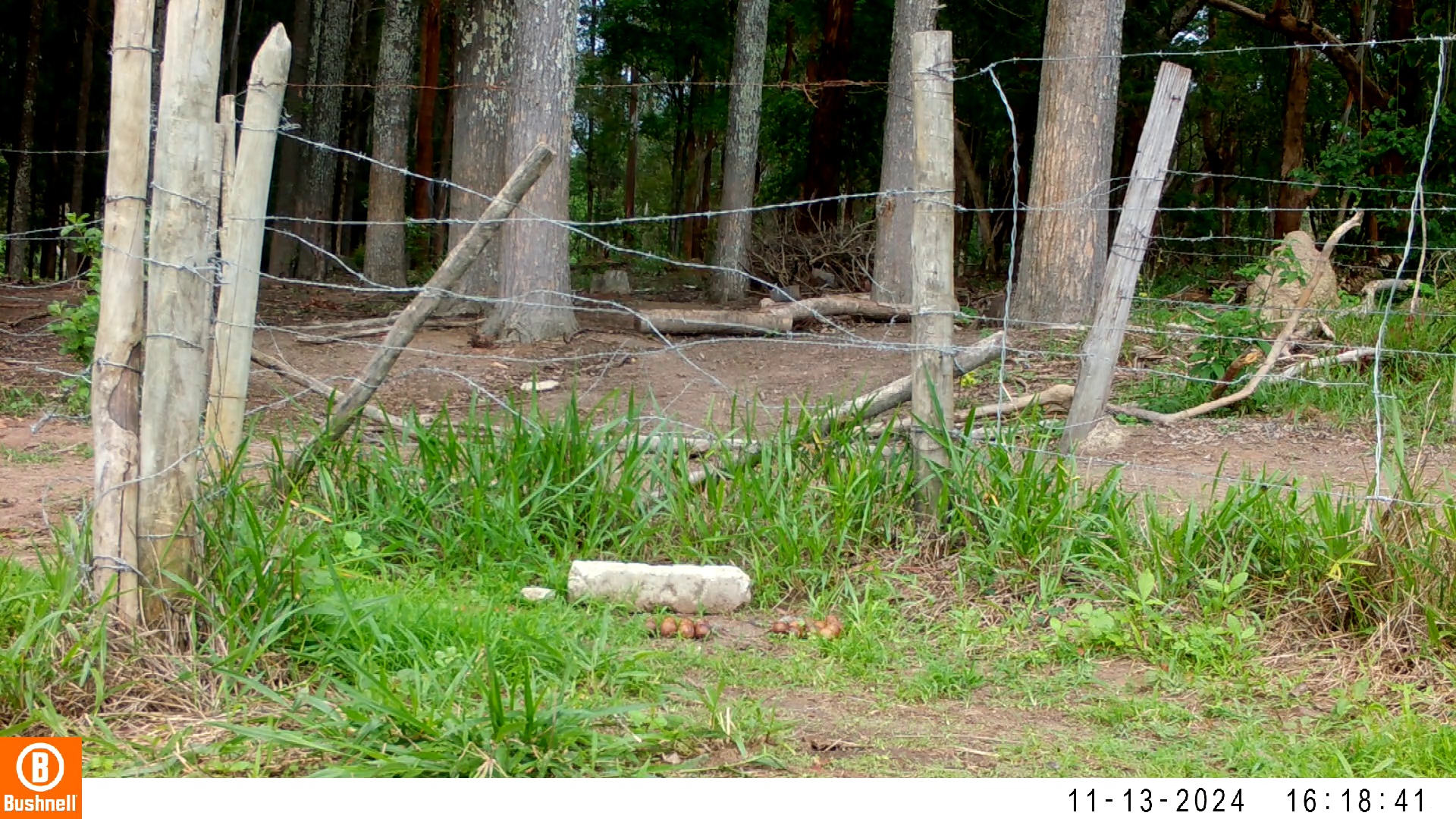

### 11130250-82886-1208.jpg

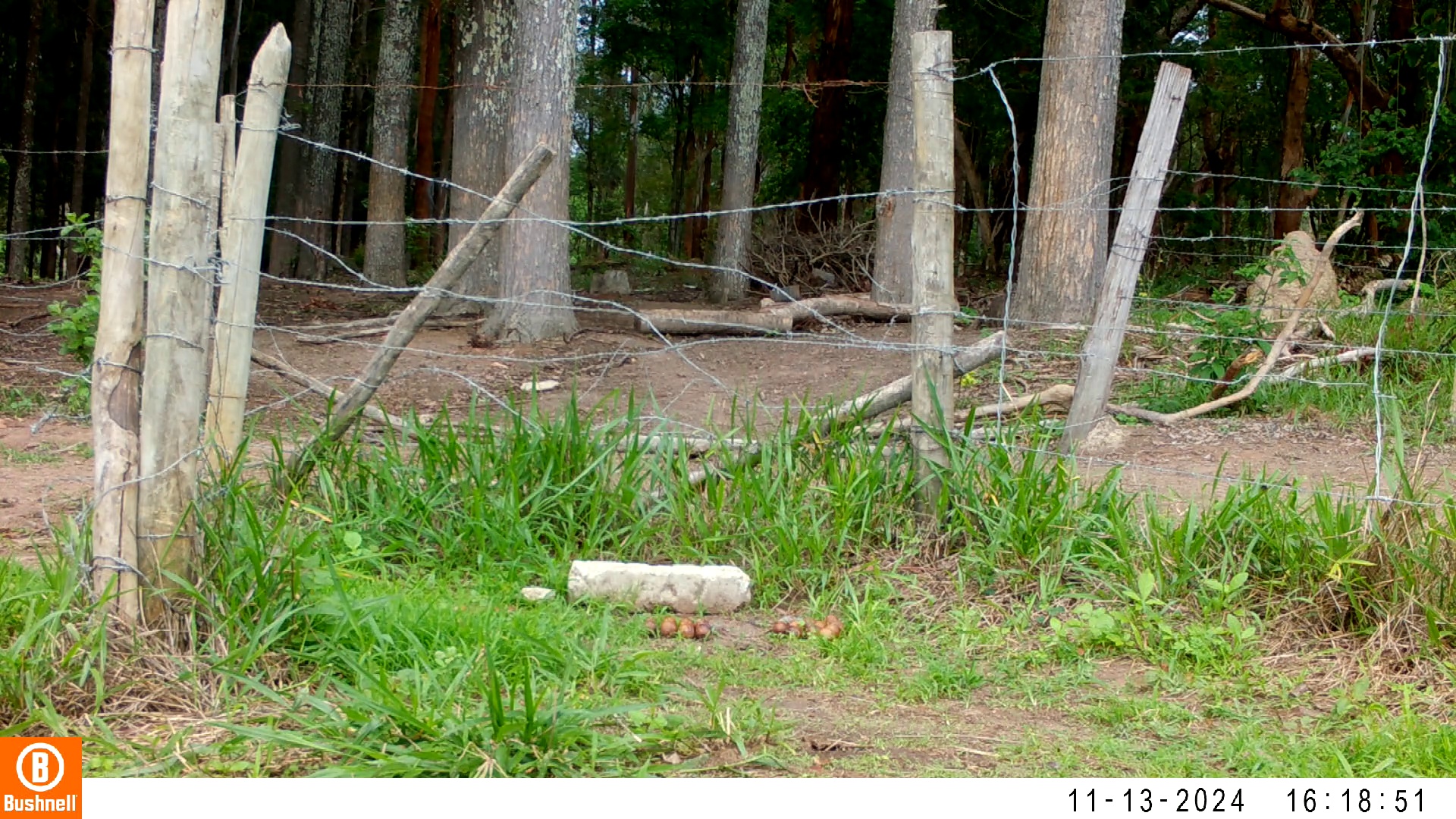

### 11130250-82886-1510.jpg

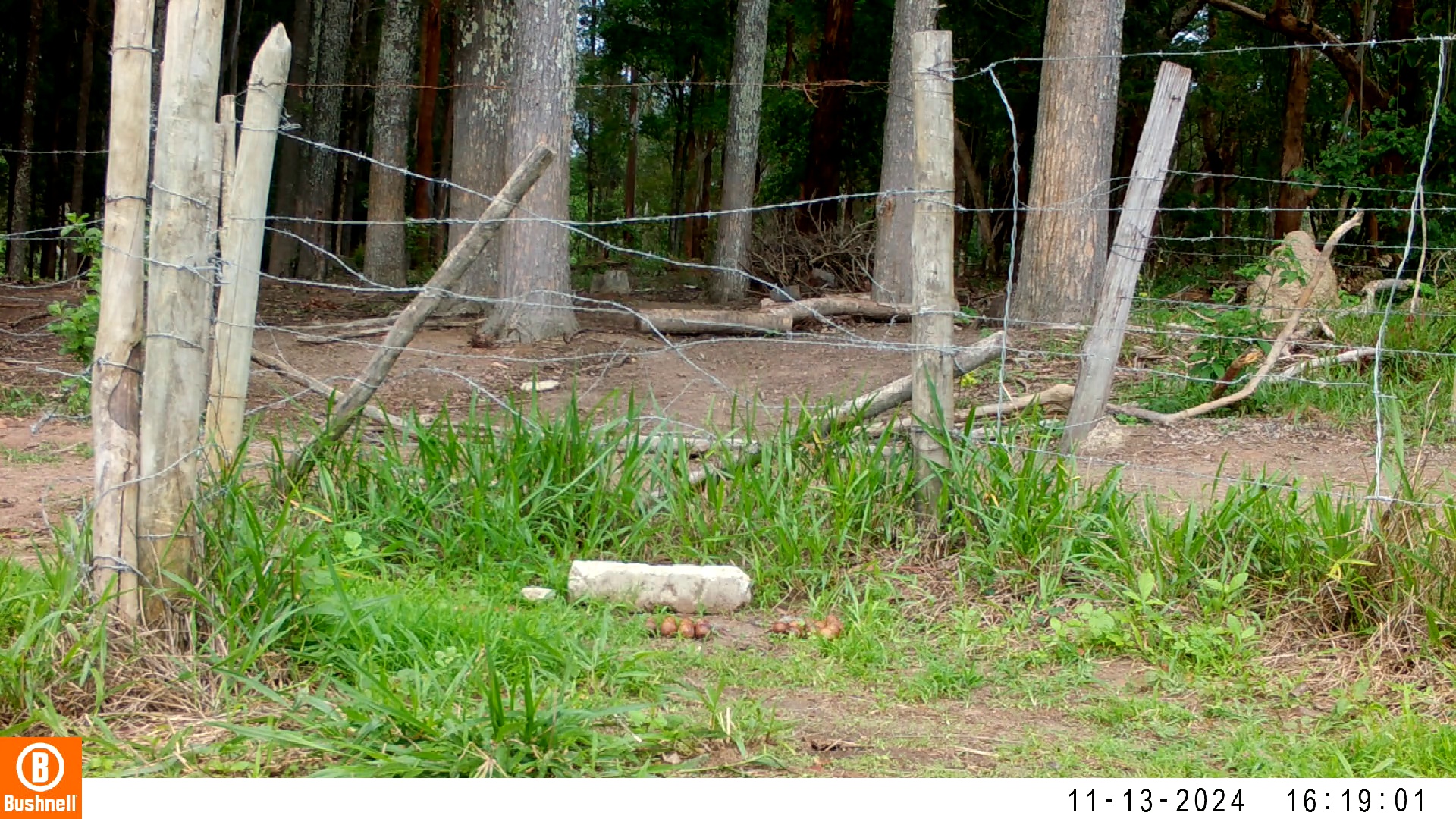

### 11150002-45038-0.jpg

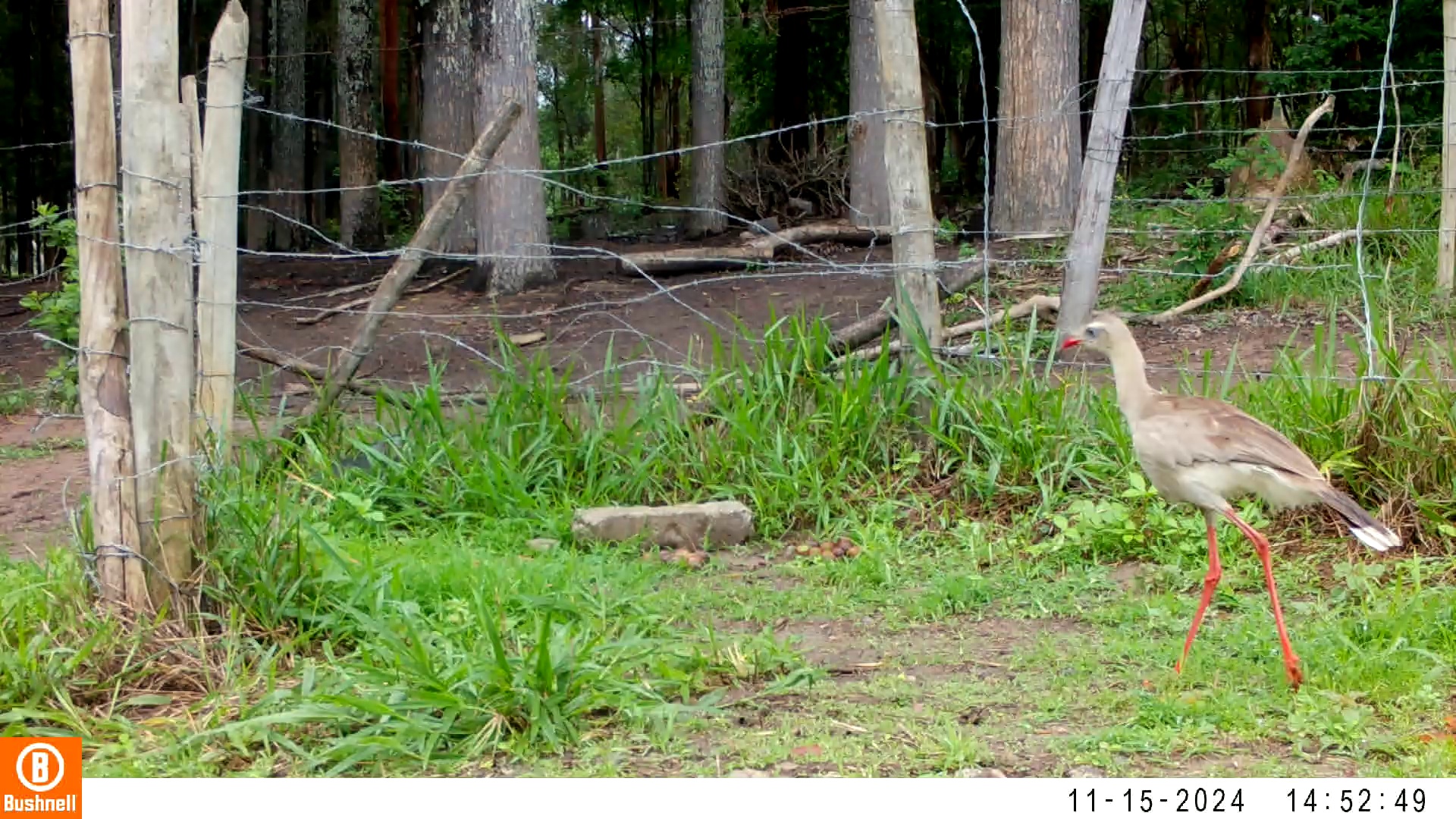

### 11150002-45038-302.jpg

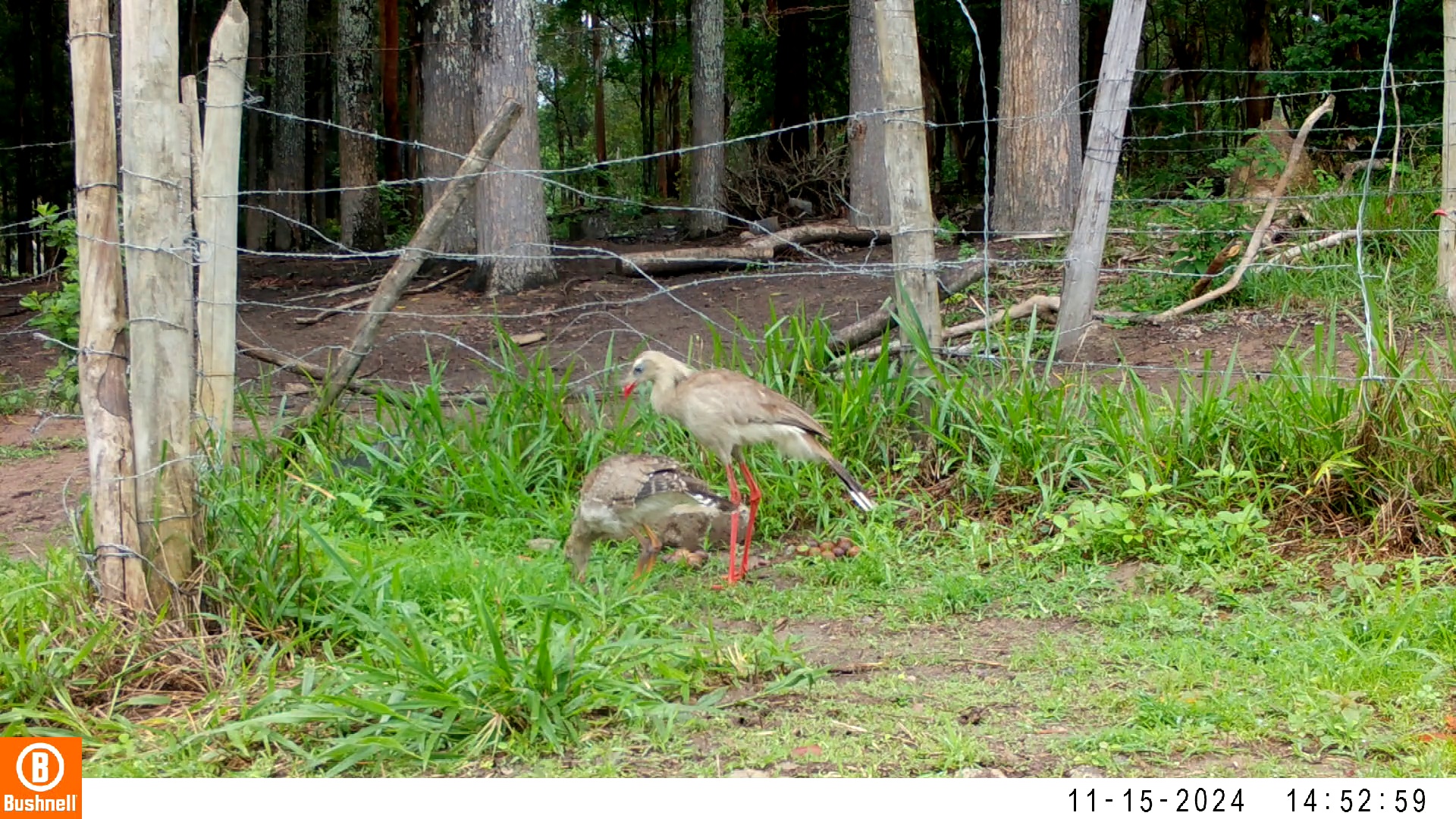

### 11150002-45038-604.jpg

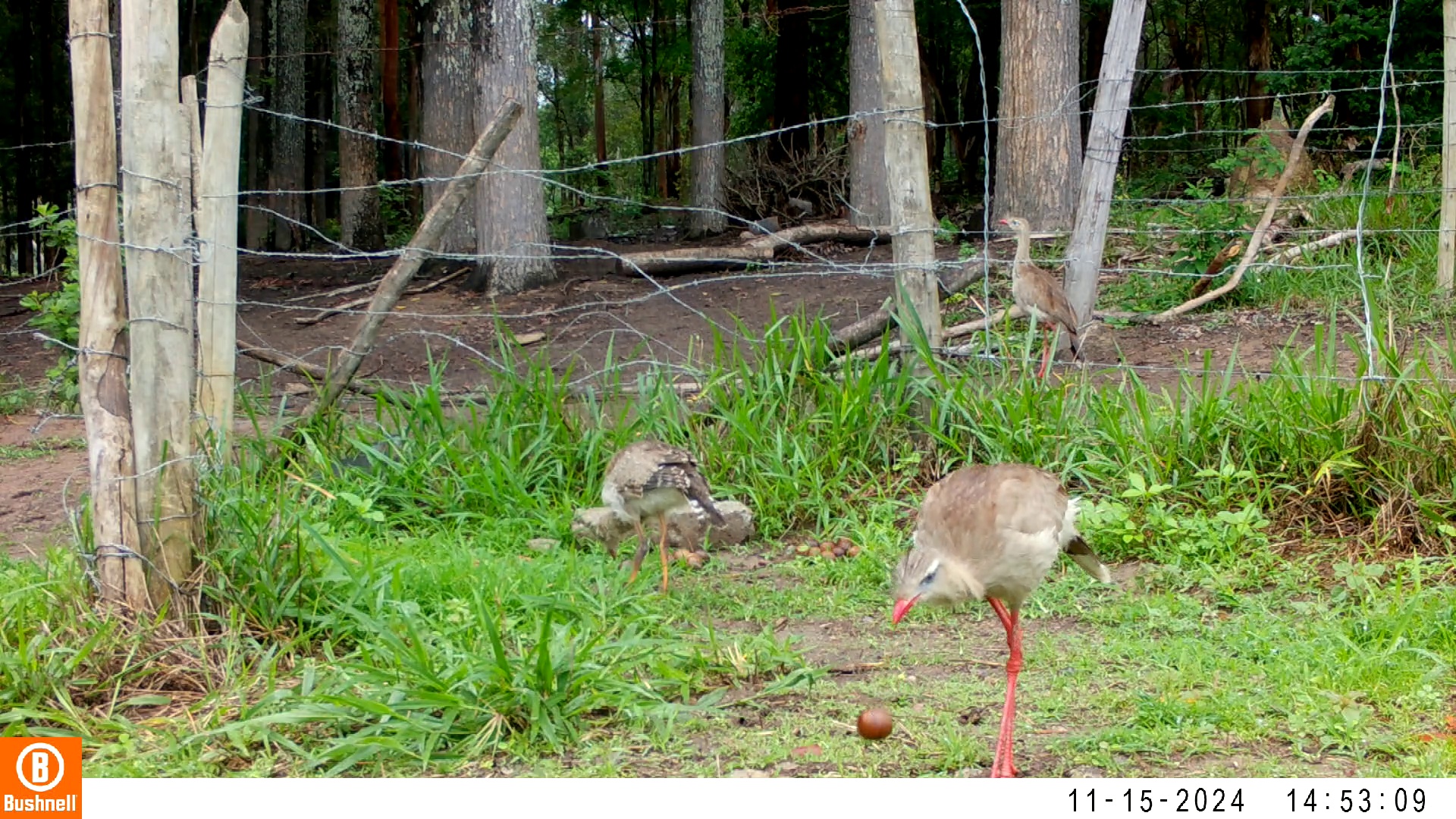

### 11150002-45038-906.jpg

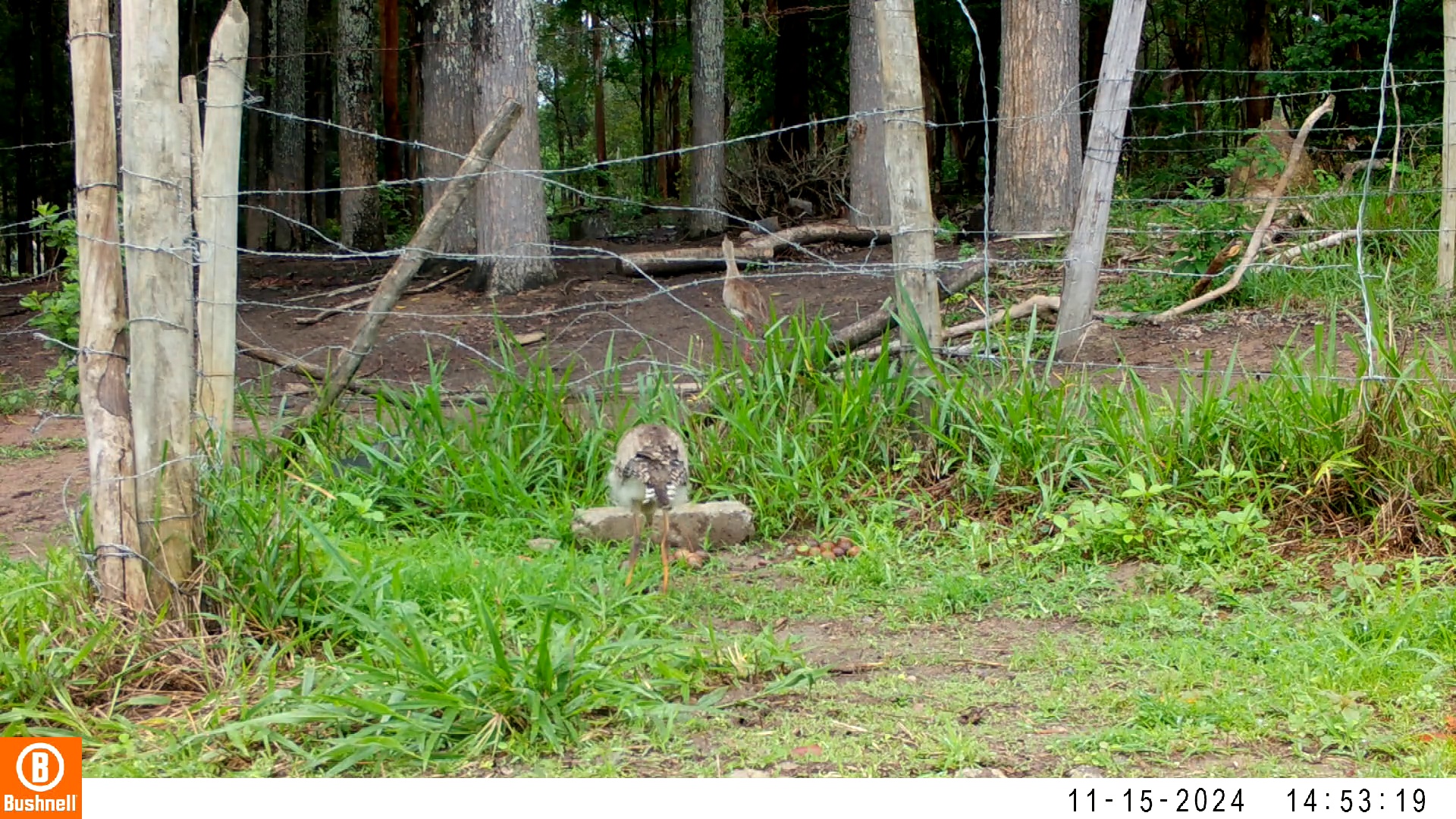

### 11150002-45038-1208.jpg

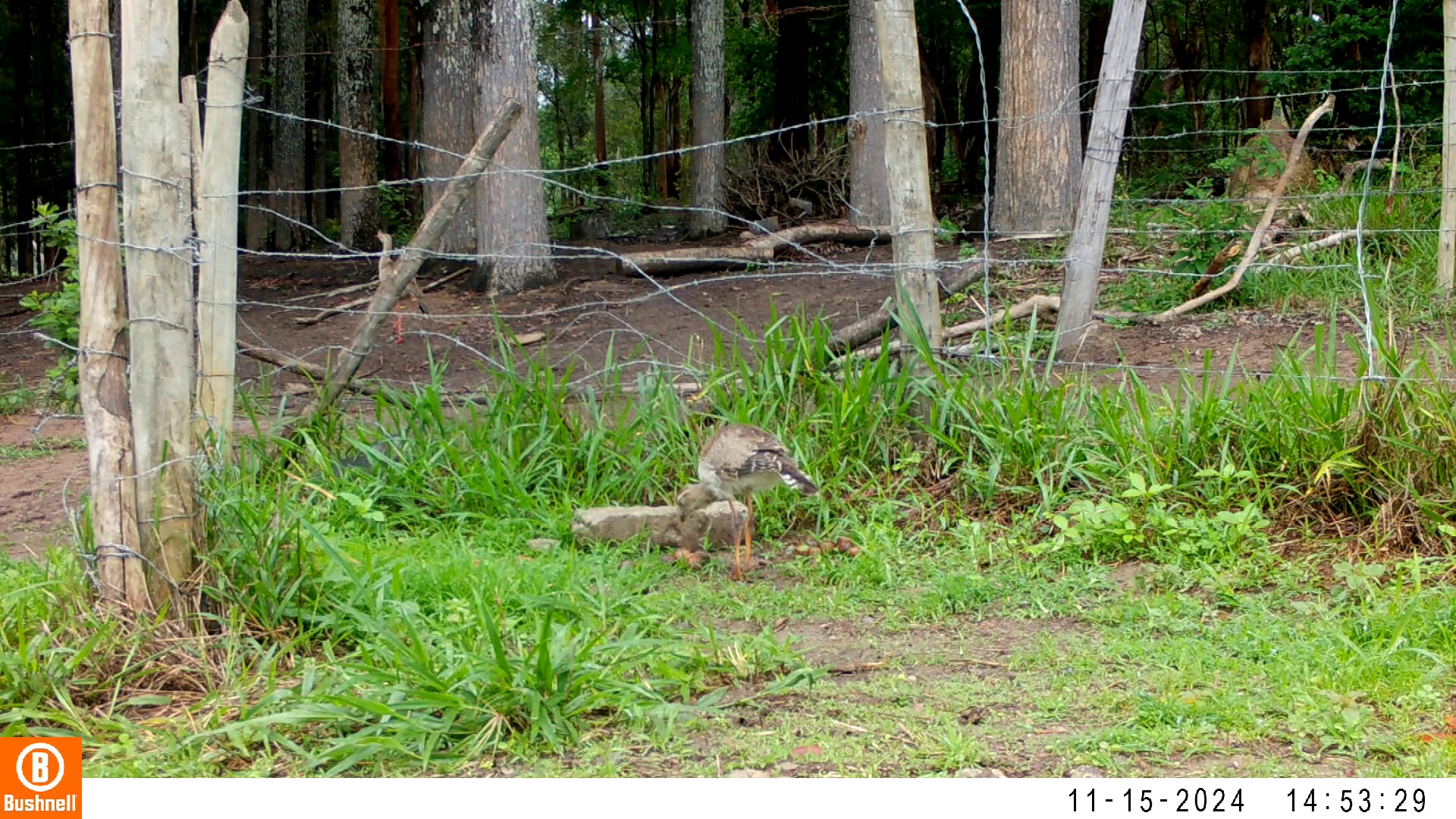

### 11160031-73776-0.jpg

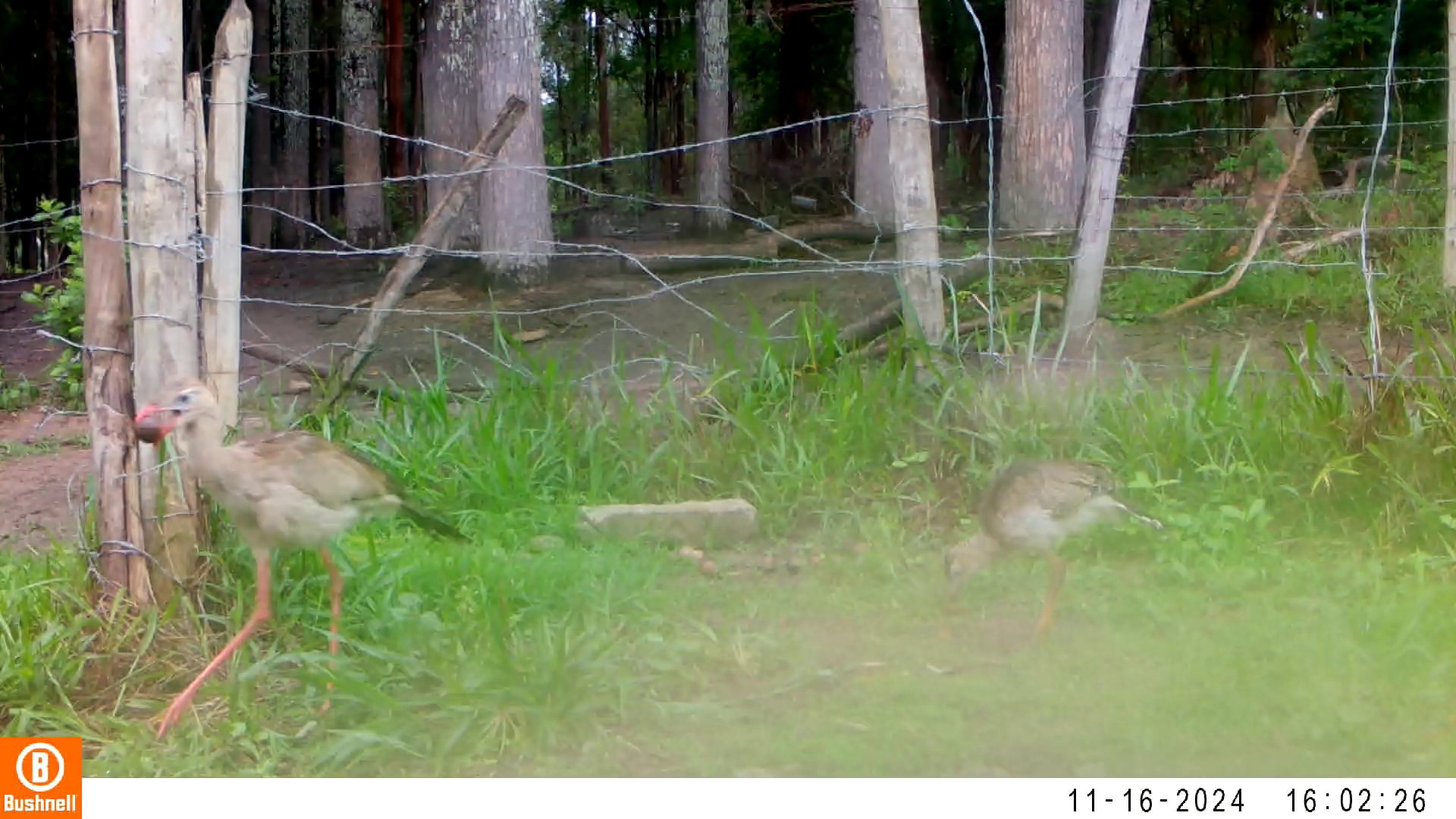

### 11160031-73776-302.jpg

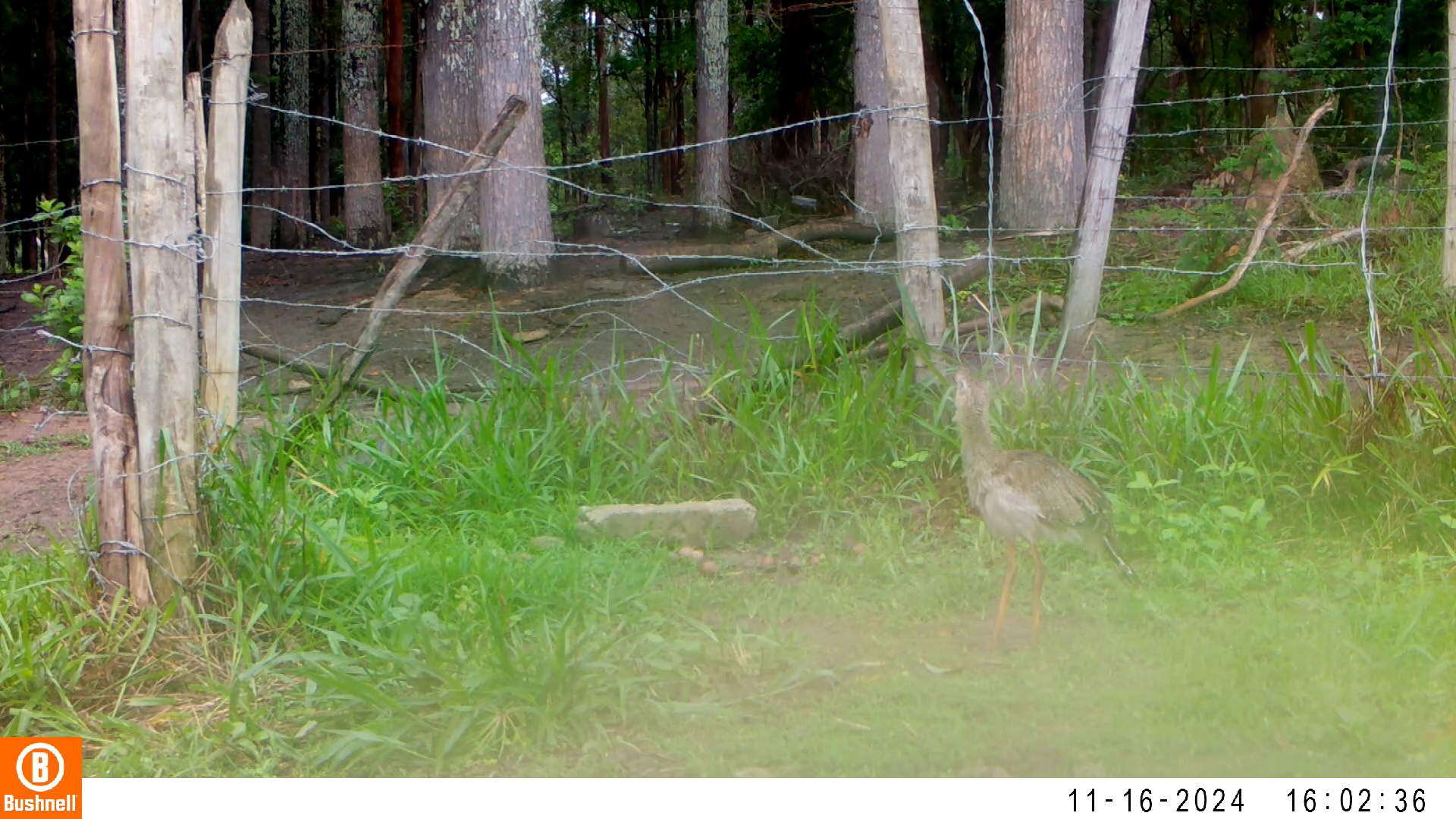

### 11160031-73776-604.jpg

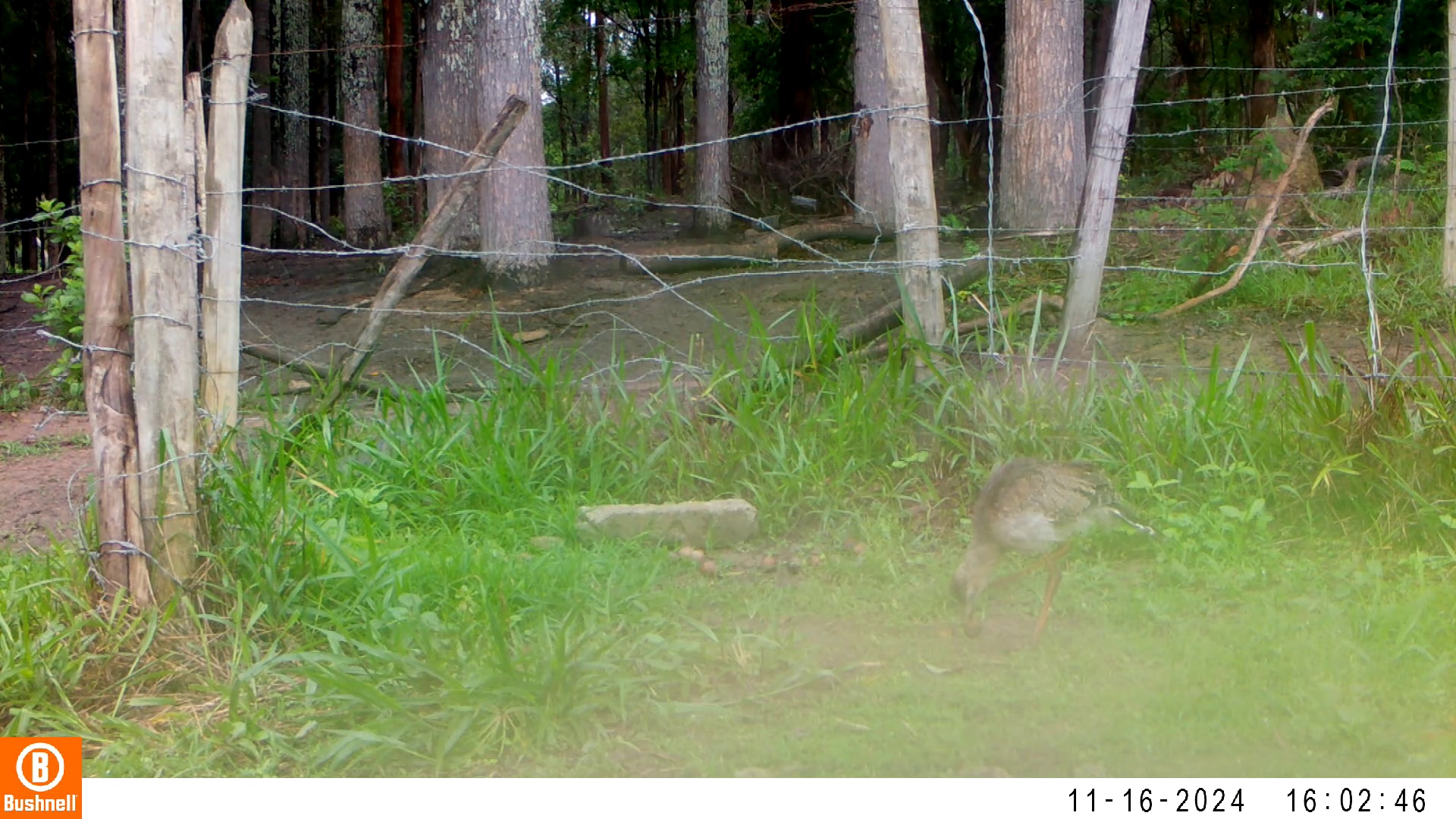

### 11160031-73776-1208.jpg

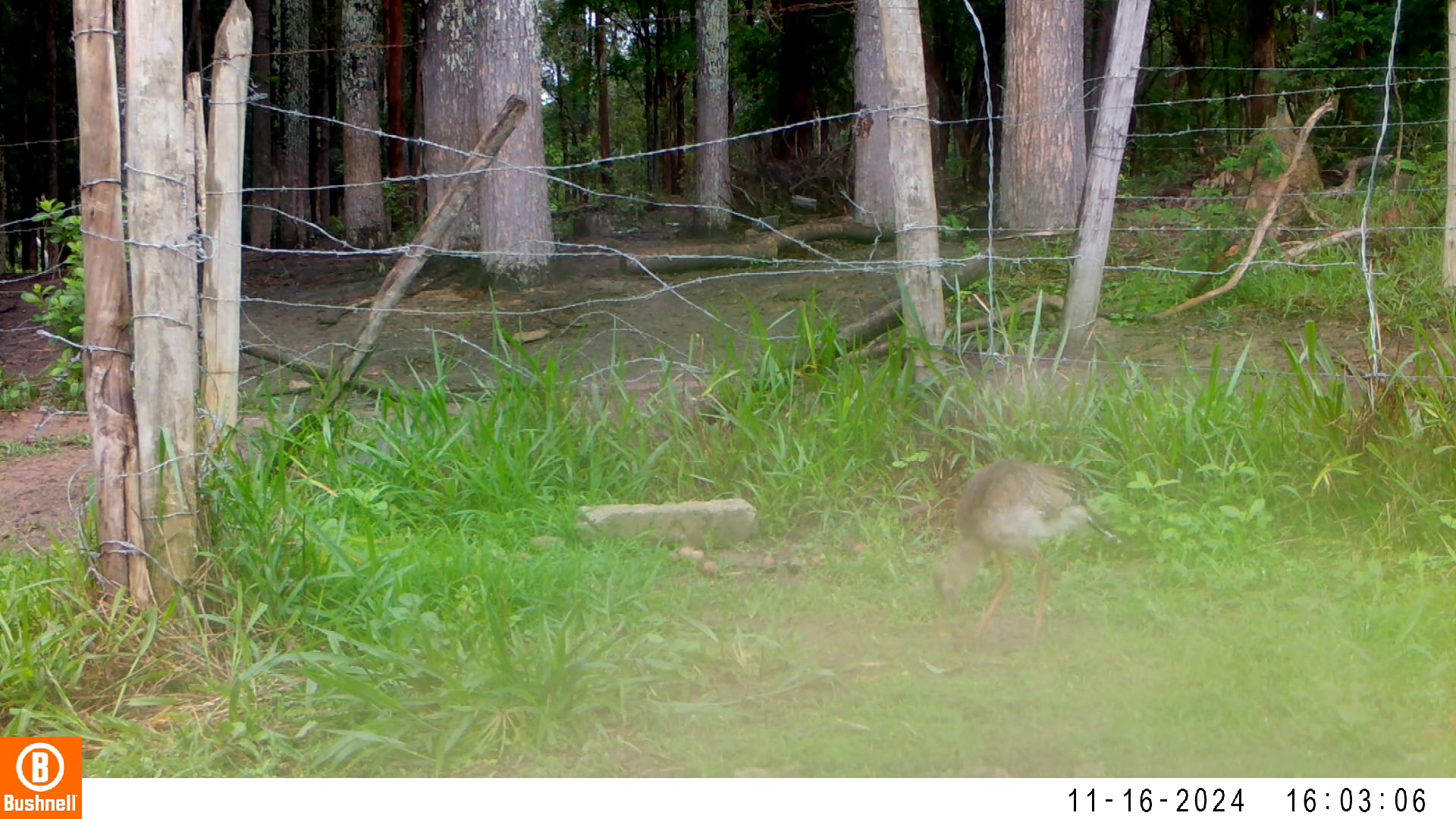

### 11160031-73776-1510.jpg

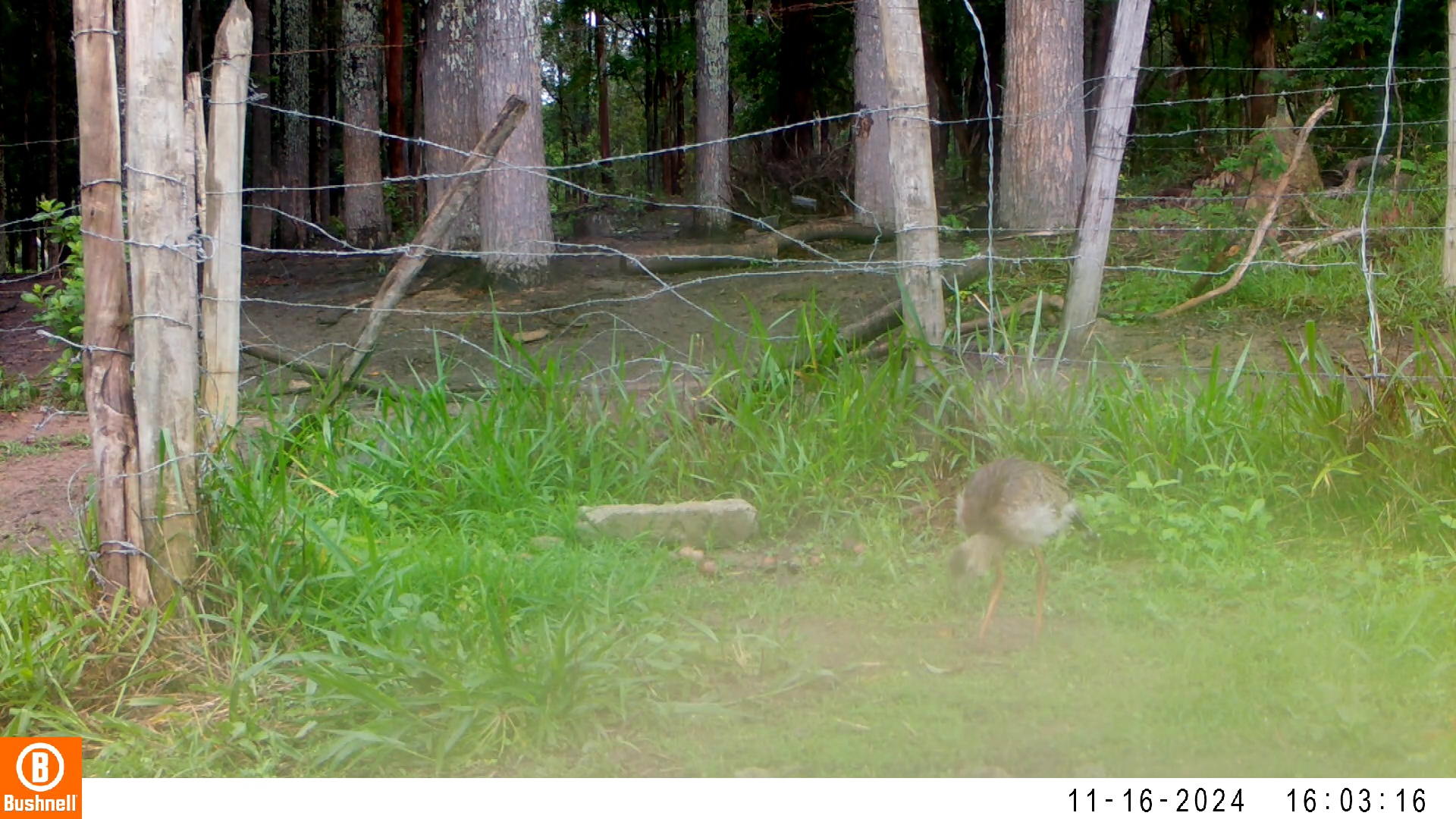

### 11160035-12753-302.jpg

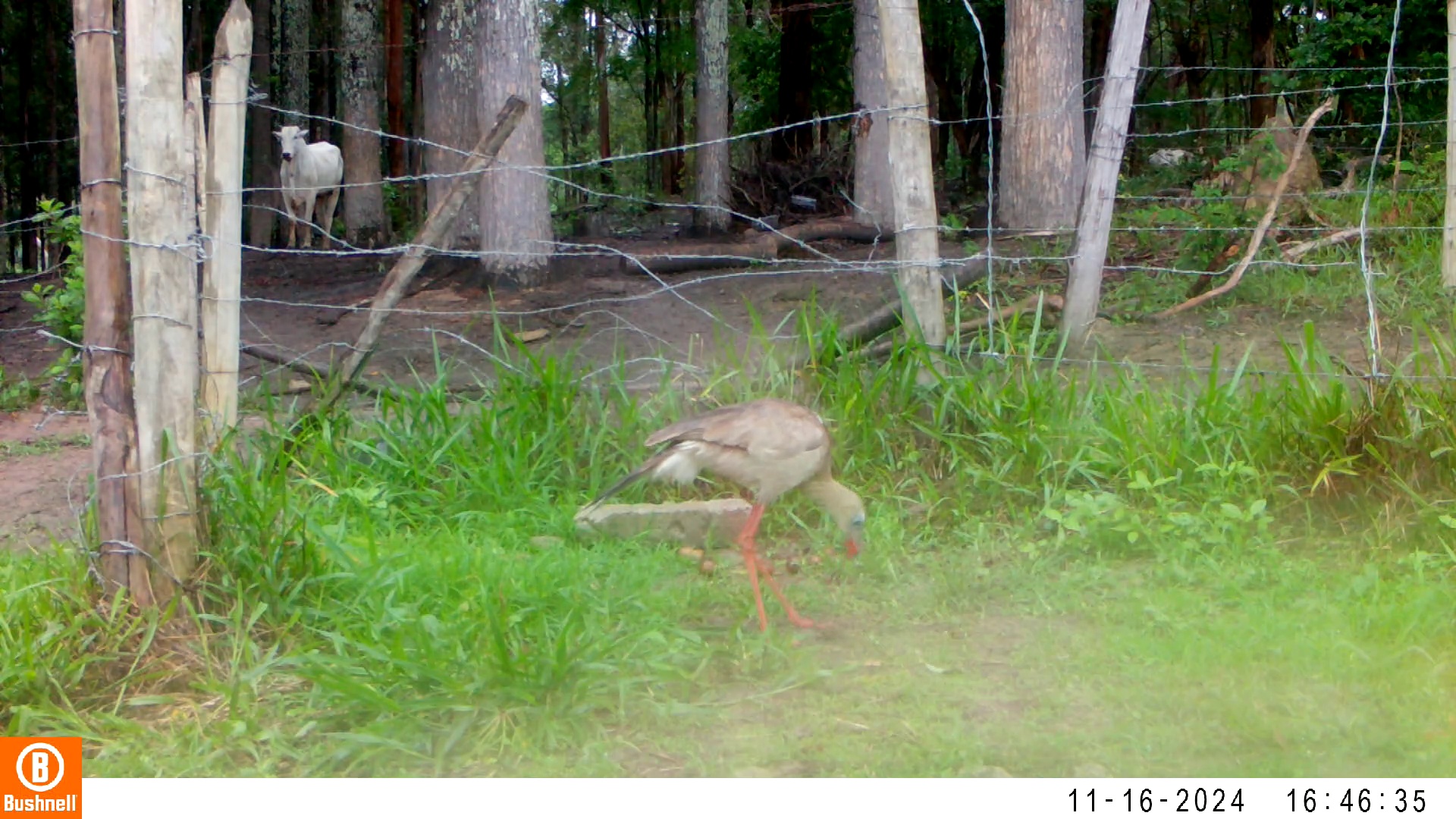

### 11160035-12753-1208.jpg

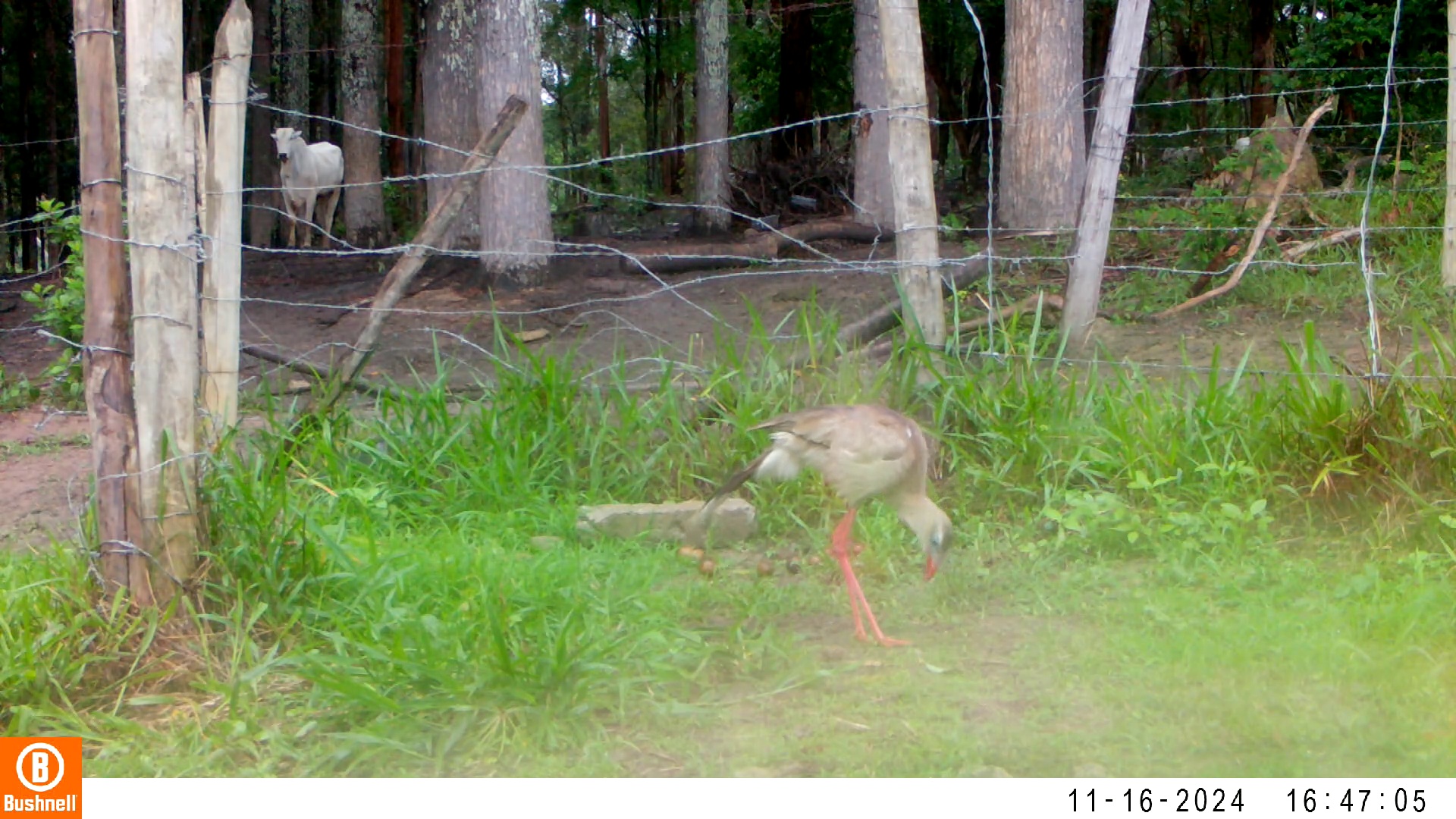

### 11160035-12753-1510.jpg

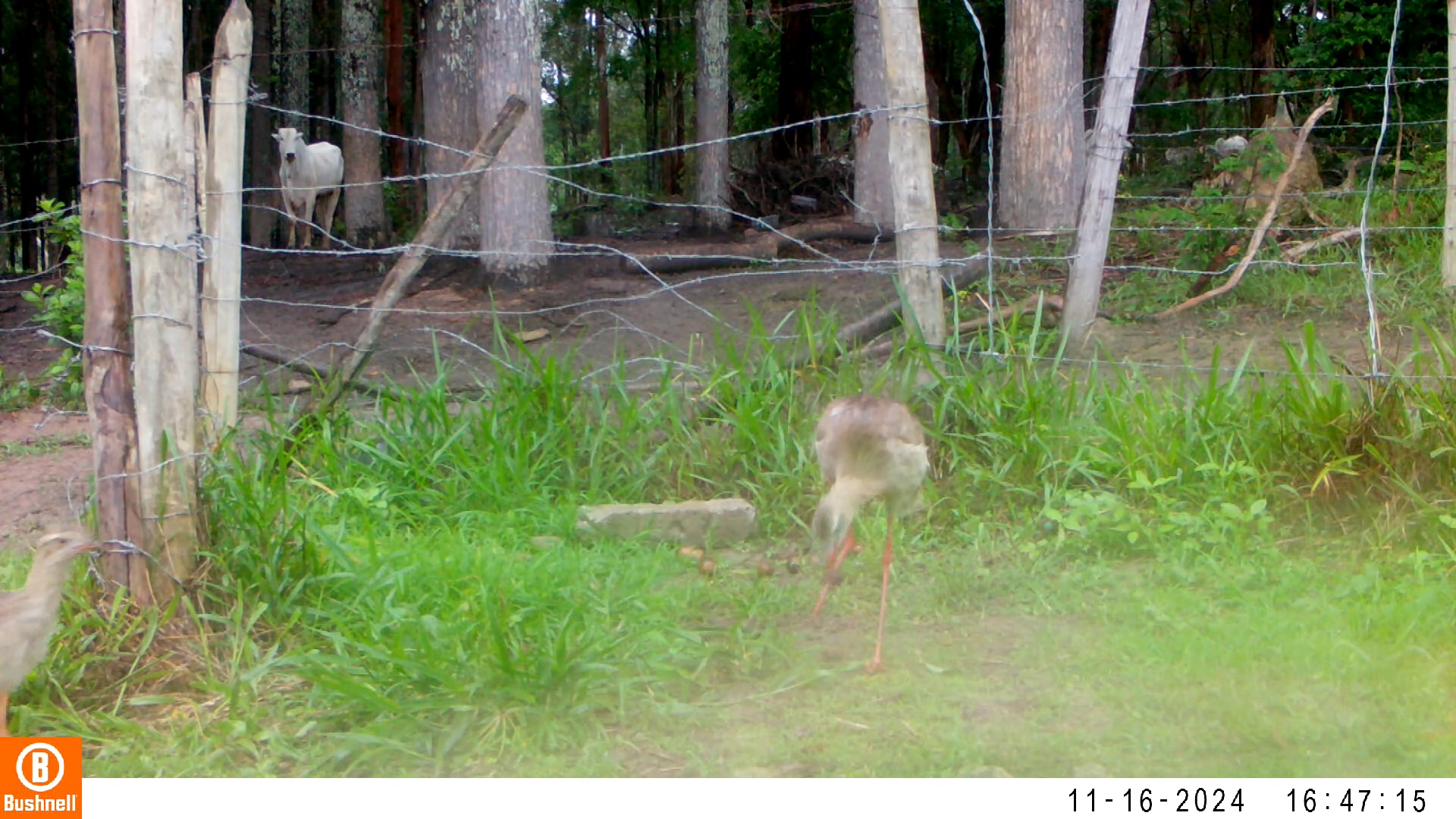

### 11160037-03390-0.jpg

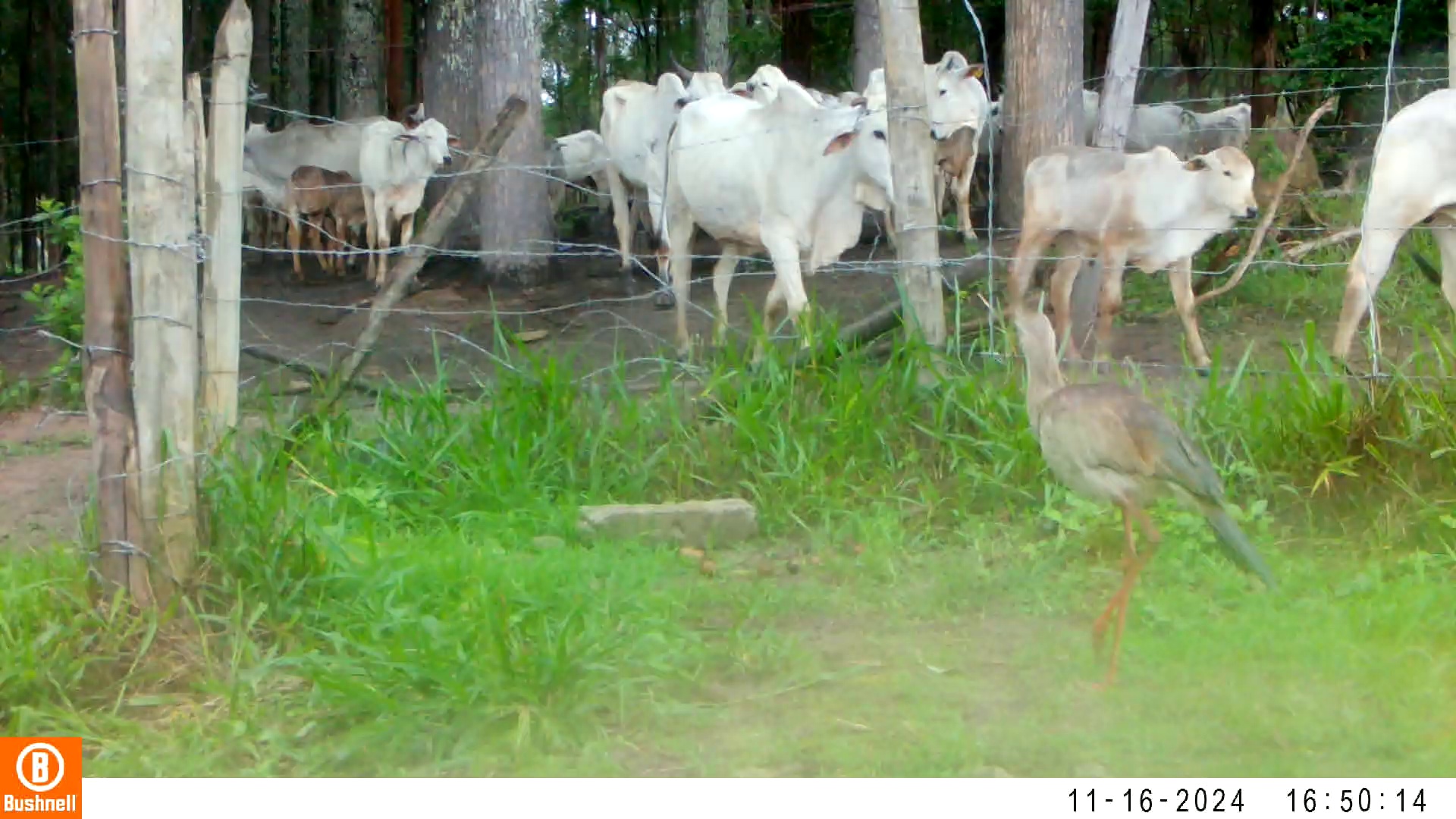

### 11160037-03390-906.jpg

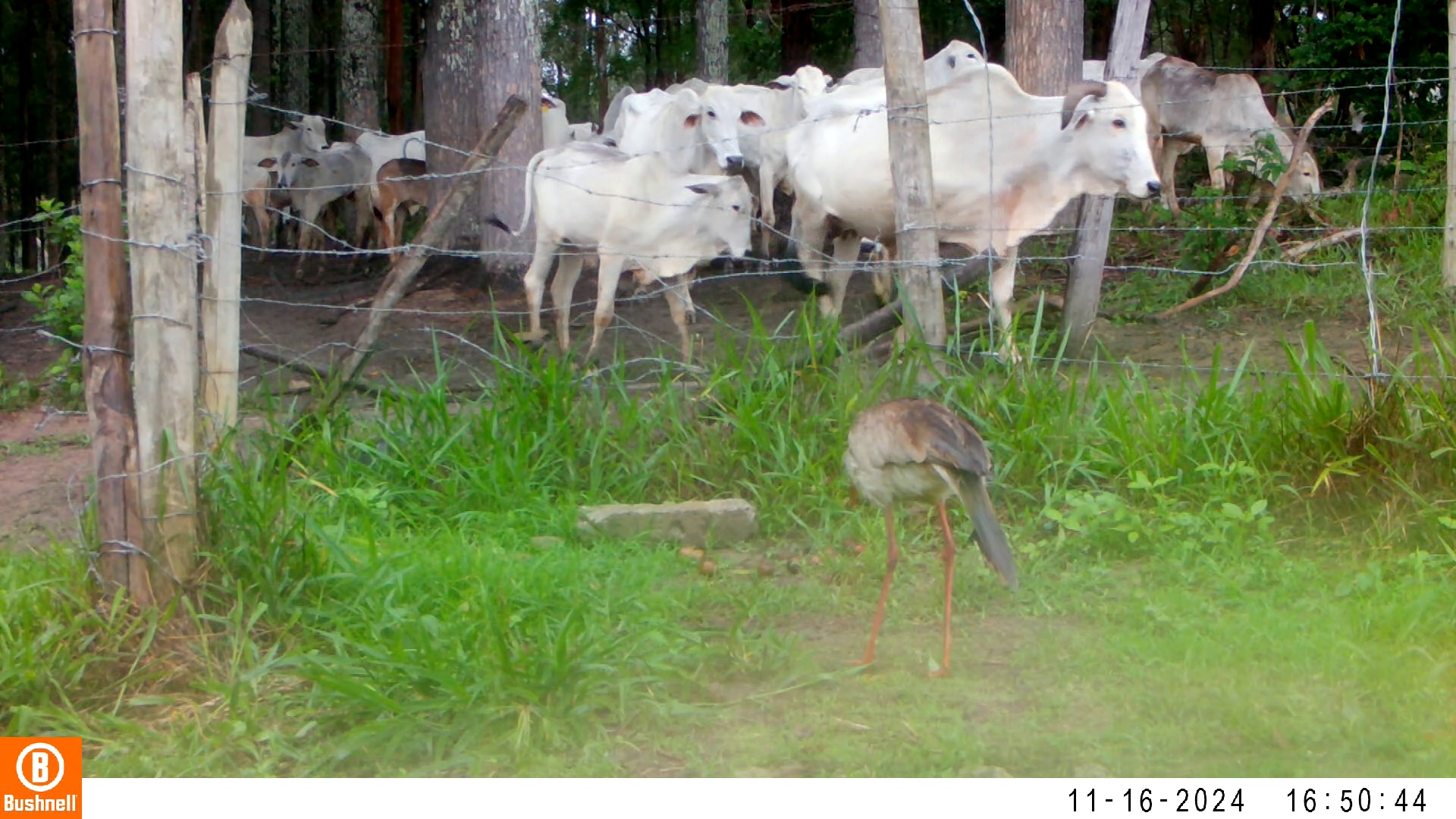

### 11160037-03390-1208.jpg

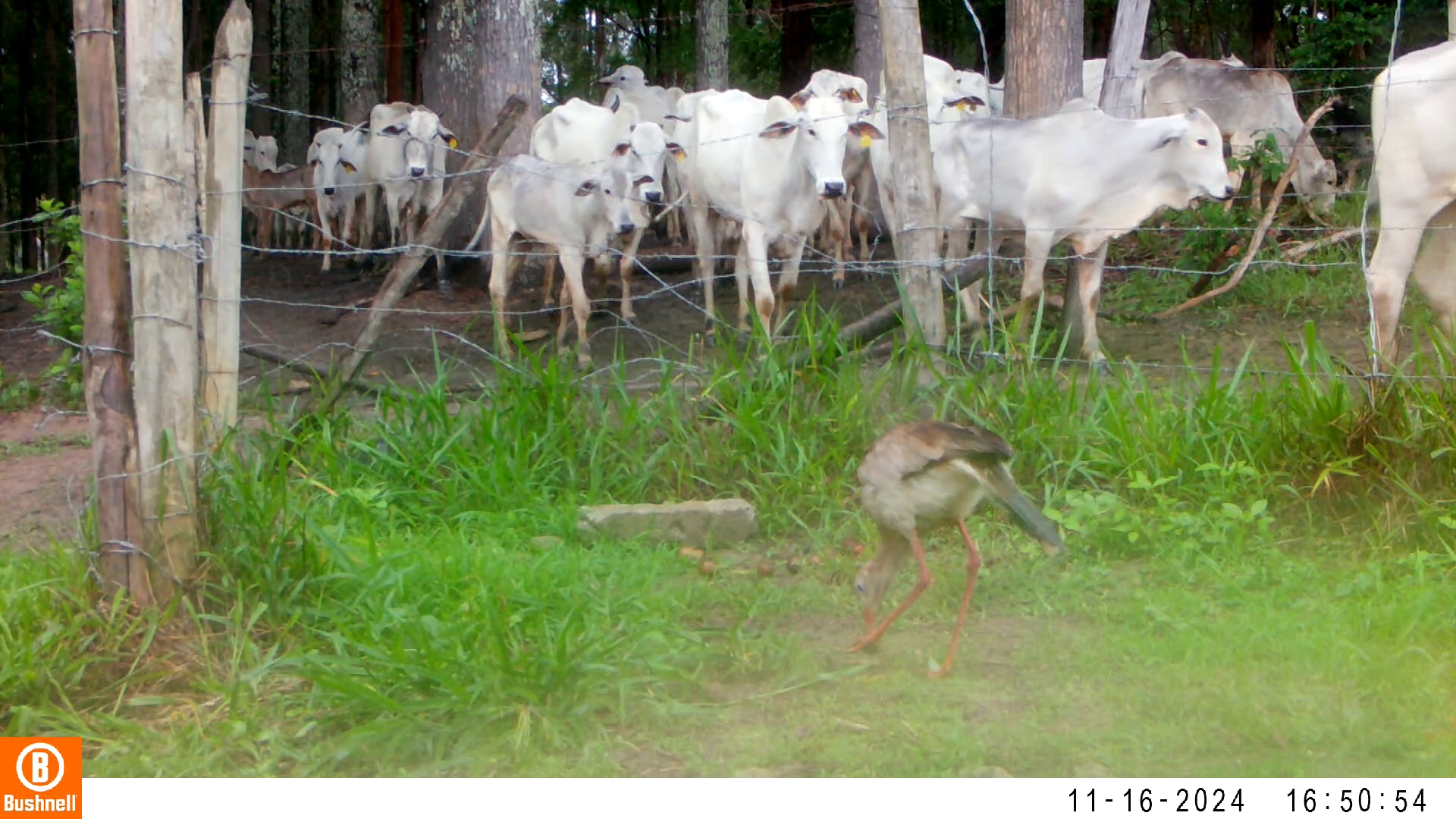

### 11160205-91464-0.jpg

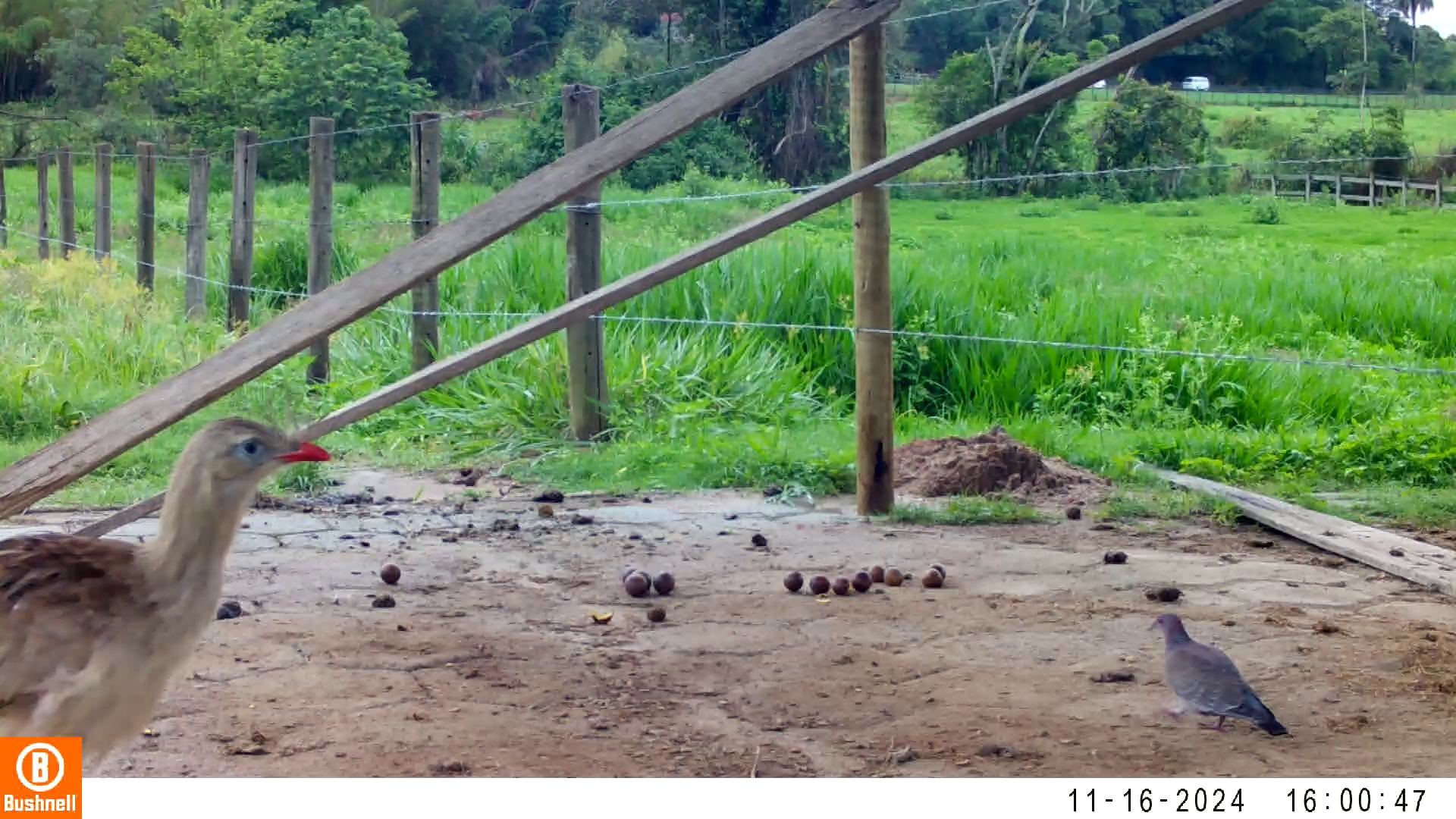

### 11160205-91464-906.jpg

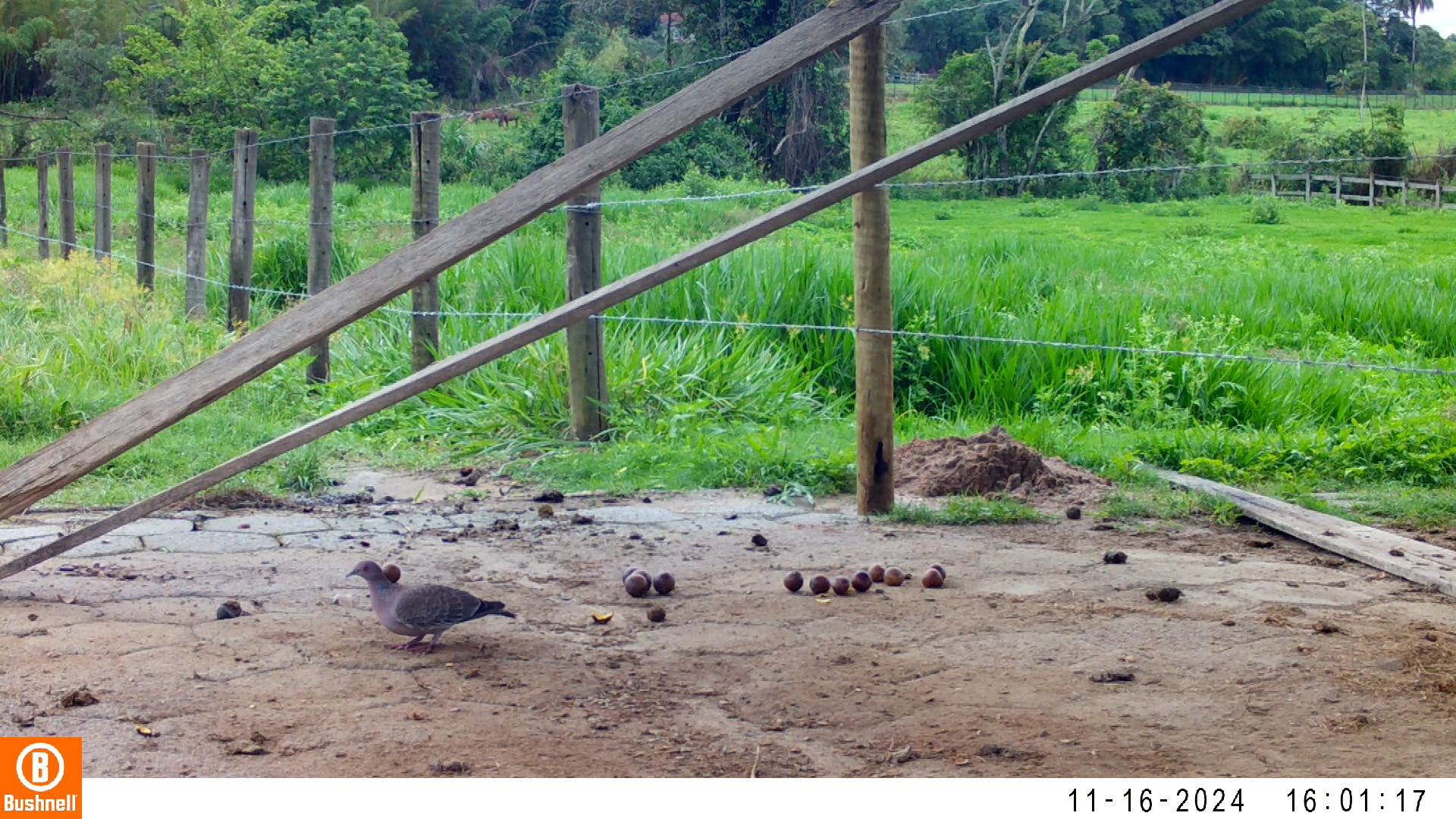

### 11170083-14255-0.jpg

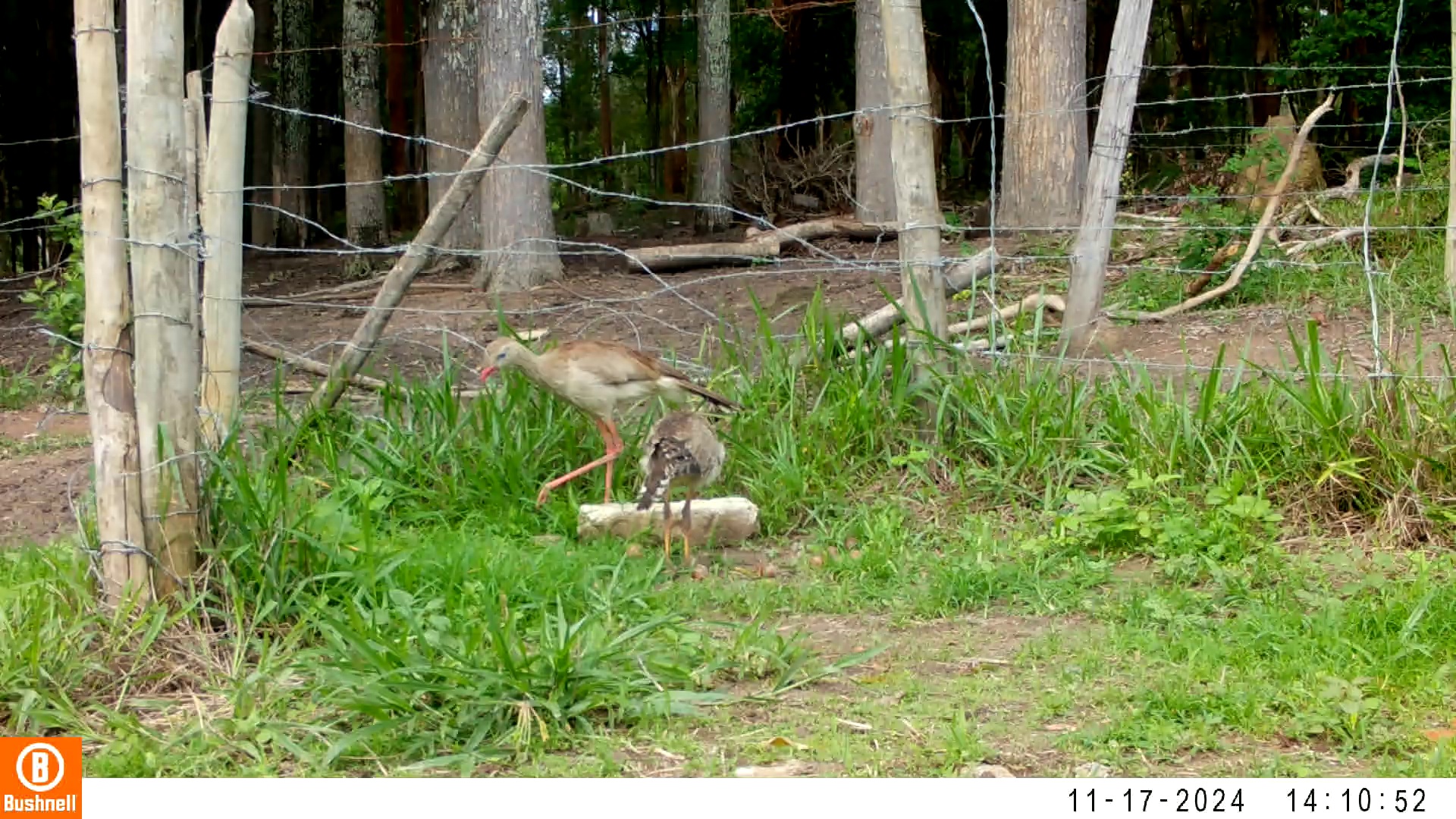

### 11170083-14255-302.jpg

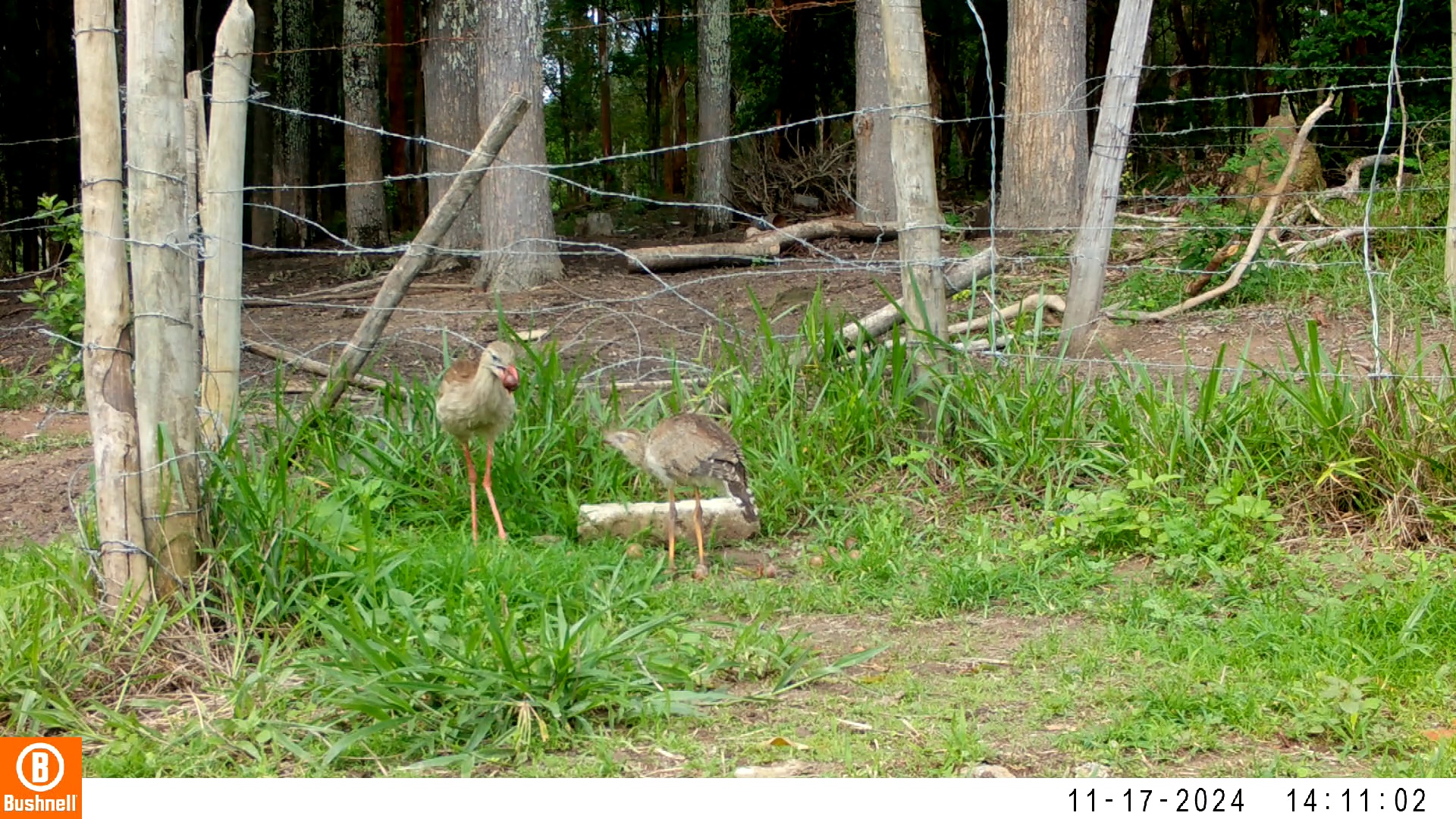

### 11170083-14255-604.jpg

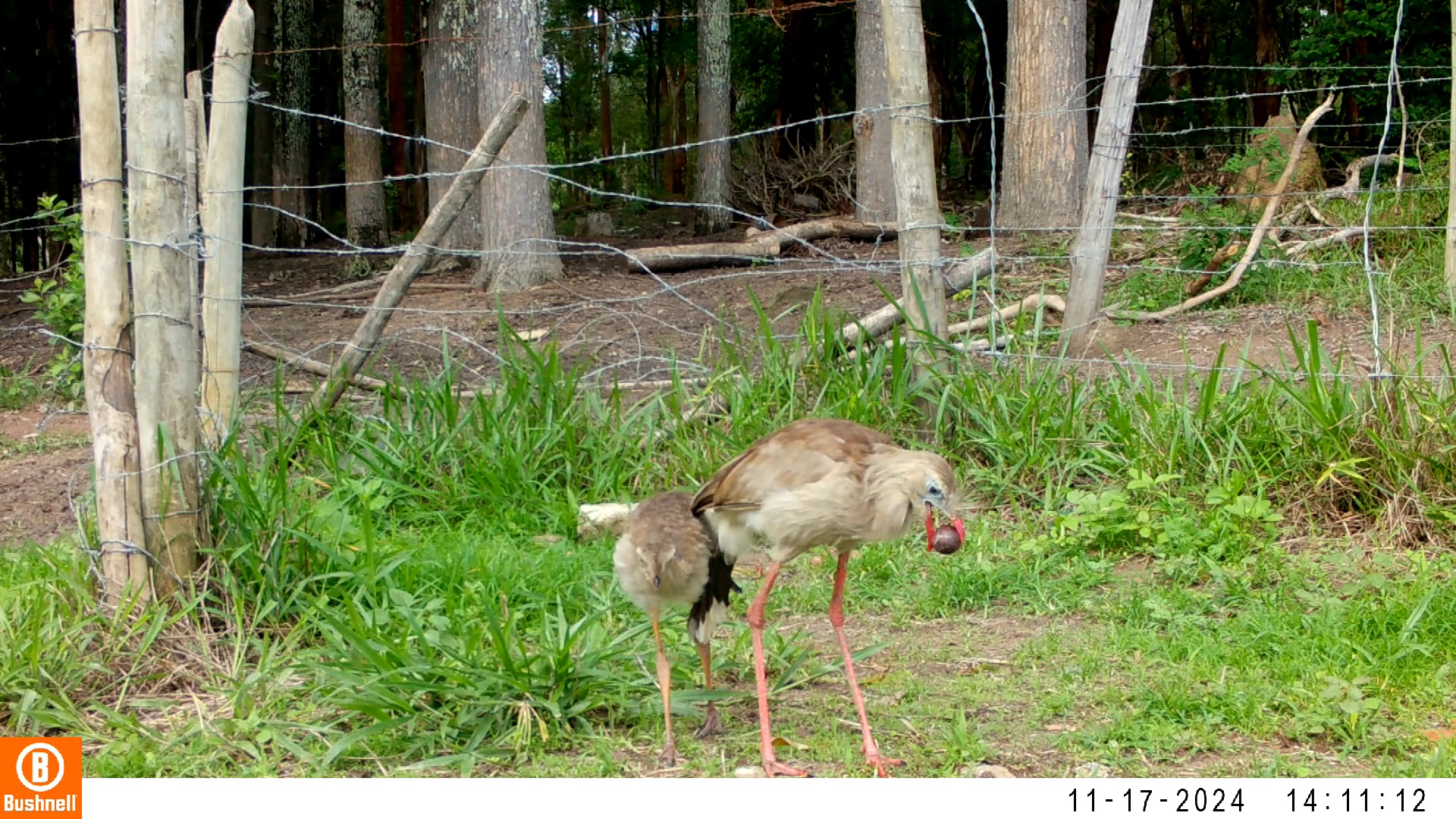

### 11170083-14255-906.jpg

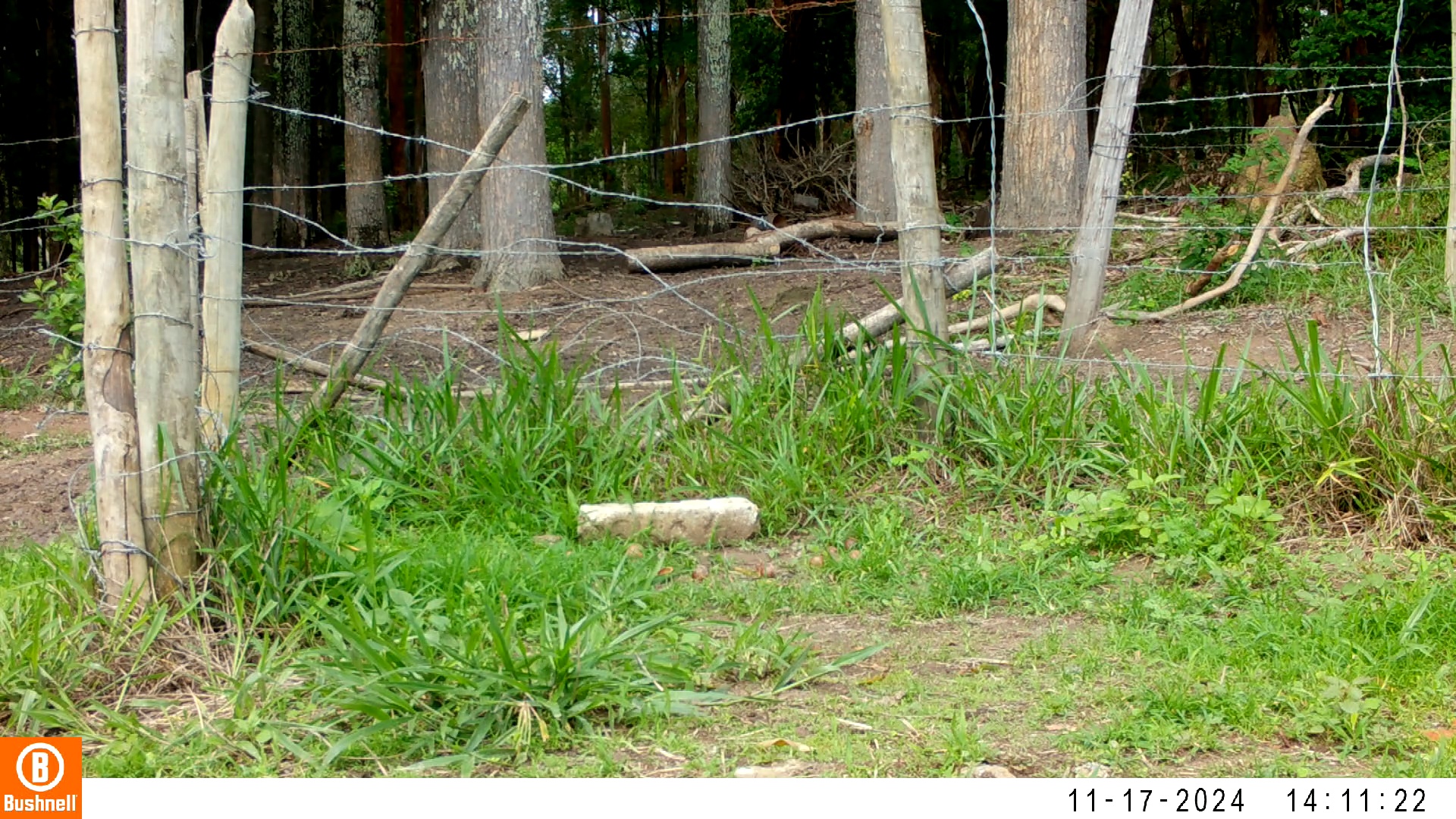

### 11170083-14255-1208.jpg

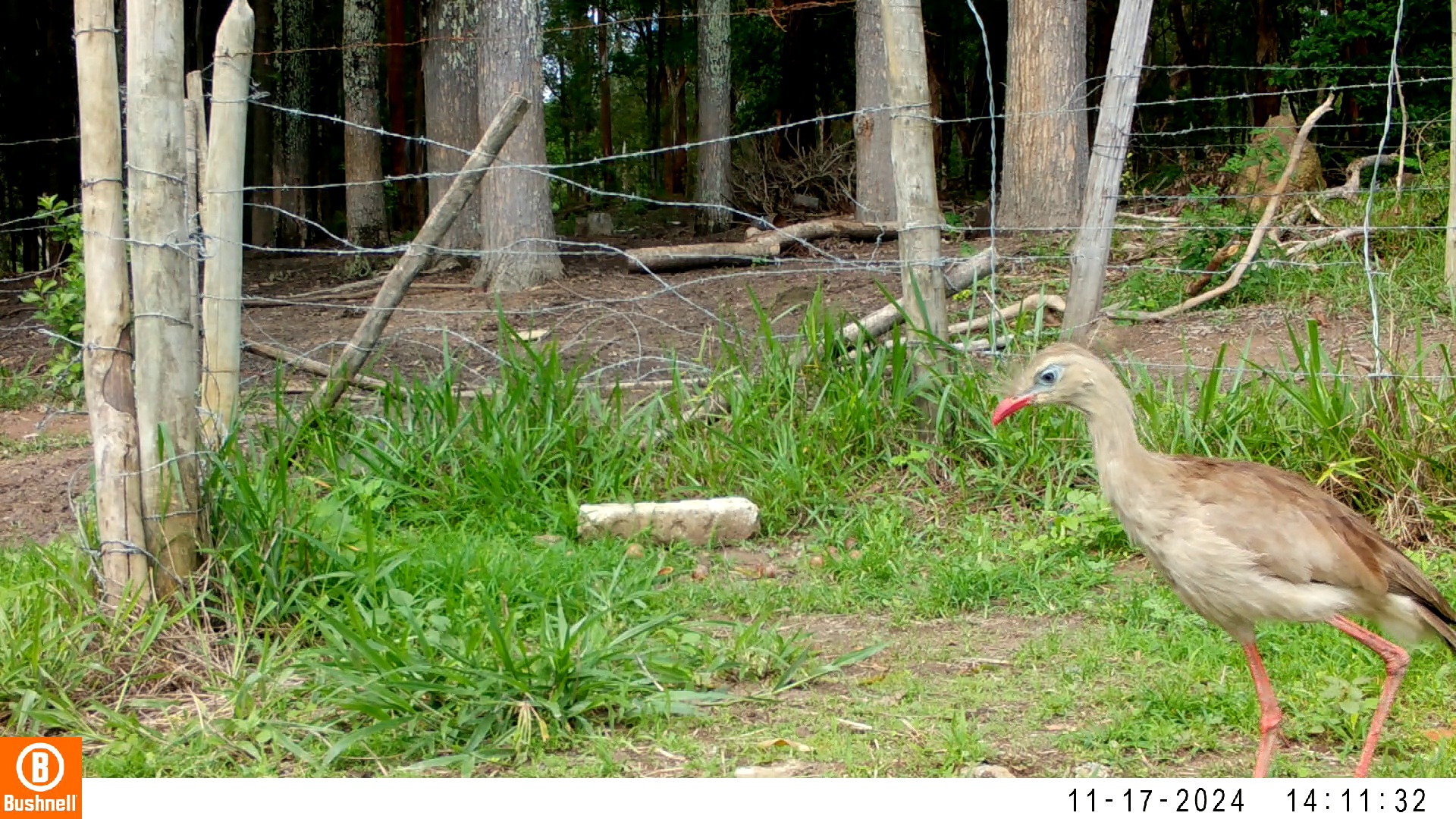

### 11170083-14255-1510.jpg

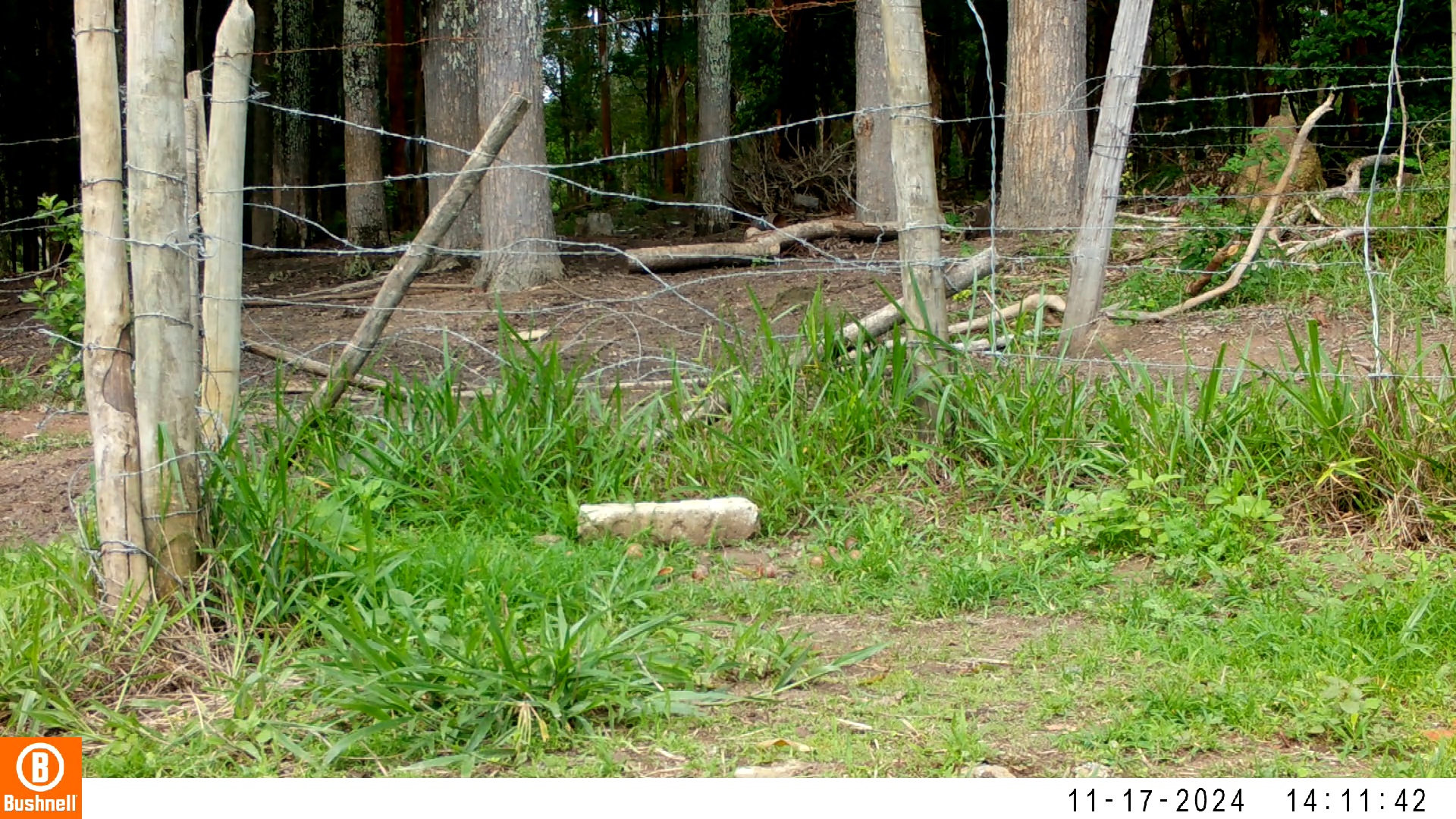

### 11170086-33174-302.jpg

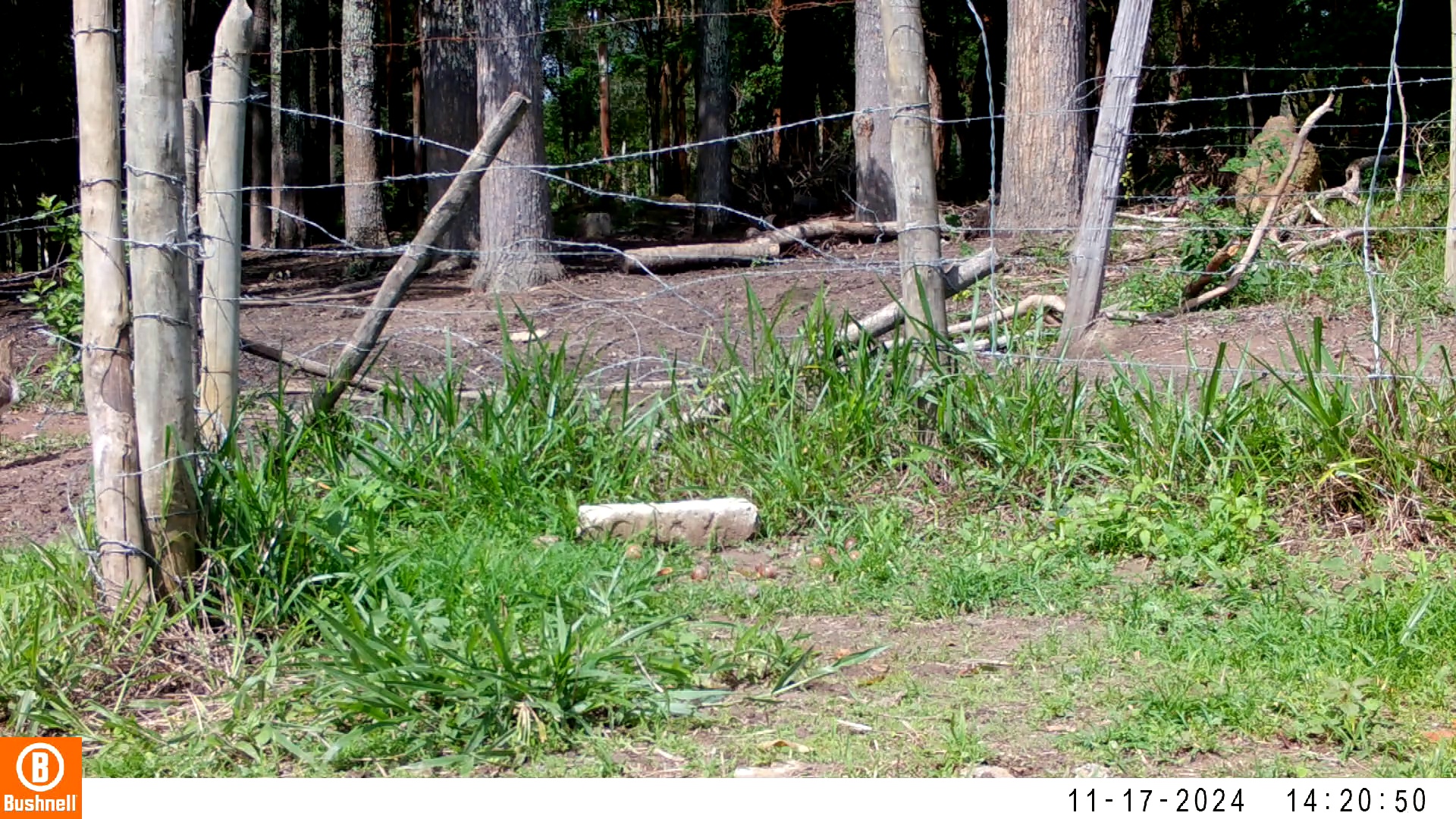

### 11170086-33174-604.jpg

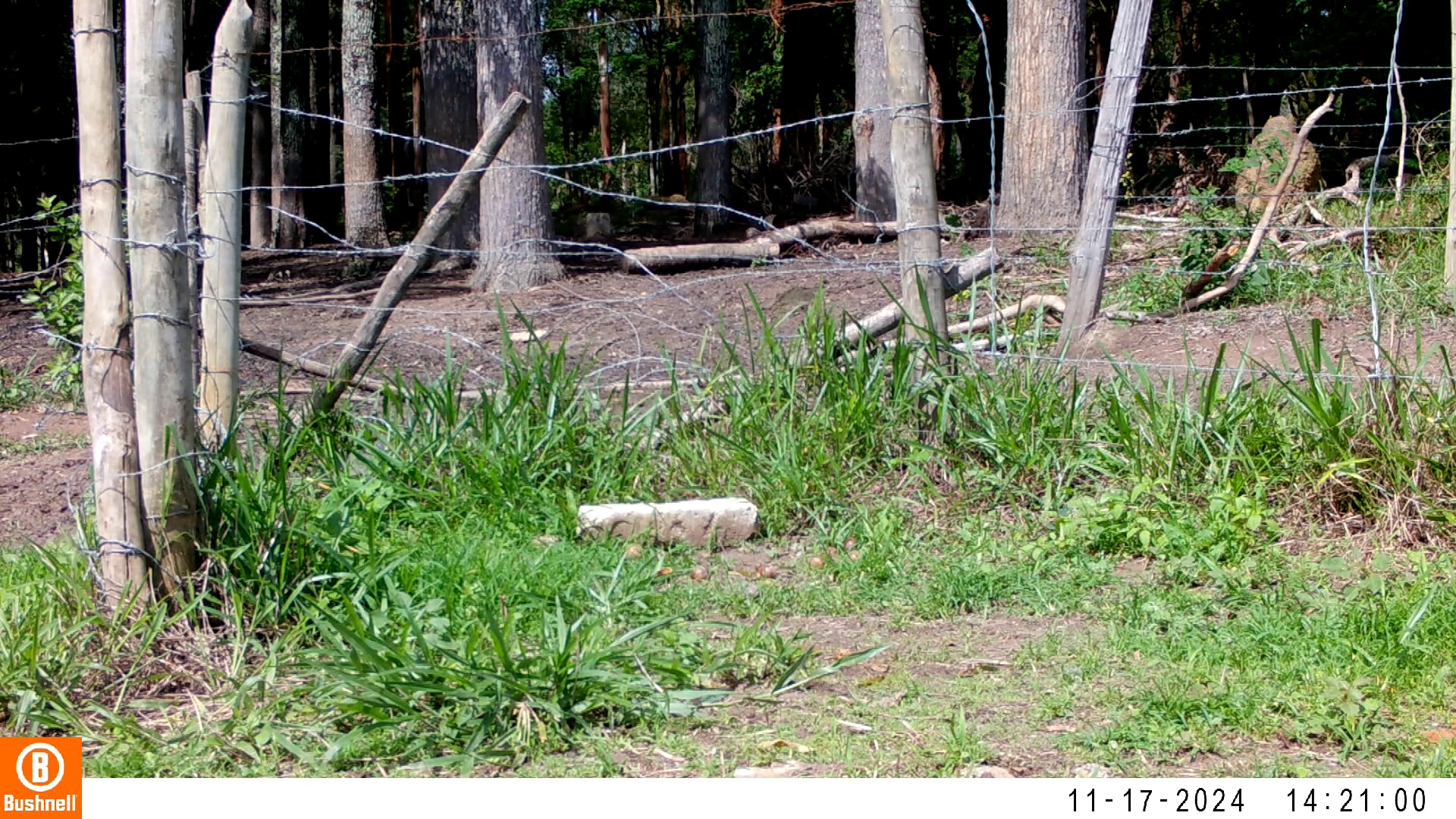

### 11170086-33174-1208.jpg

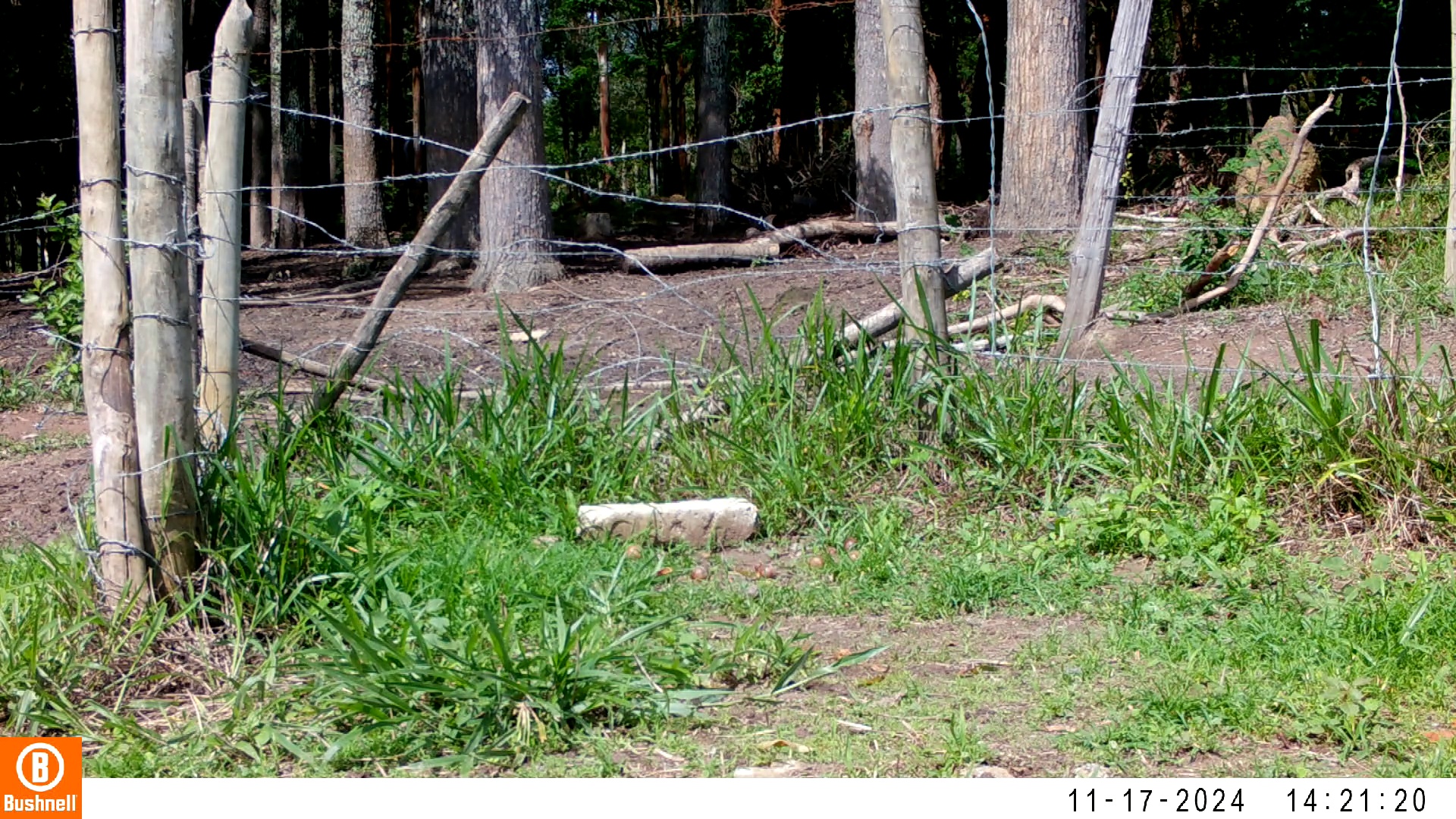

### 11170086-33174-1510.jpg

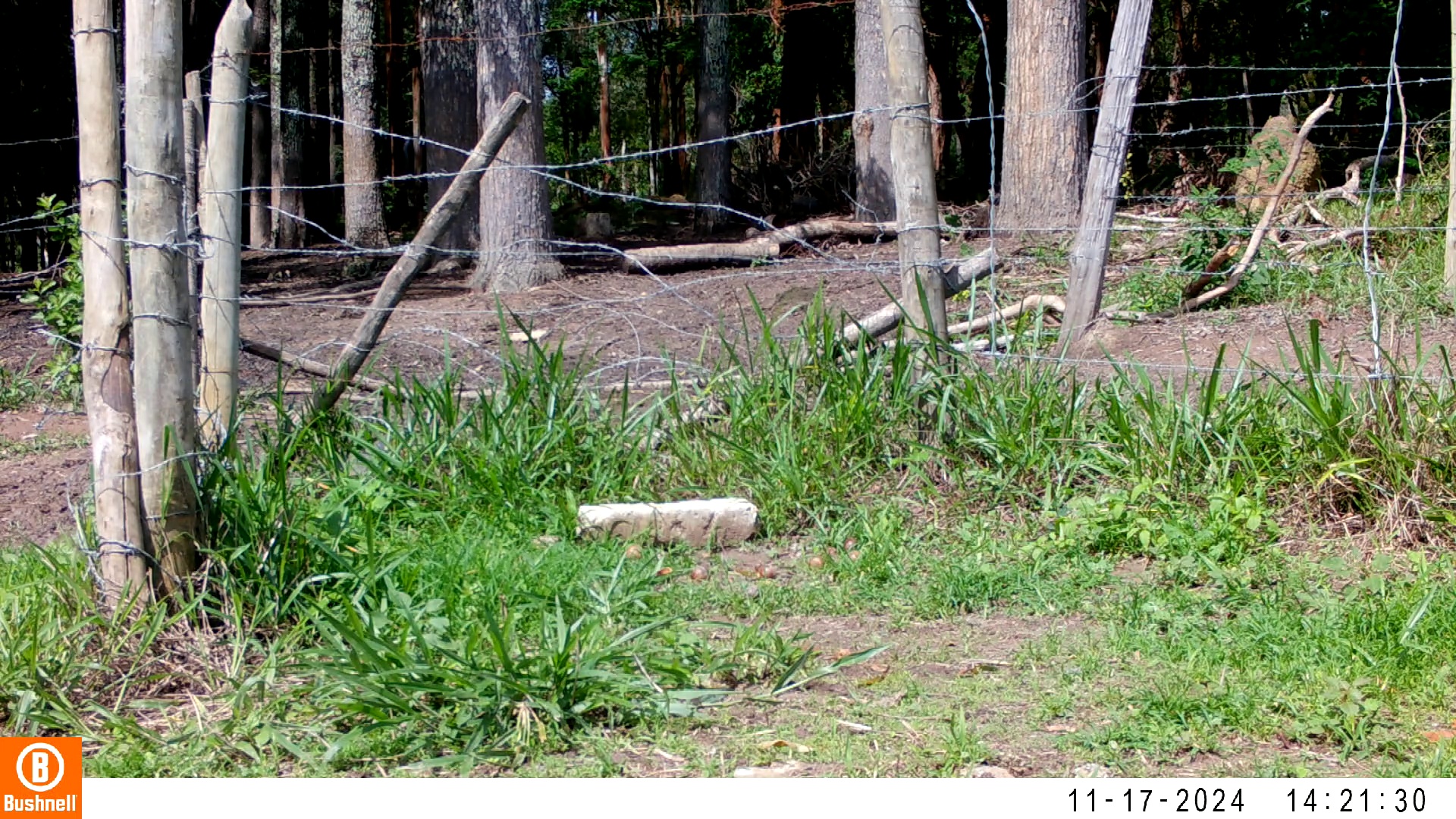

### 11170087-48404-0.jpg

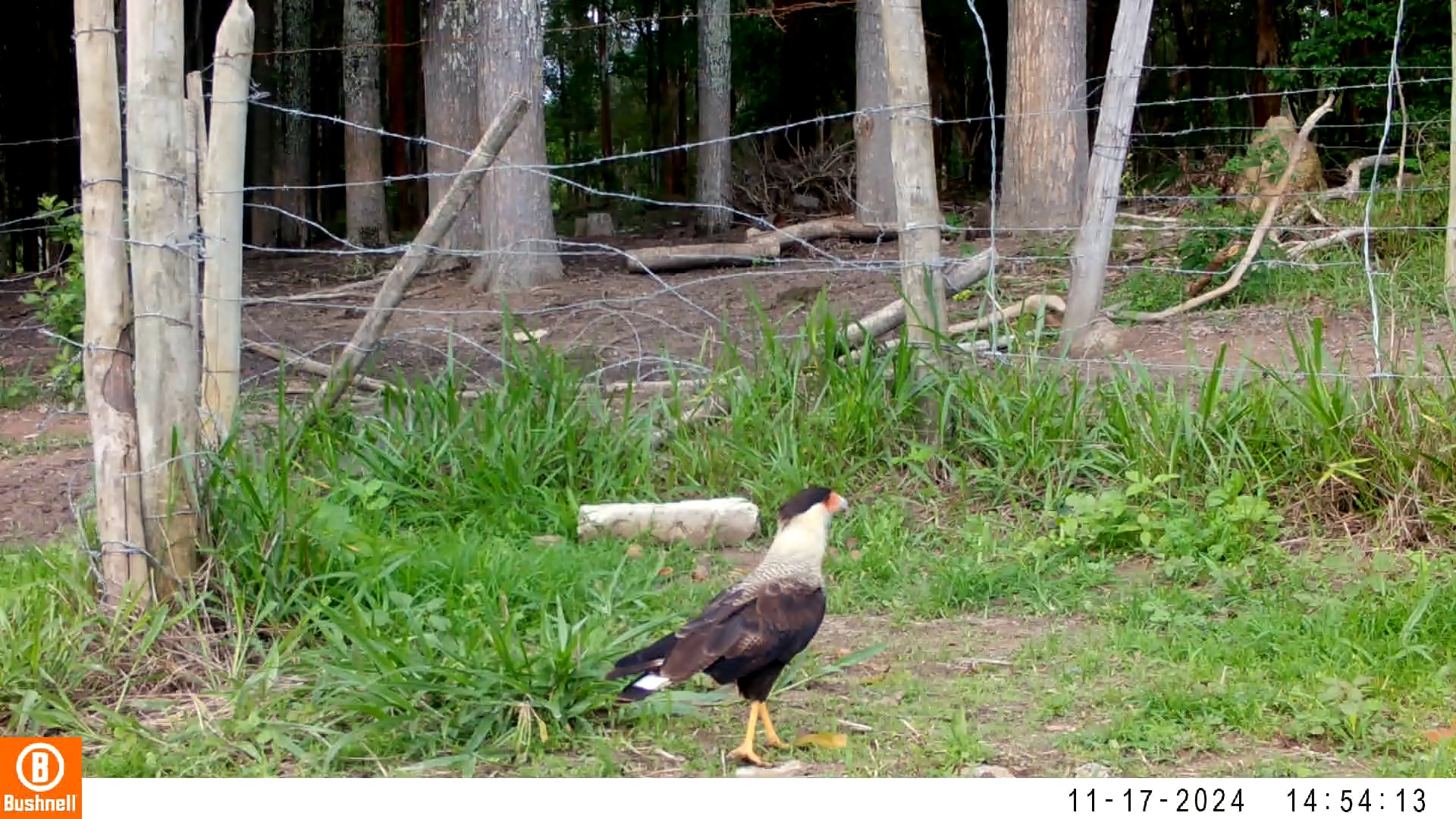

### 11170087-48404-906.jpg

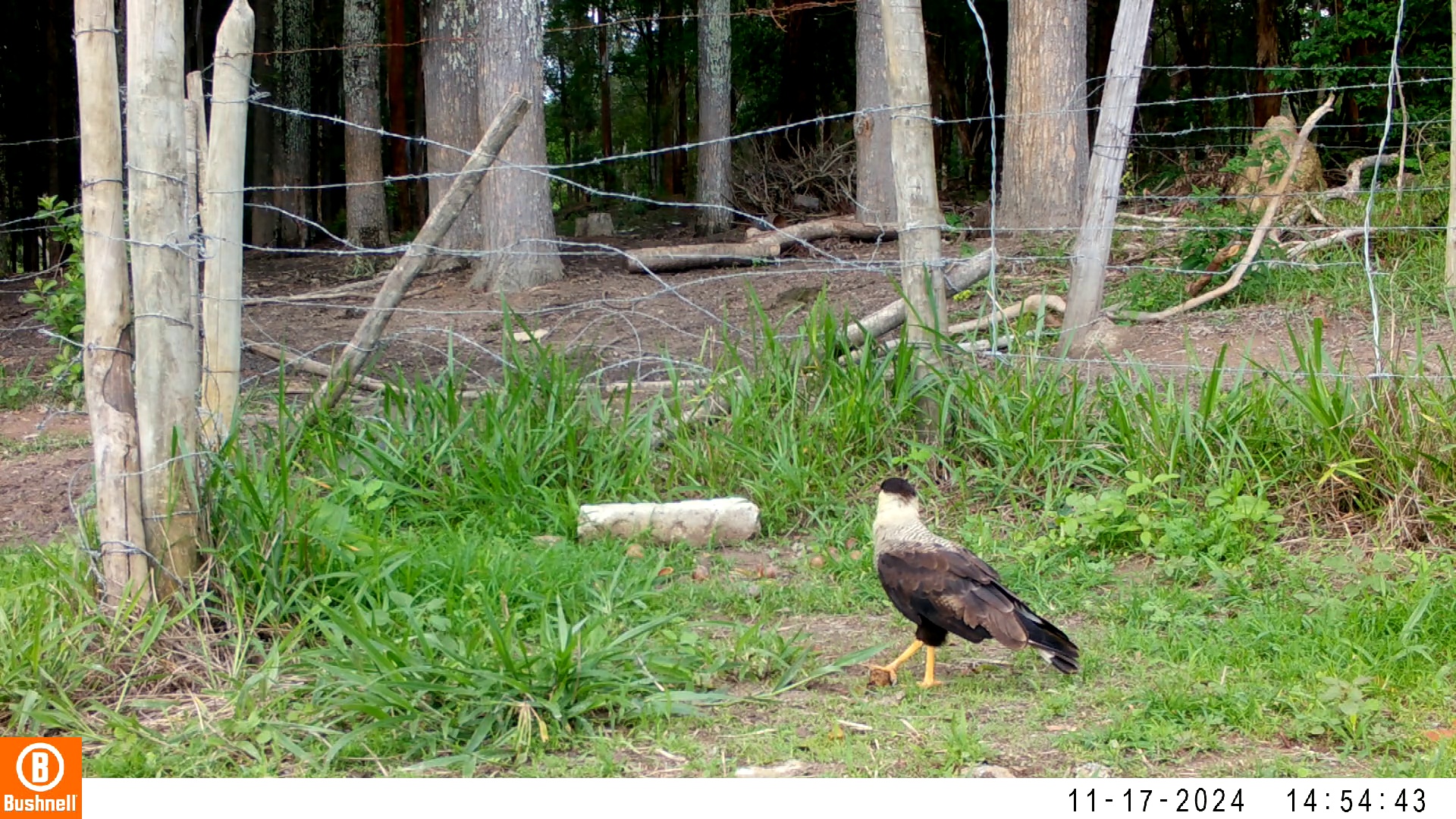

### 11170087-48404-1510.jpg

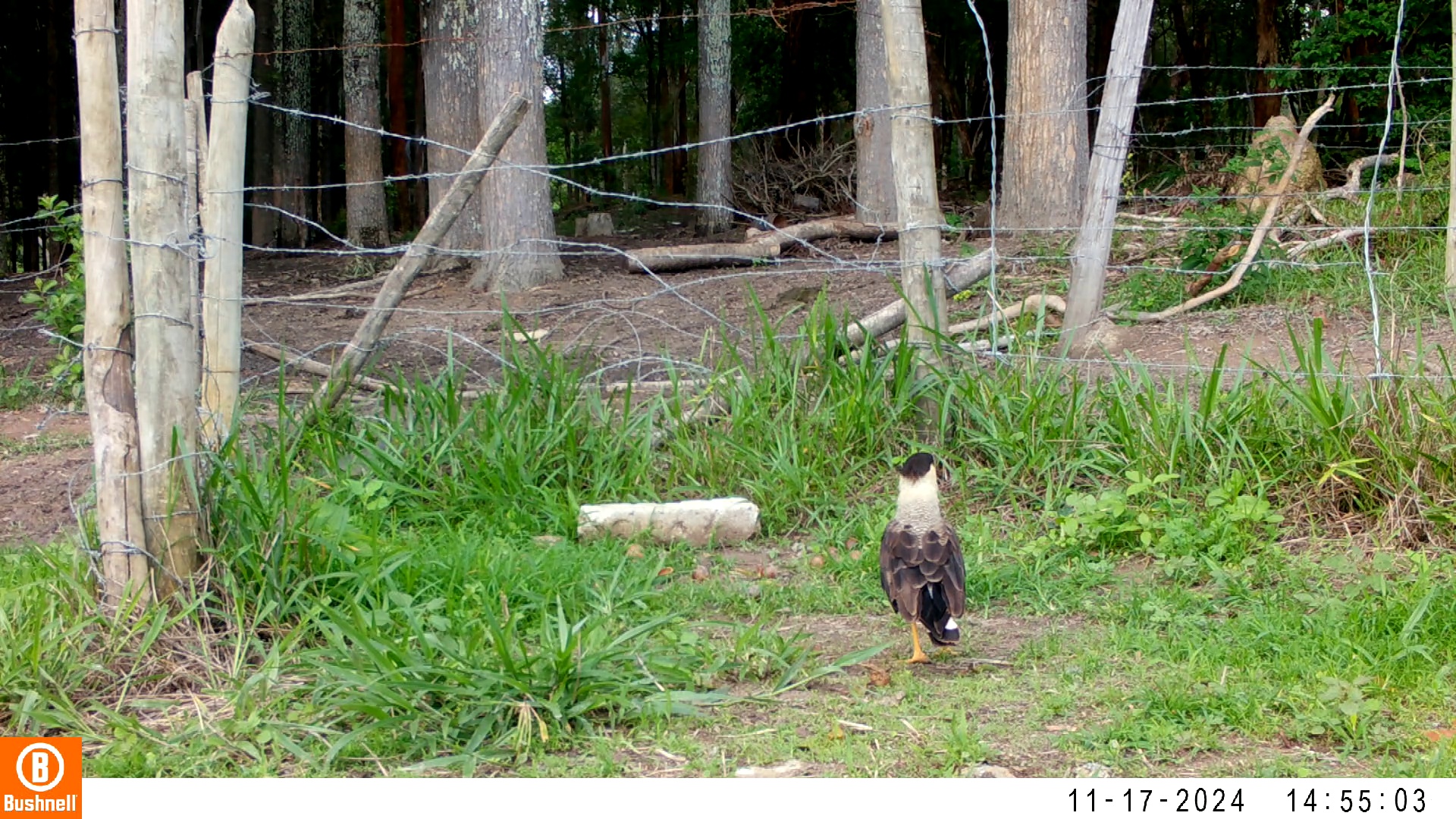

### 11170289-75290-0.jpg

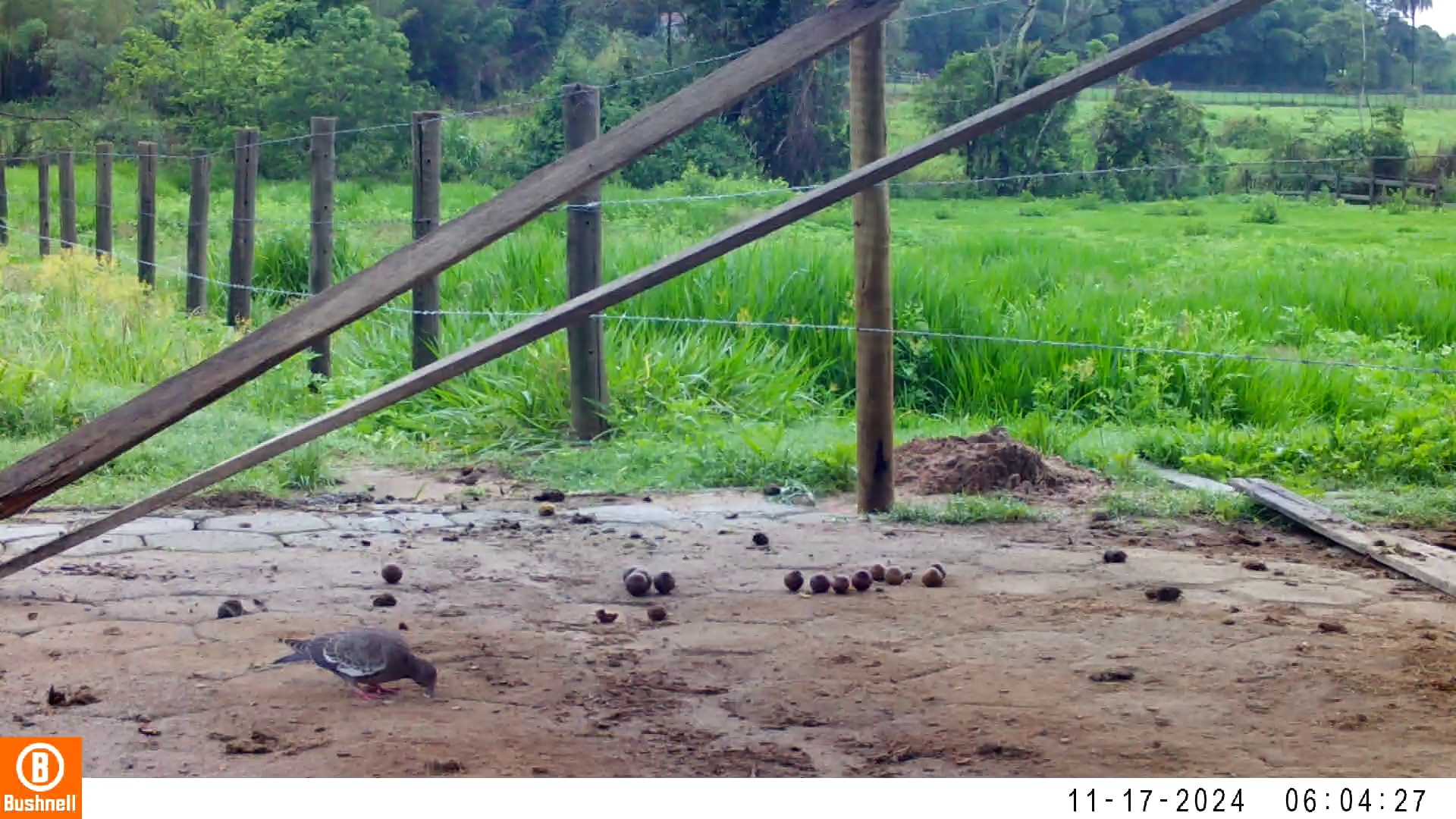

### 11170289-75290-302.jpg

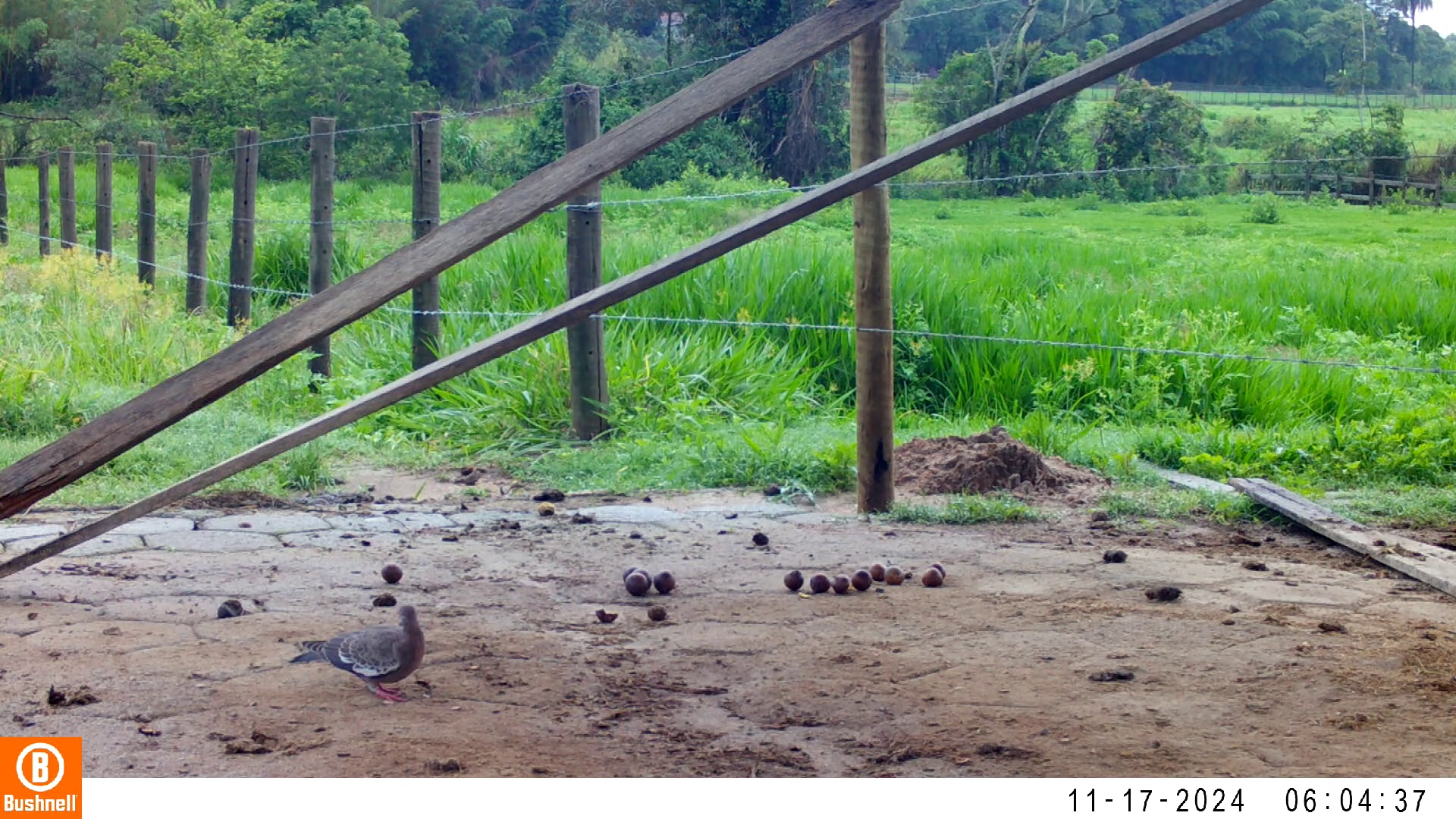

### 11170289-75290-604.jpg

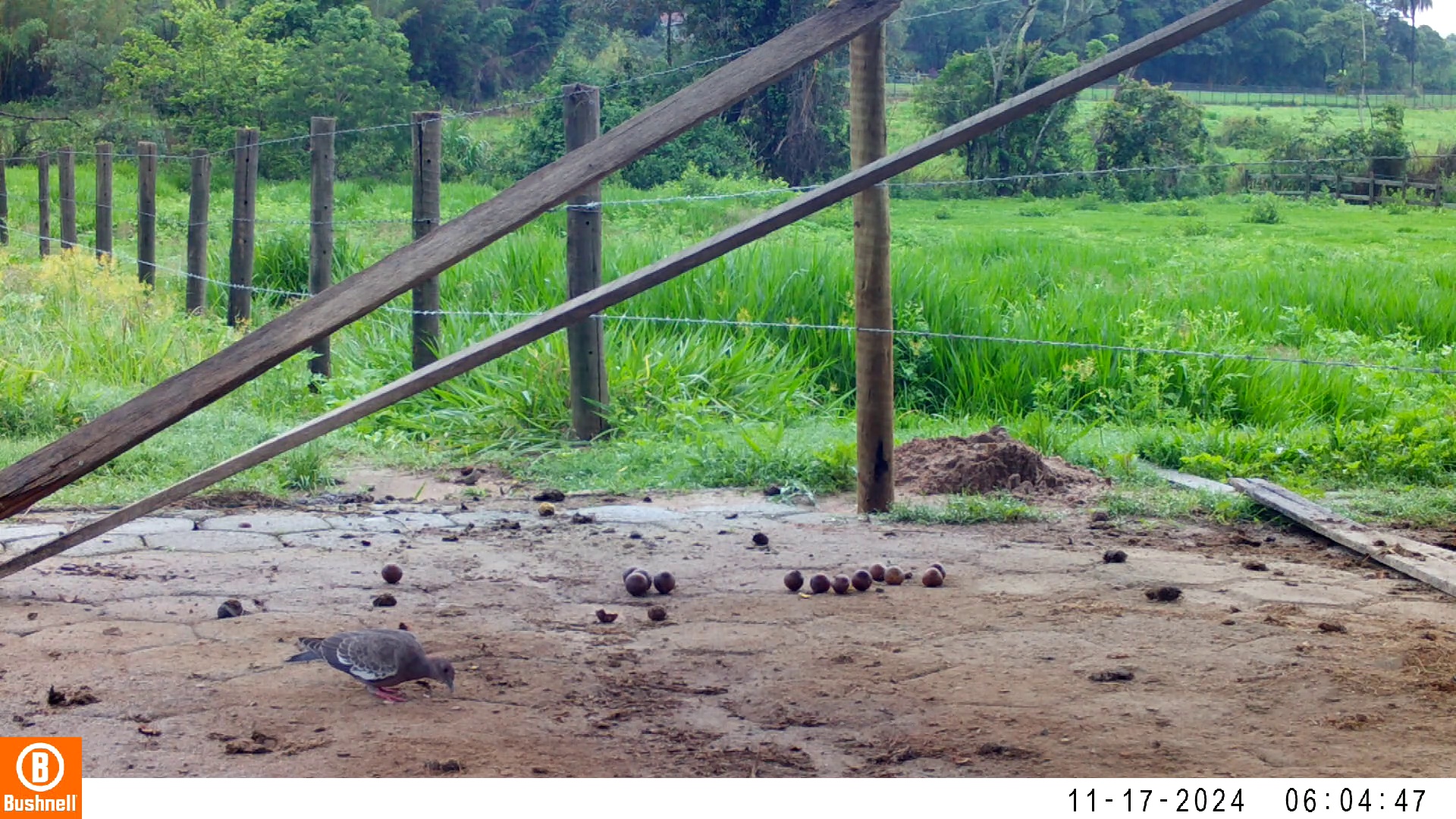

### 11170289-75290-1208.jpg

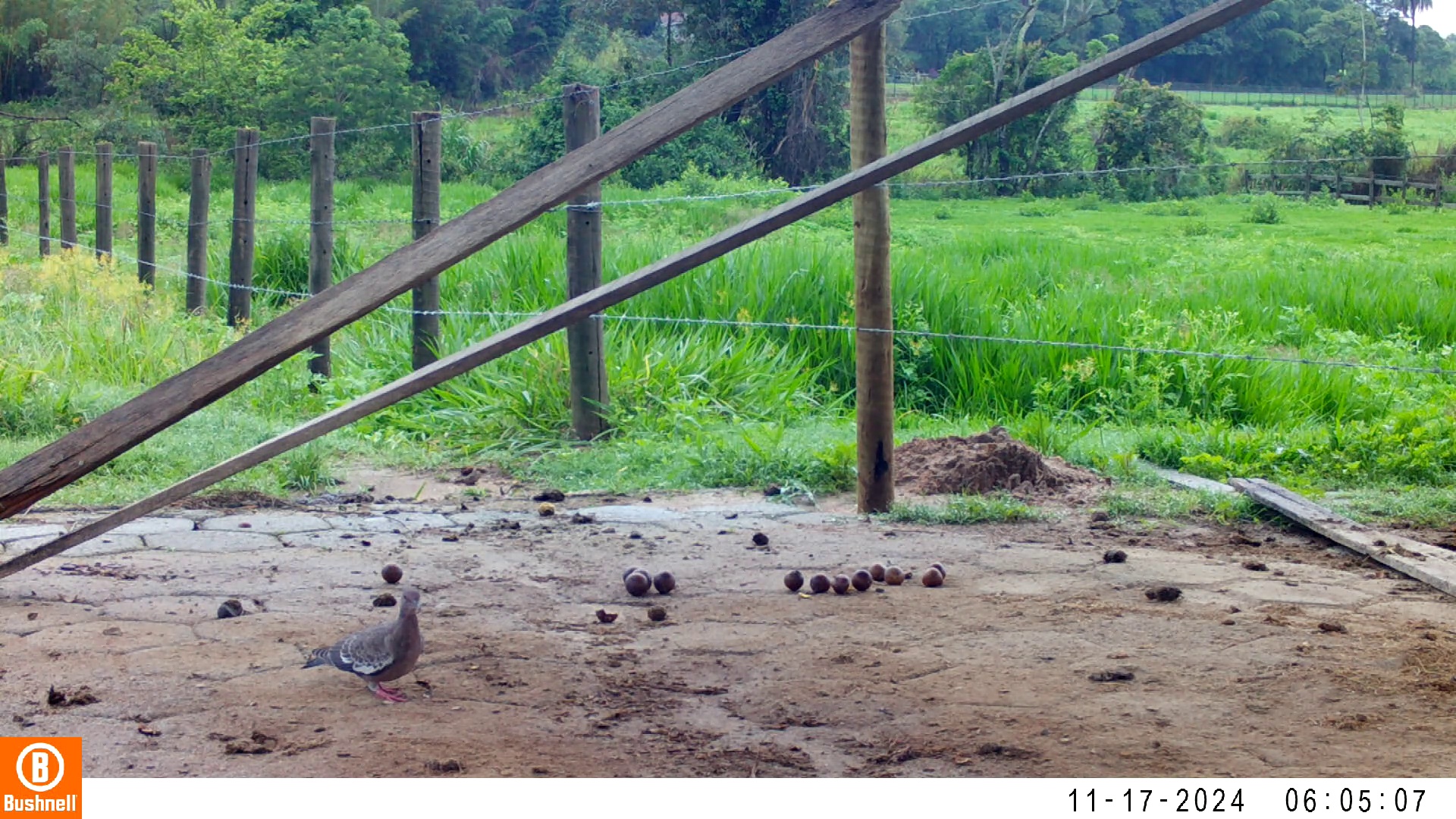

### 11170289-75290-1510.jpg

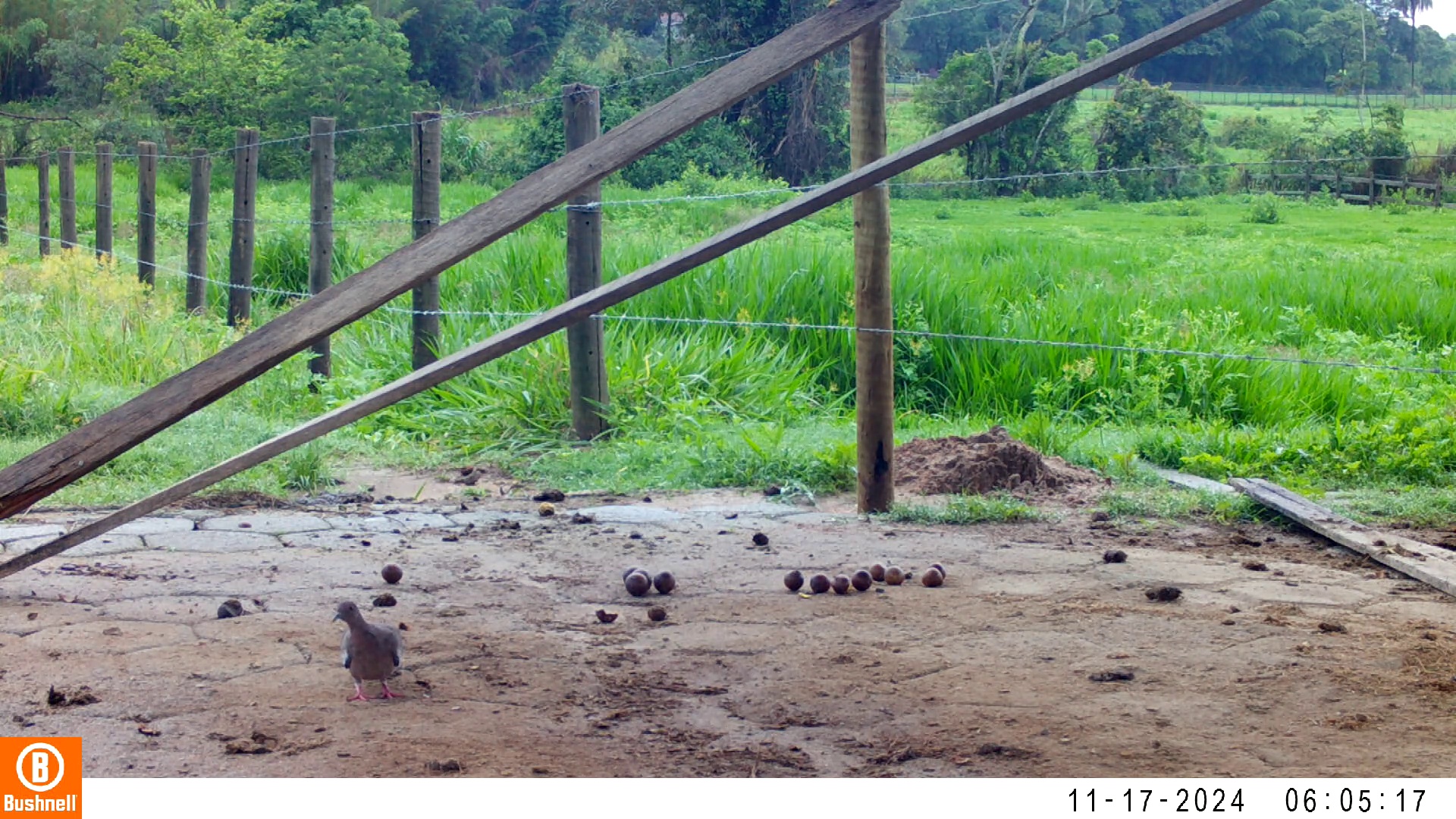

### 11180004-61866-302.jpg

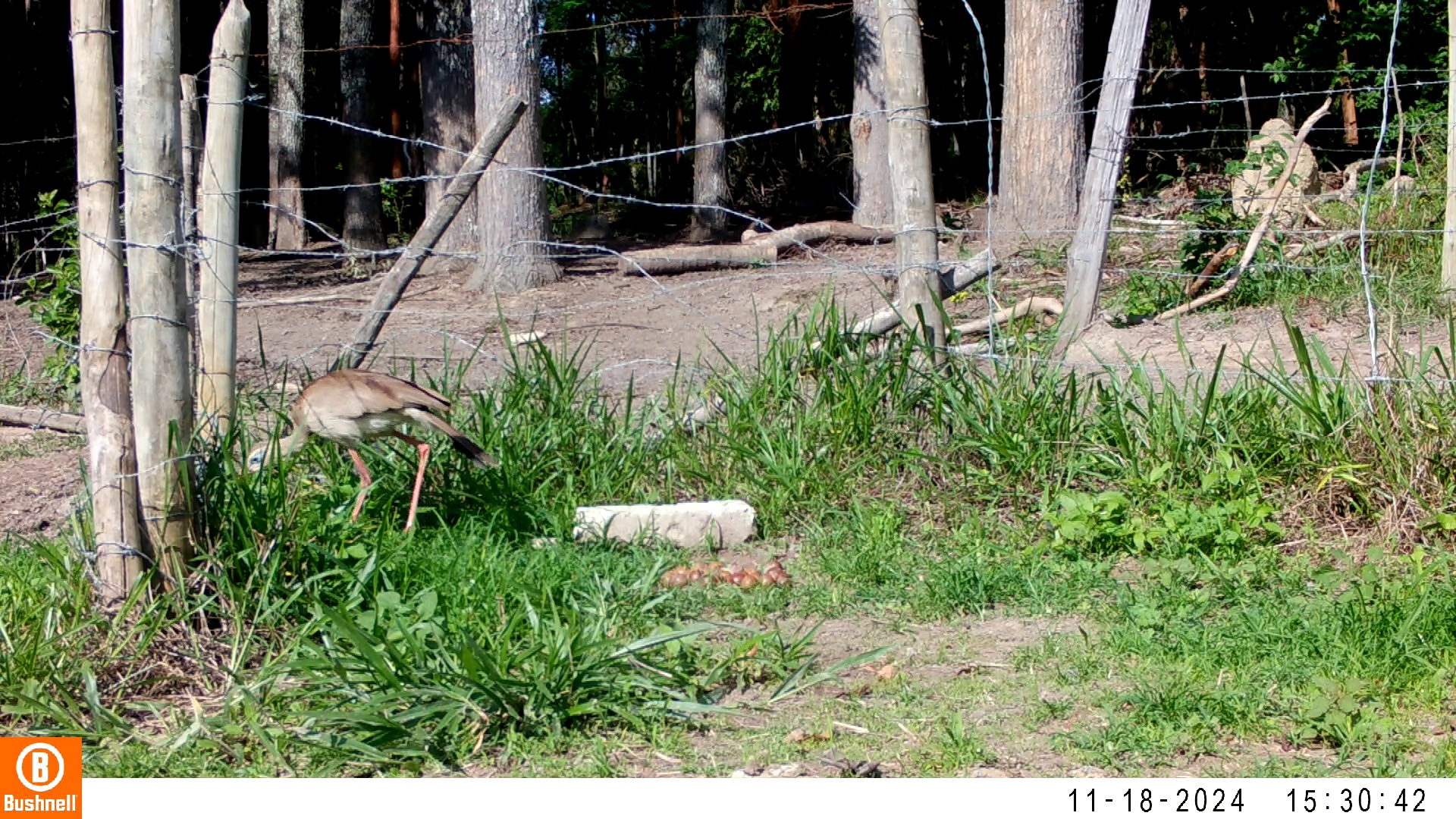

### 11180004-61866-604.jpg

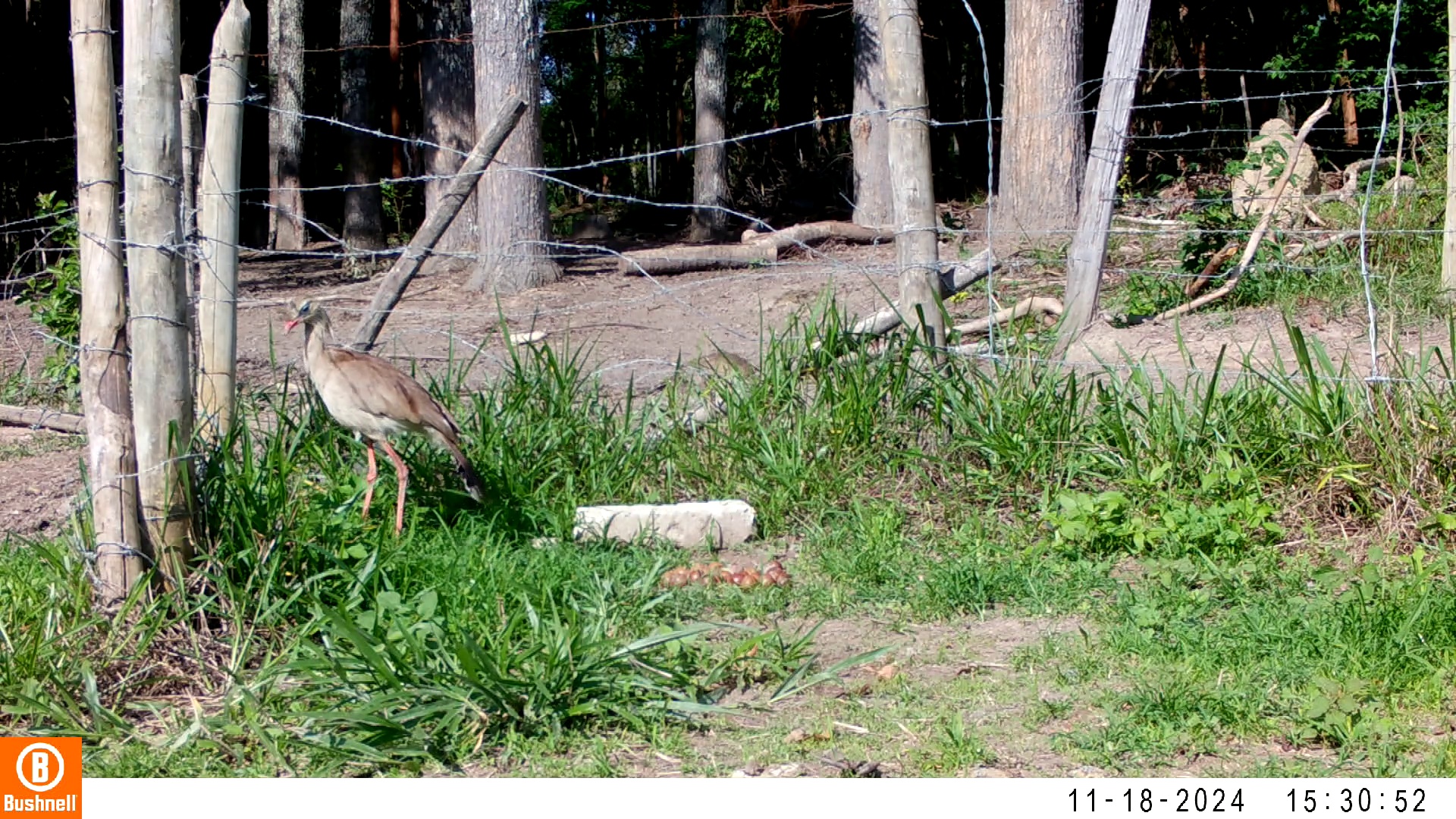

### 11180004-61866-1208.jpg

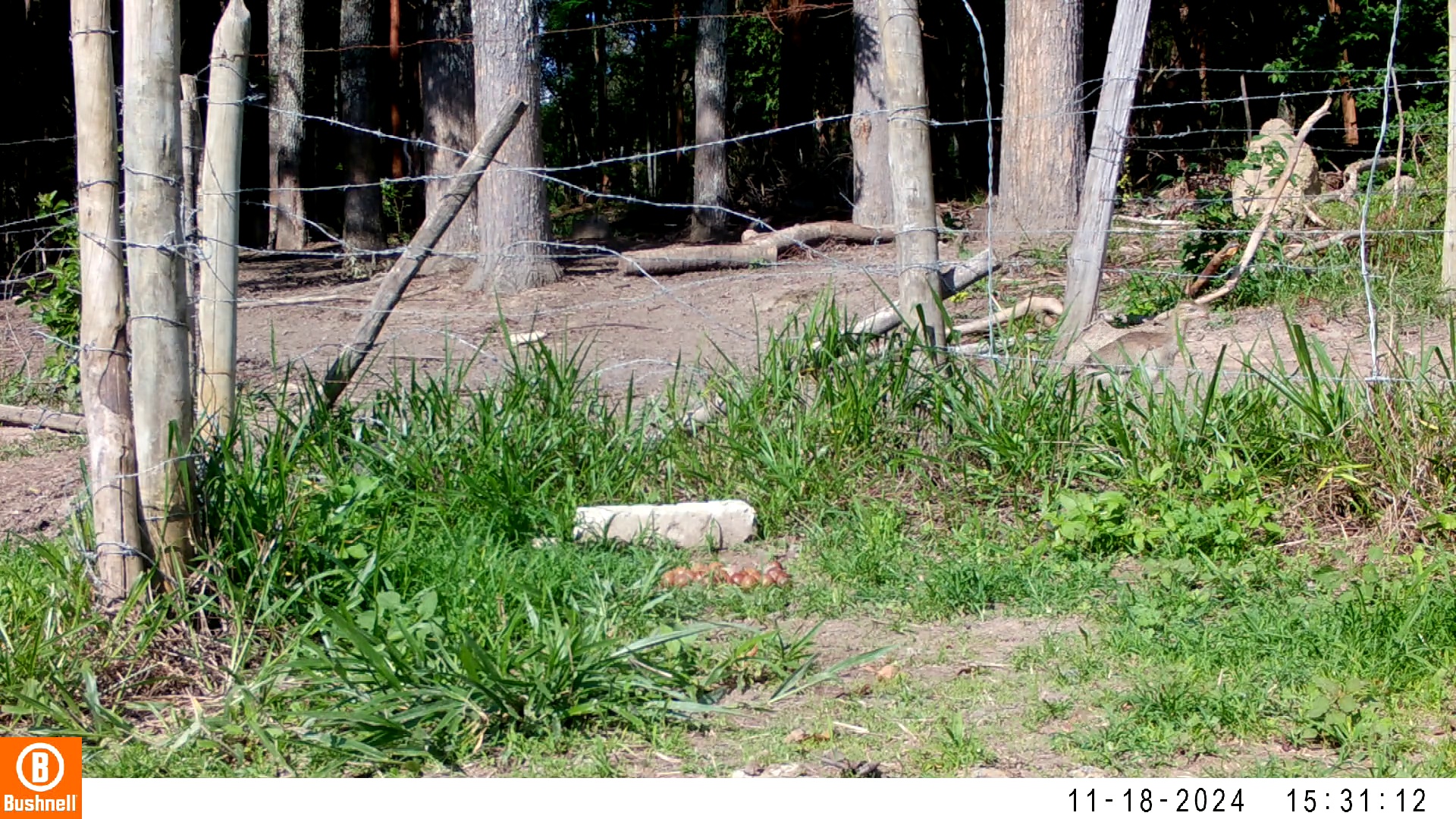

### 11180004-61866-1510.jpg

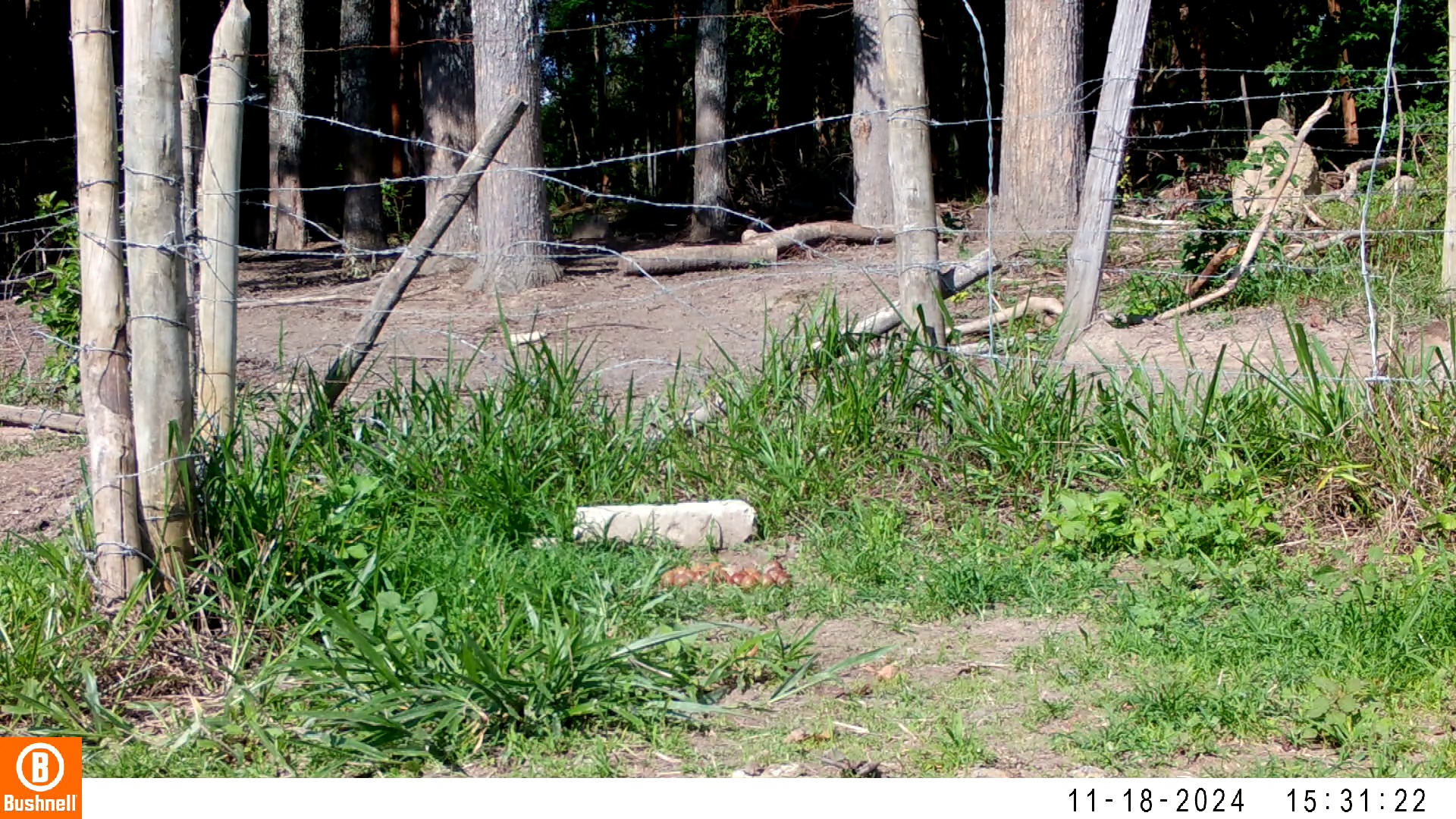

### 11180023-93292-0.jpg

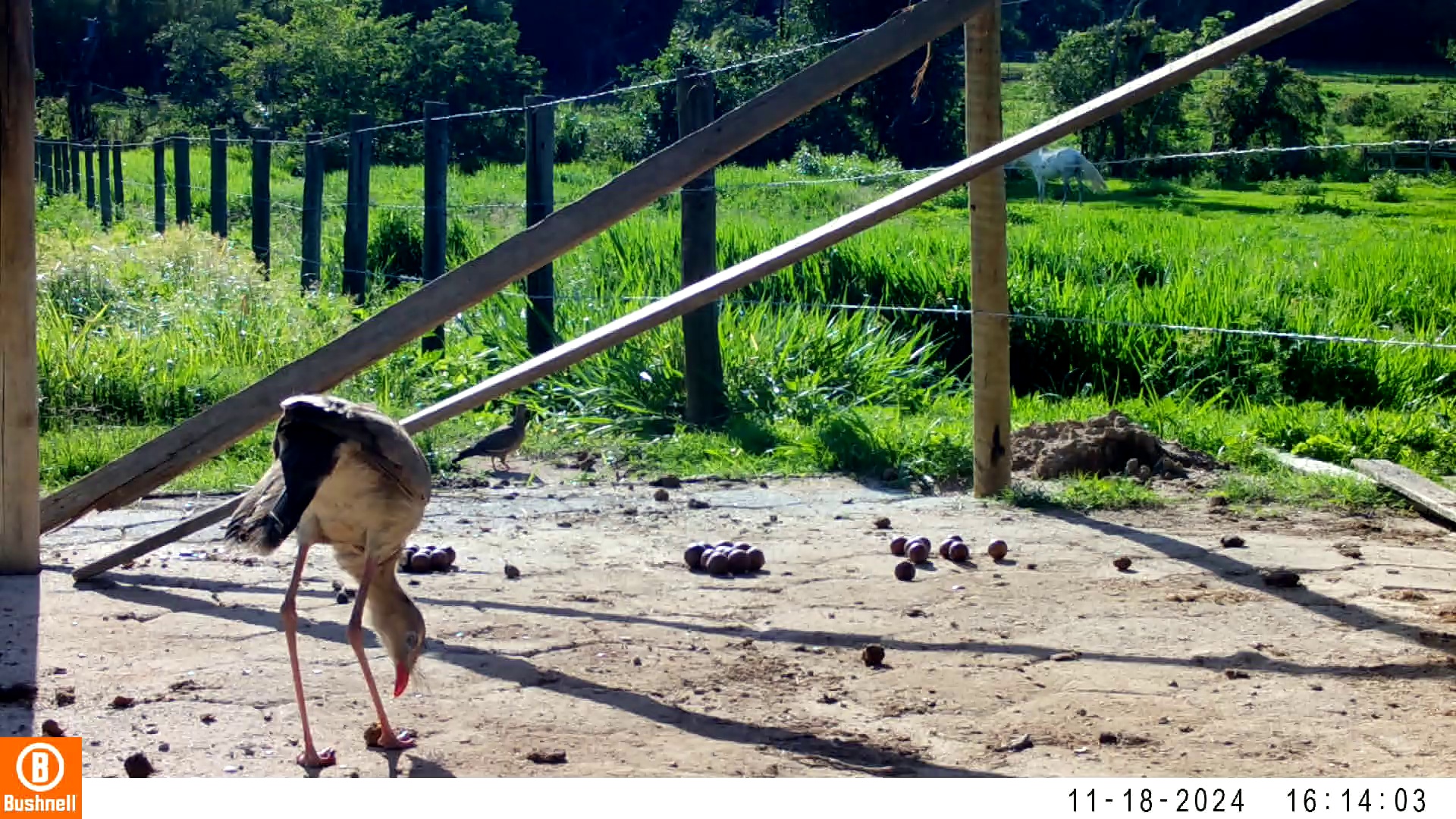

### 11180023-93292-302.jpg

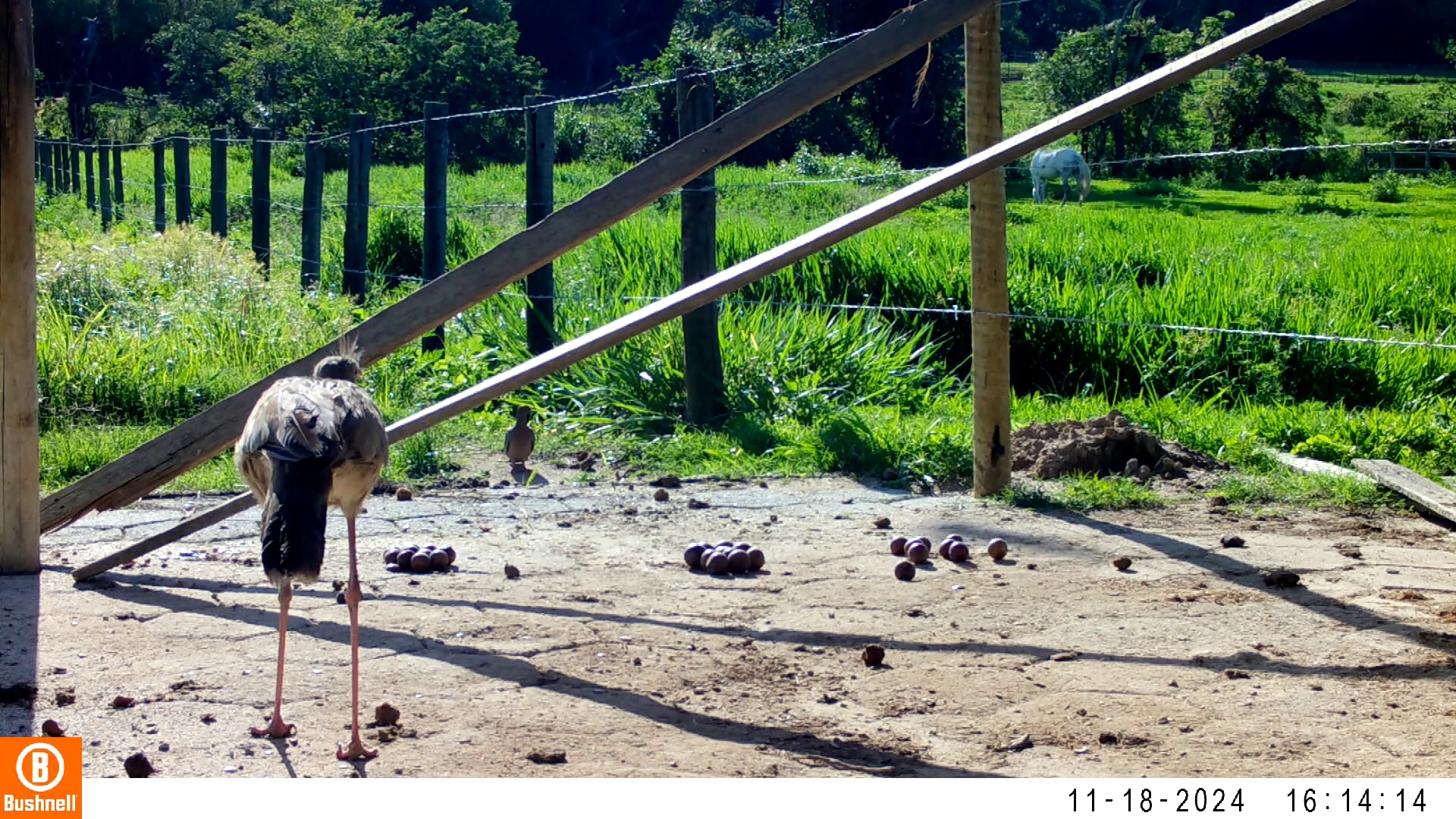

### 11180023-93292-906.jpg

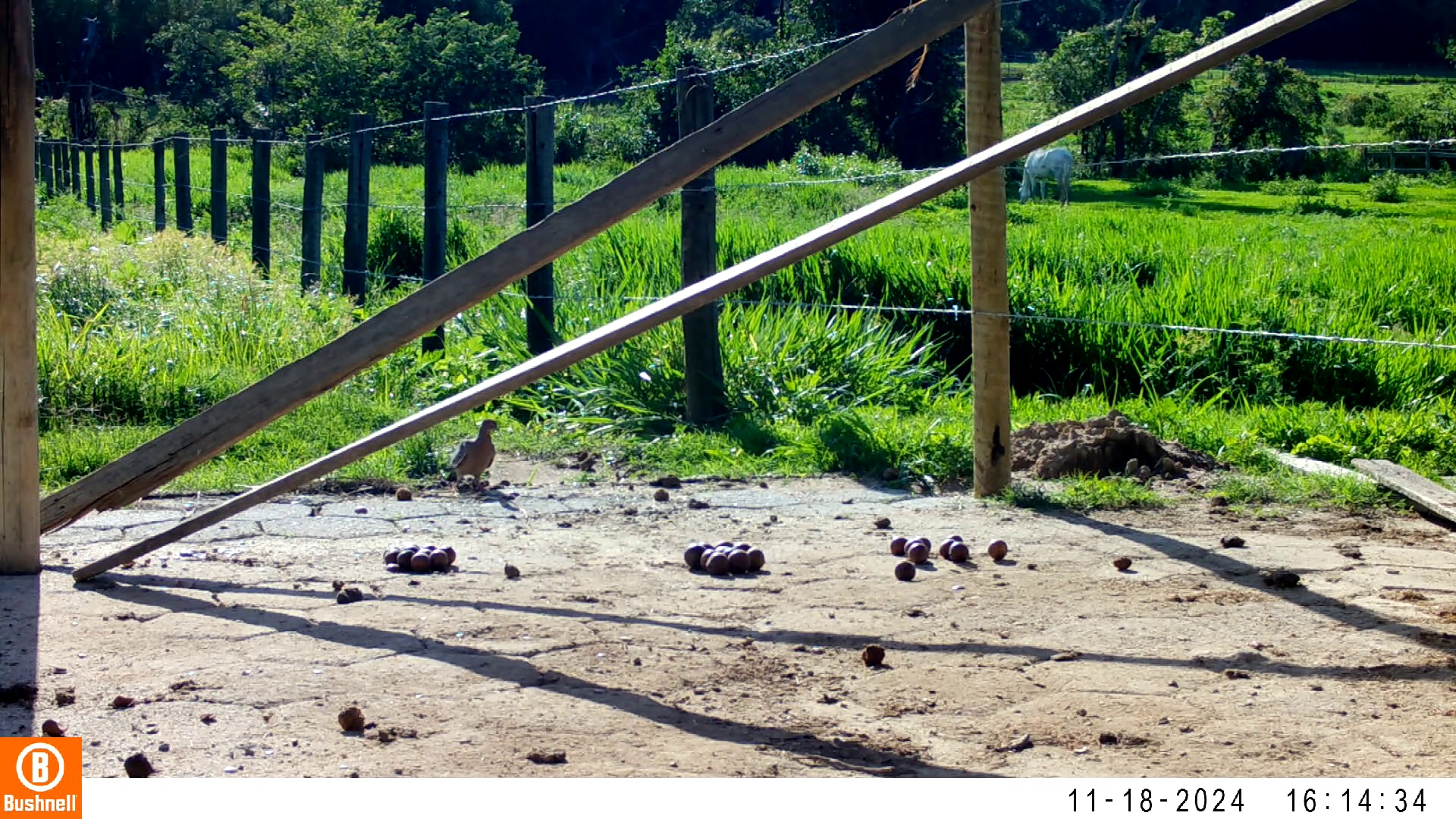

### 11180023-93292-1208.jpg

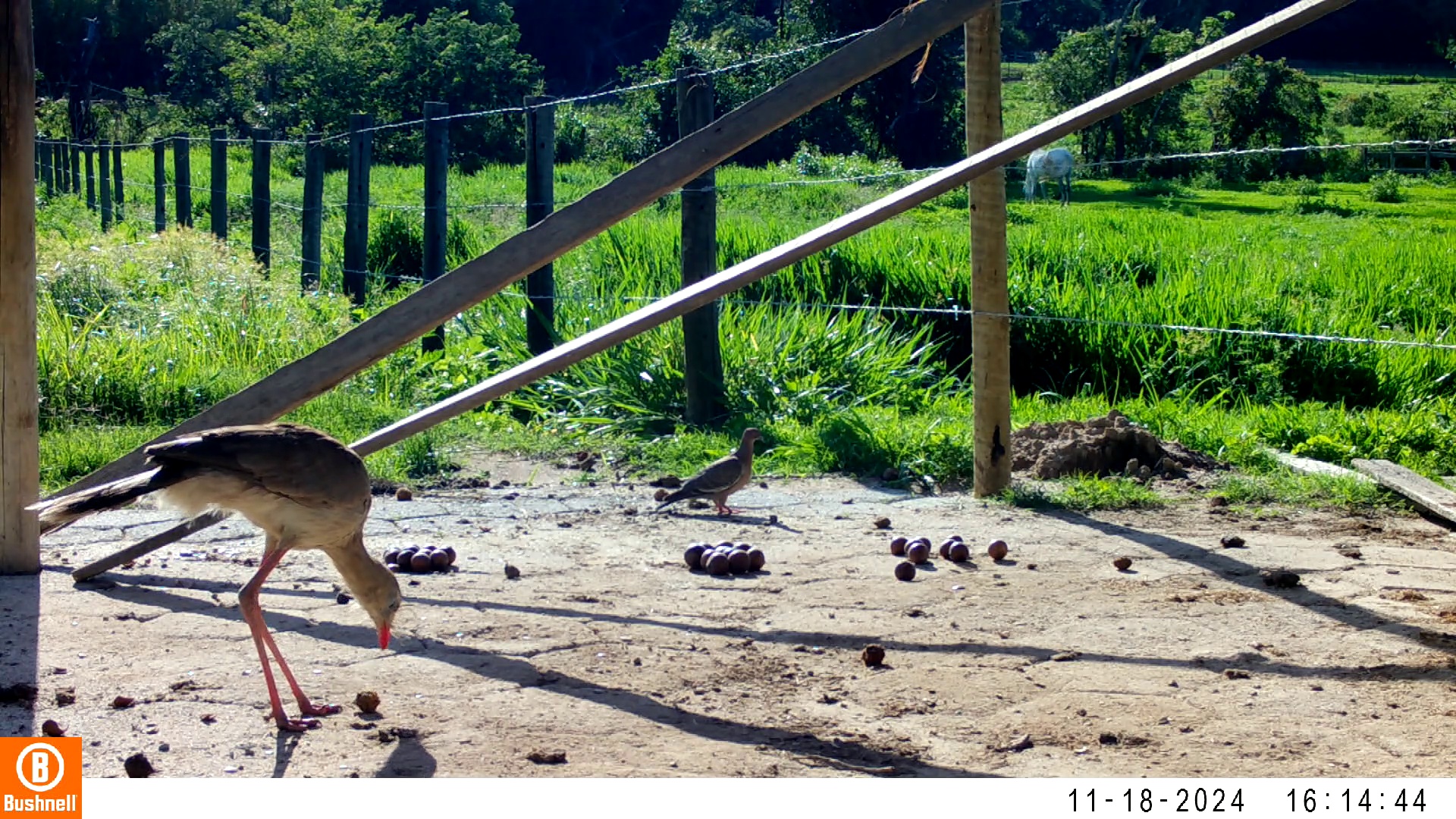

### 11180023-93292-1510.jpg

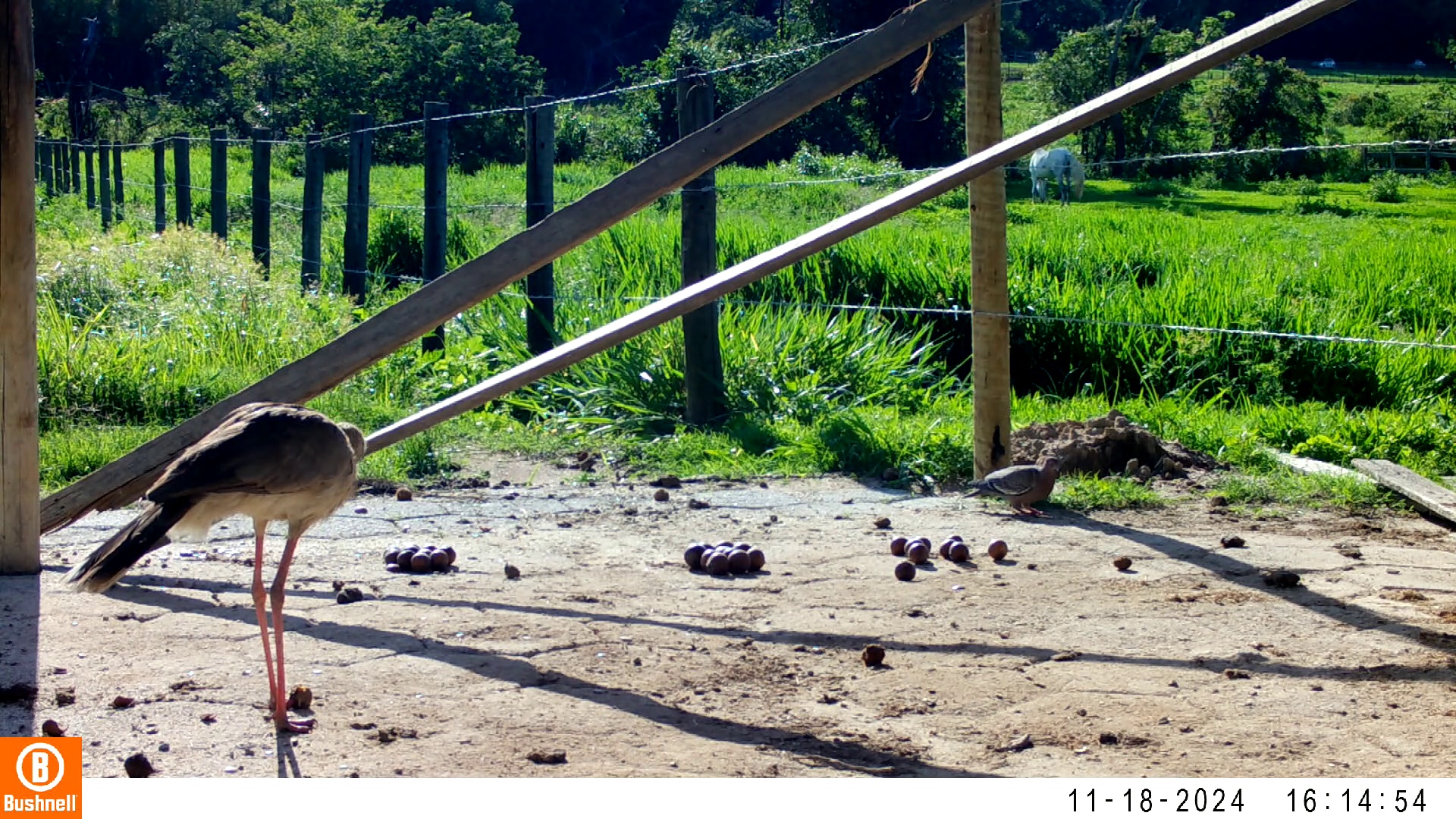

### 11180024-48638-0.jpg

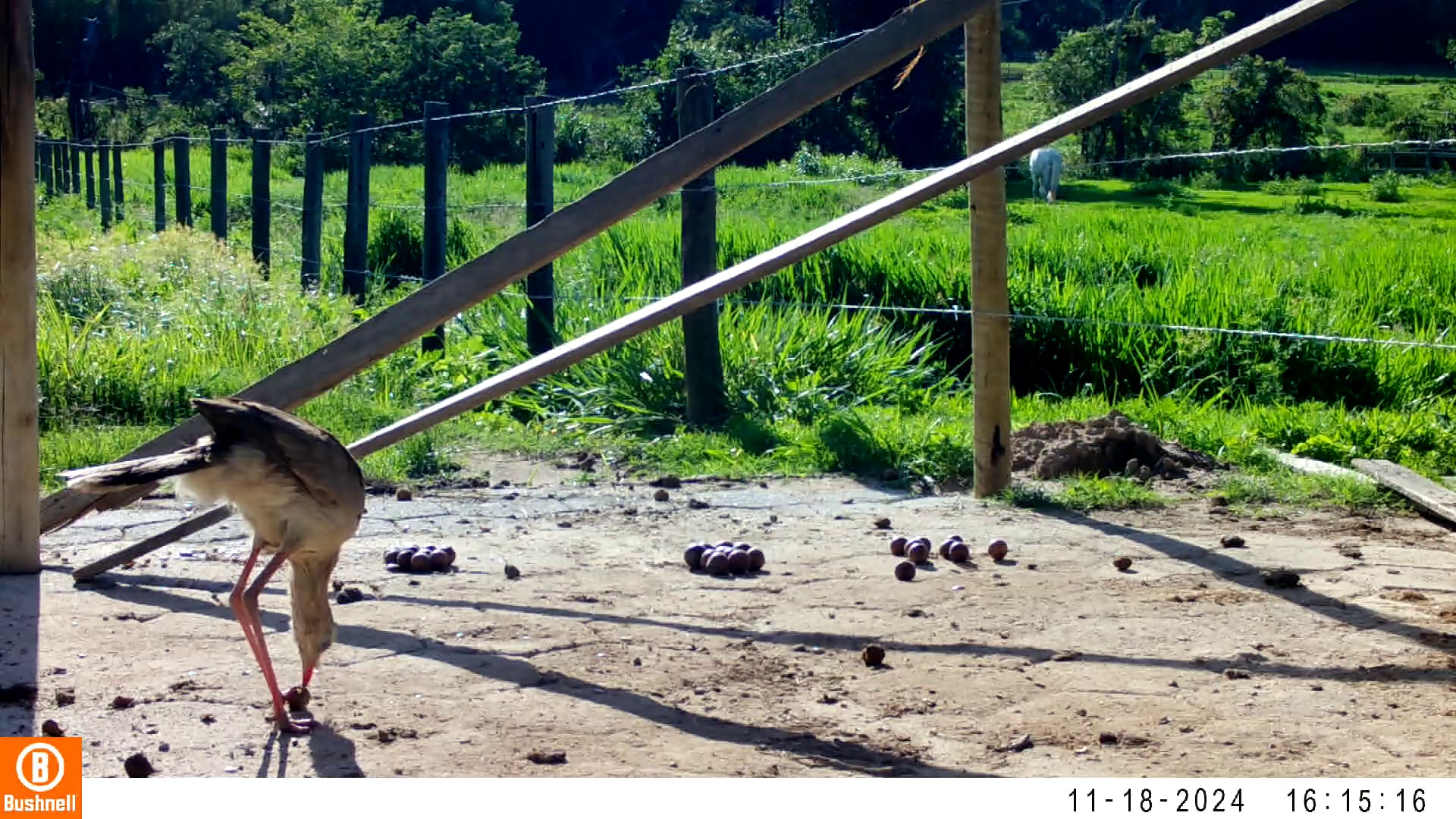

### 11180024-48638-302.jpg

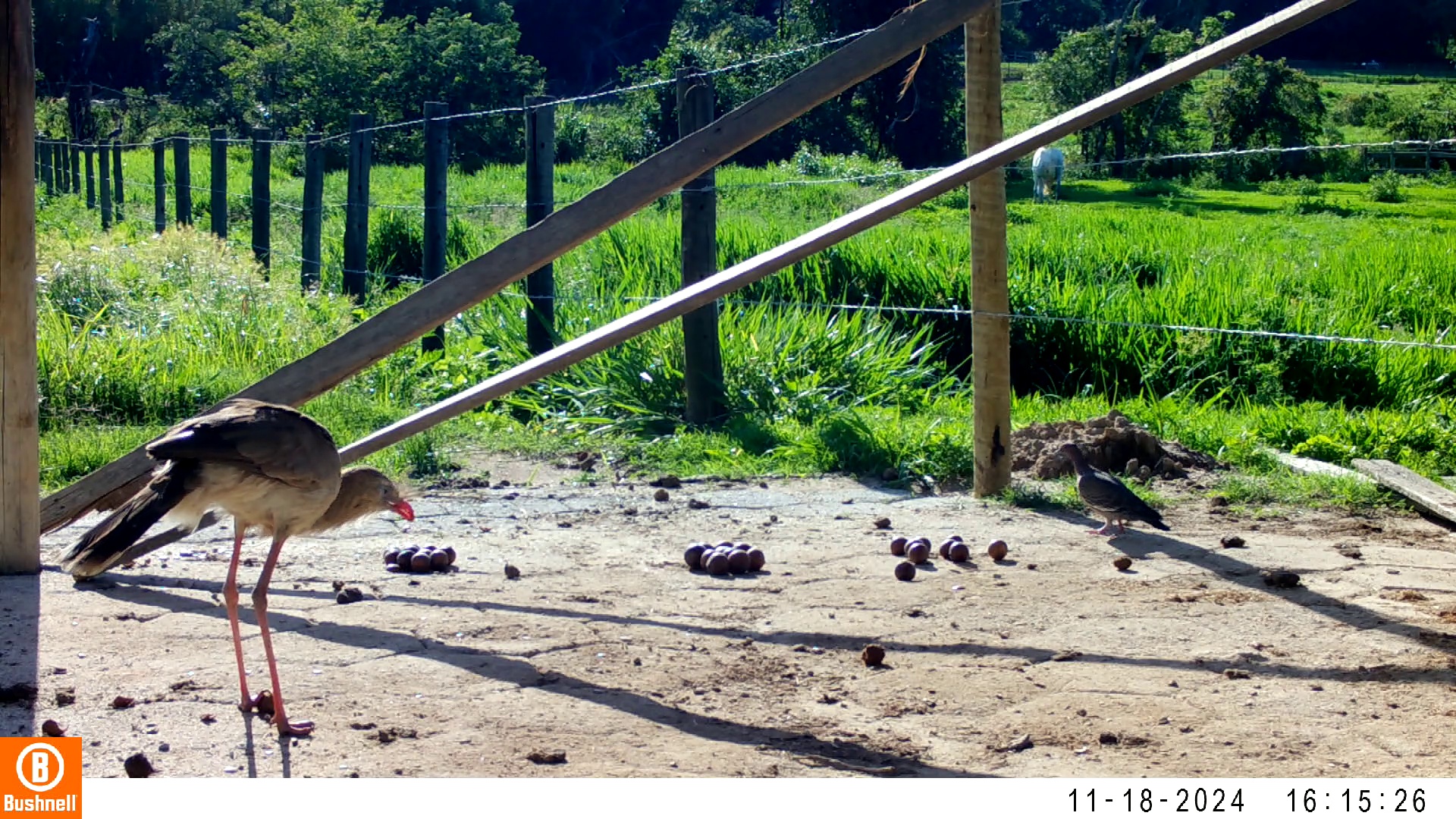

### 11180024-48638-604.jpg

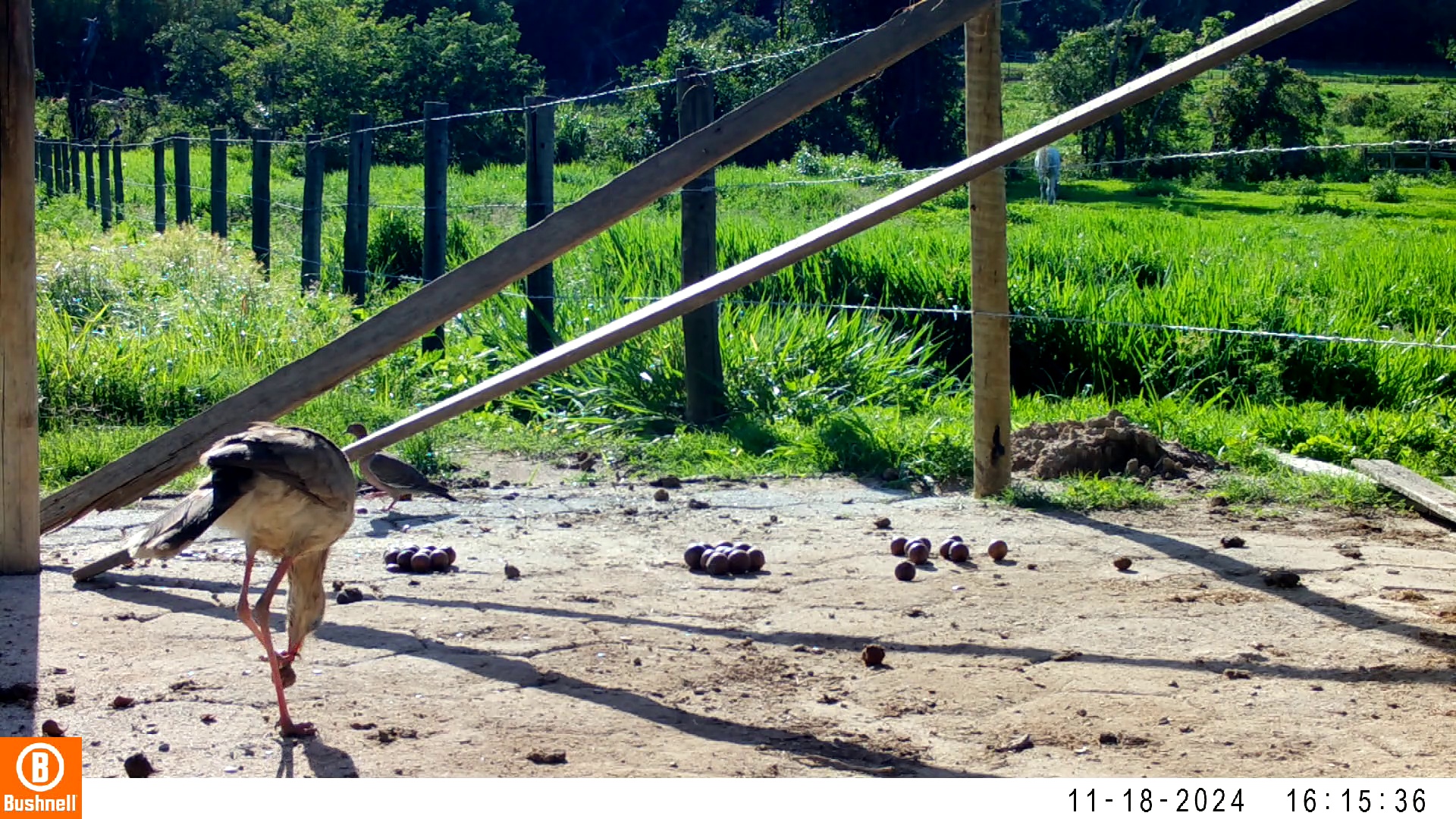

### 11180024-48638-1208.jpg

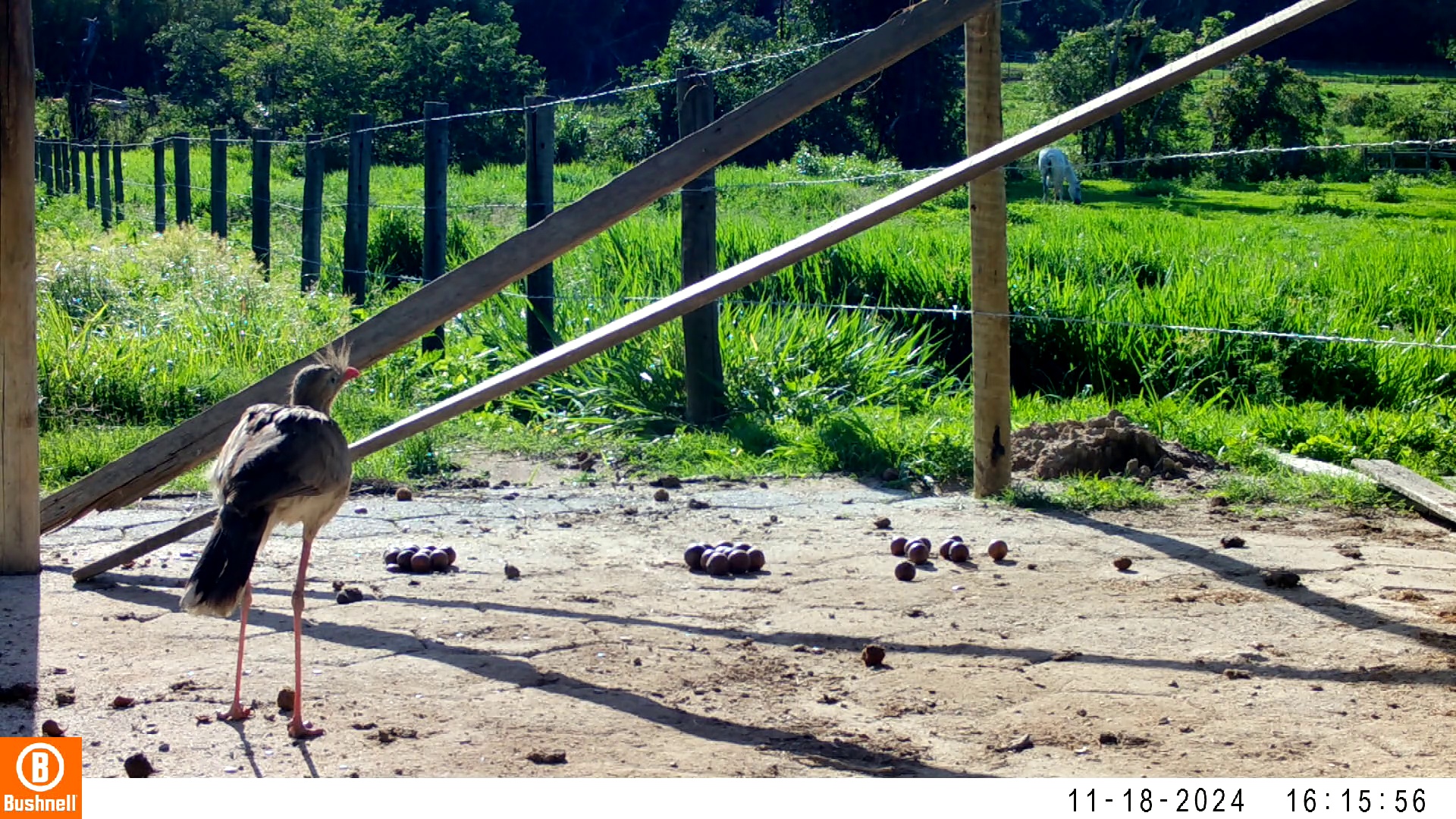

### 11180027-07363-0.jpg

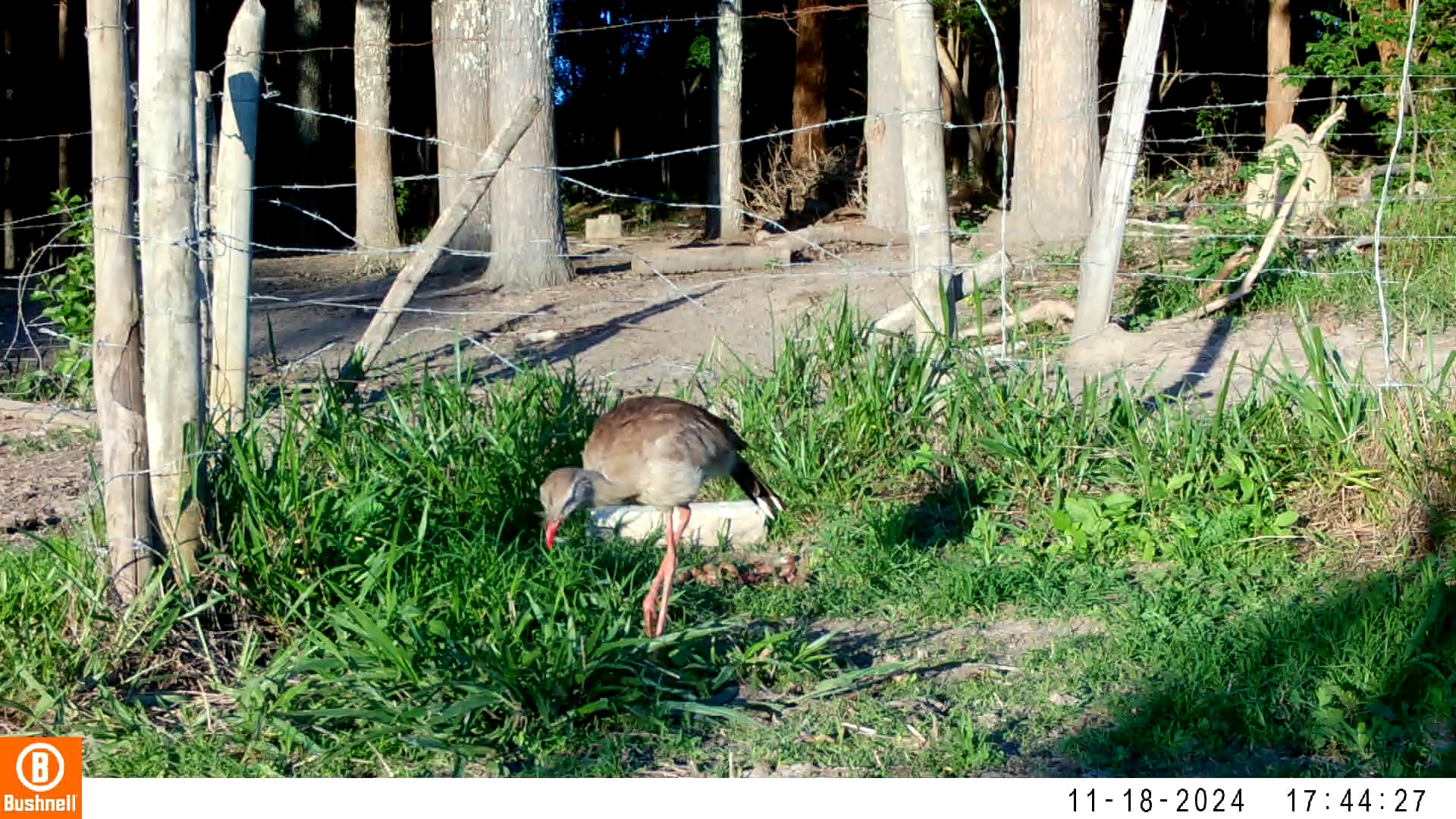

### 11180027-07363-302.jpg

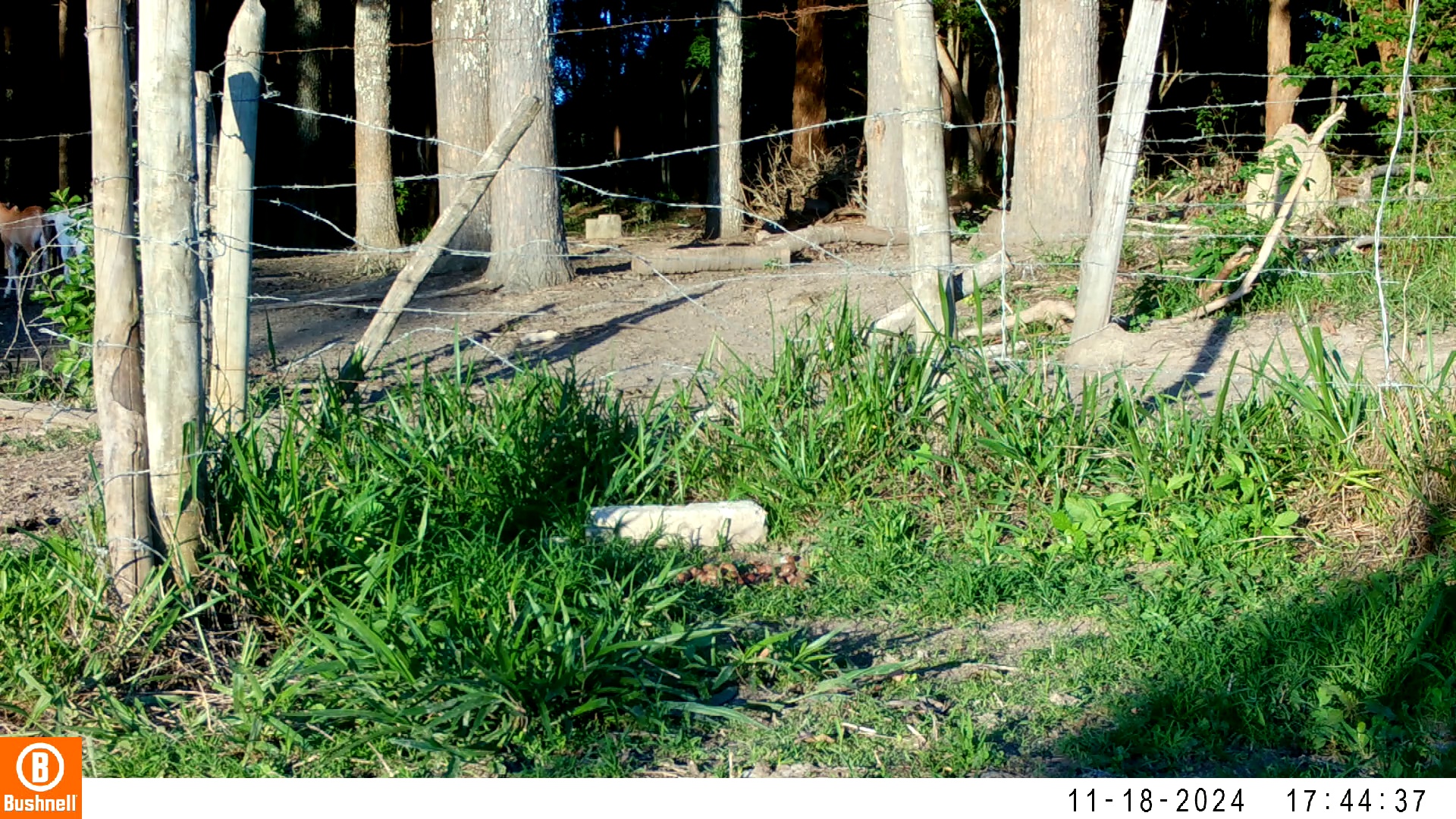

### 11180027-07363-604.jpg

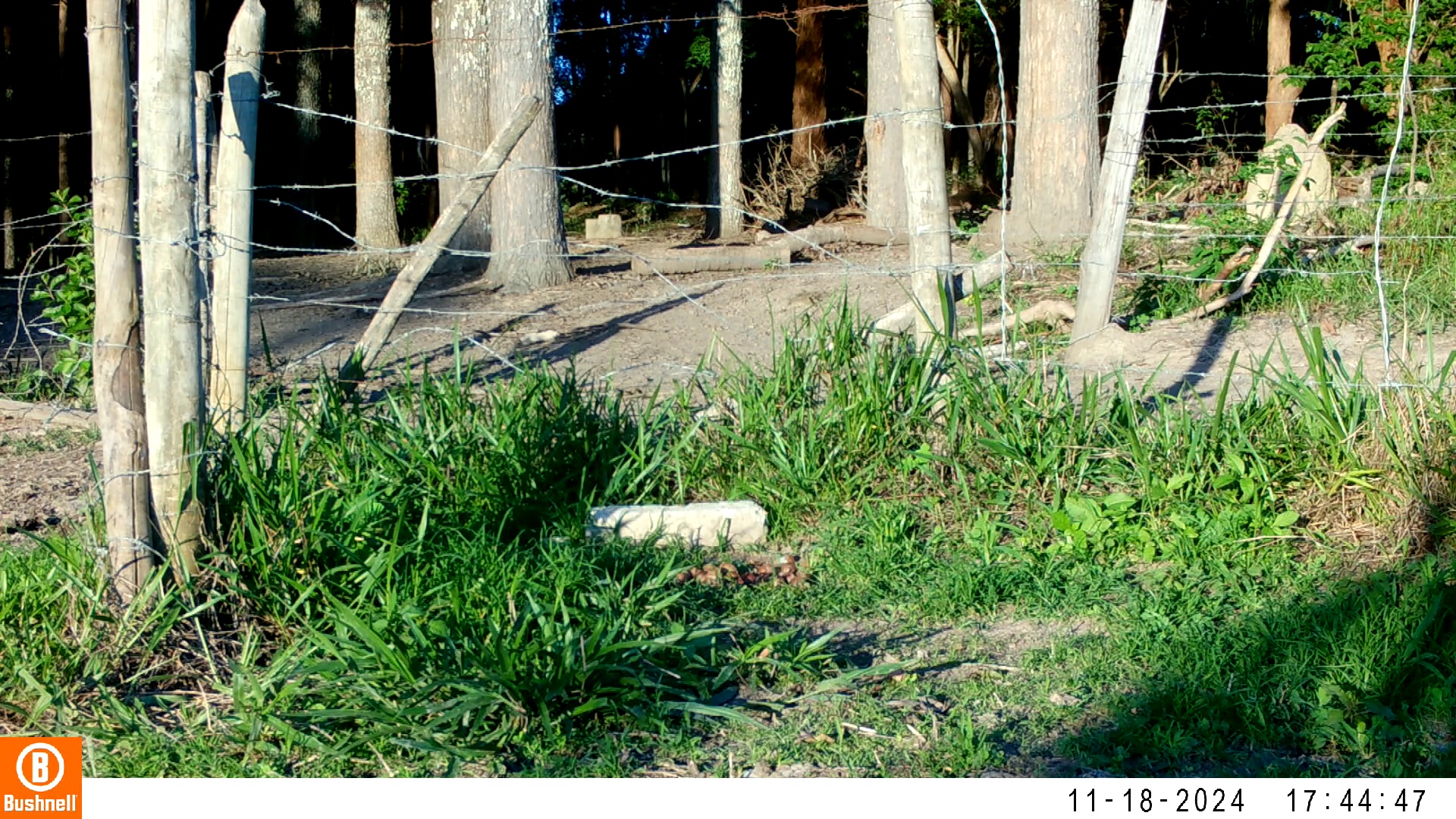

### 11180027-07363-1208.jpg

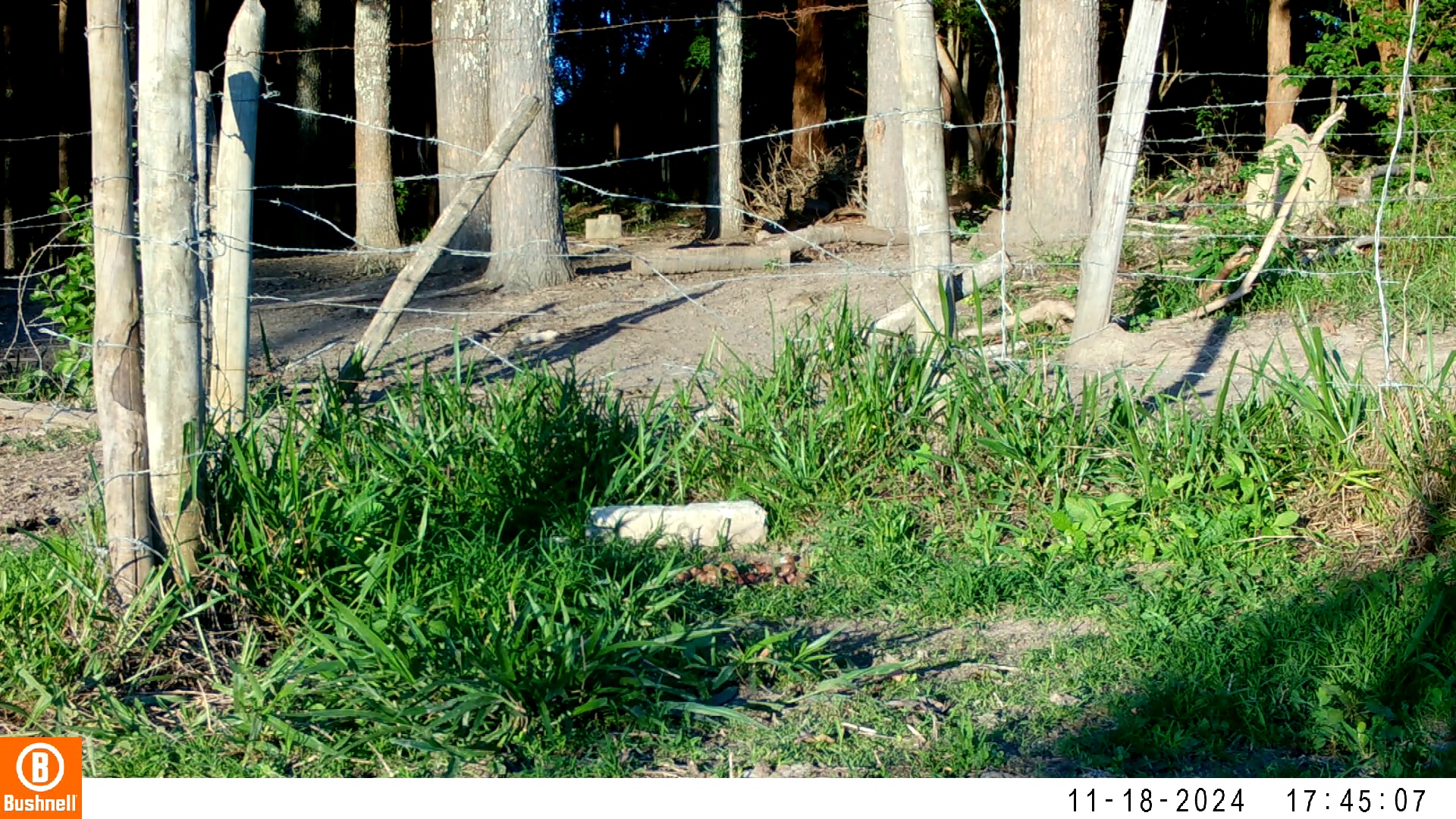

### 11180027-07363-1510.jpg

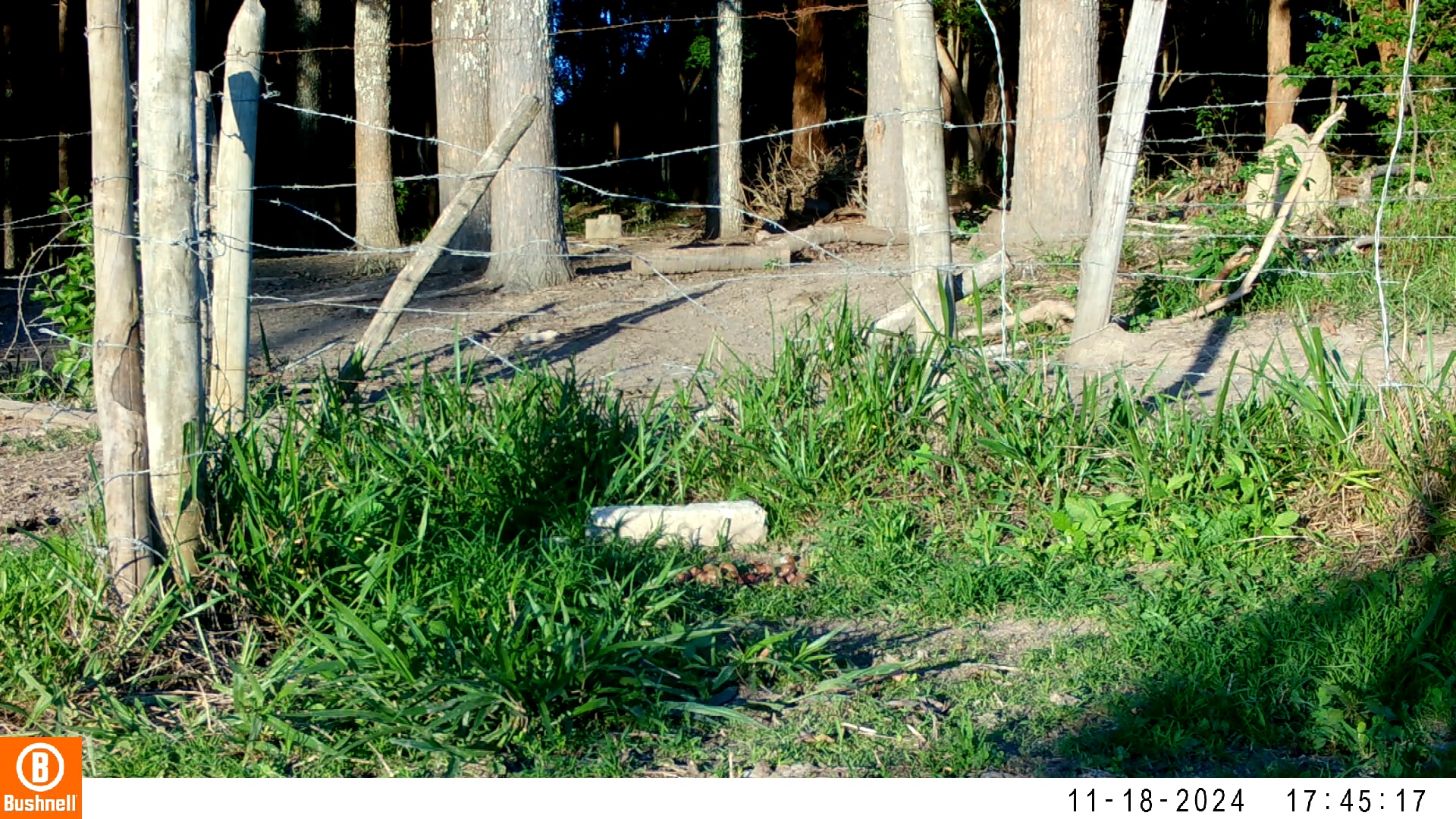

### 11190043-43338-302.jpg

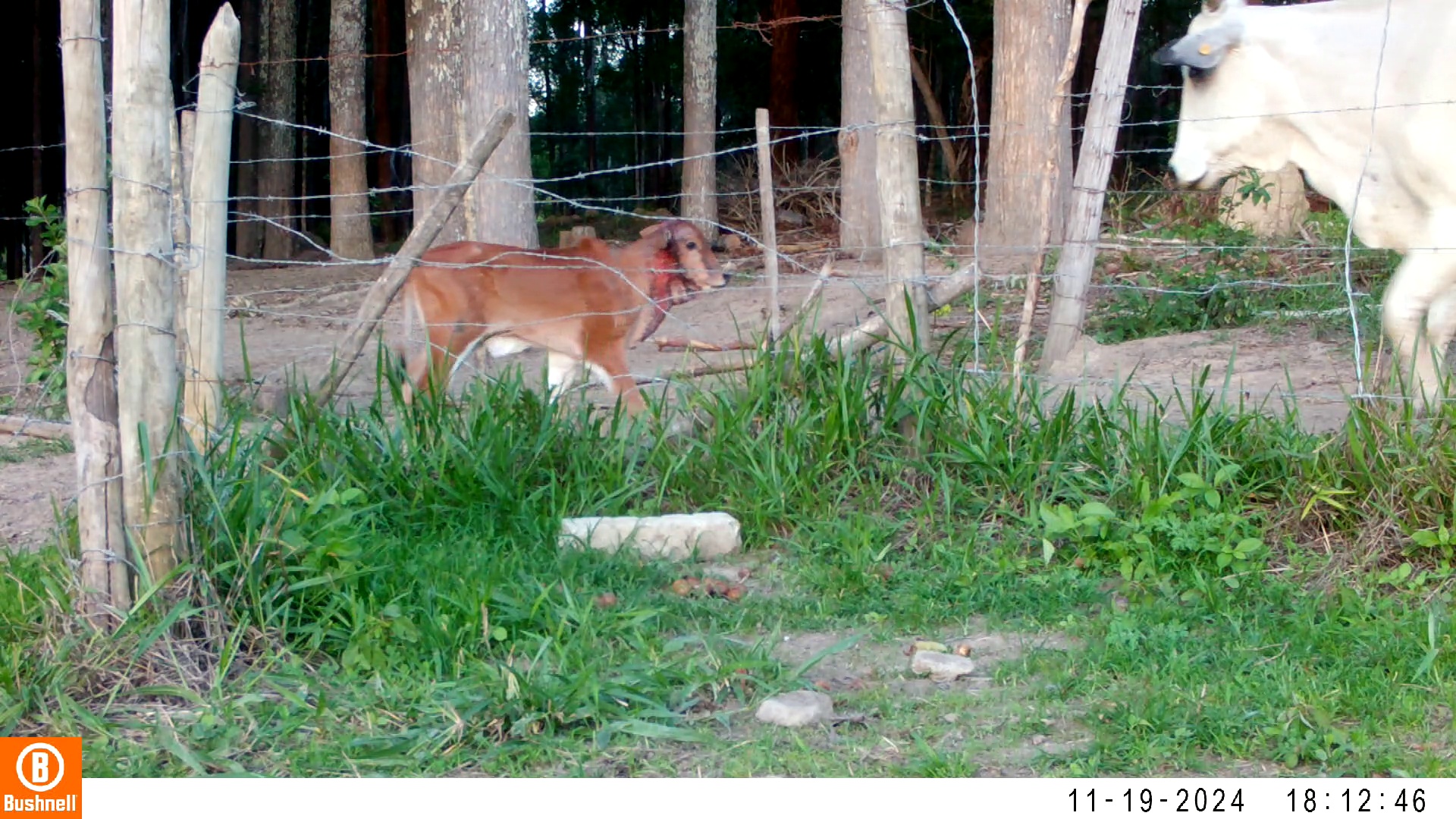

### 11190043-43338-604.jpg

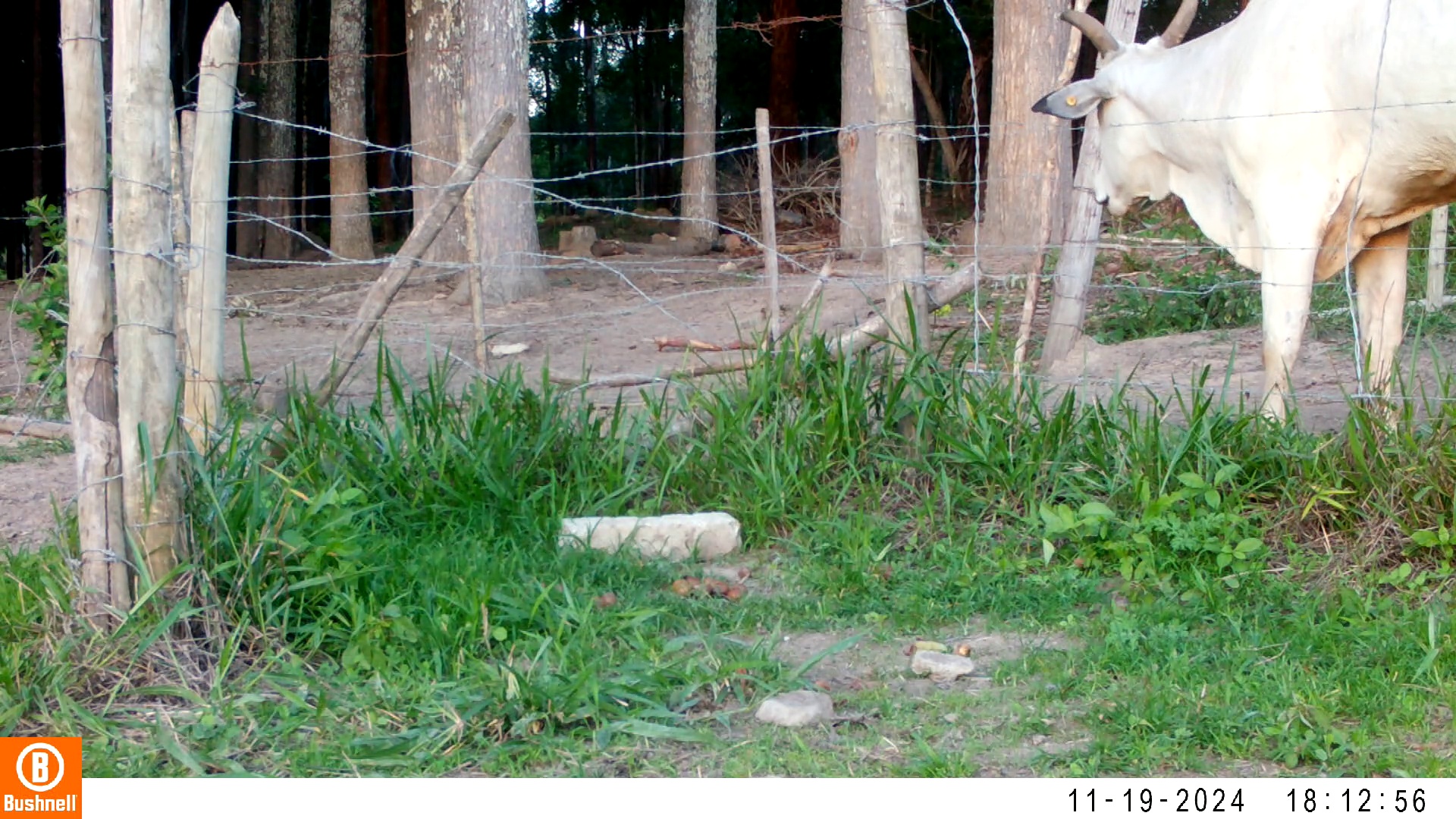

### 11190043-43338-906.jpg

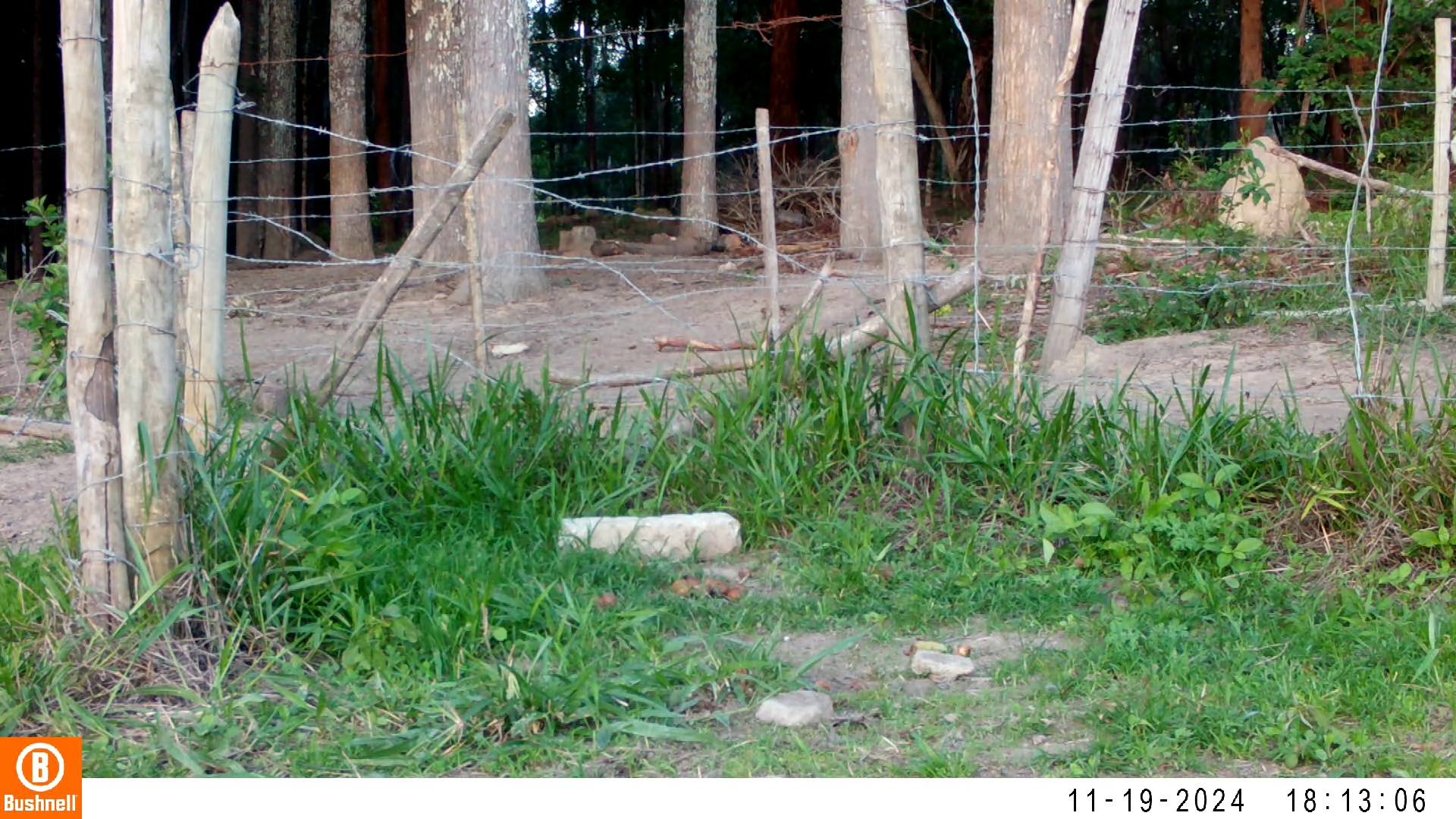

### 11190043-43338-1208.jpg

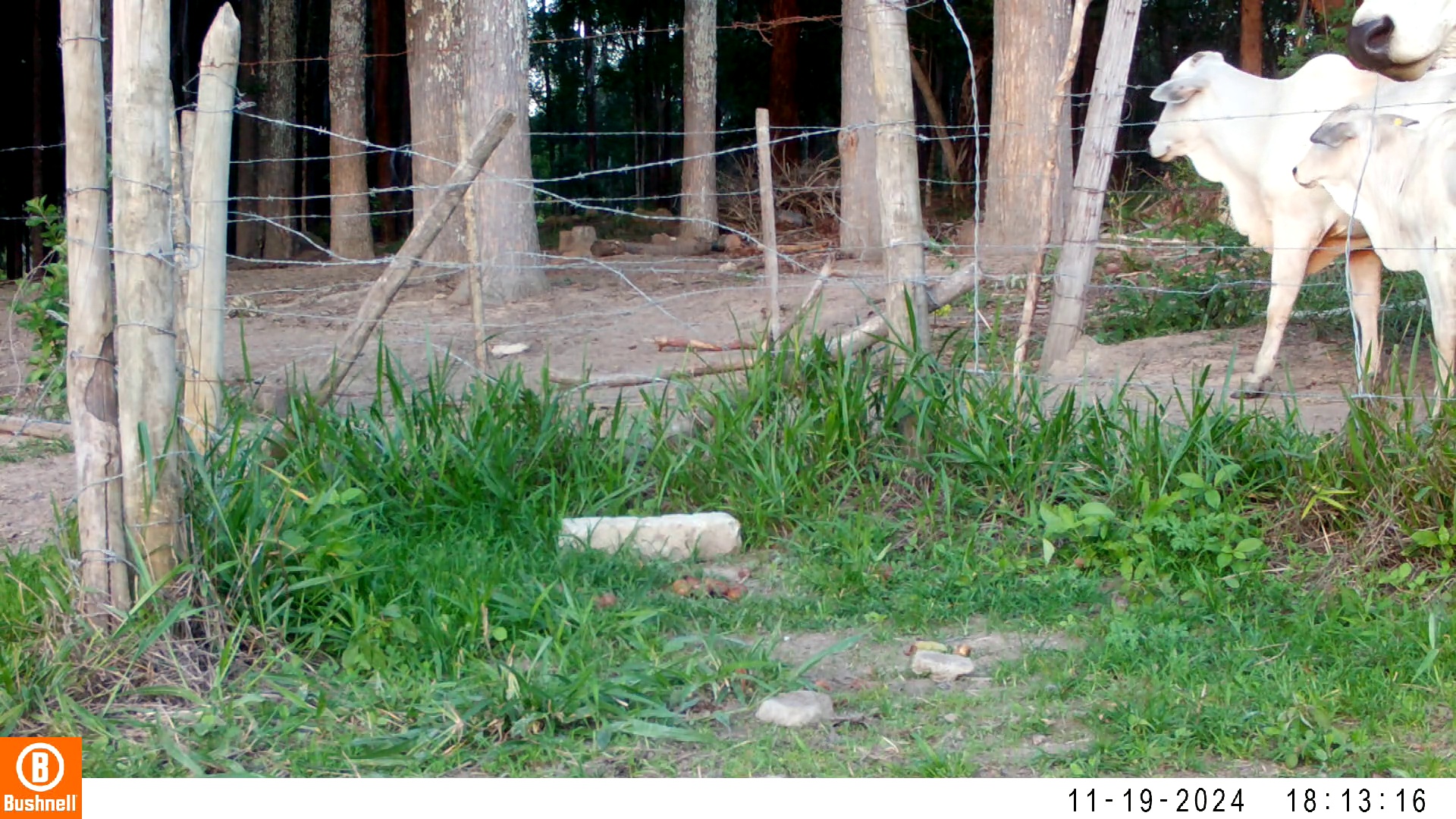

### 11190043-43338-1510.jpg

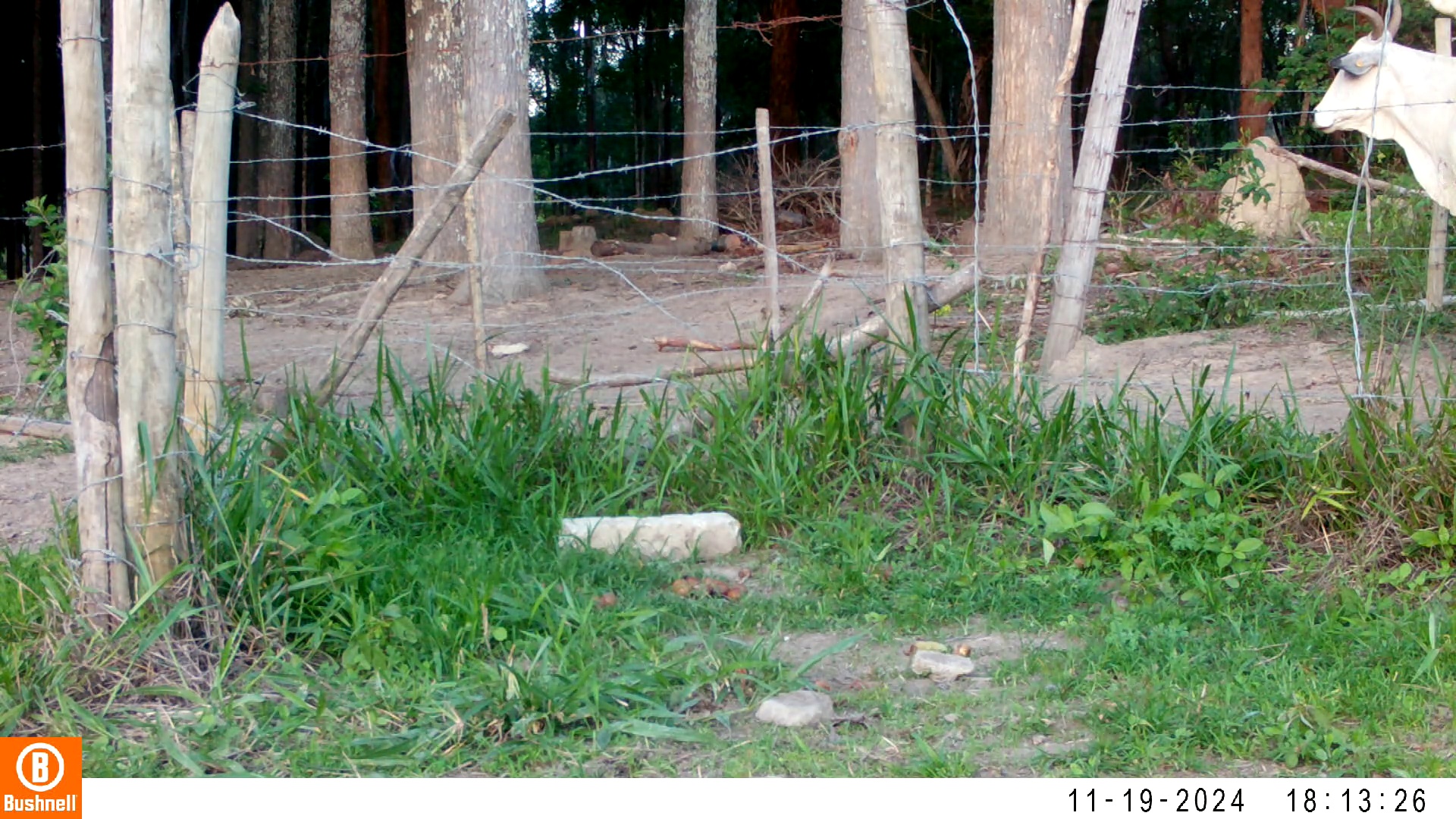

### 11190080-02410-1208.jpg

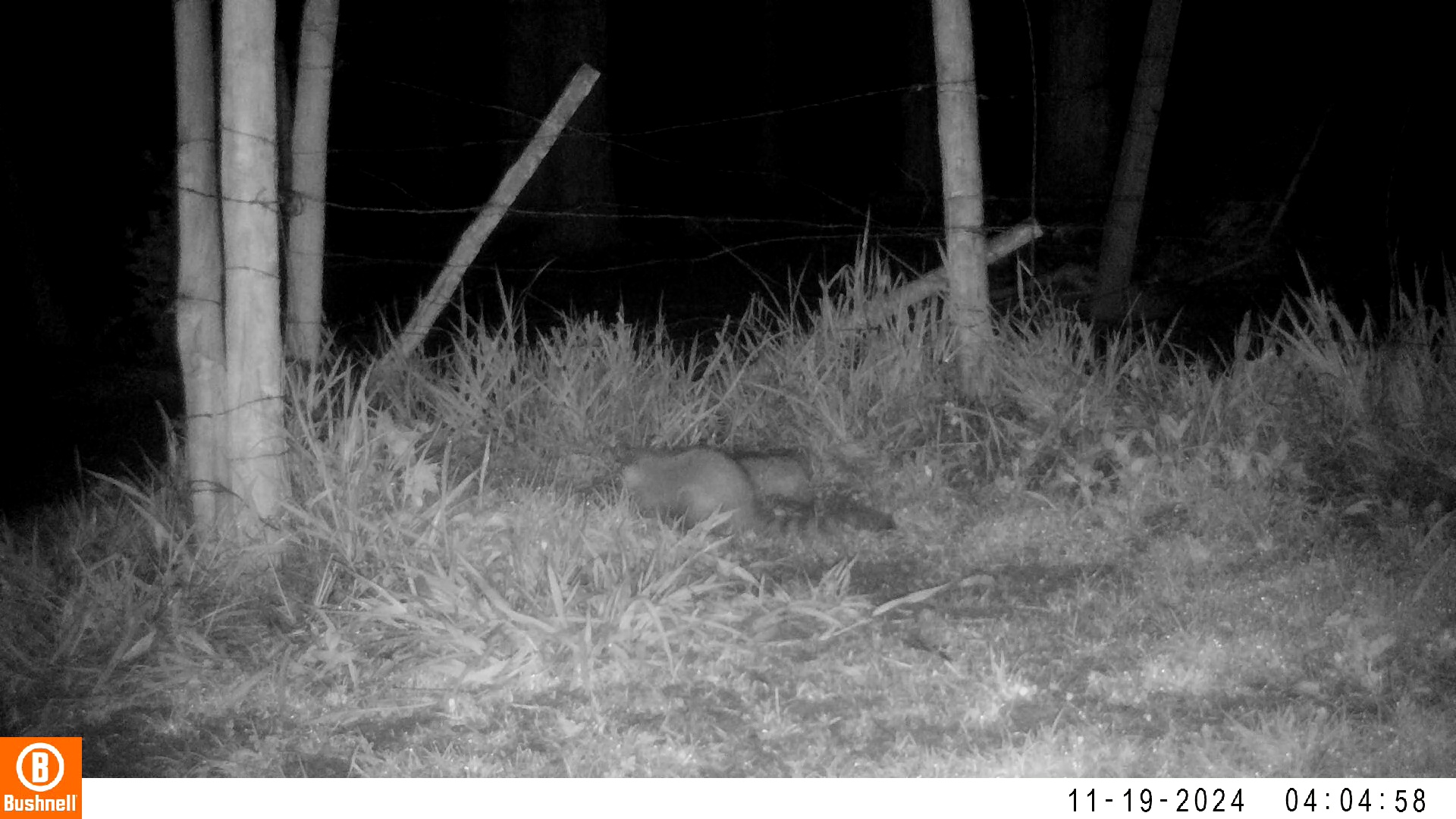

### 11190101-41263-0.jpg

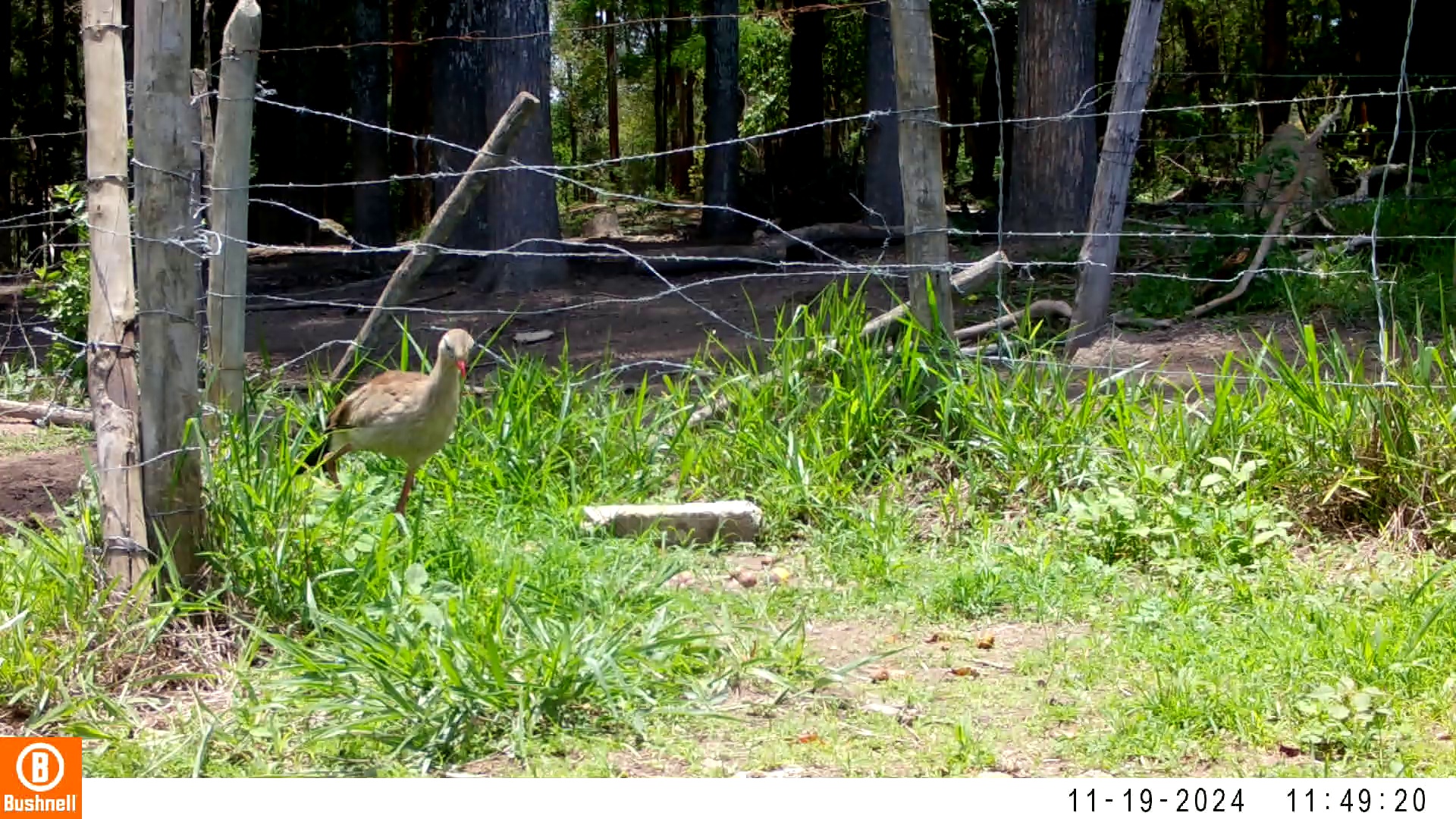

### 11190101-41263-1208.jpg

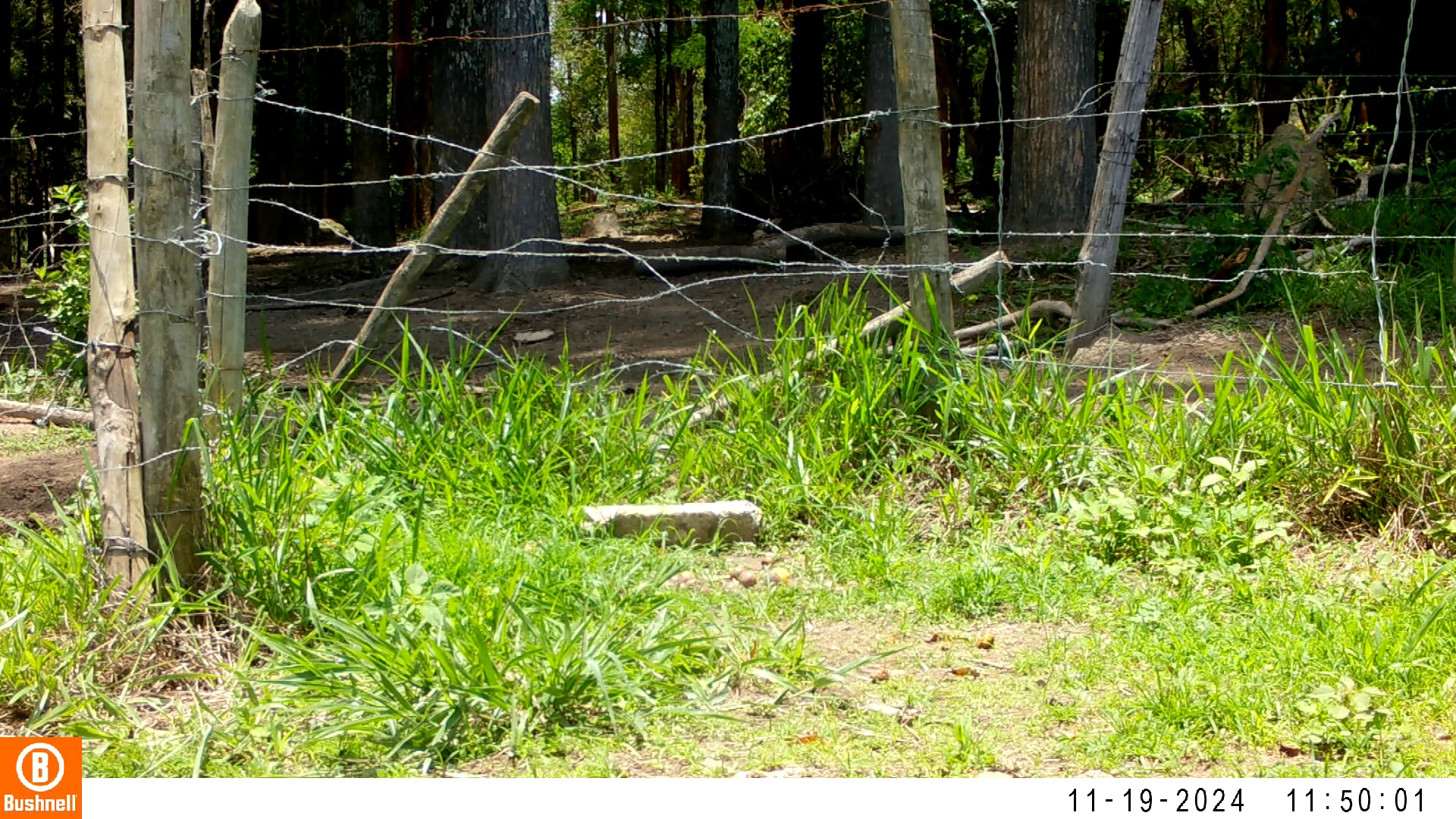
