## Supplementary figures and images for "Trade-off between foraging and vigilance in the Red-legged Seriema"

### 1280-04209-600.jpg

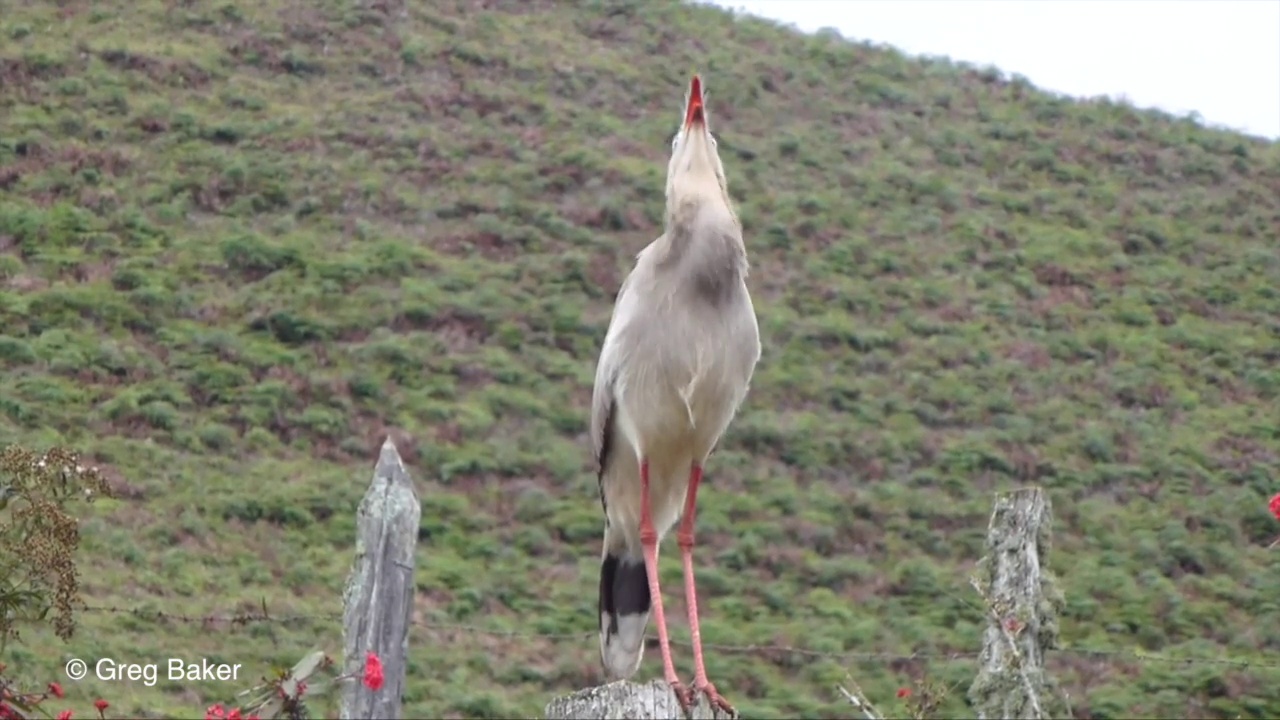

### 01130007-74058-0.jpg

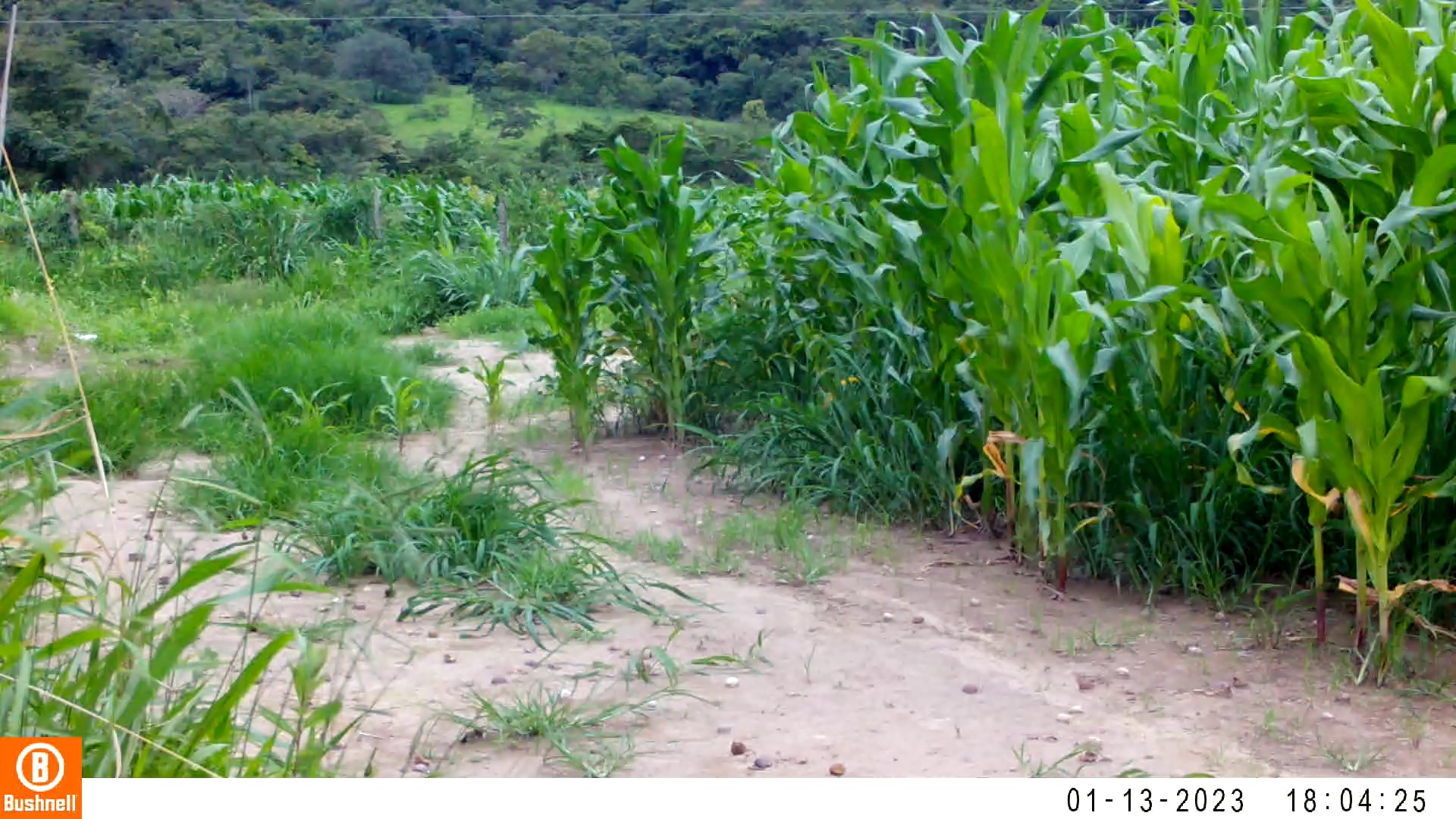

### 01130007-74058-202.jpg

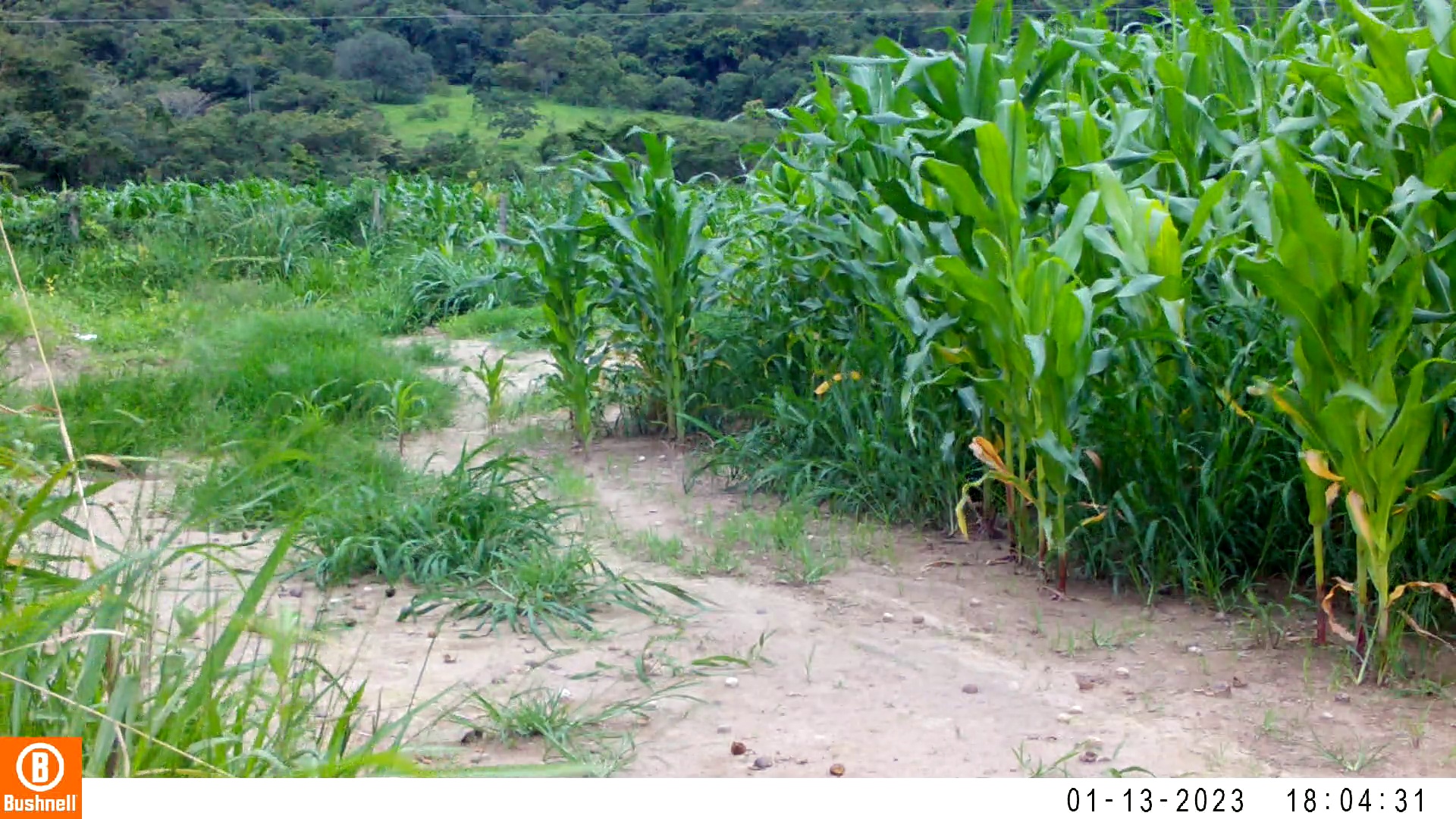

### 01130007-74058-404.jpg

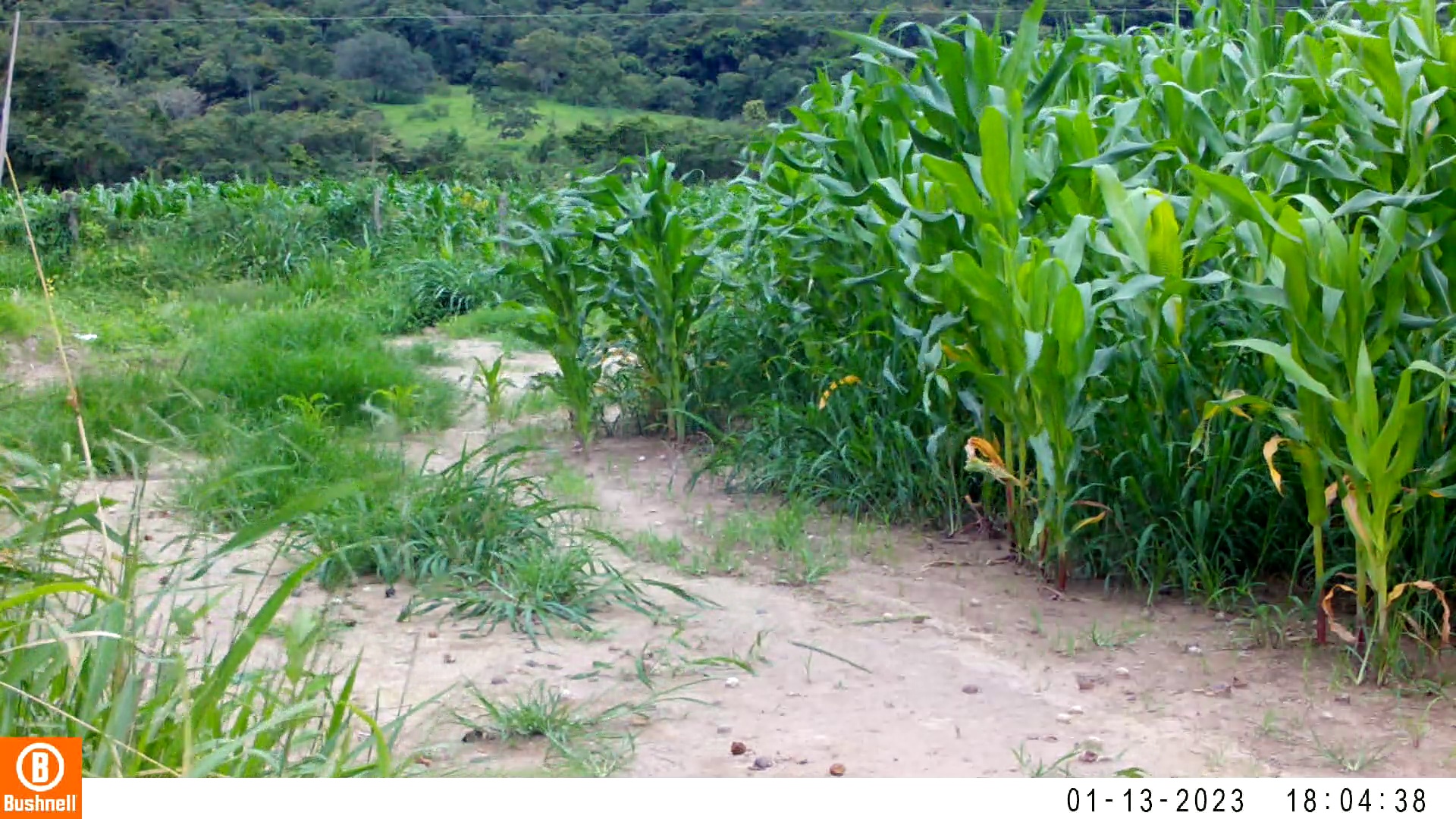

### 01130007-74058-606.jpg

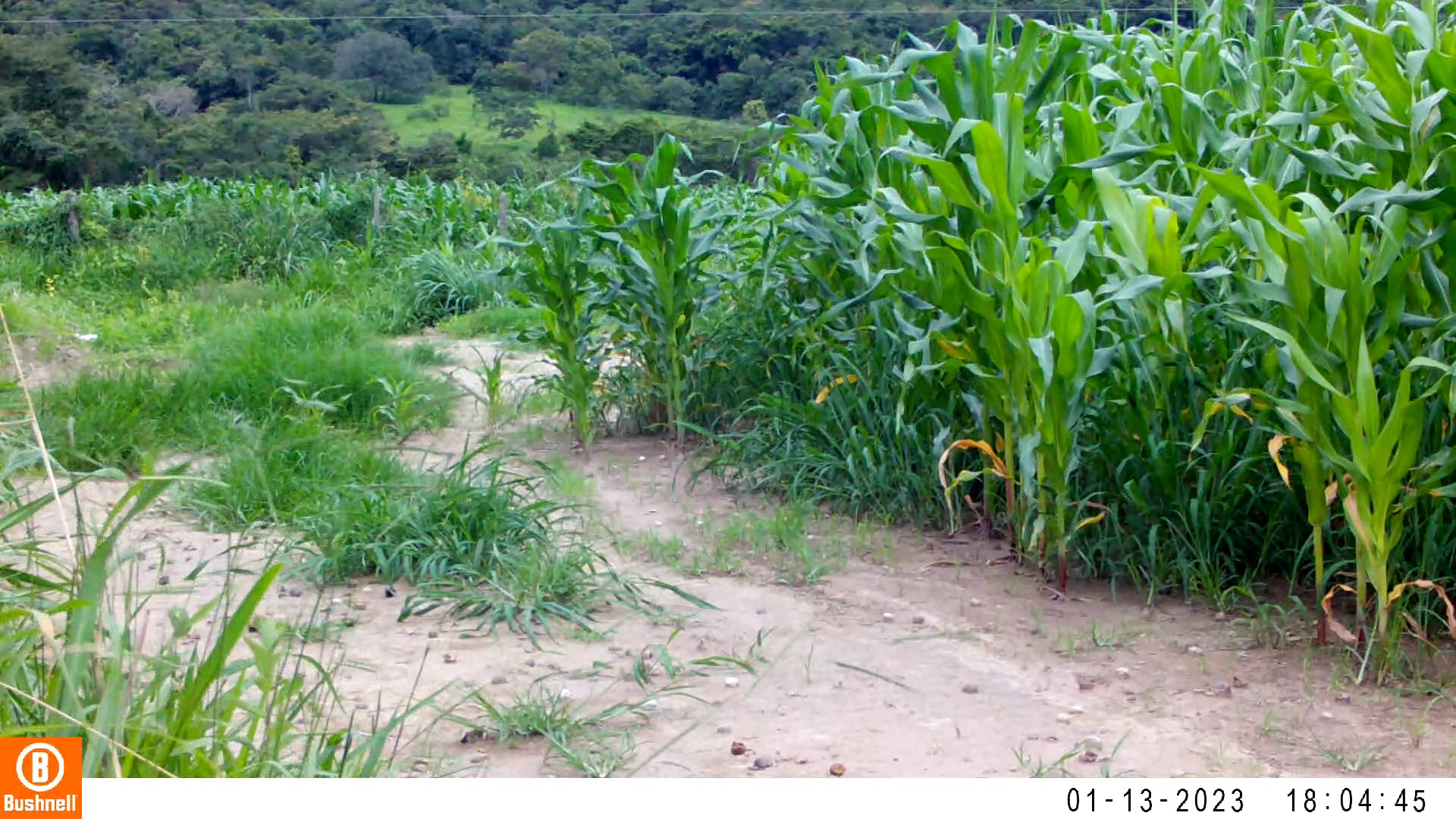

### 01130007-74058-808.jpg

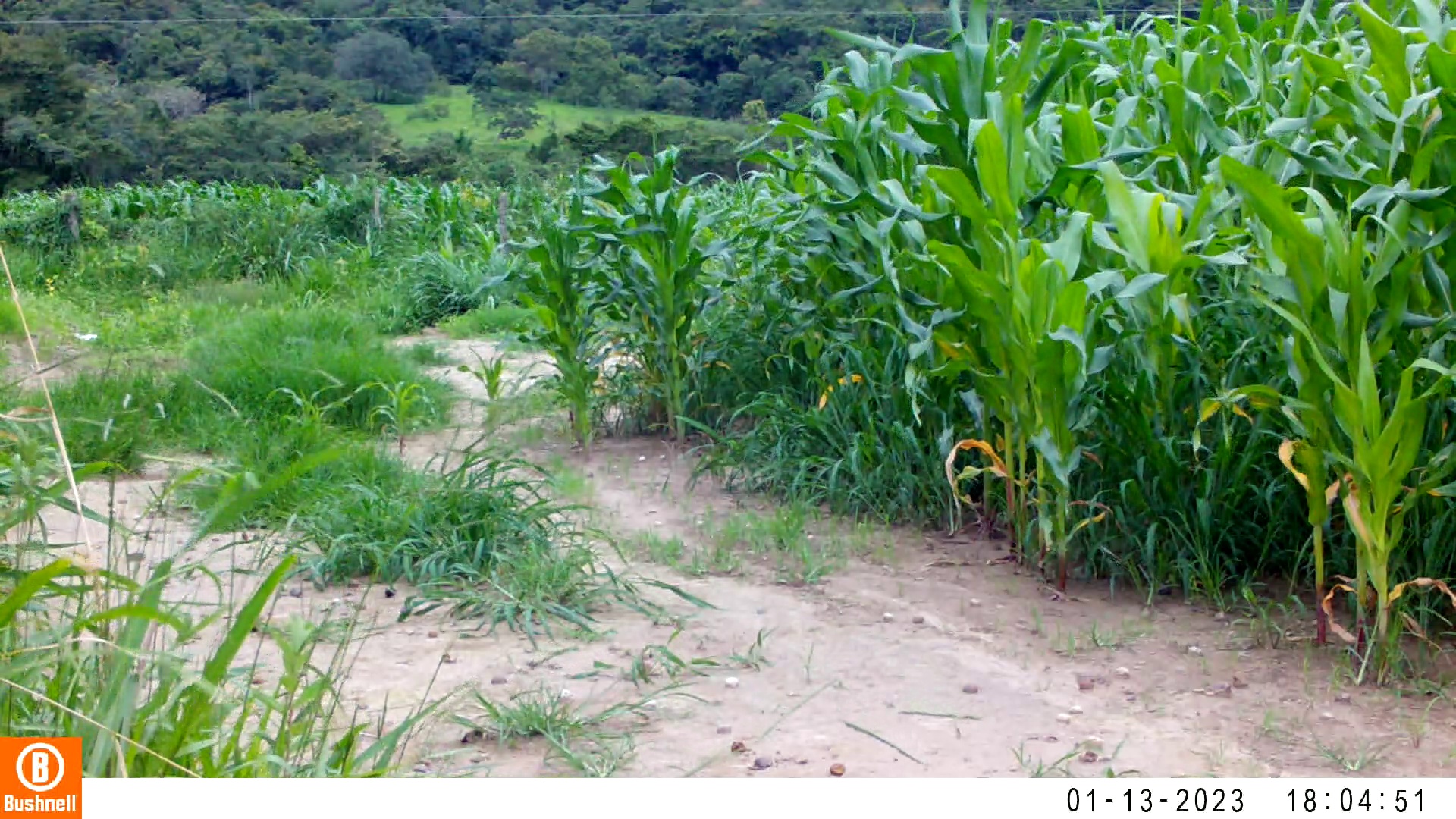

### 01130007-74058-1010.jpg

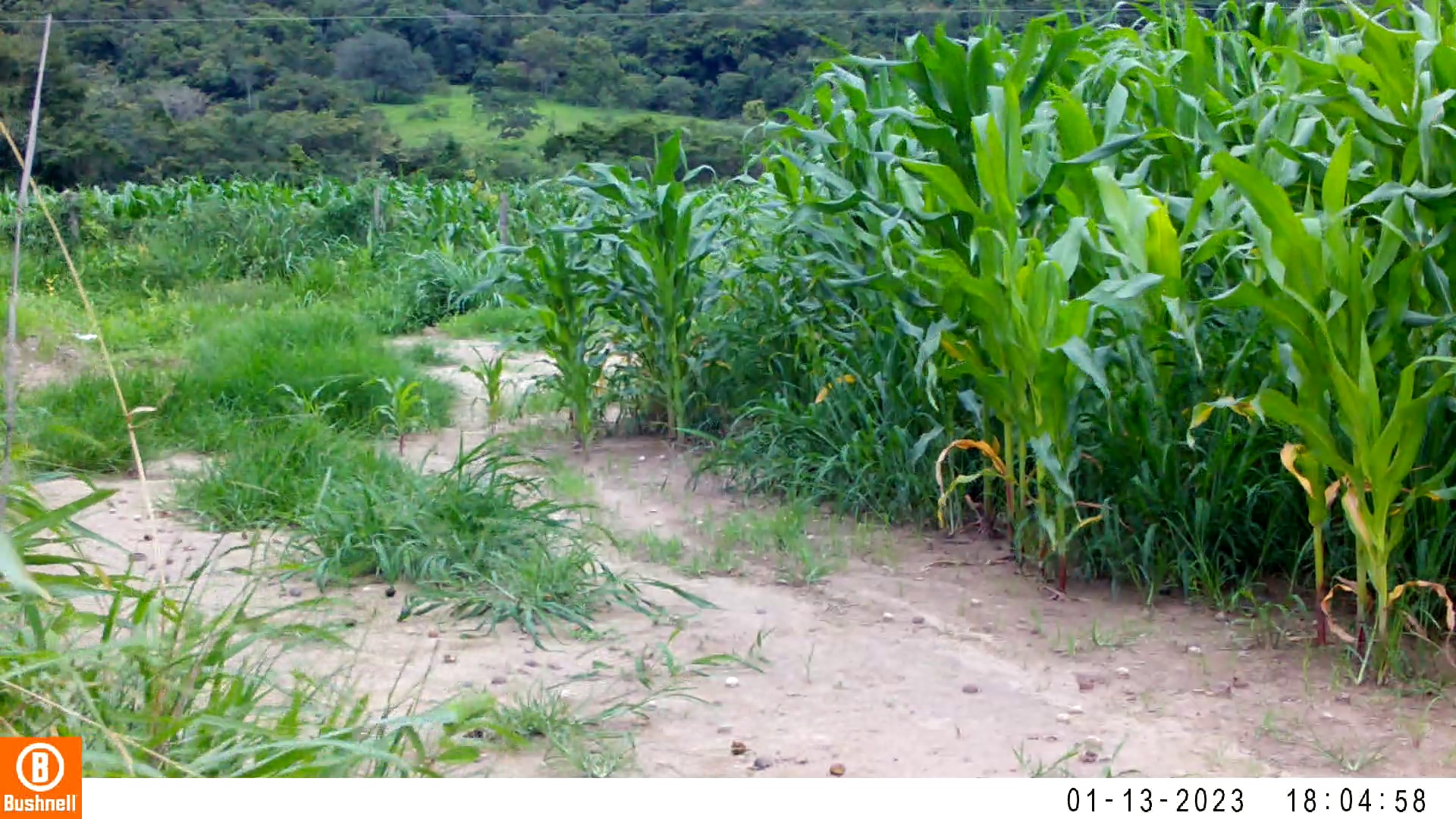

### 01140009-76782-0.jpg

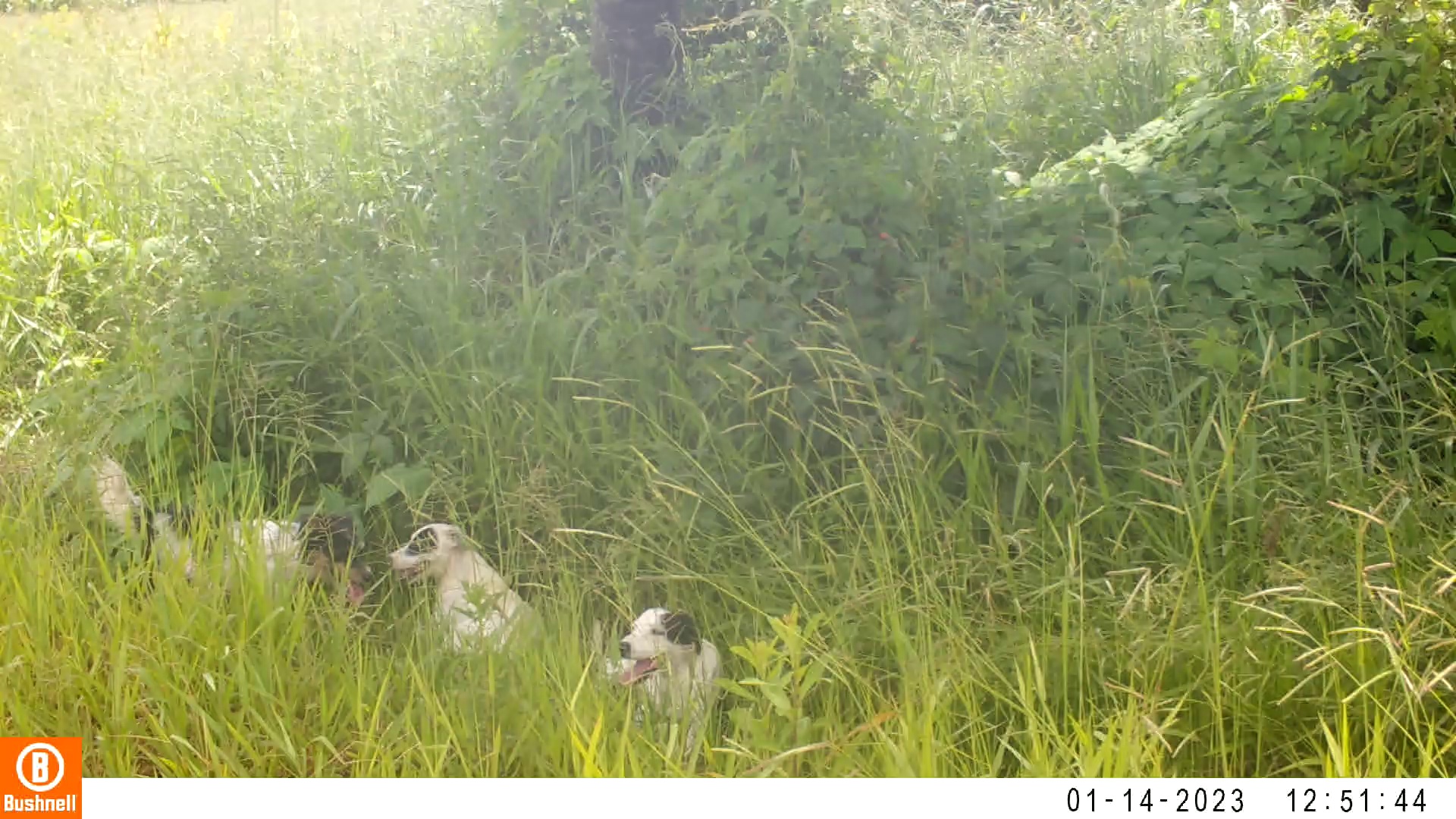

### 01140009-76782-152.jpg

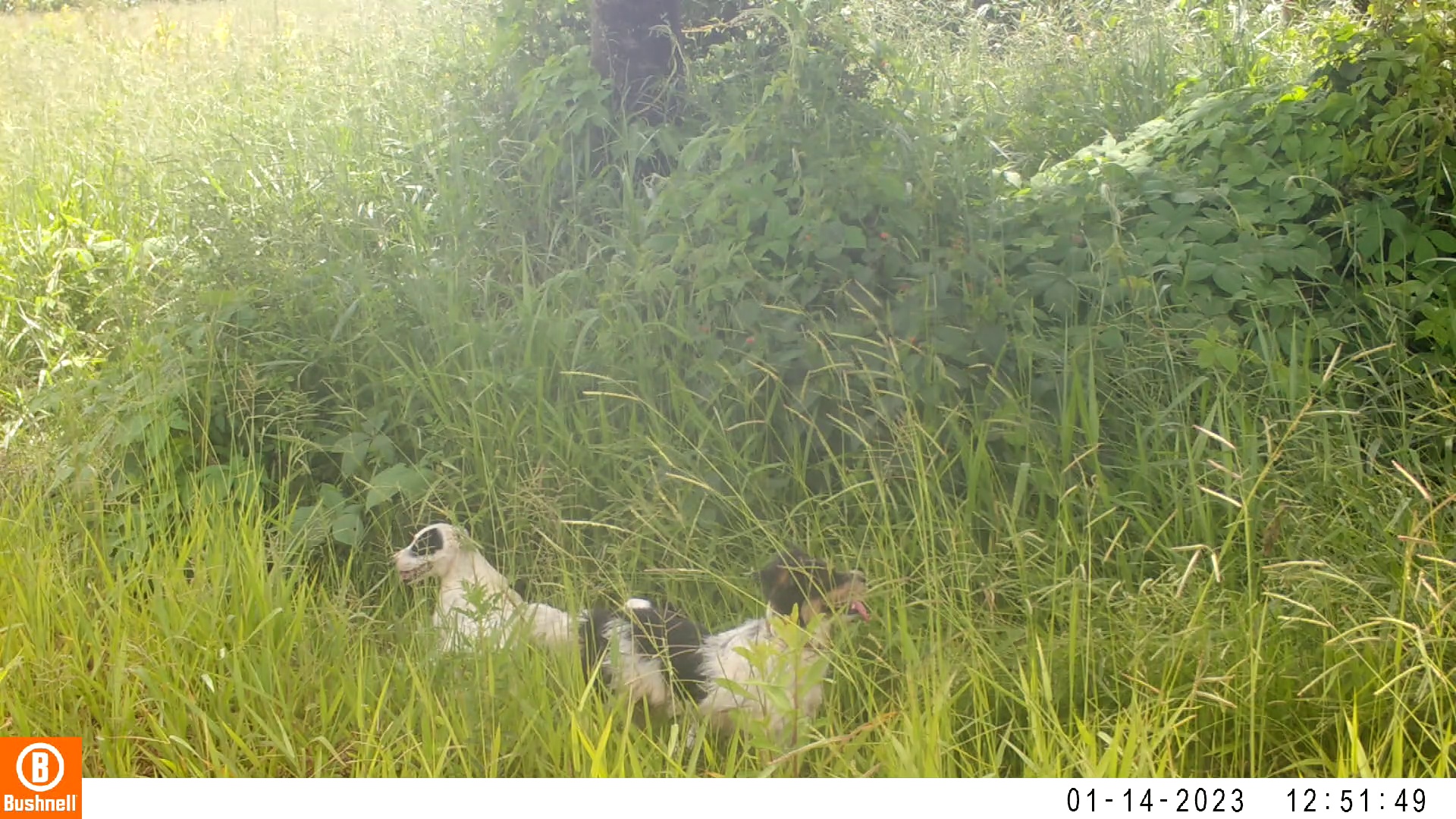

### 01140009-76782-304.jpg

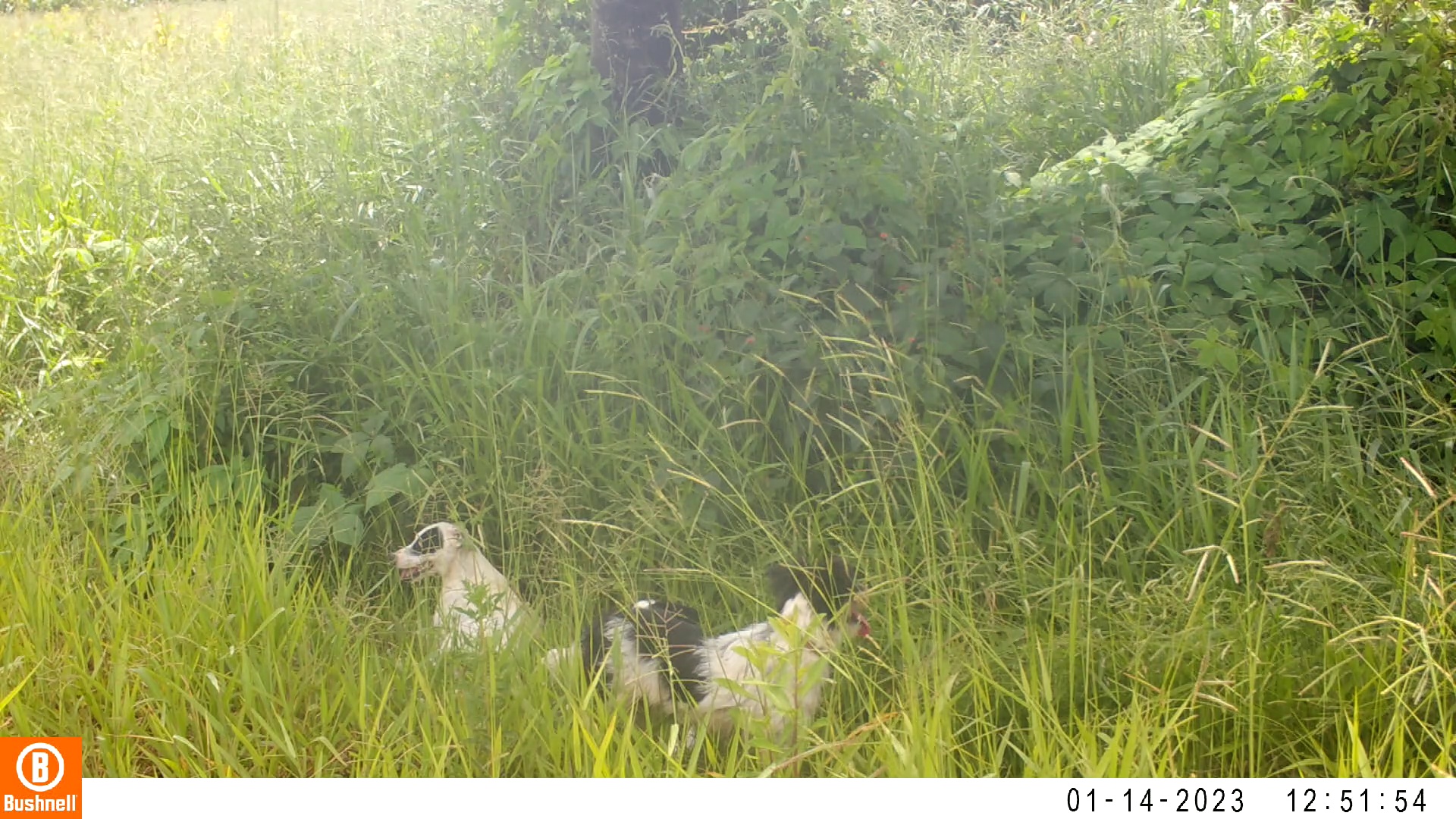

### 01140009-76782-456.jpg

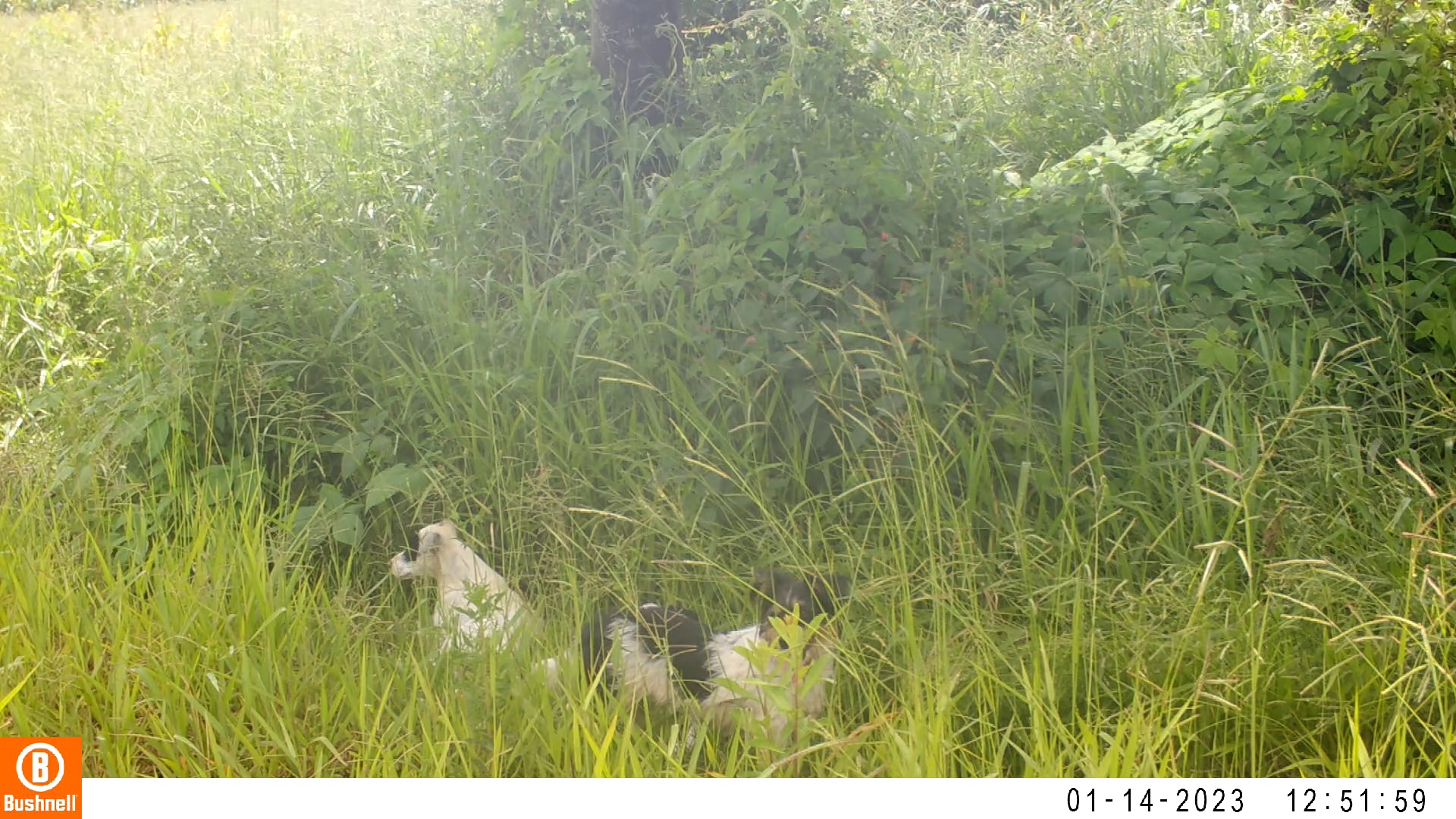

### 01140009-76782-608.jpg

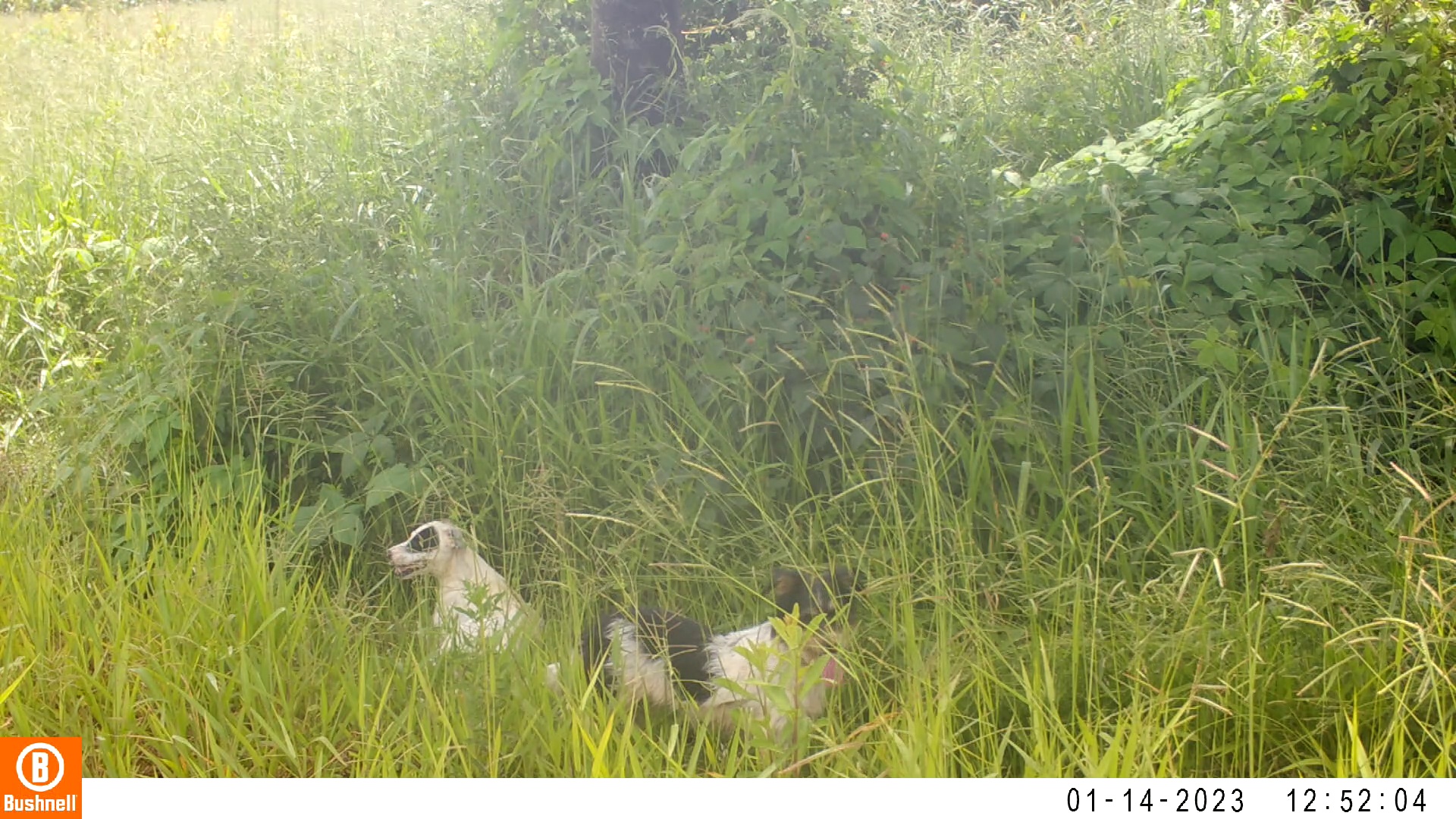

### 01140009-76782-760.jpg

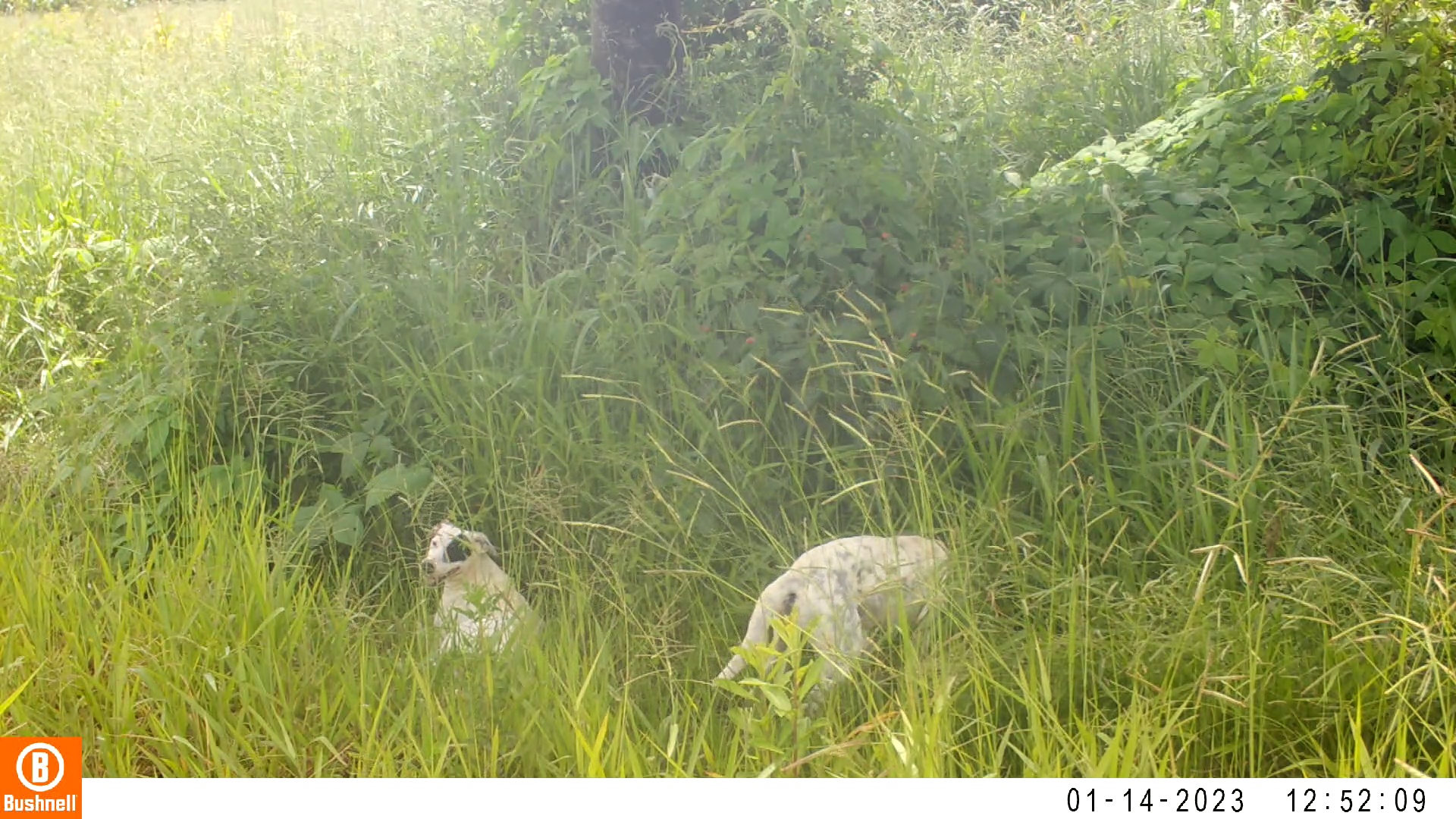

### 01140021-02265-0.jpg

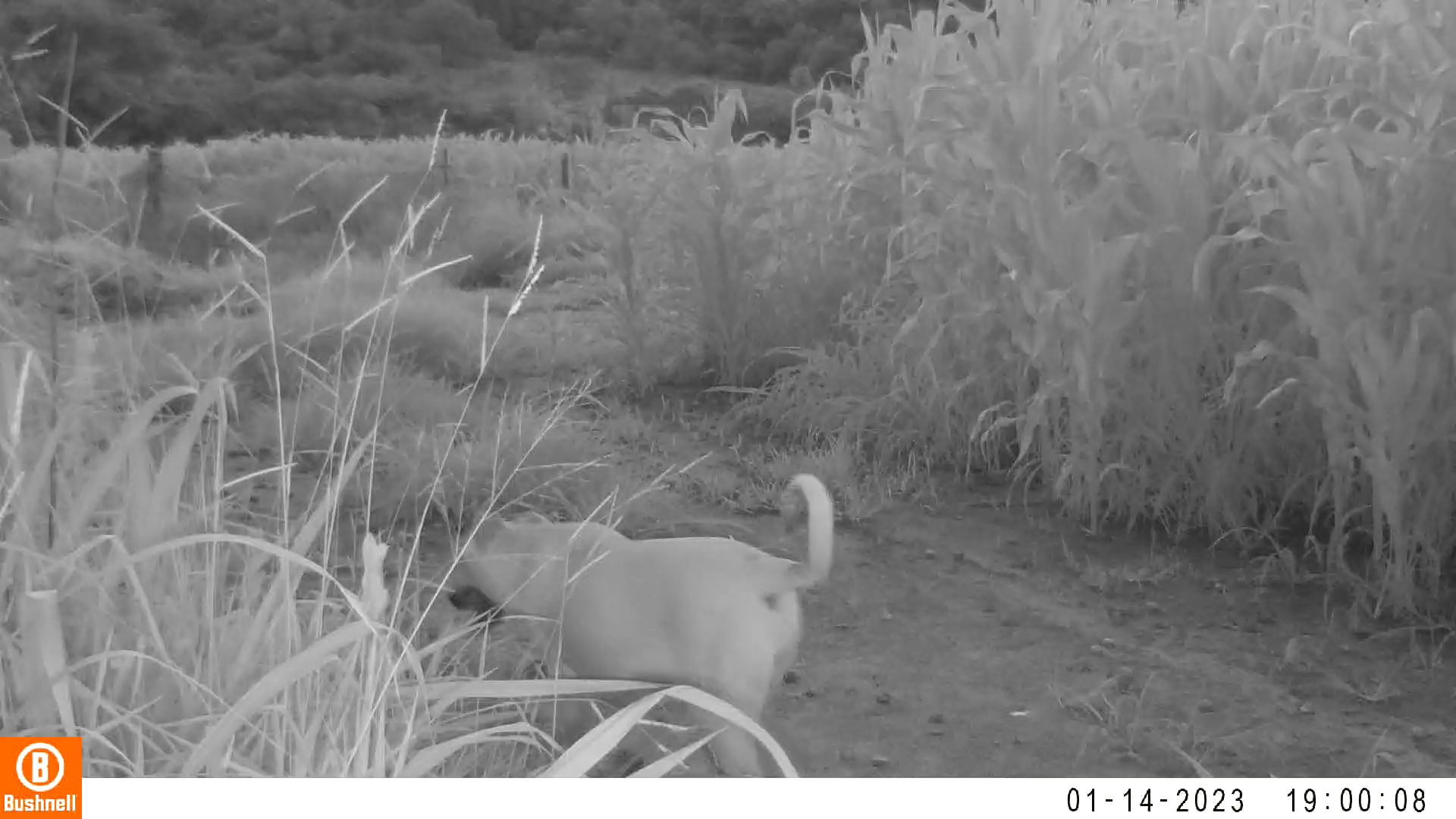

### 01140021-02265-154.jpg

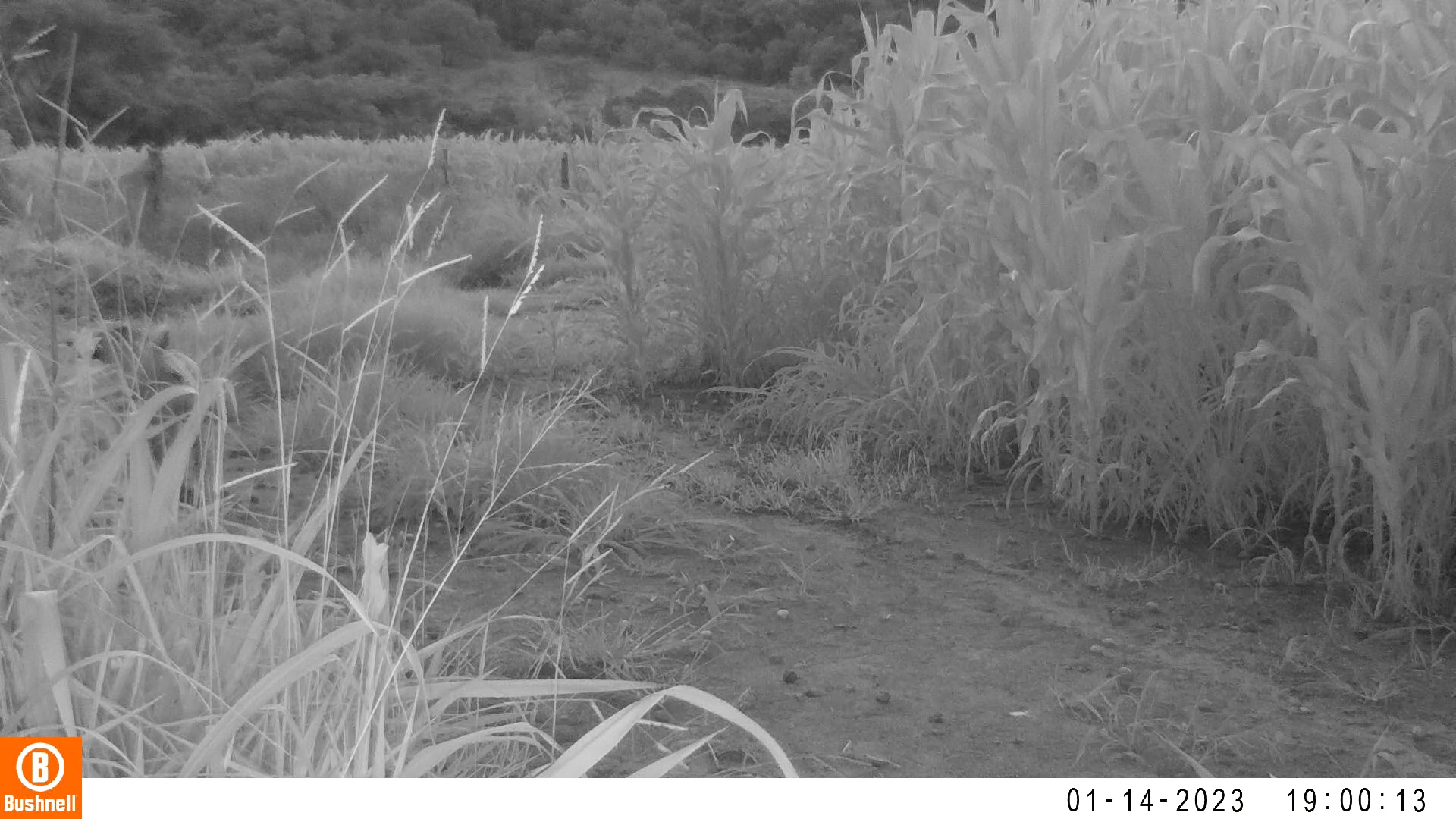

### 01140021-02265-231.jpg

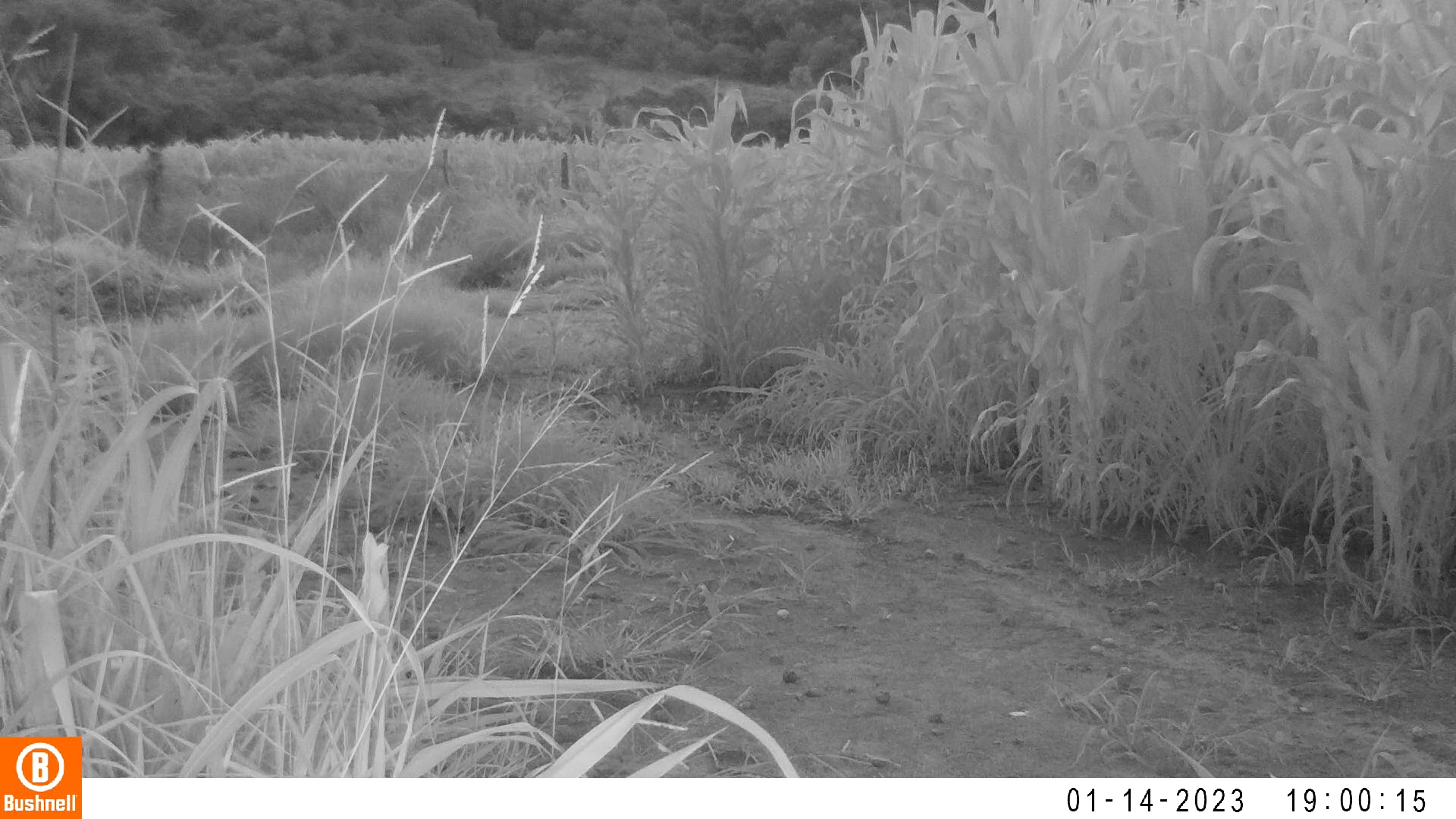

### 01140021-02265-308.jpg

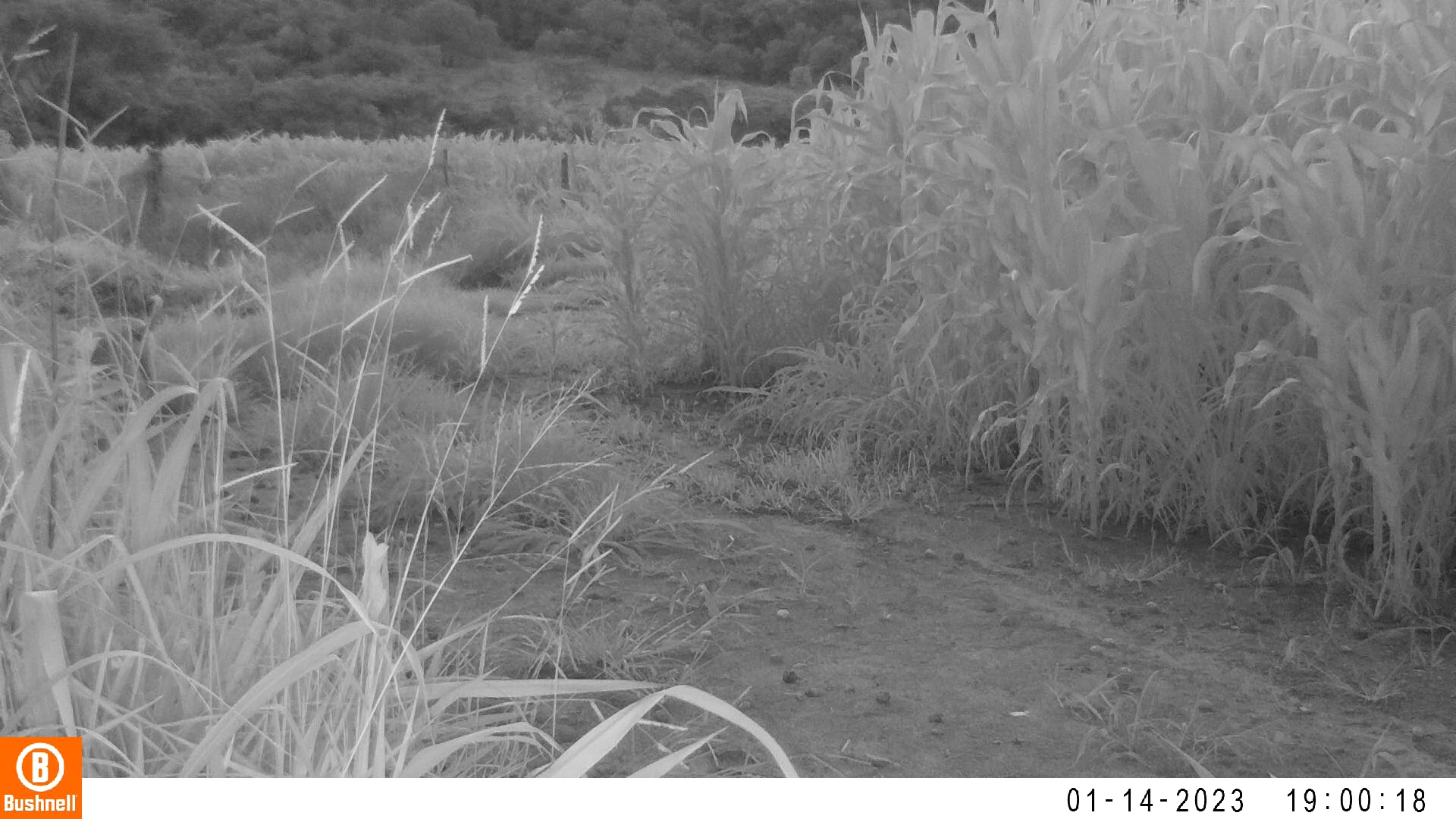

### 01140021-02265-385.jpg

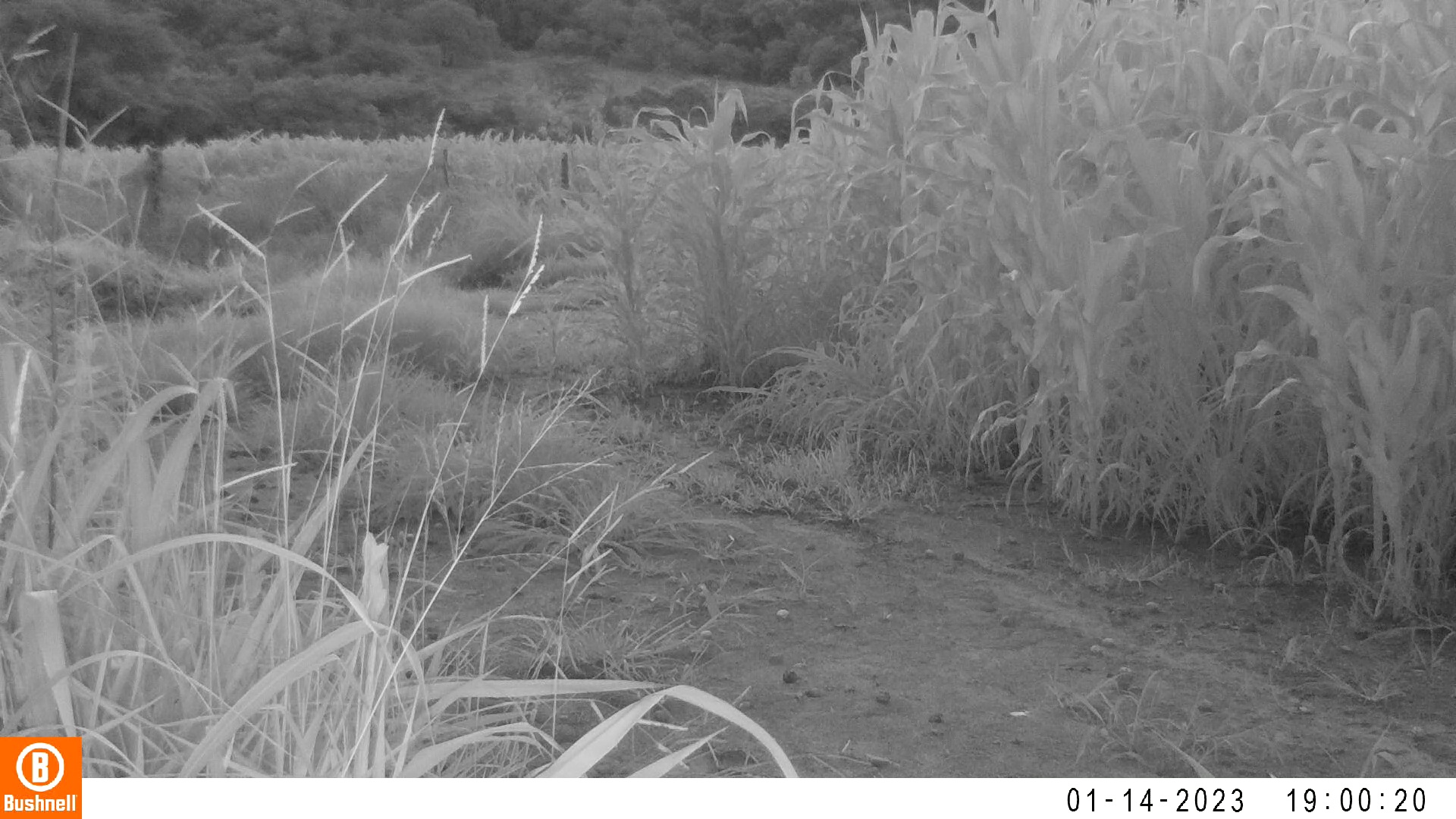

### 01150016-02464-0.jpg

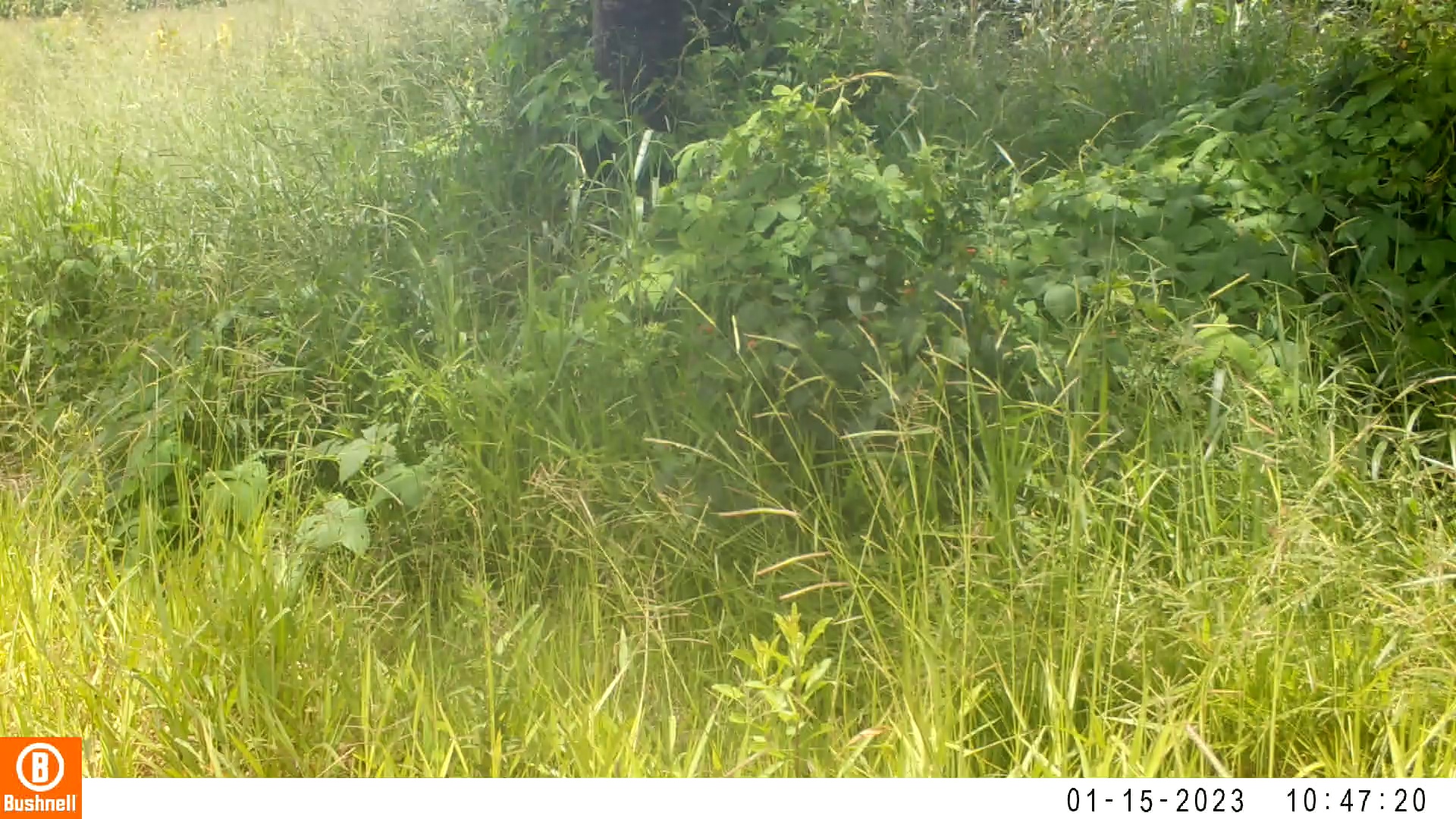

### 01150016-02464-152.jpg

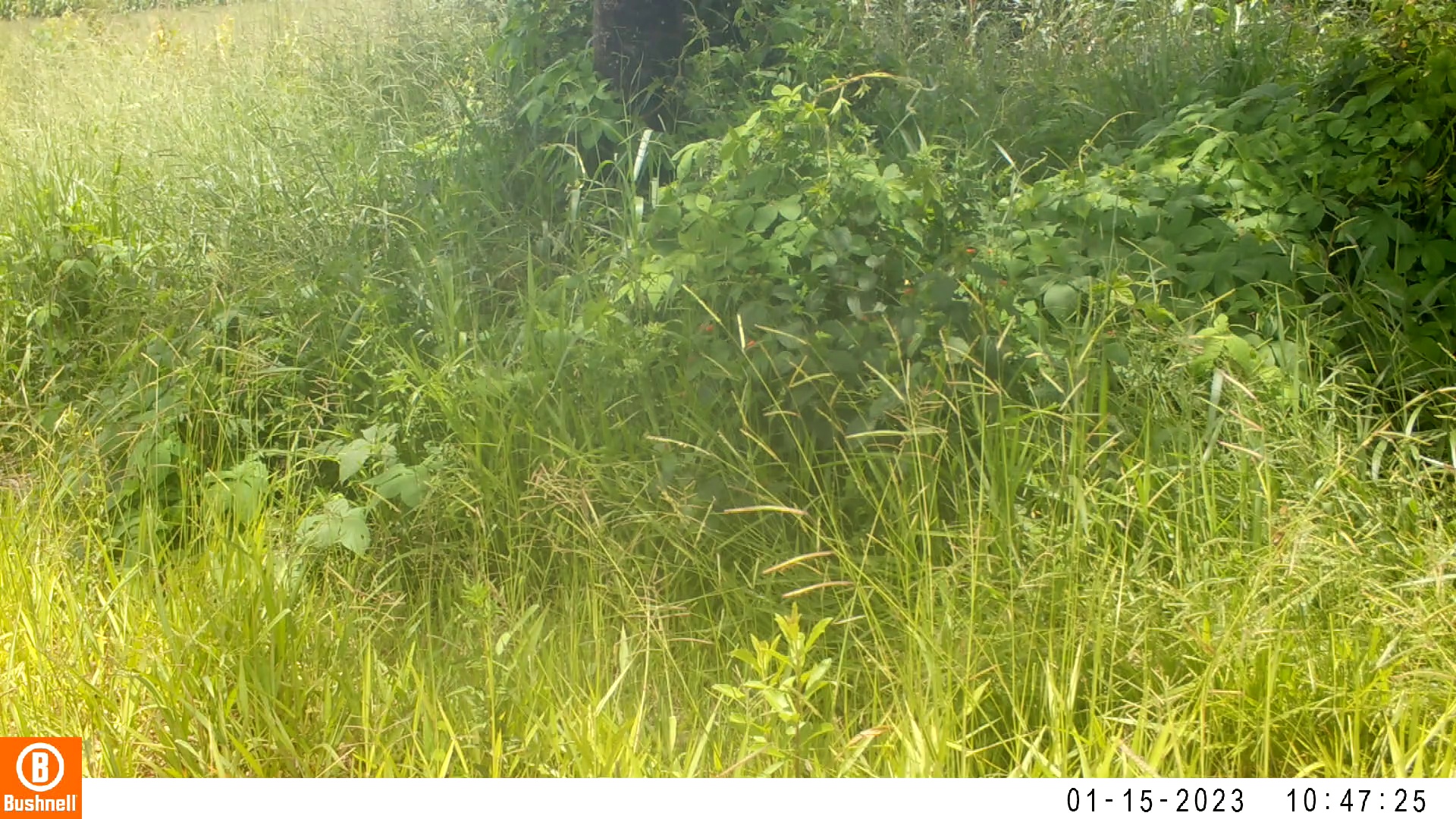

### 01150016-02464-304.jpg

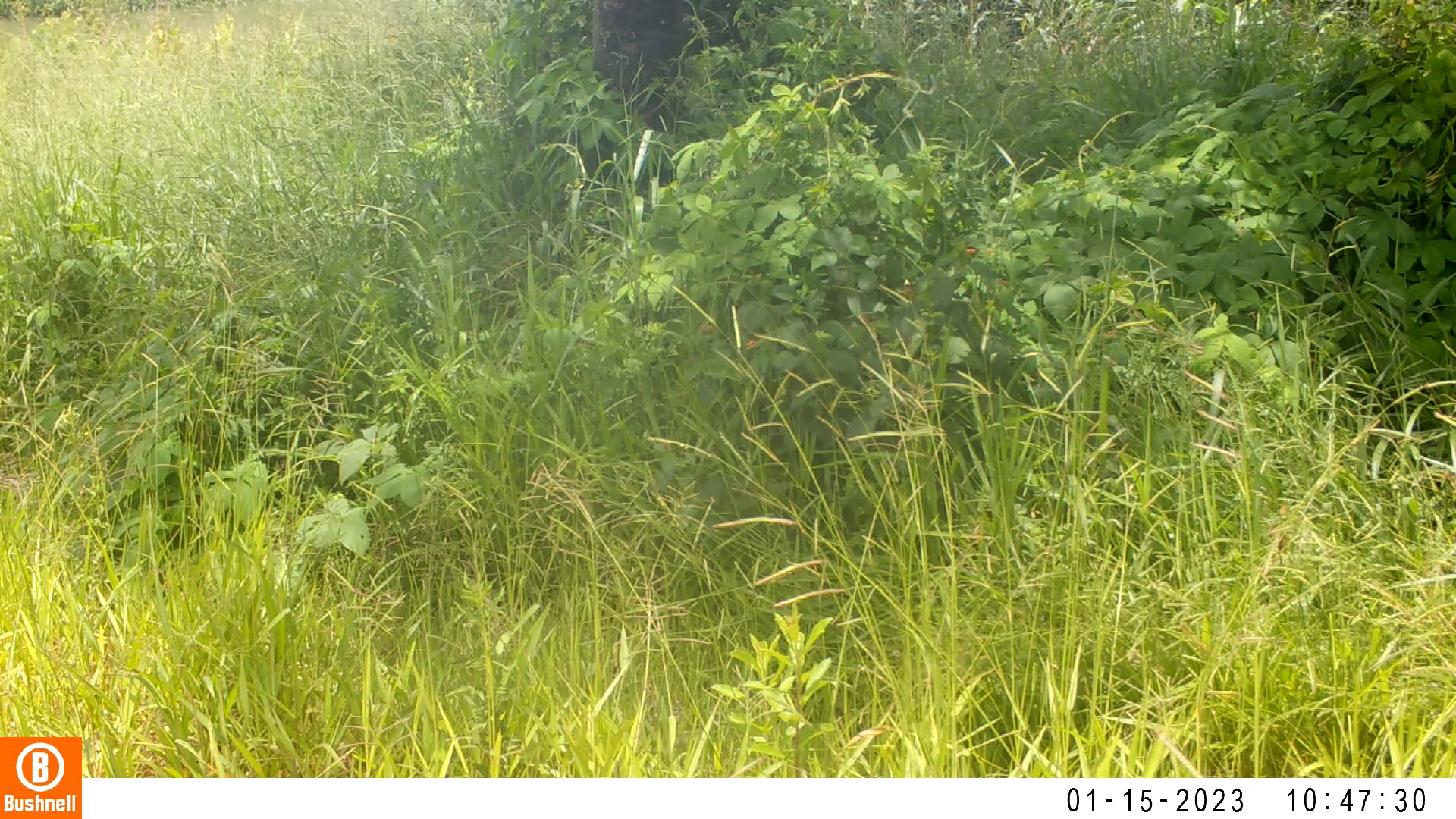

### 01150016-02464-456.jpg

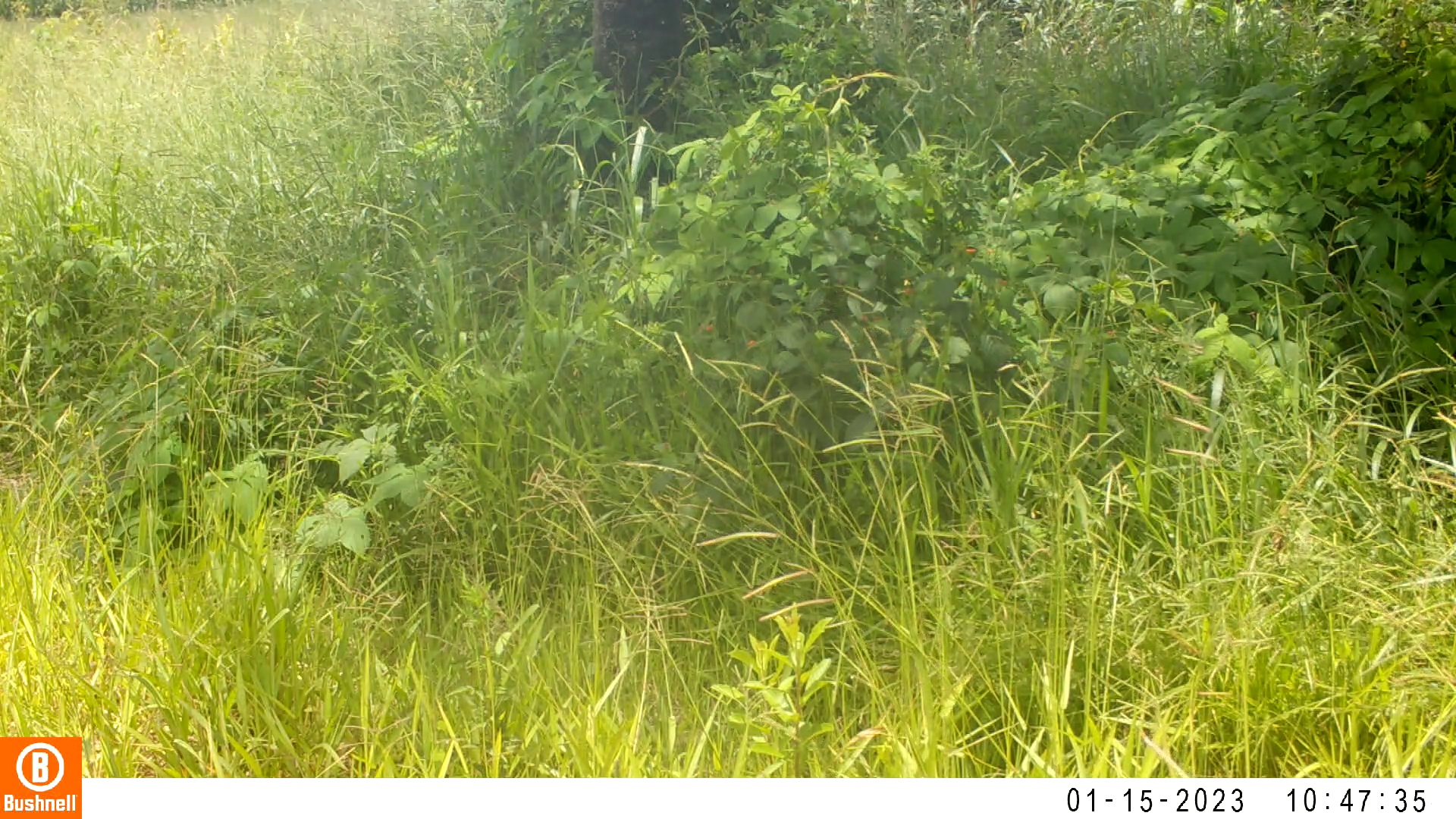

### 01150016-02464-608.jpg

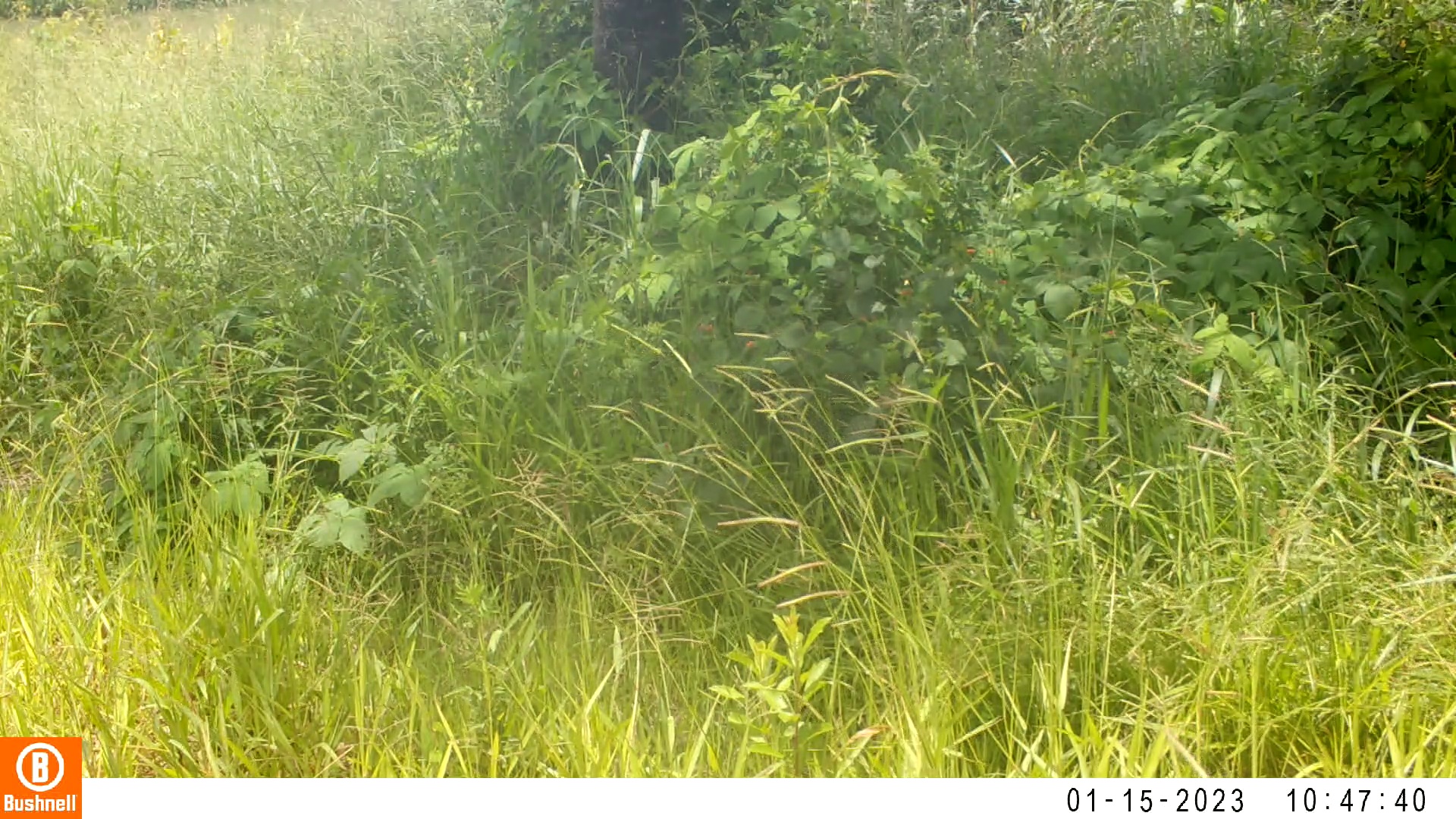

### 01150016-02464-760.jpg

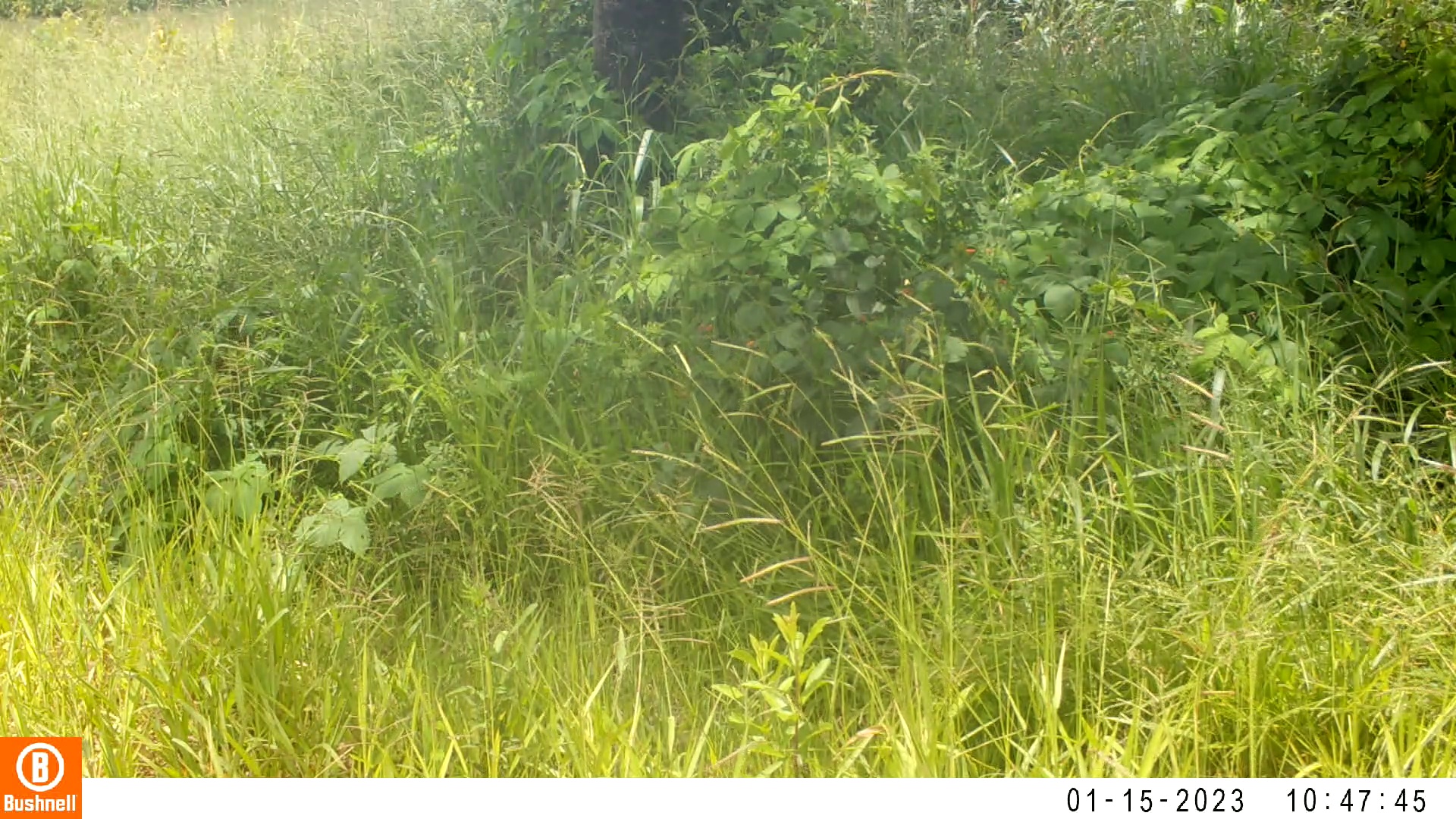

### 01150027-01940-0.jpg

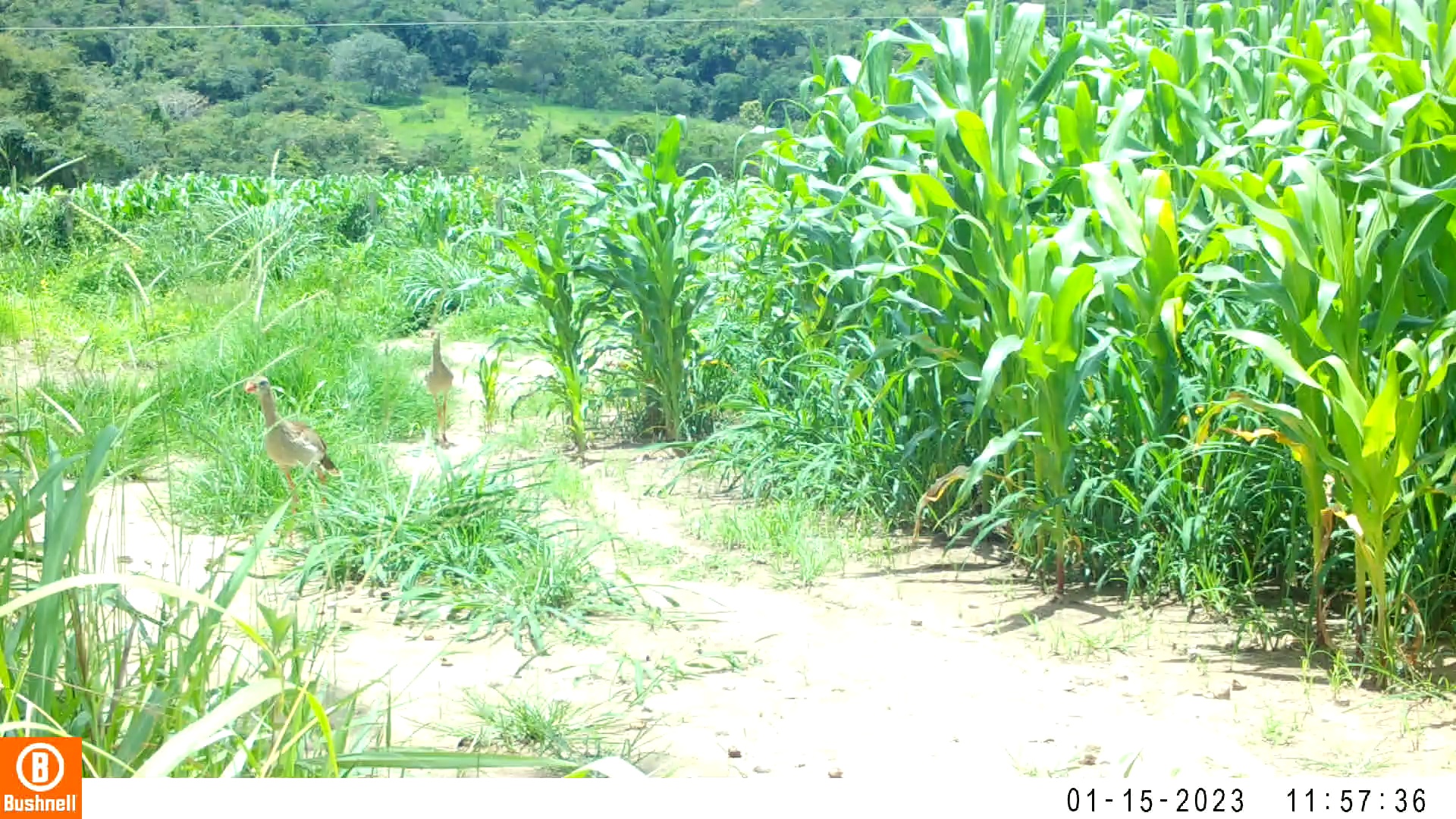

### 01150027-01940-404.jpg

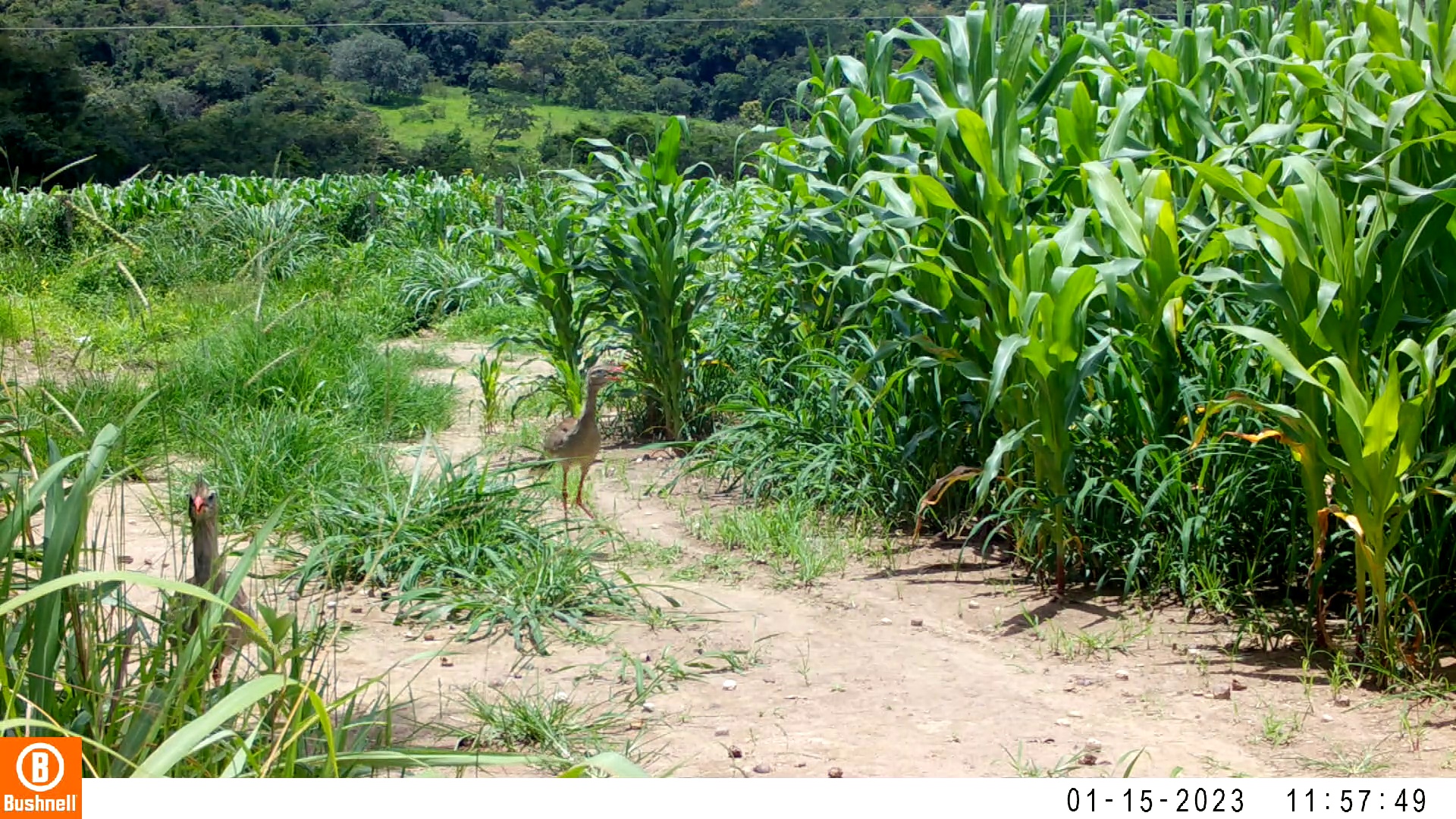

### 01150027-01940-808.jpg

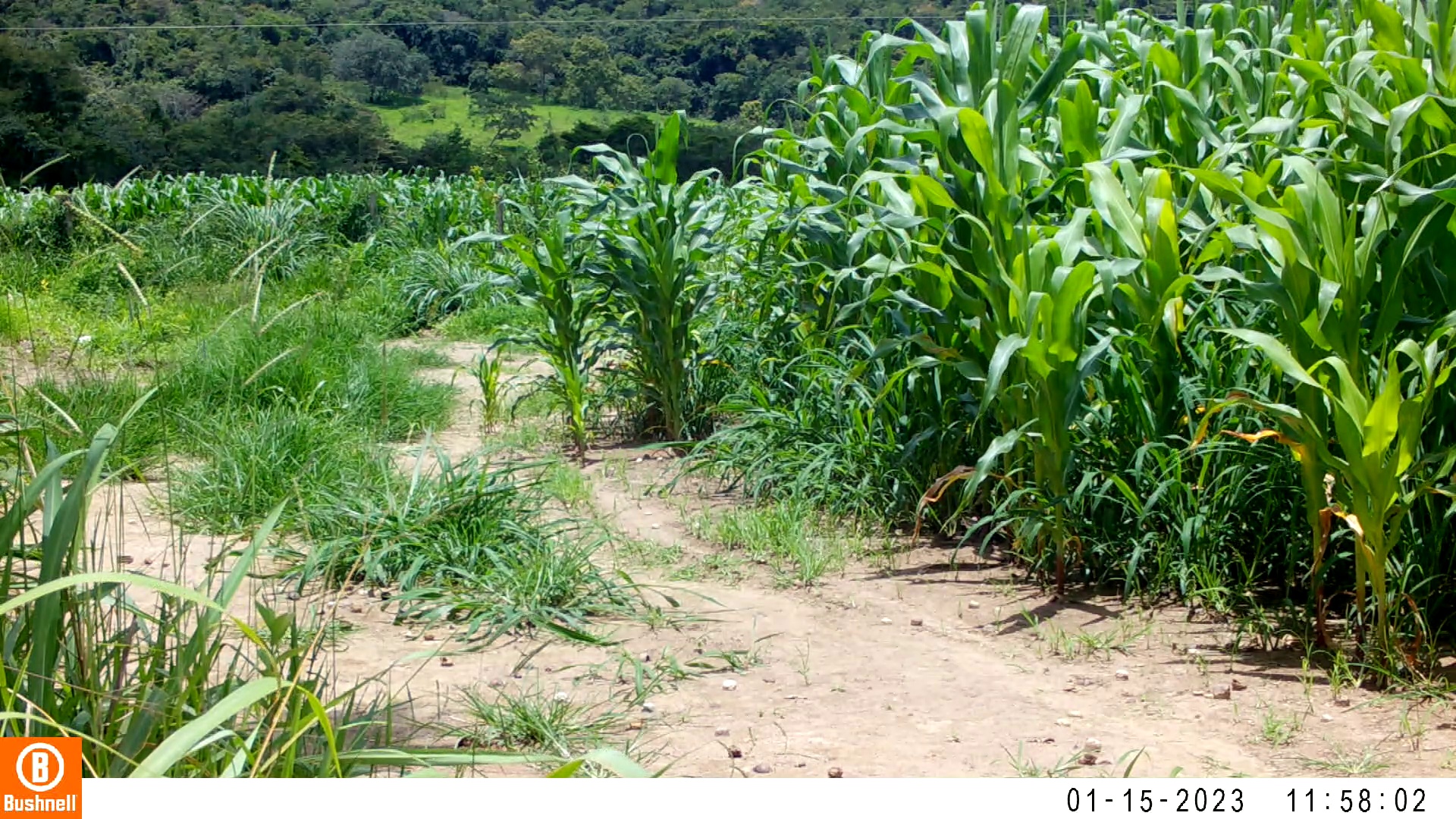

### 01150027-01940-1010.jpg

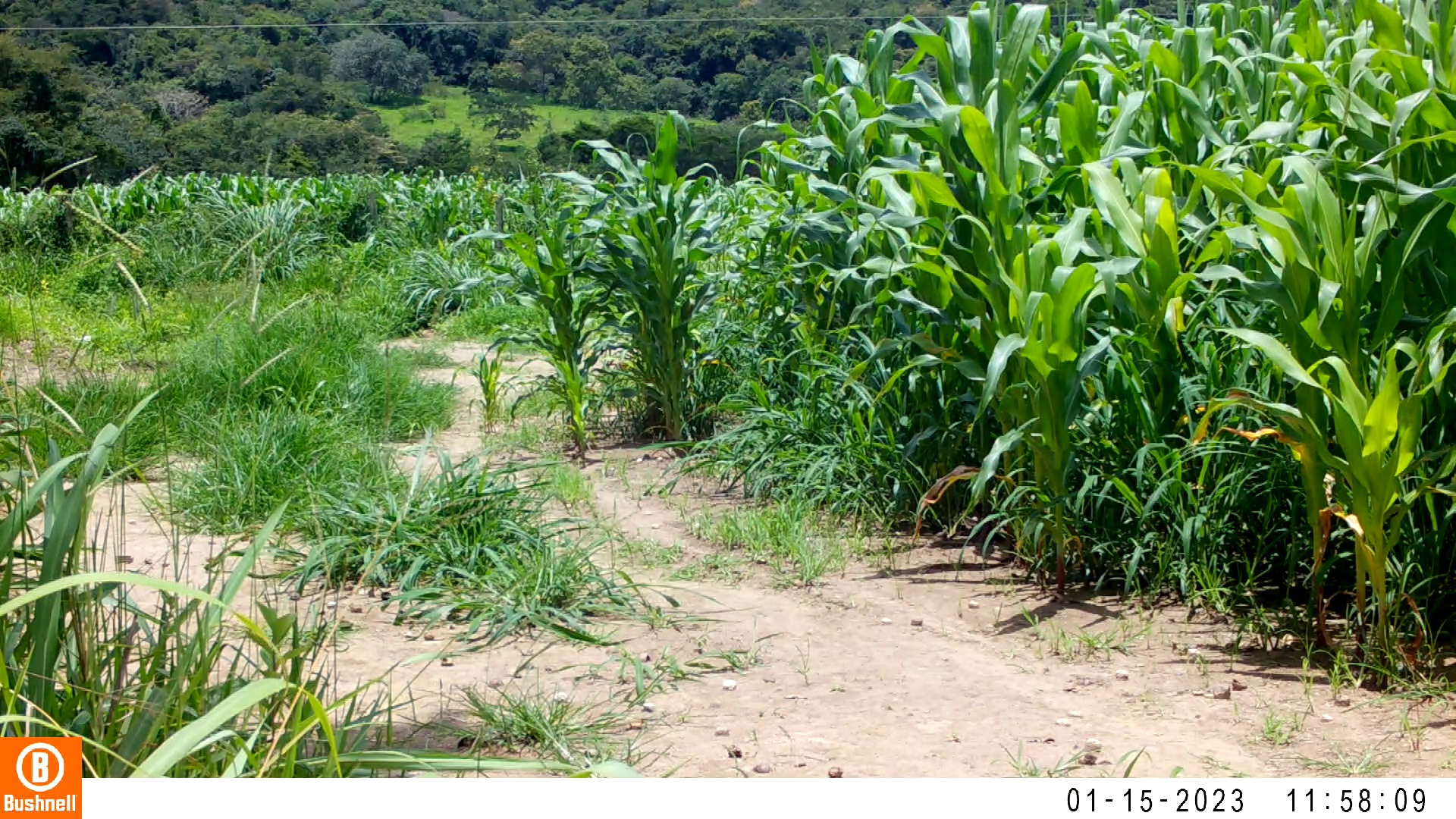

### 01150030-10180-0.jpg

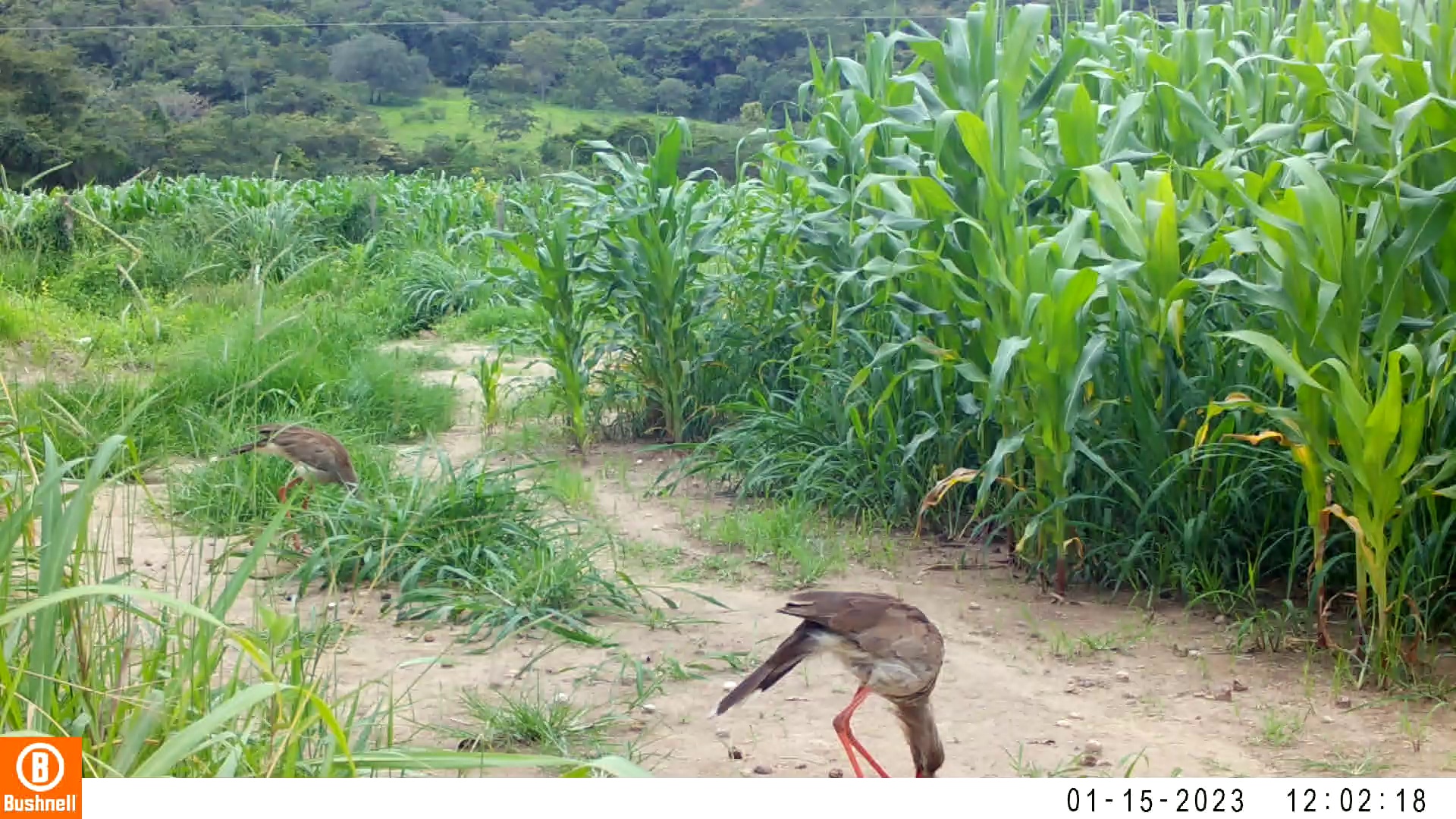

### 01150030-10180-202.jpg

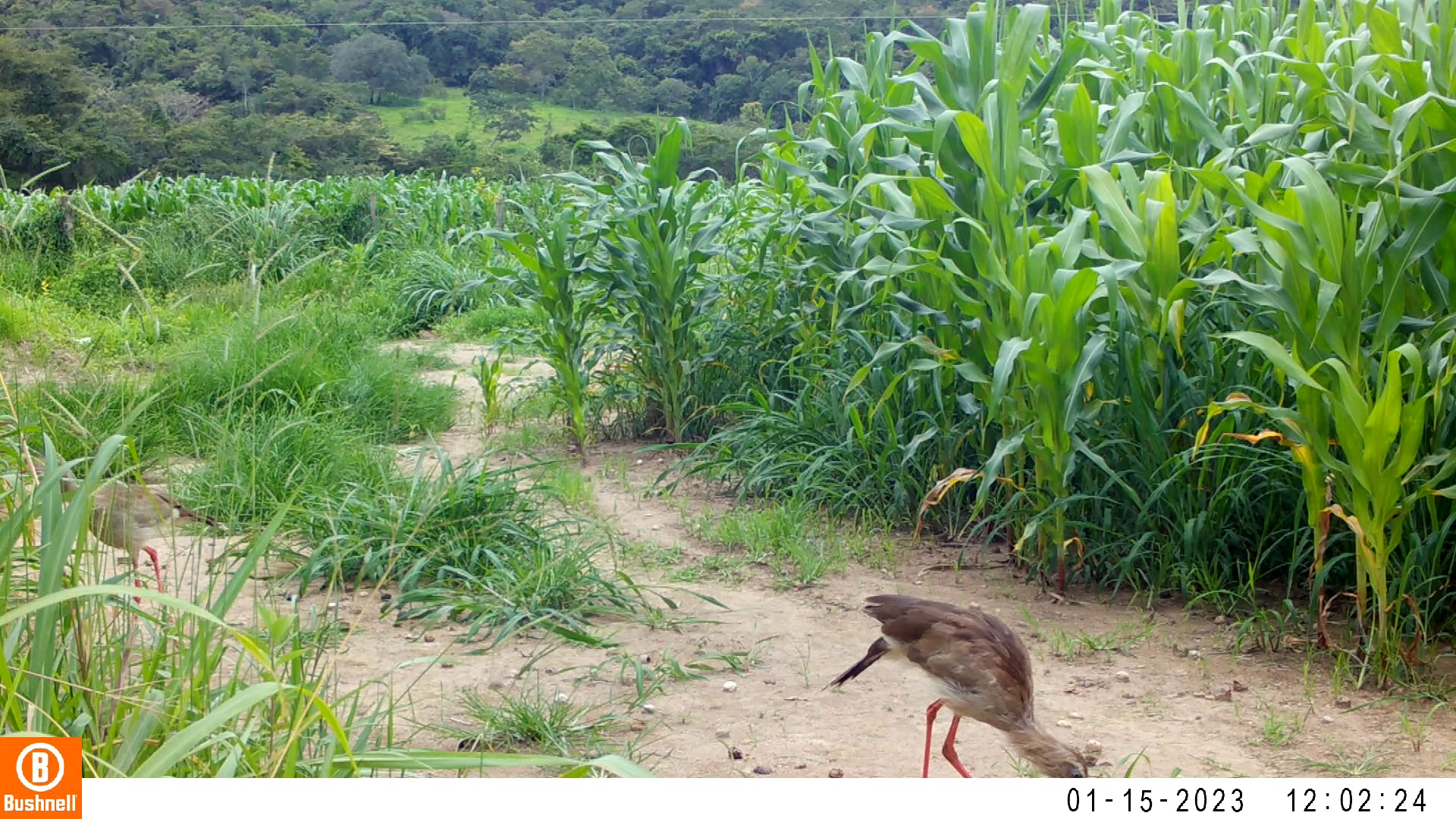

### 01150030-10180-404.jpg

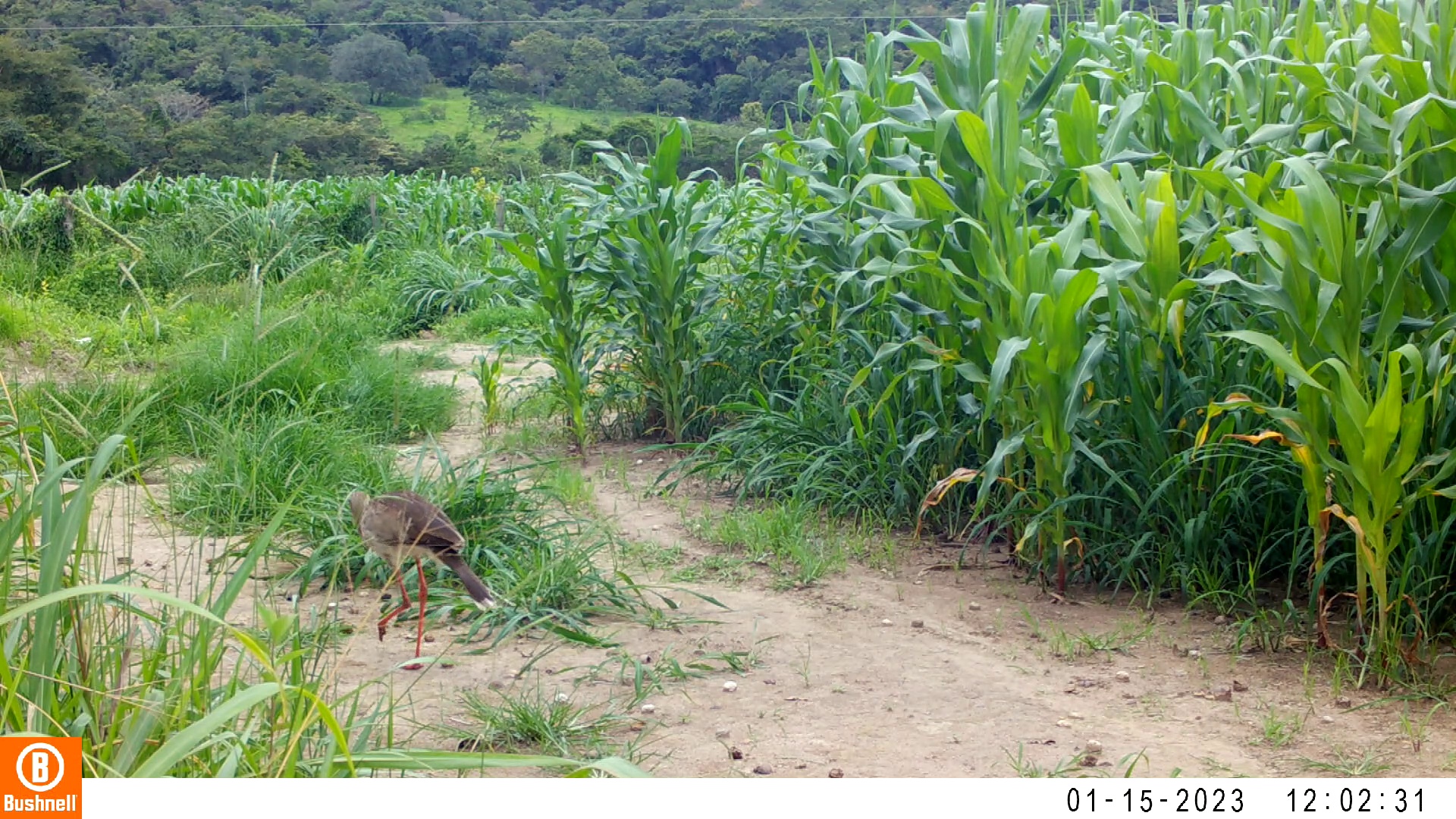

### 01150030-10180-606.jpg

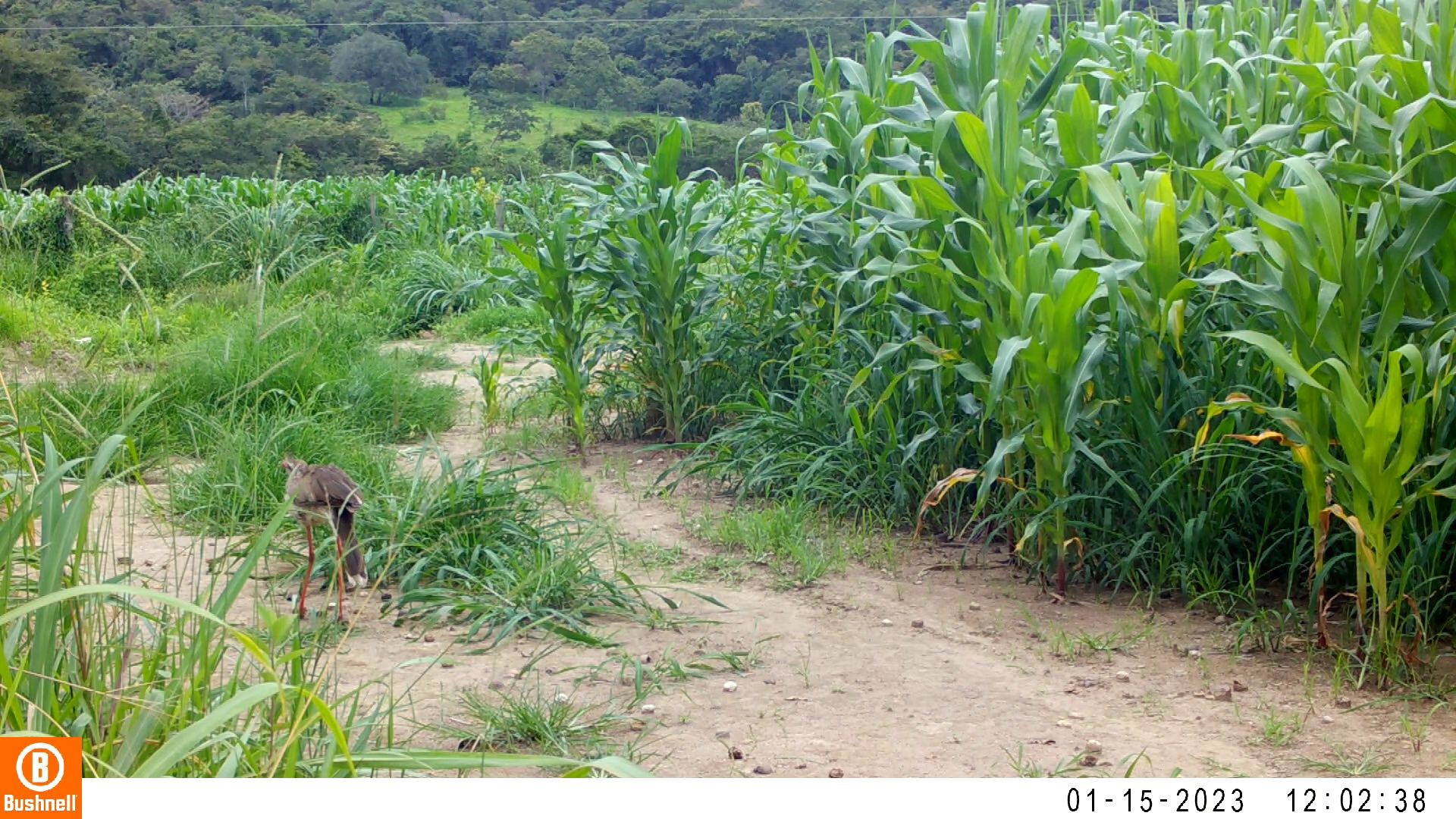

### 01150030-10180-808.jpg

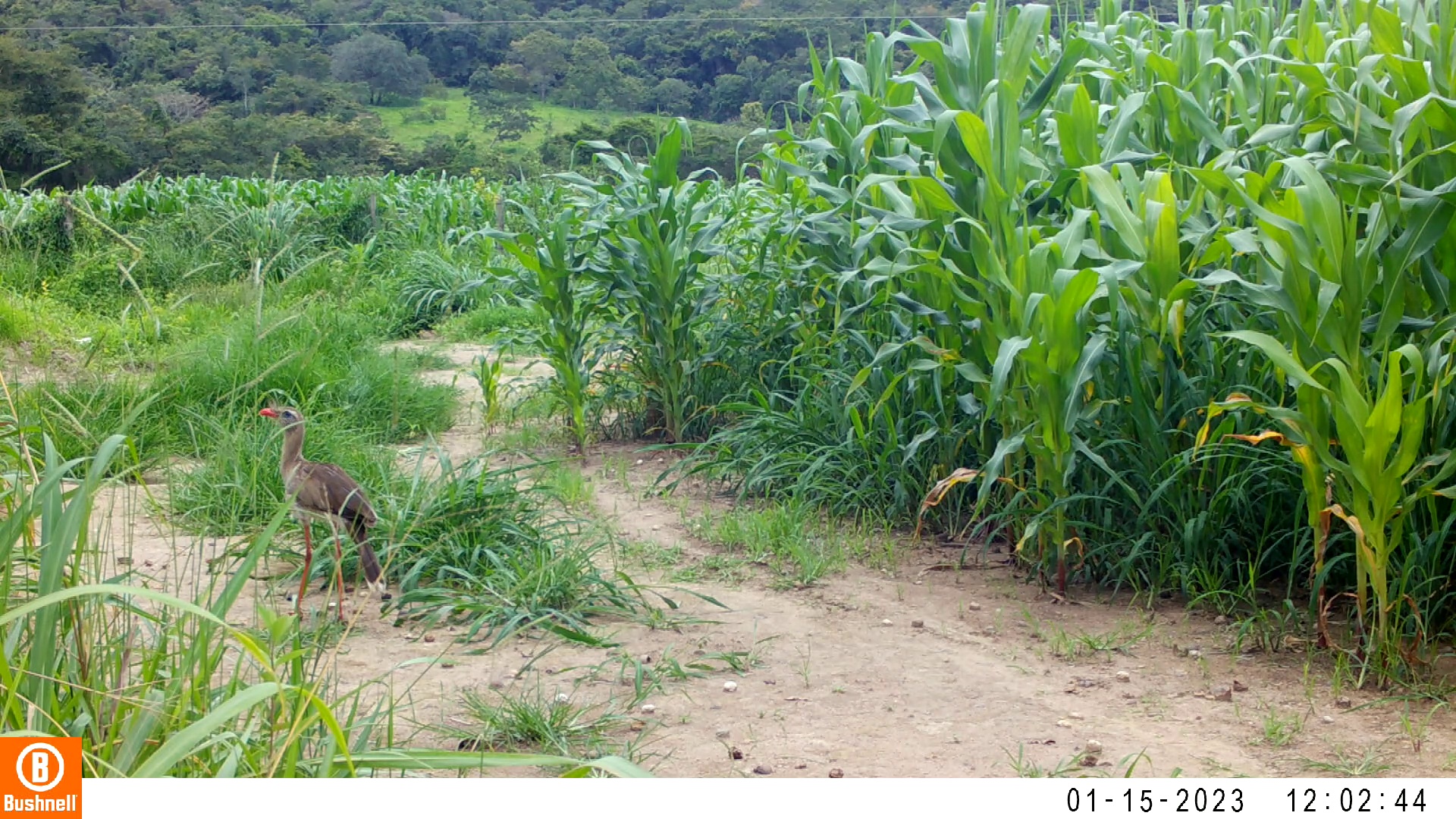

### 01150030-10180-1010.jpg

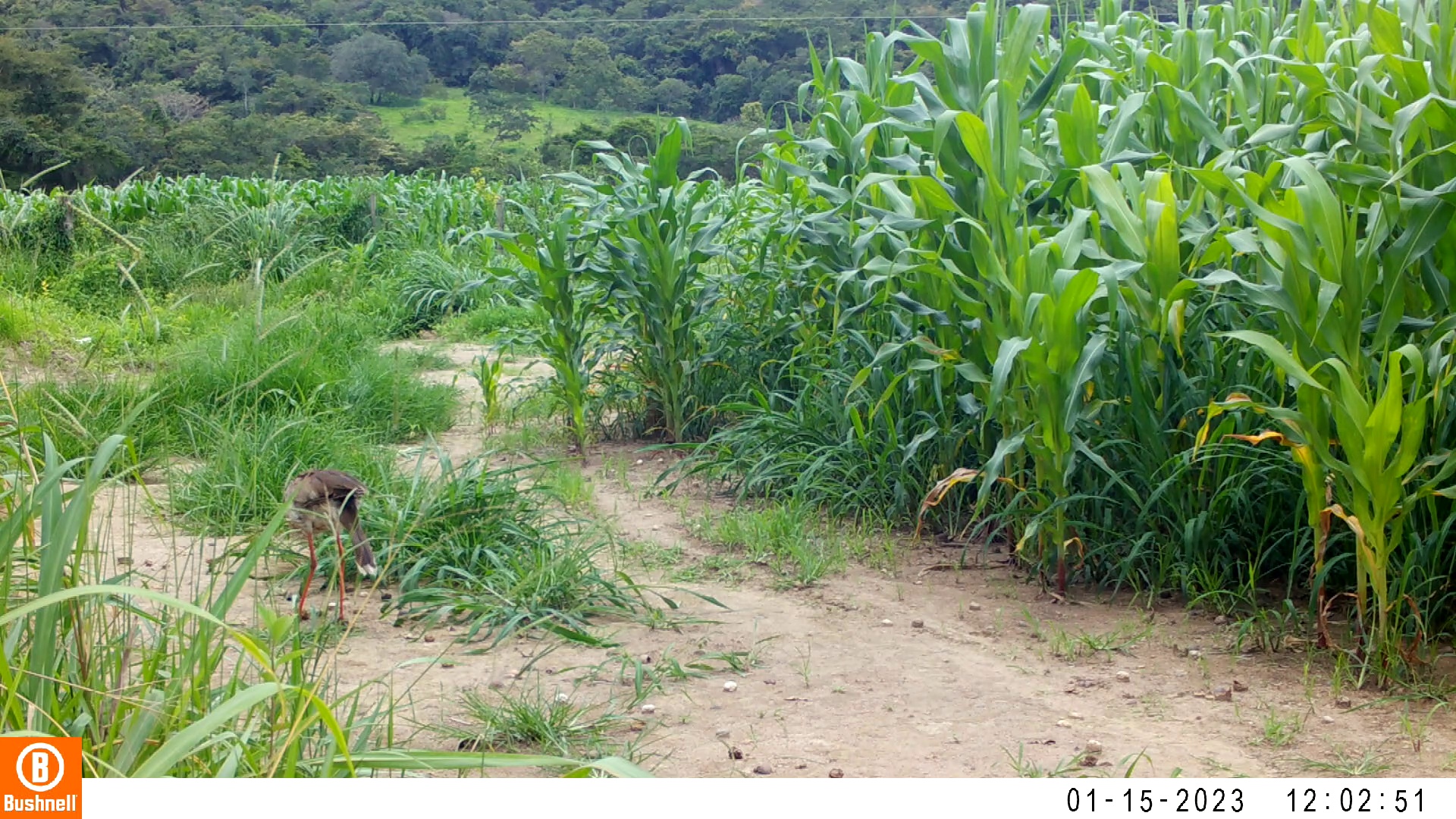

### 01150035-67464-150.jpg

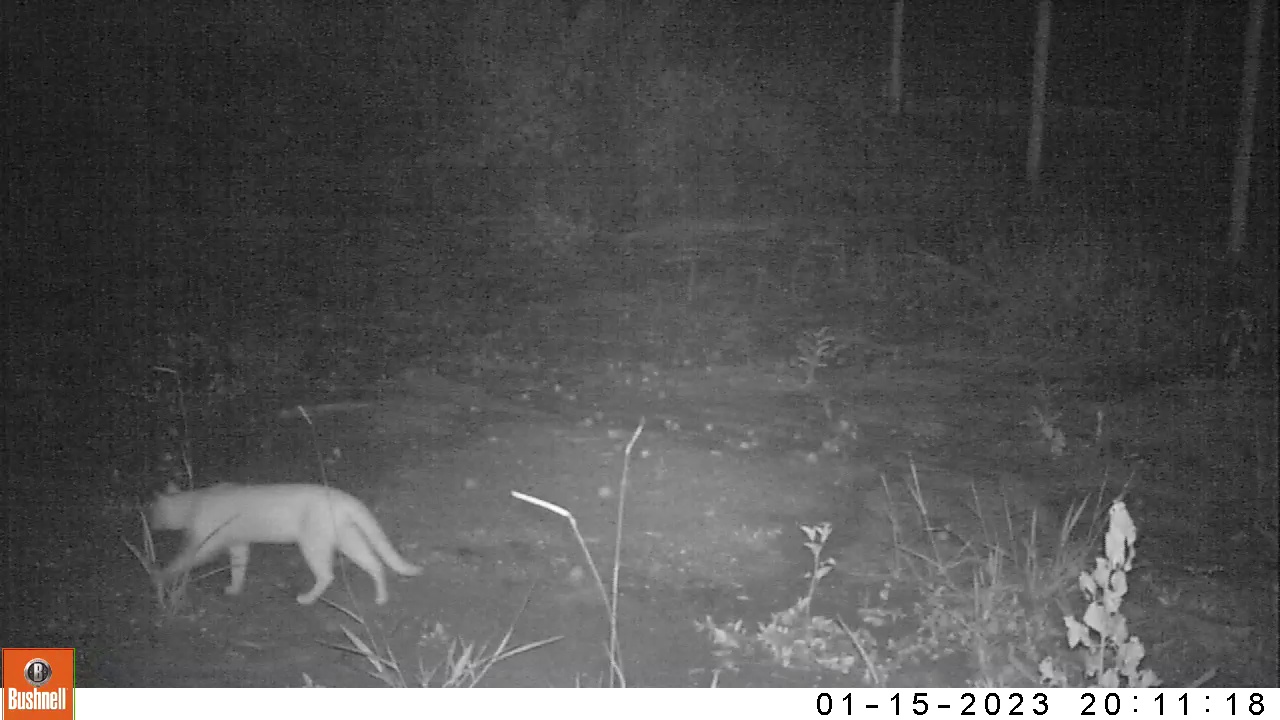

### 01150035-67464-300.jpg

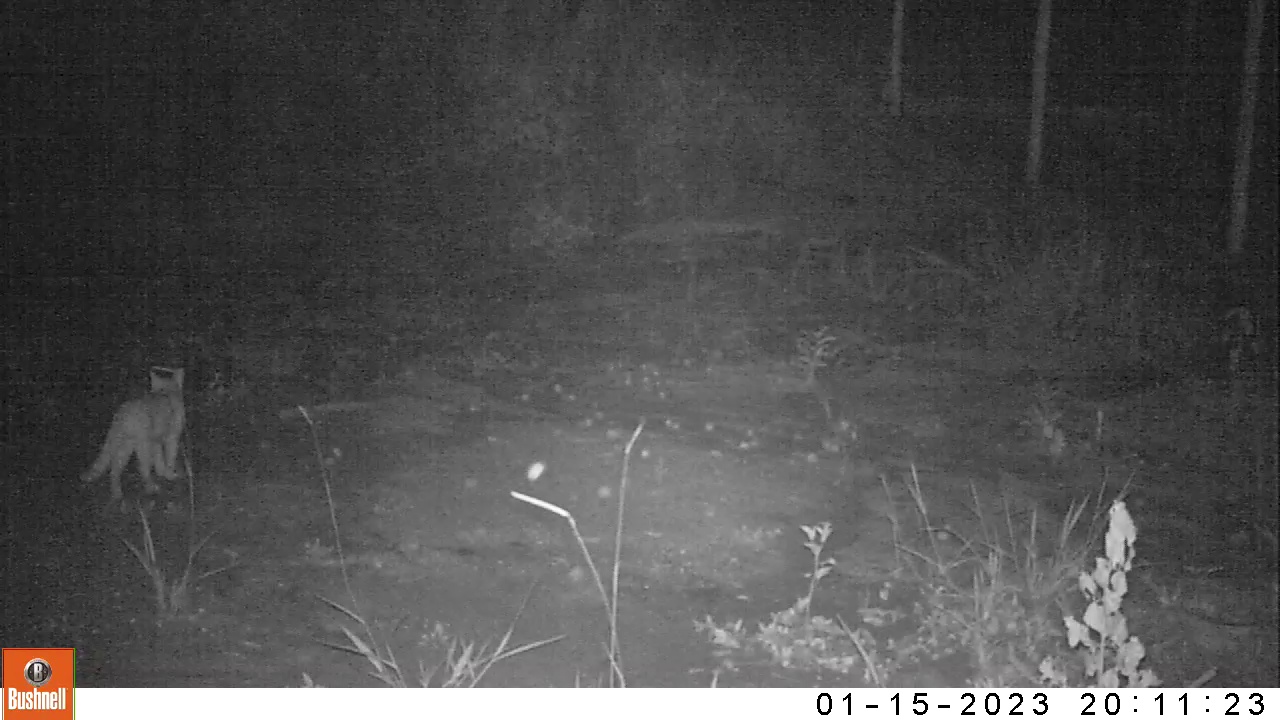

### 01150035-67464-450.jpg

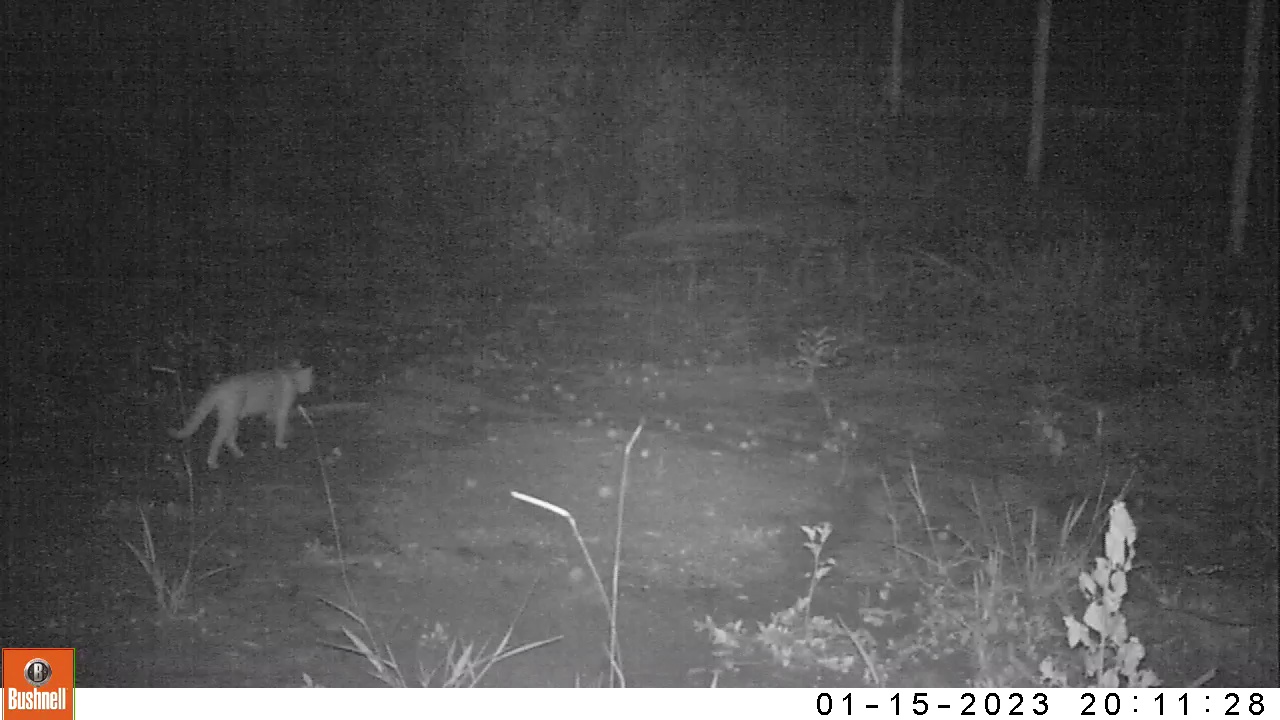

### 01150035-67464-600.jpg

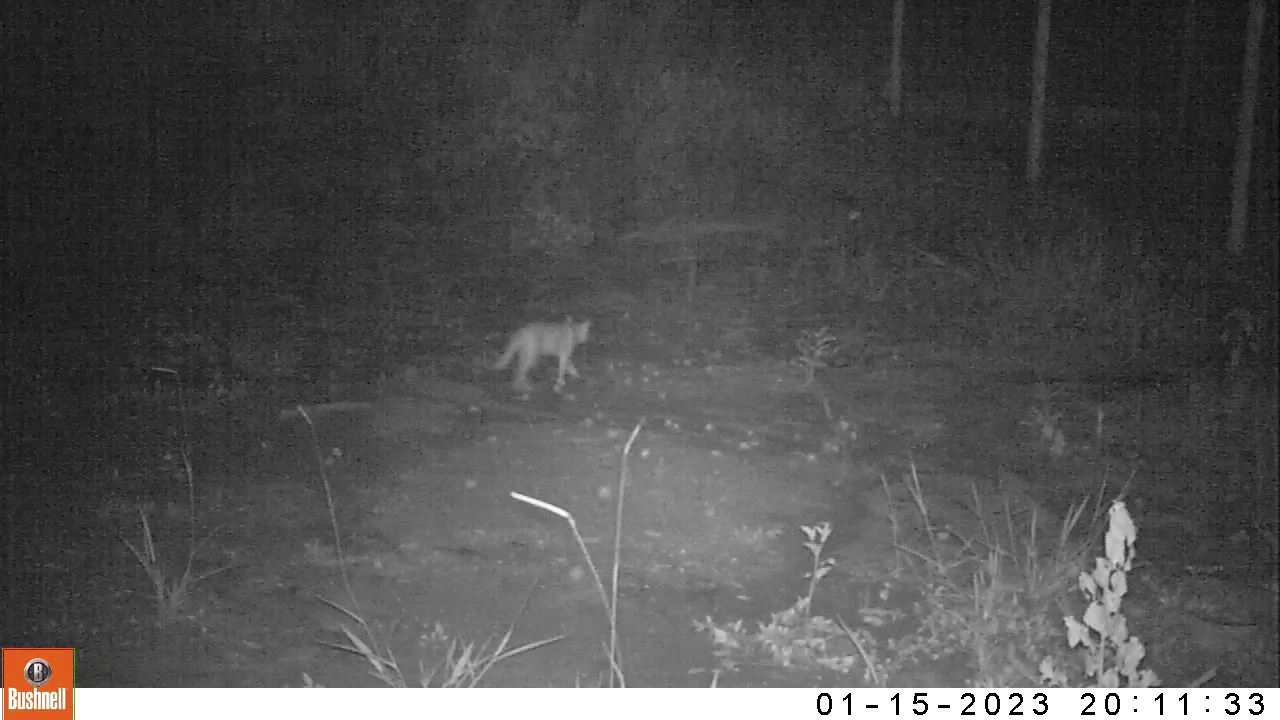

### 01150035-67464-750.jpg

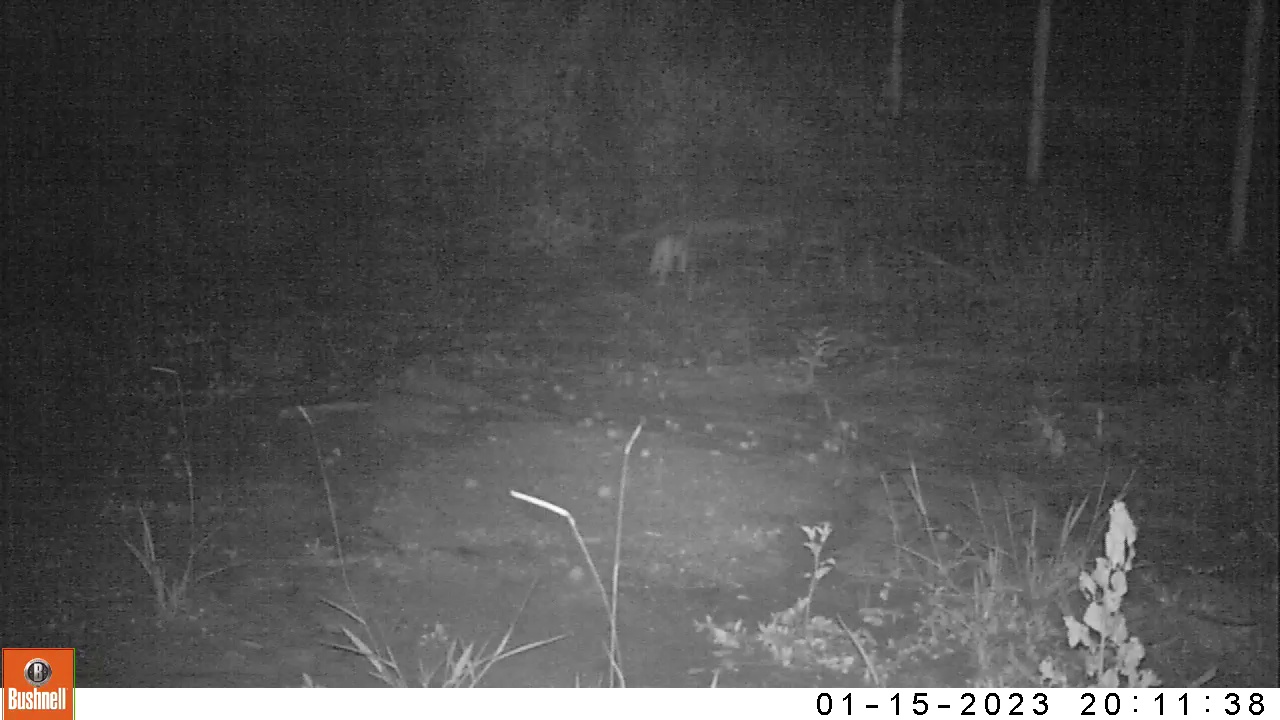

### 01160040-03309-0.jpg

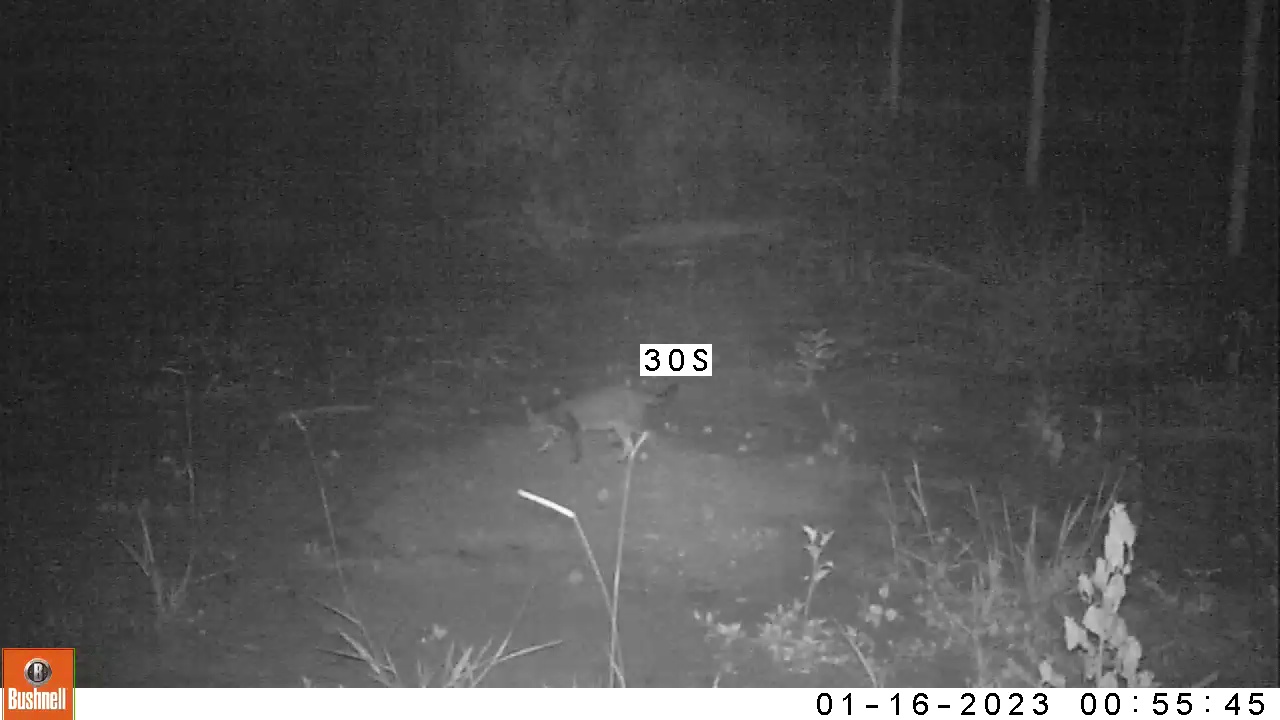

### 01160040-03309-150.jpg

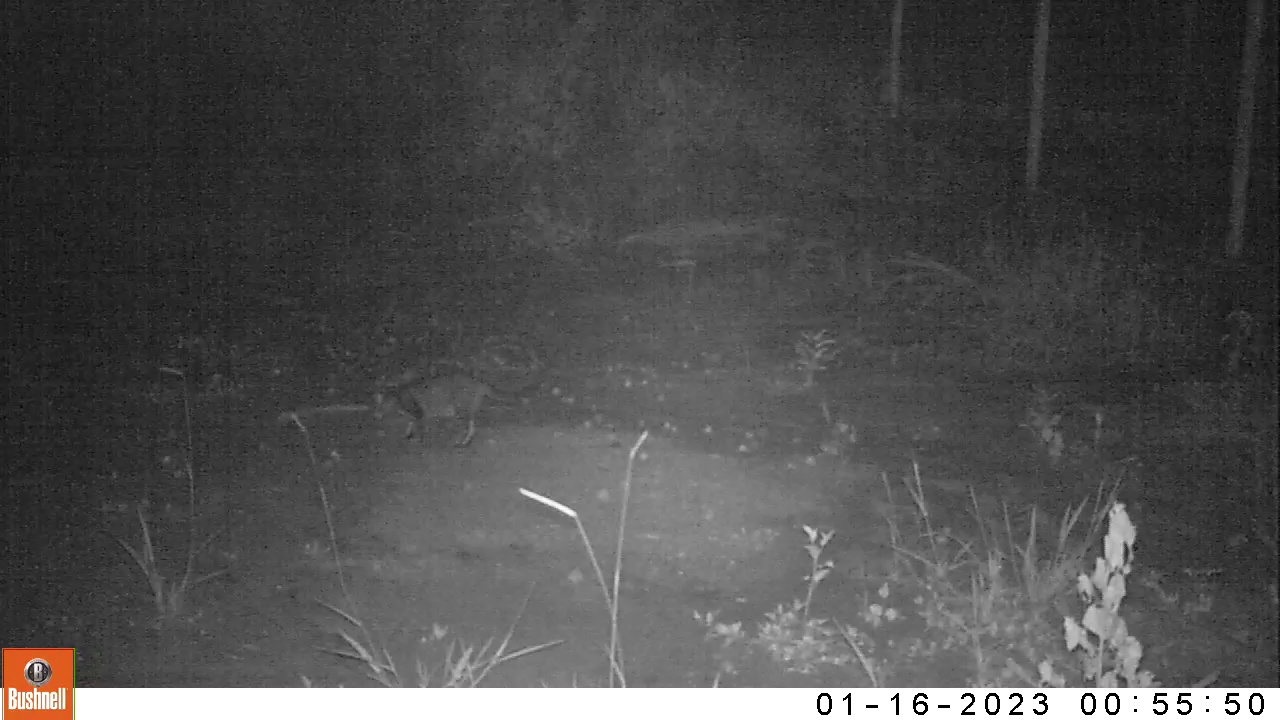

### 01160040-03309-300.jpg

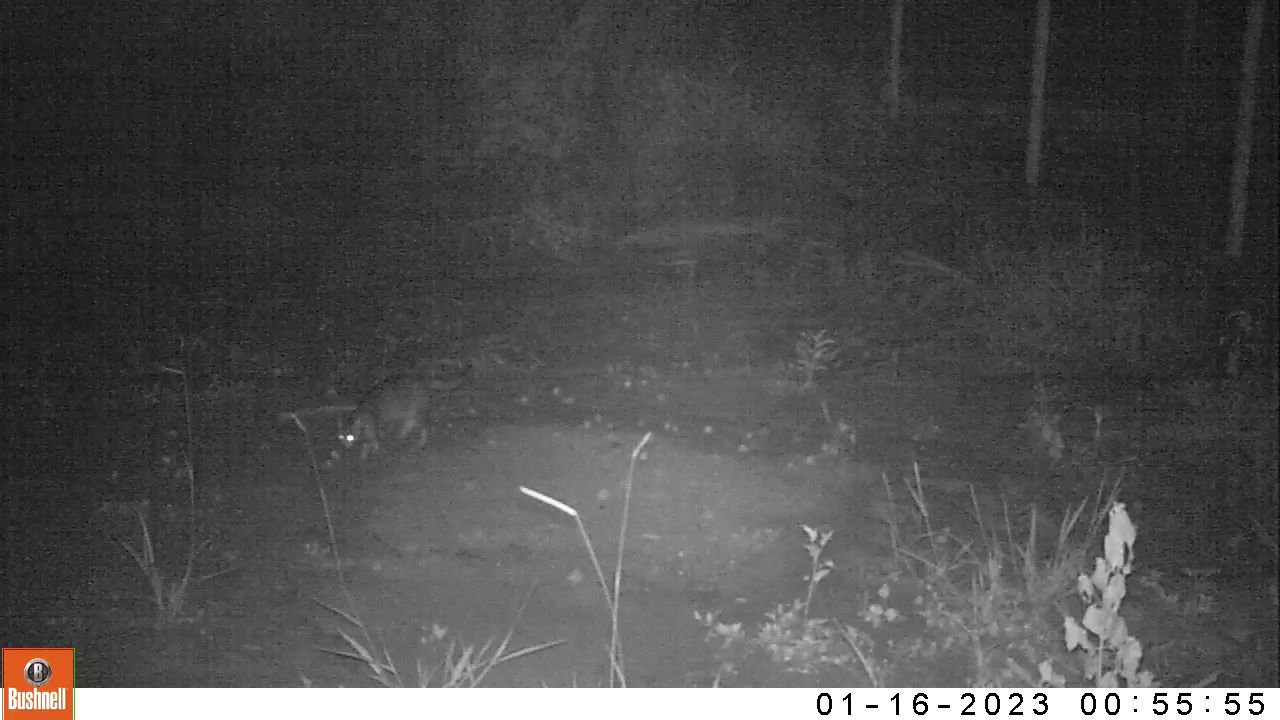

### 01160040-03309-450.jpg

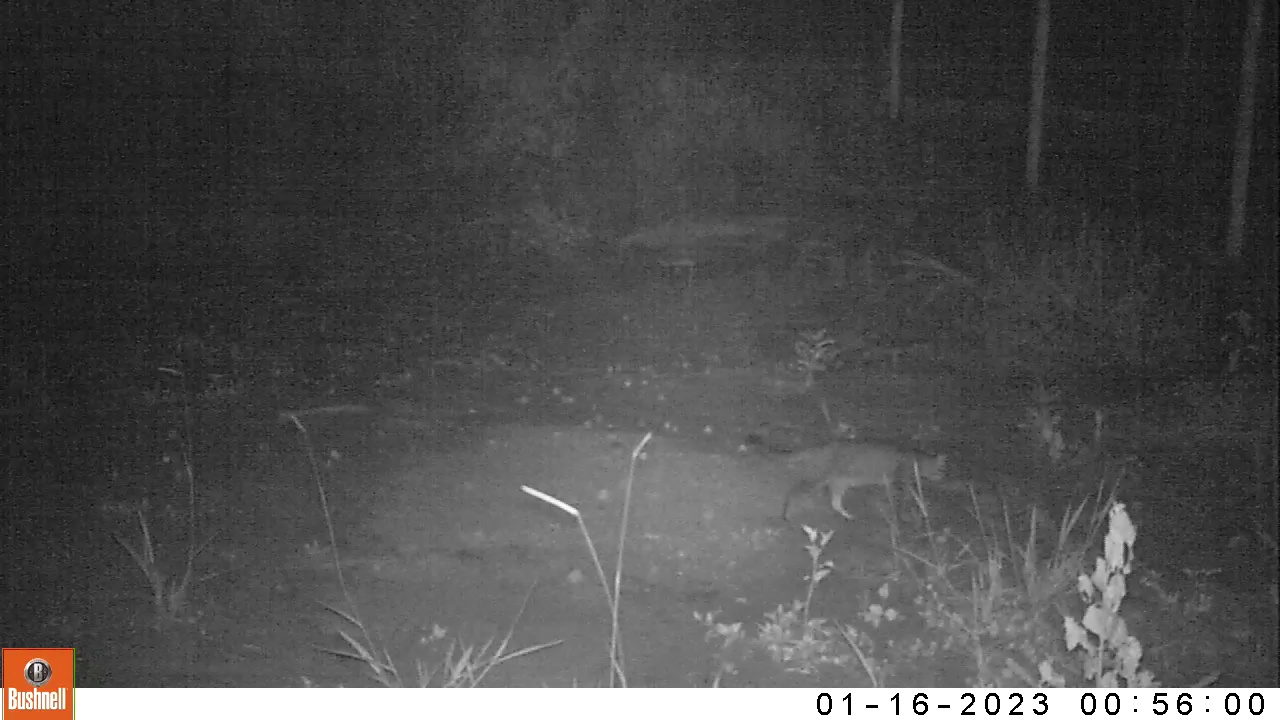

### 01160040-03309-600.jpg

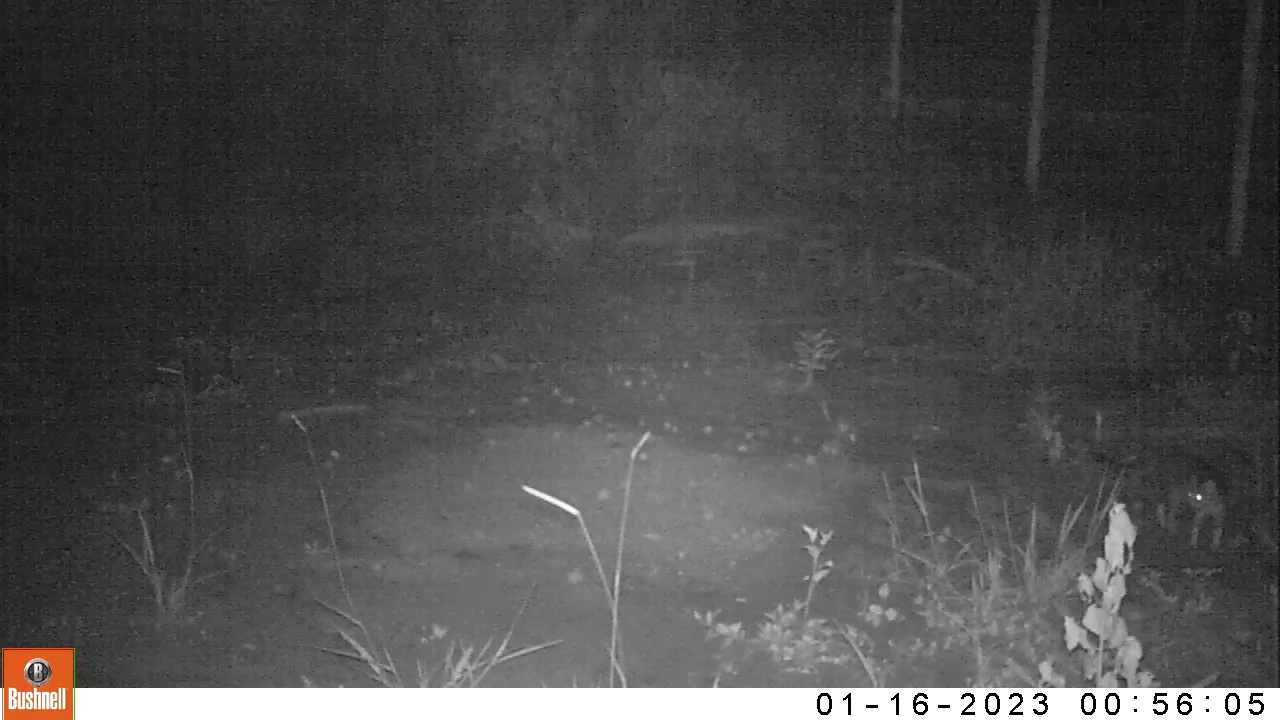

### 01160040-03309-750.jpg

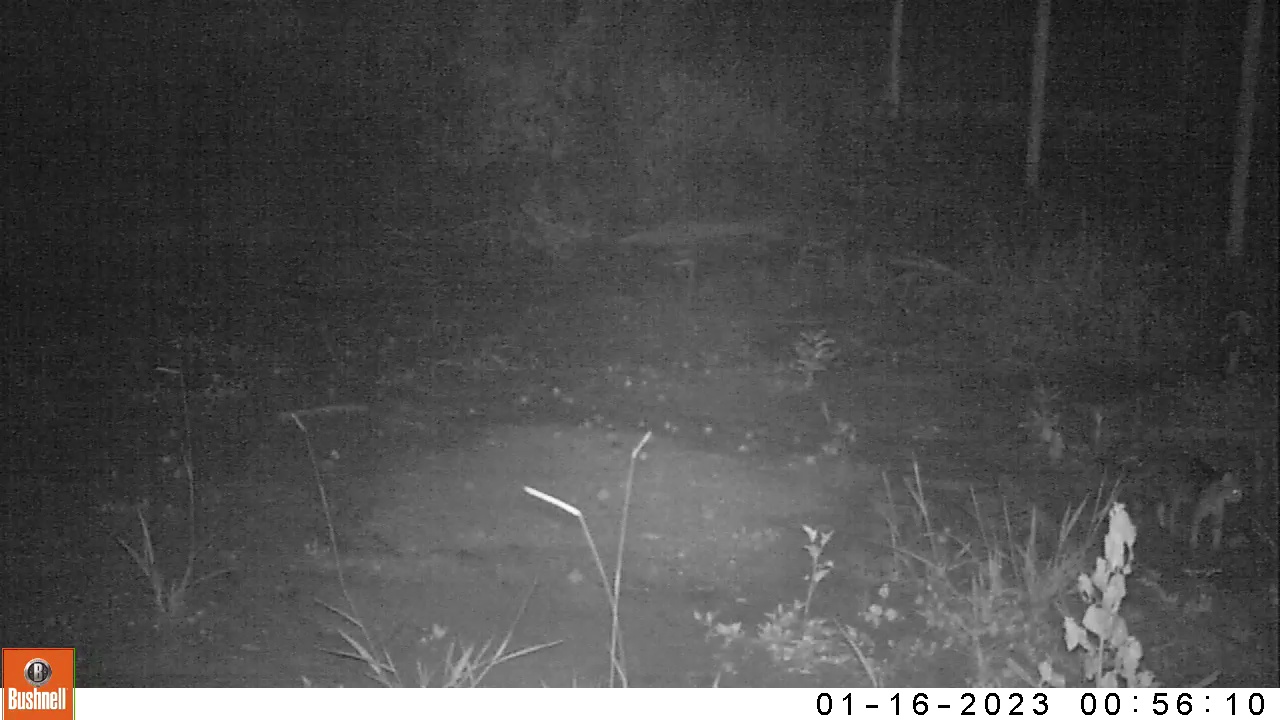

### 01170026-18885-0.jpg

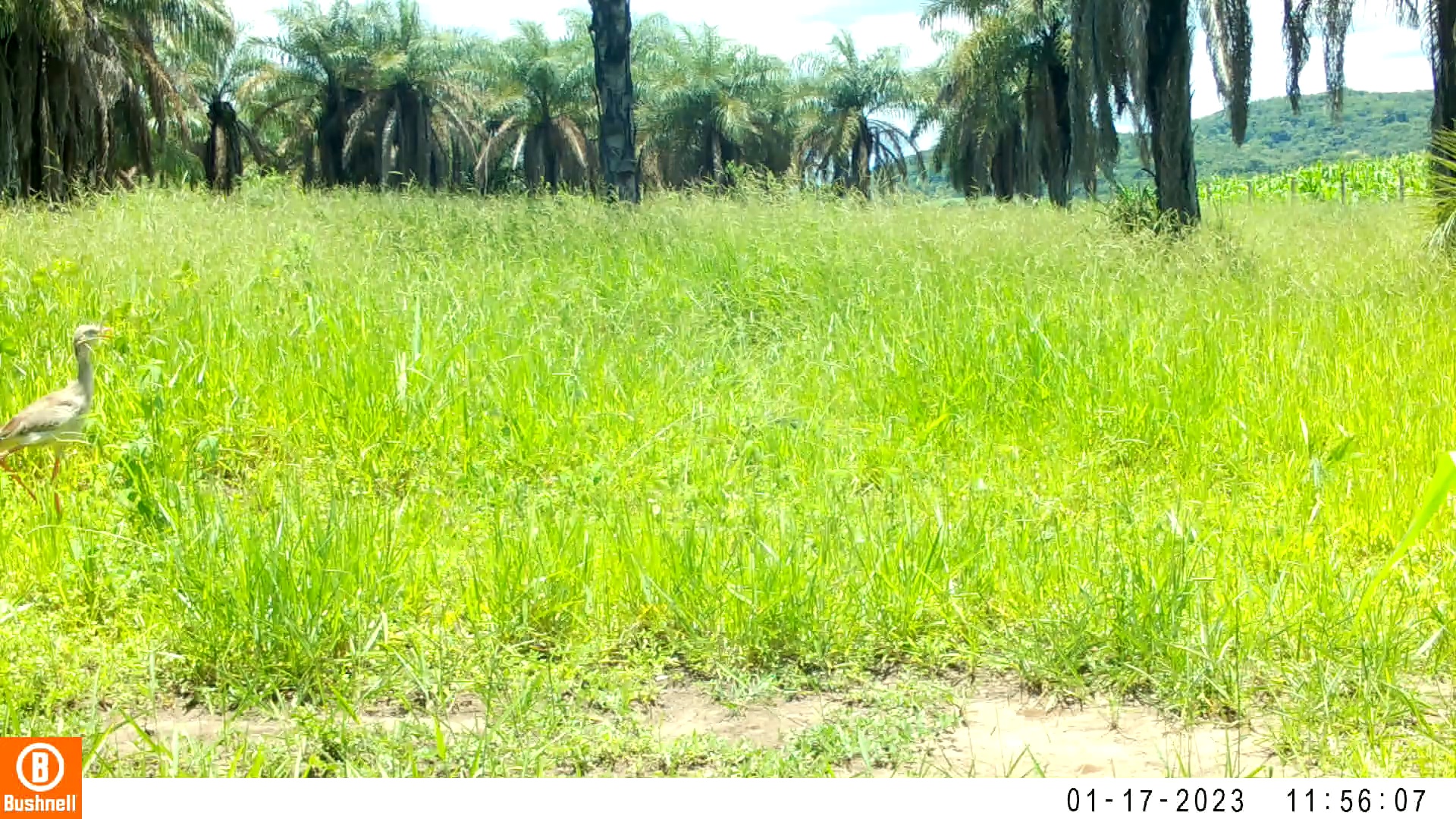

### 01170026-18885-52.jpg

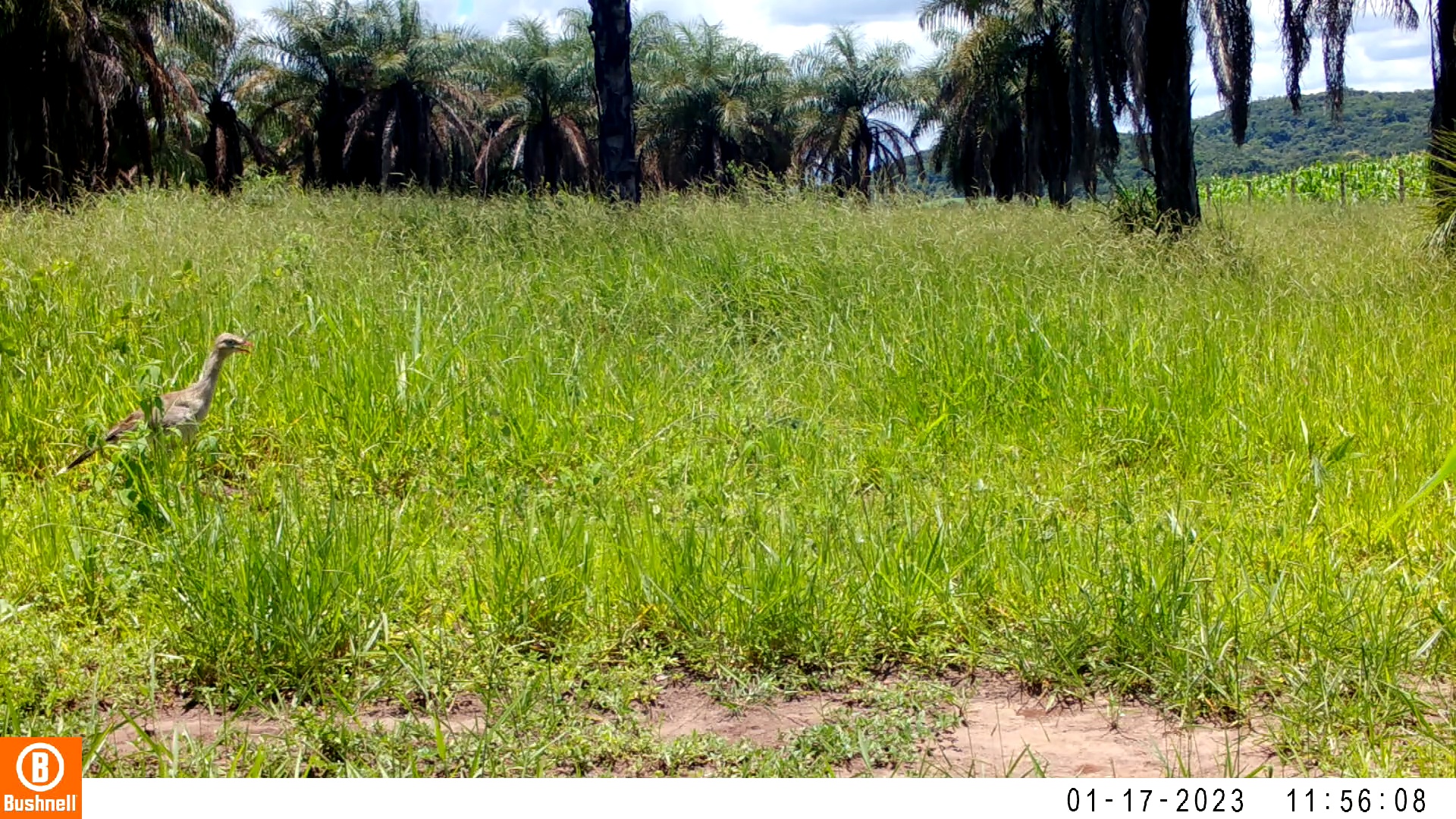

### 01170026-18885-104.jpg

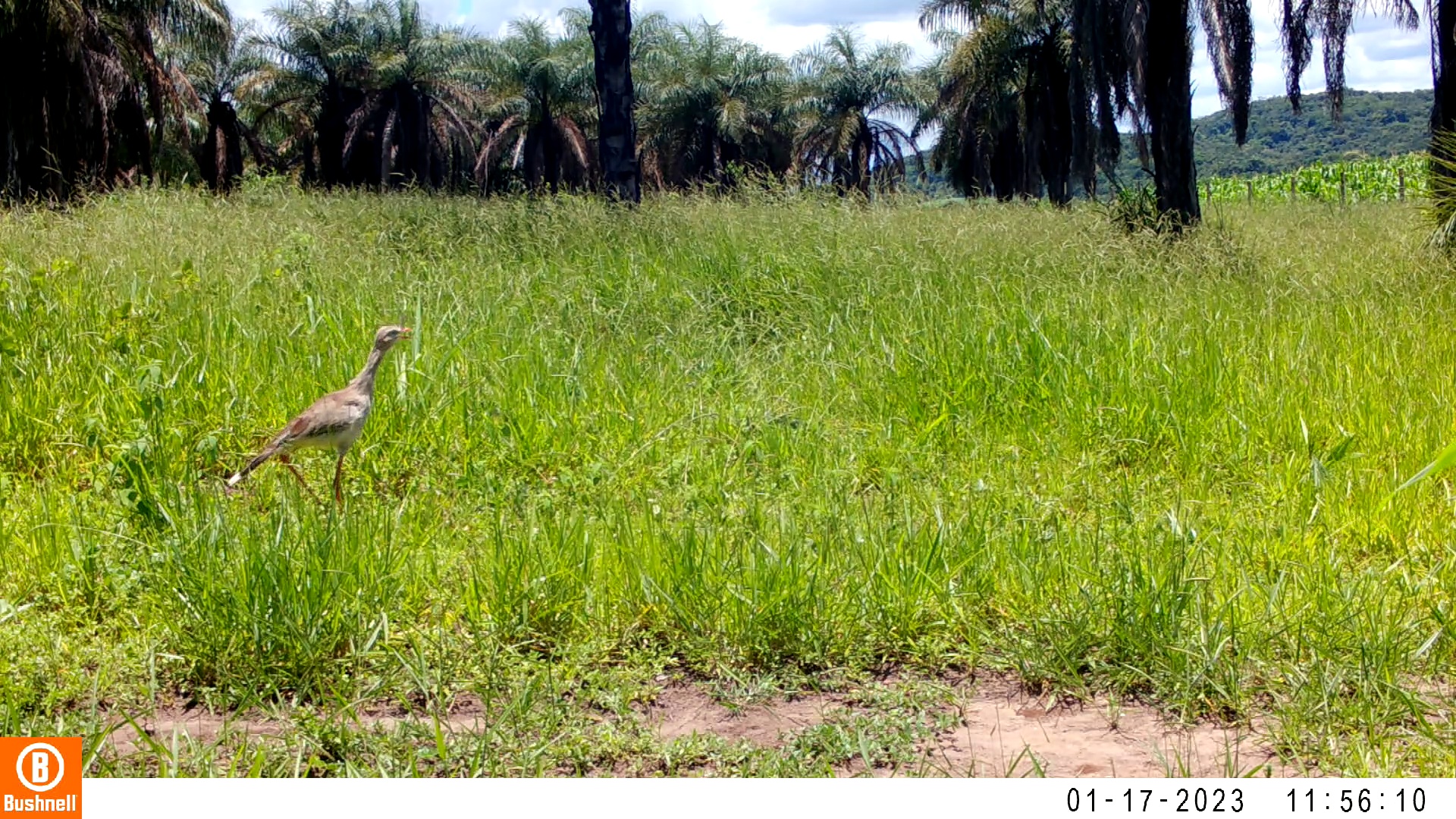

### 01170026-18885-156.jpg

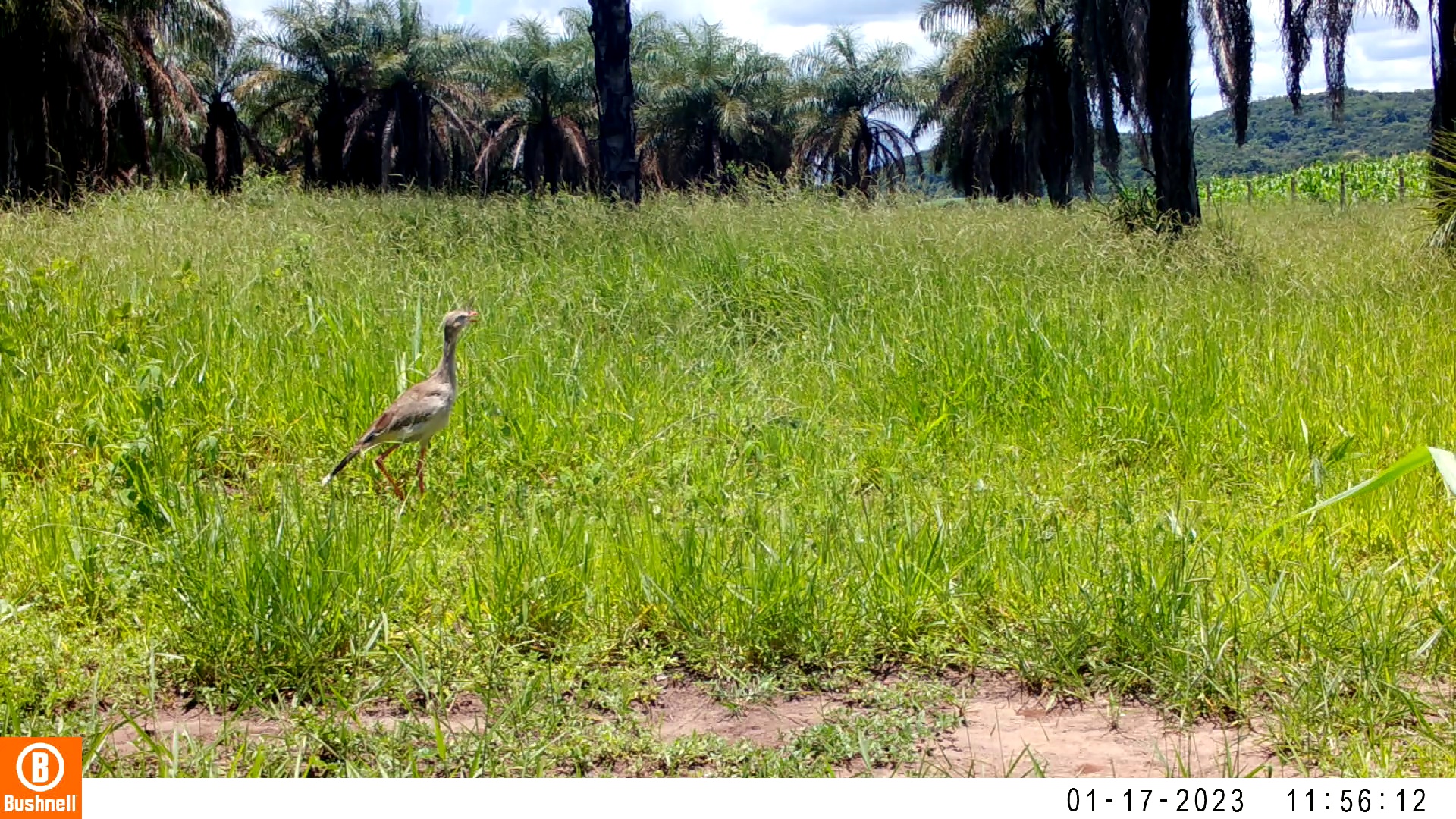

### 01170026-18885-208.jpg

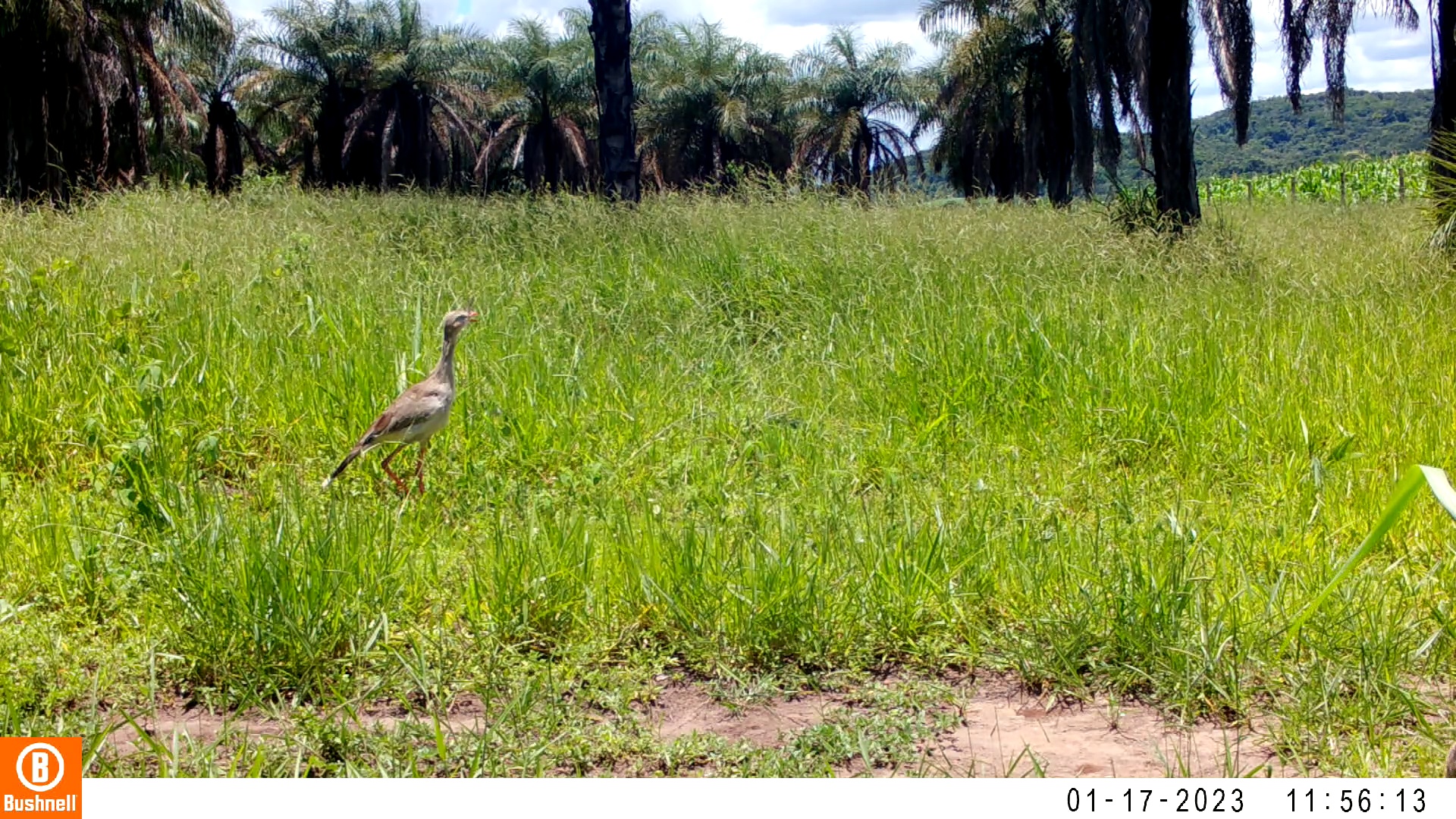

### 01170026-18885-260.jpg

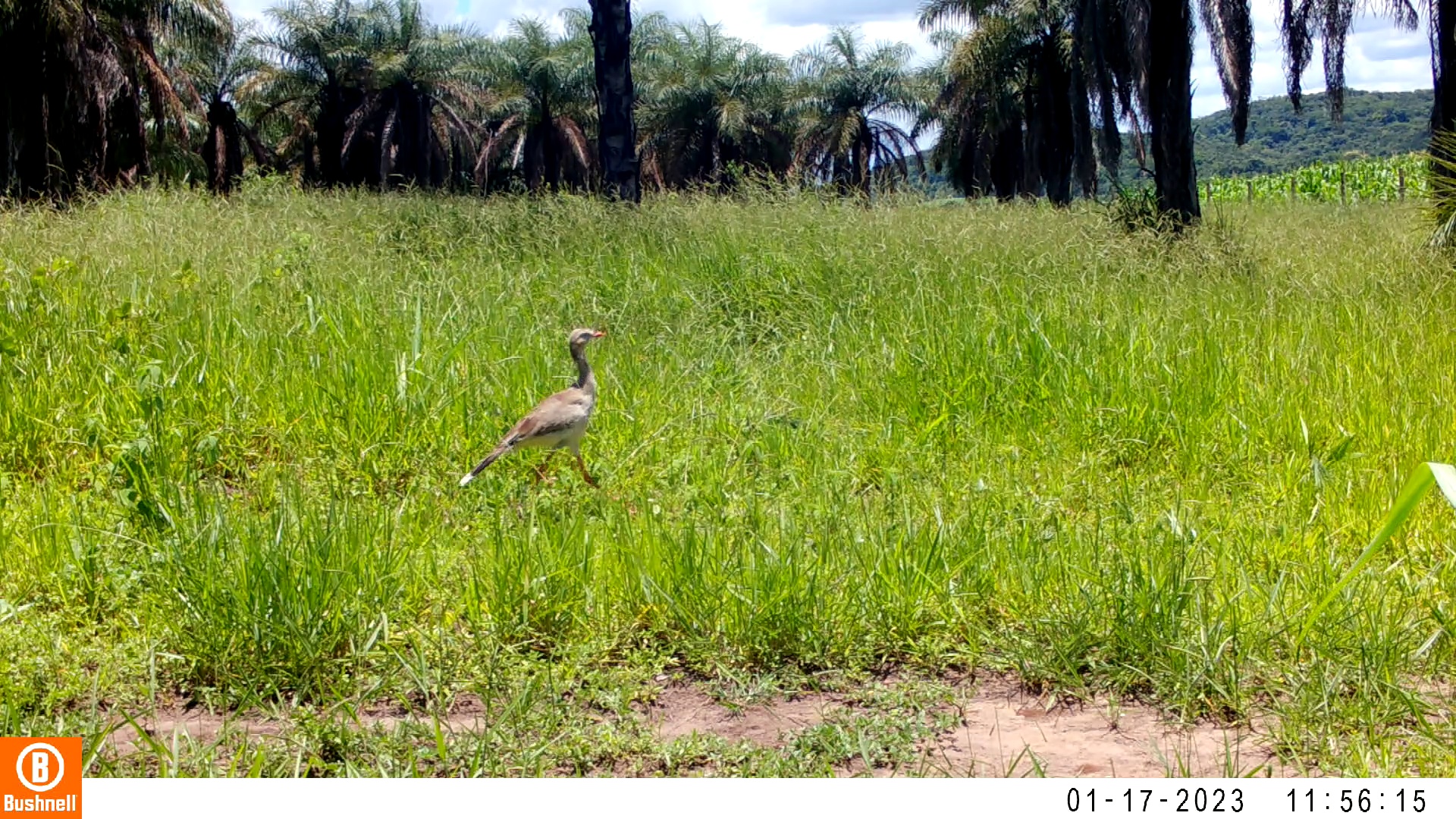

### 01170027-52181-0.jpg

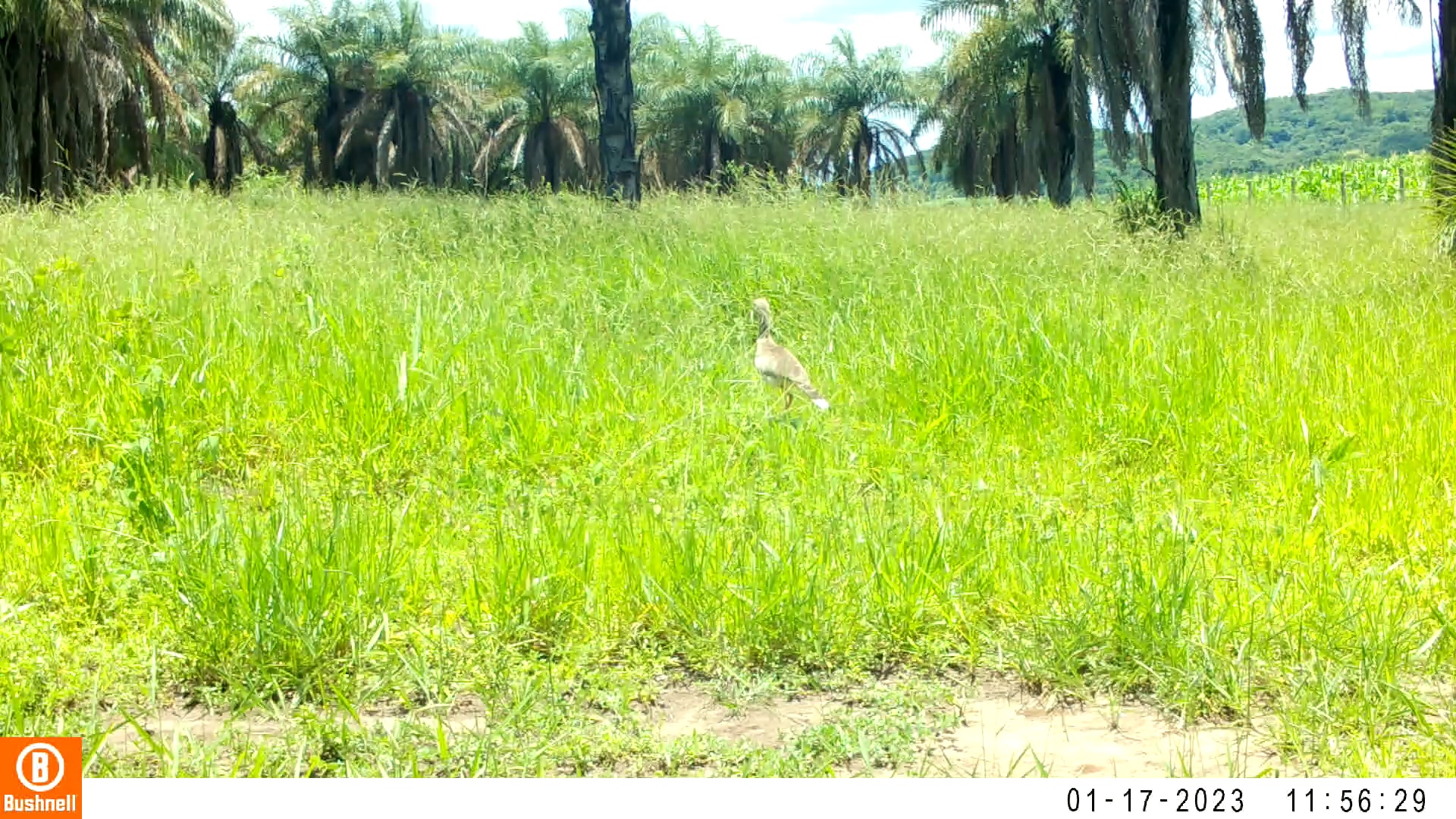

### 01170027-52181-52.jpg

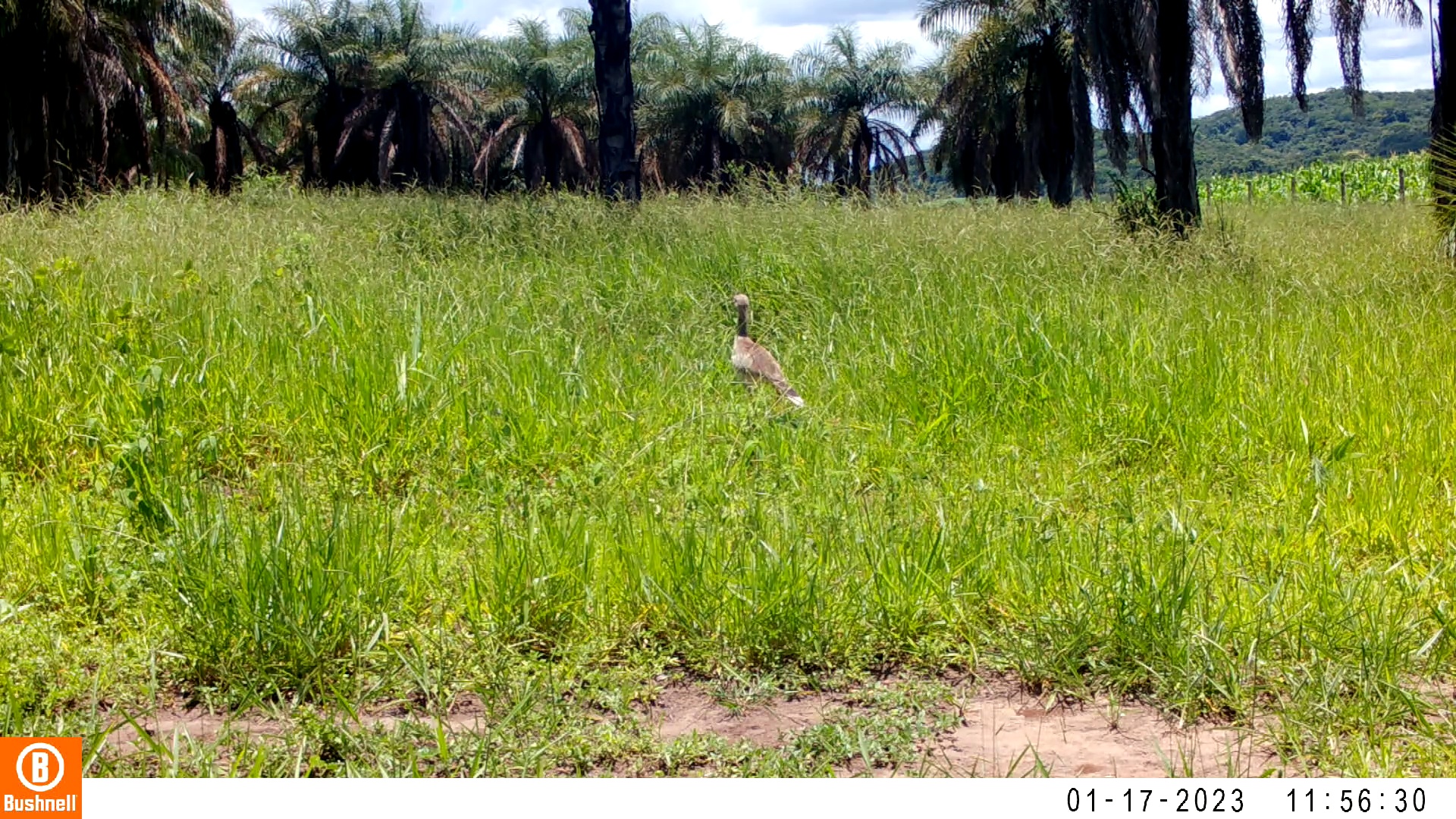

### 01170027-52181-260.jpg

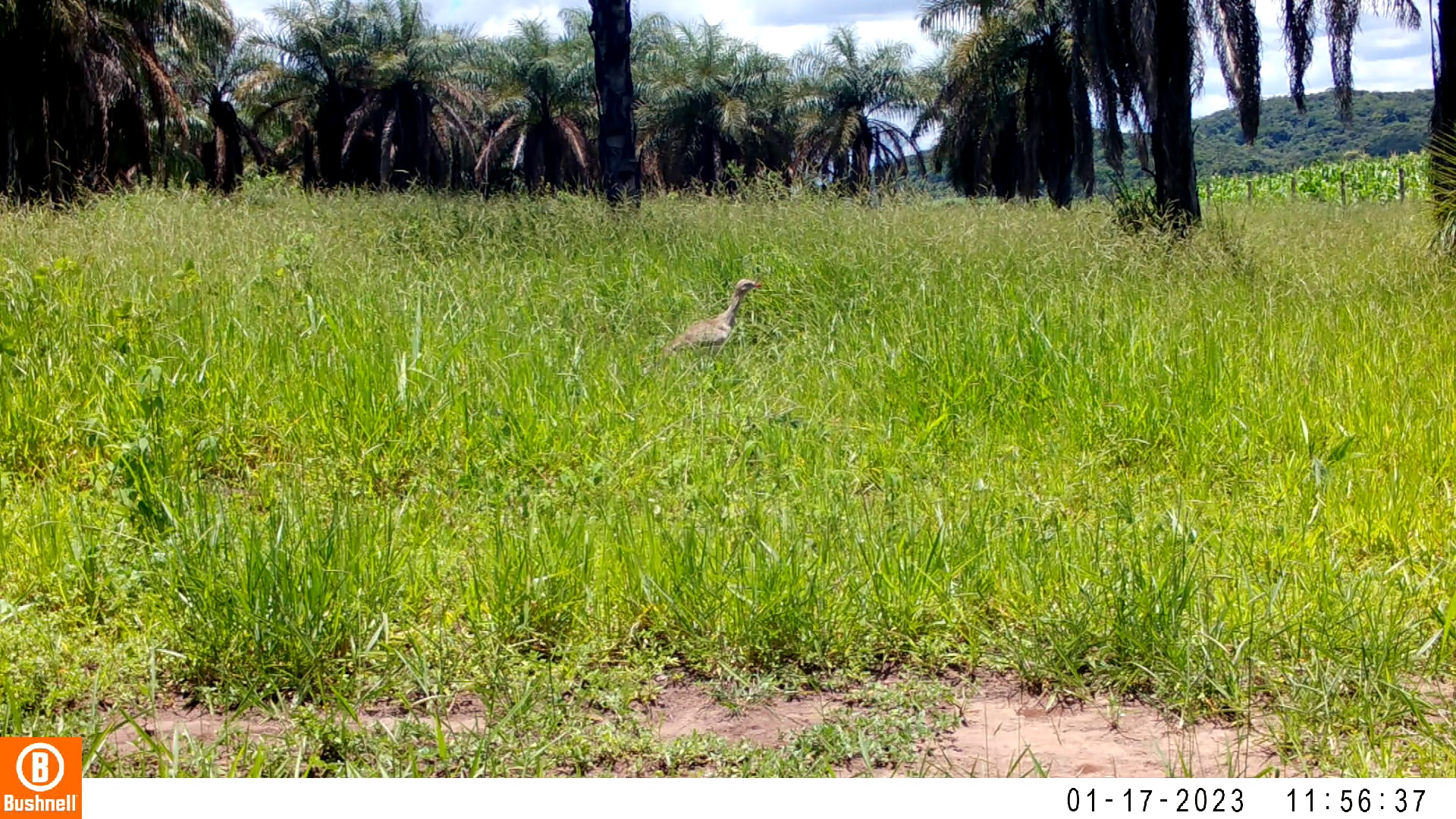

### 01170028-88262-52.jpg

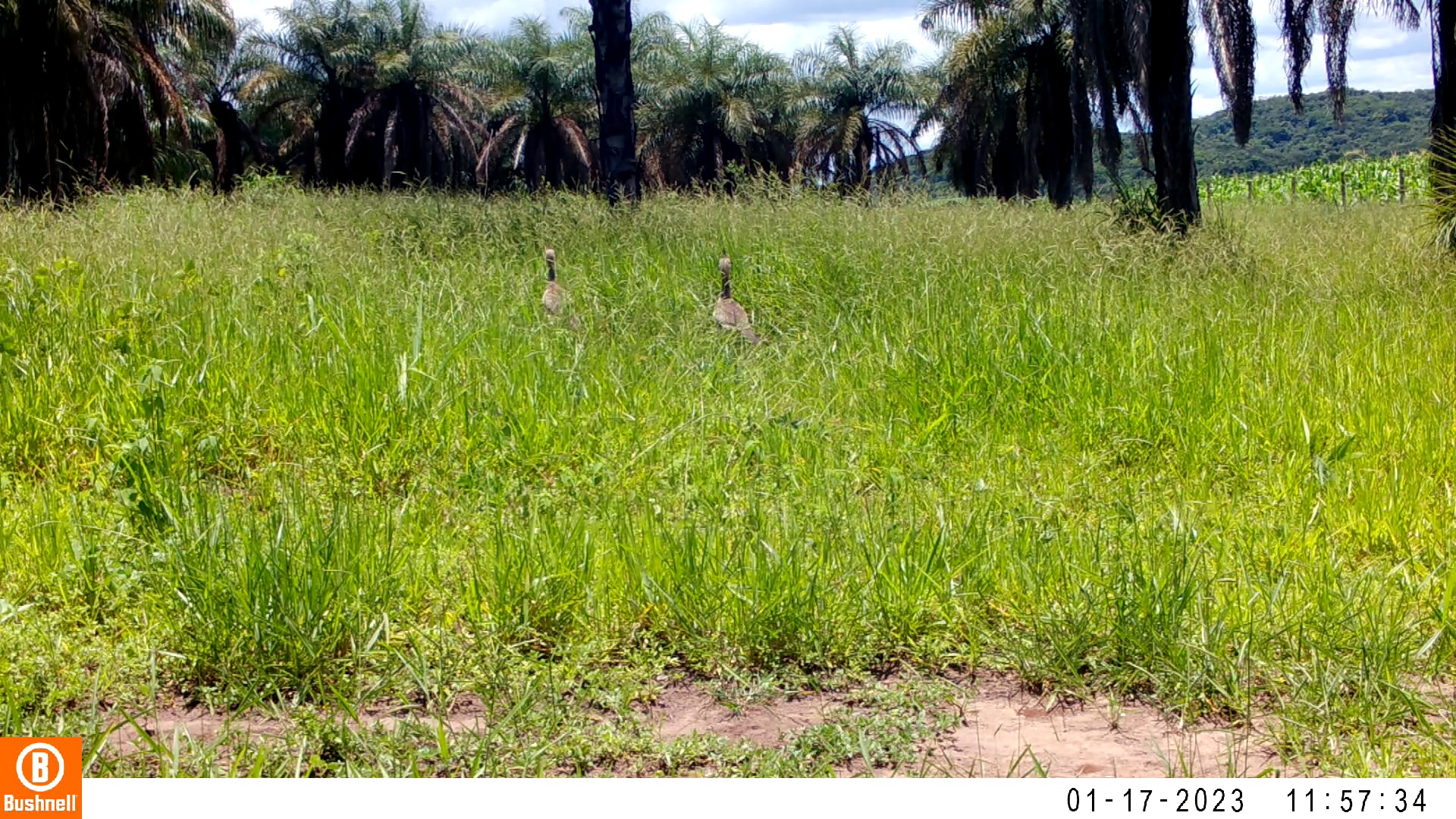

### 01170028-88262-104.jpg

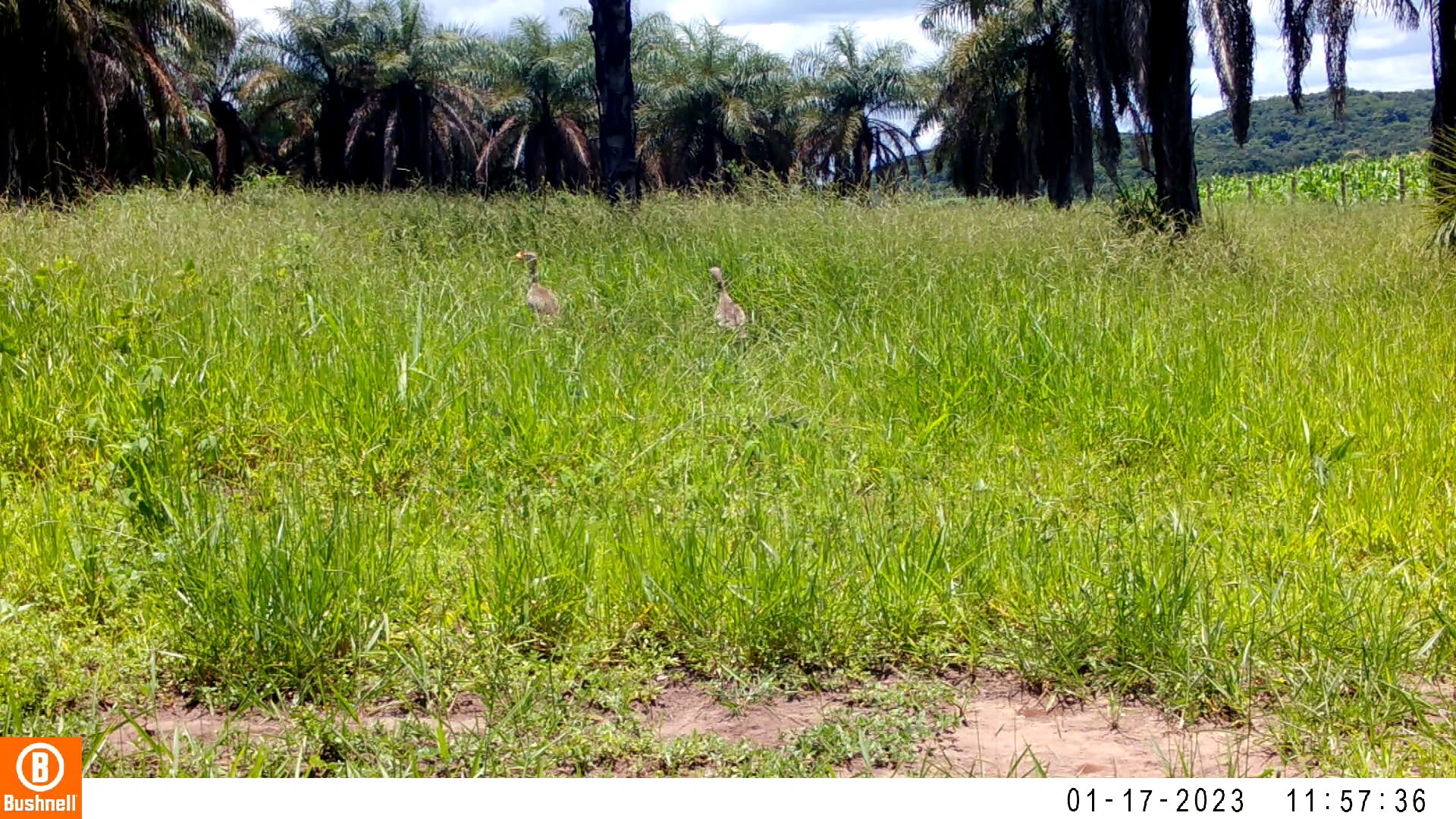

### 01170028-88262-156.jpg

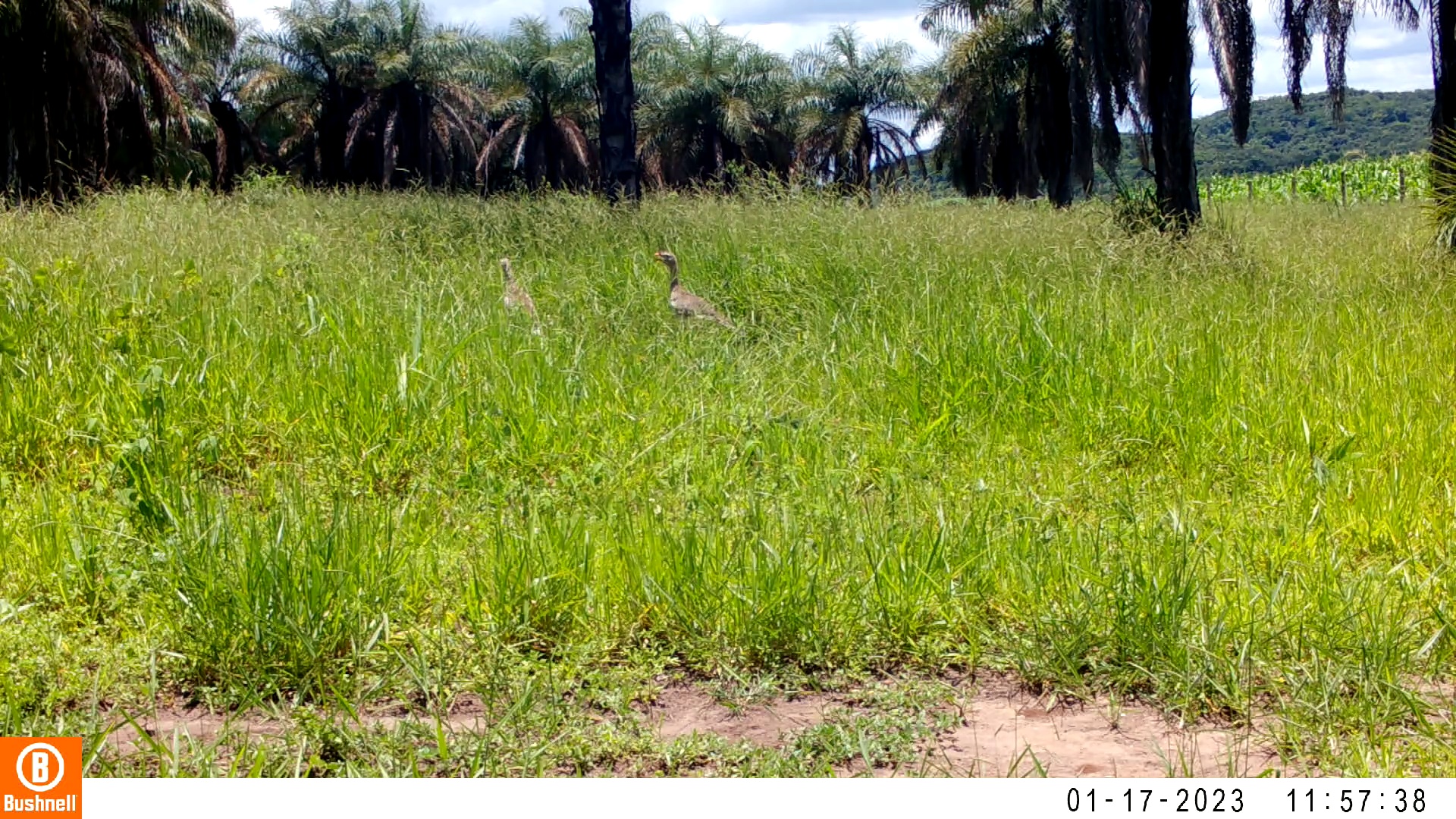

### 01170028-88262-208.jpg

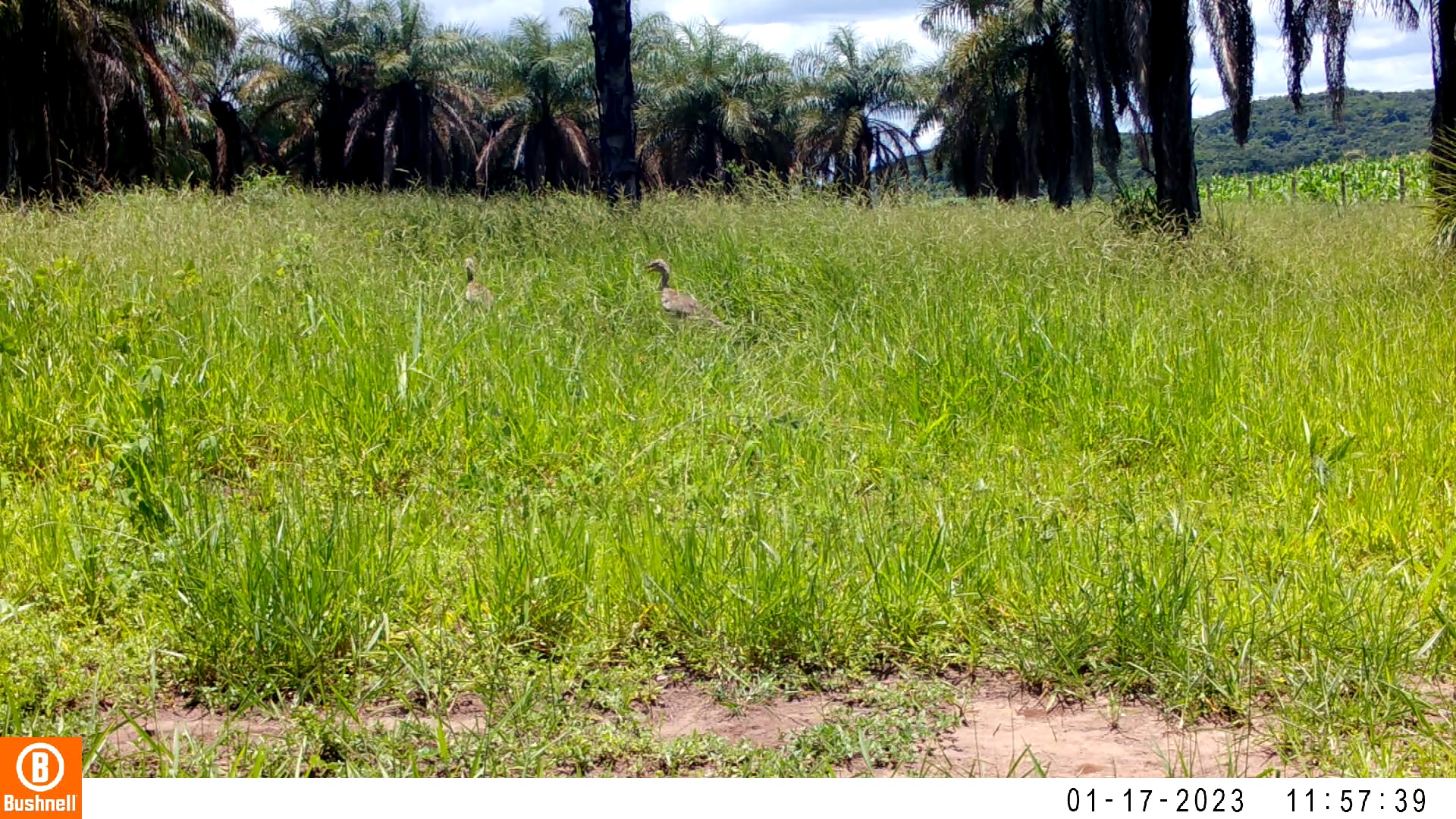

### 01170028-88262-260.jpg

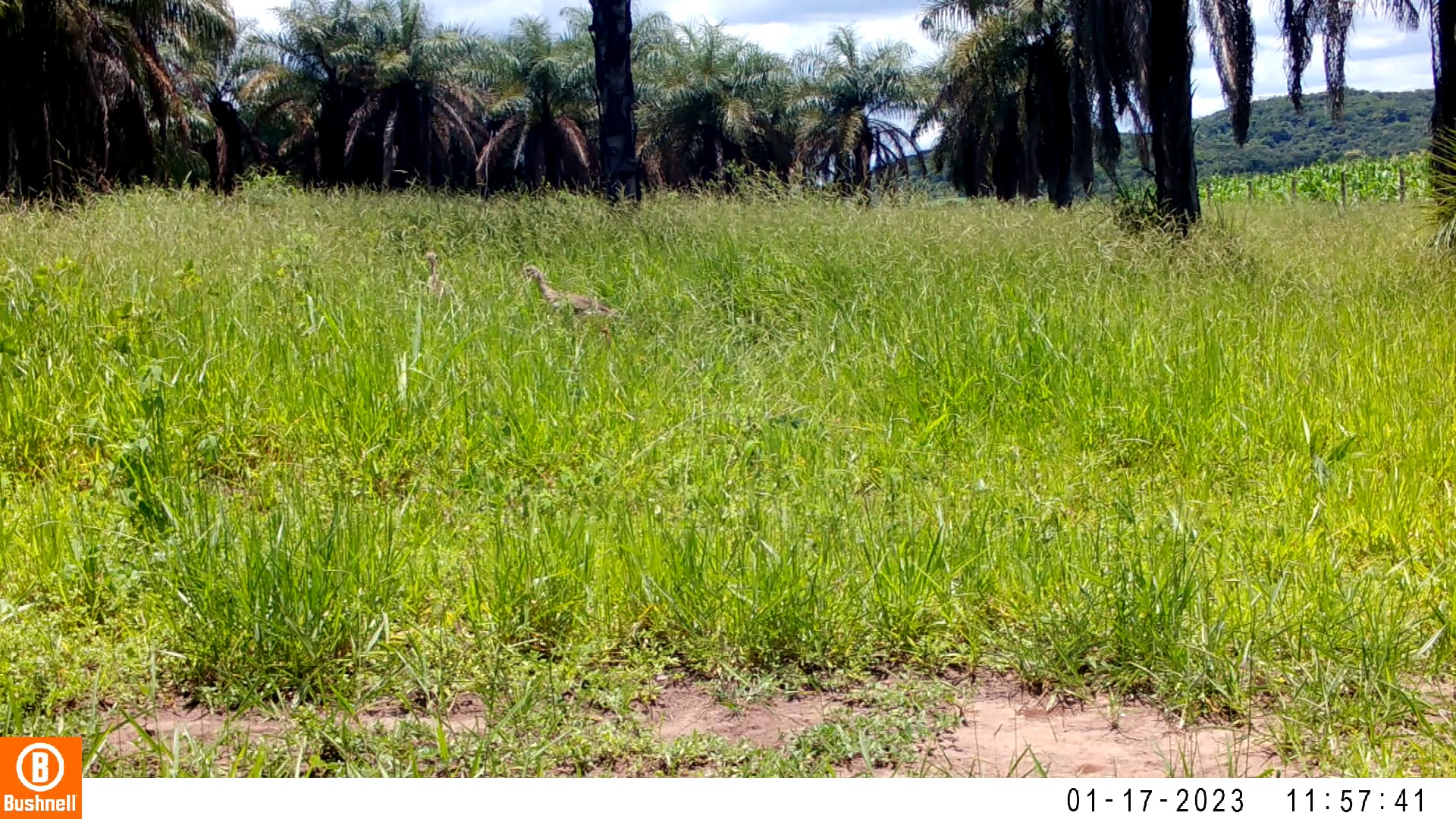

### 01170030-44304-0.jpg

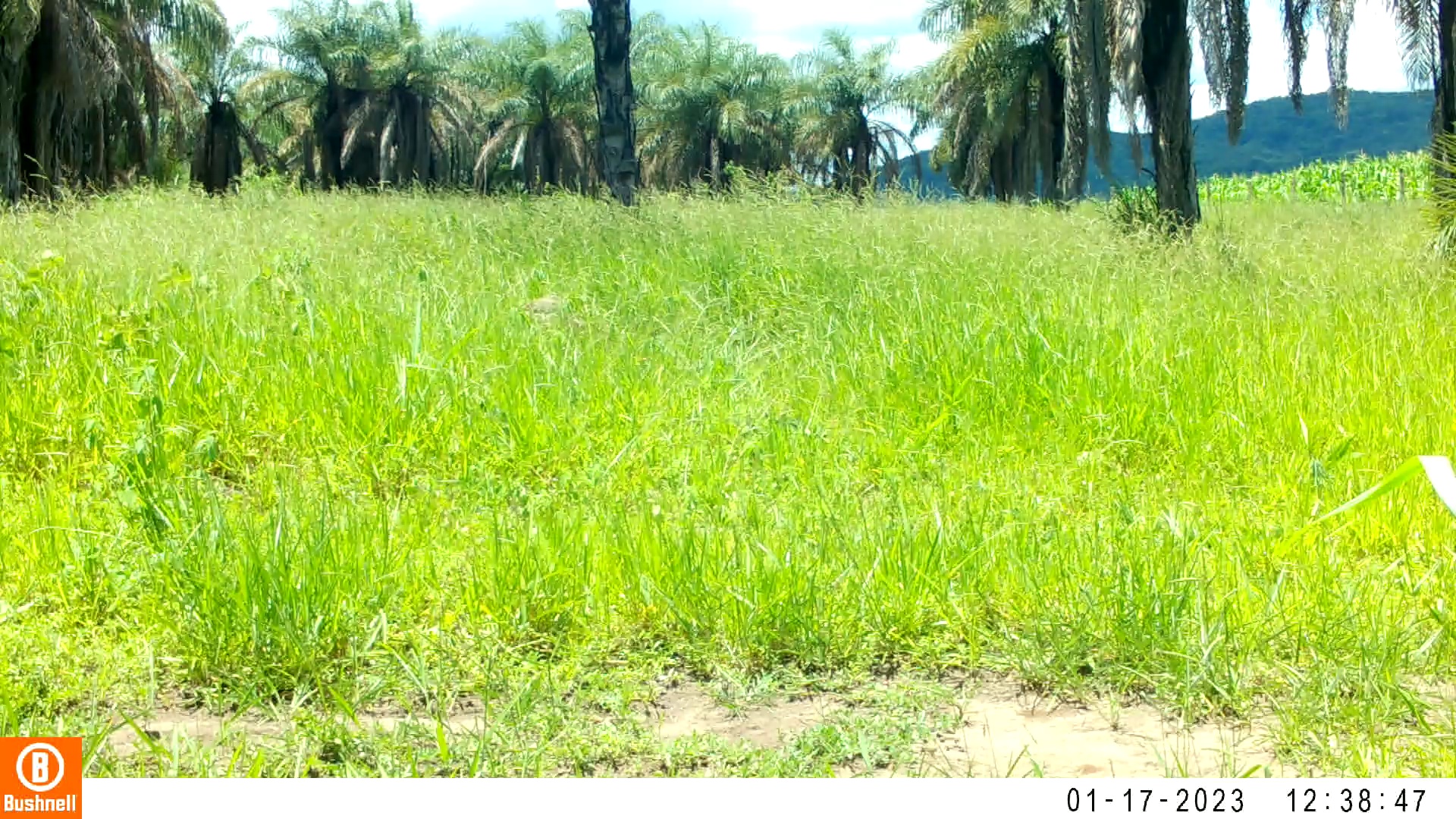

### 01170030-44304-52.jpg

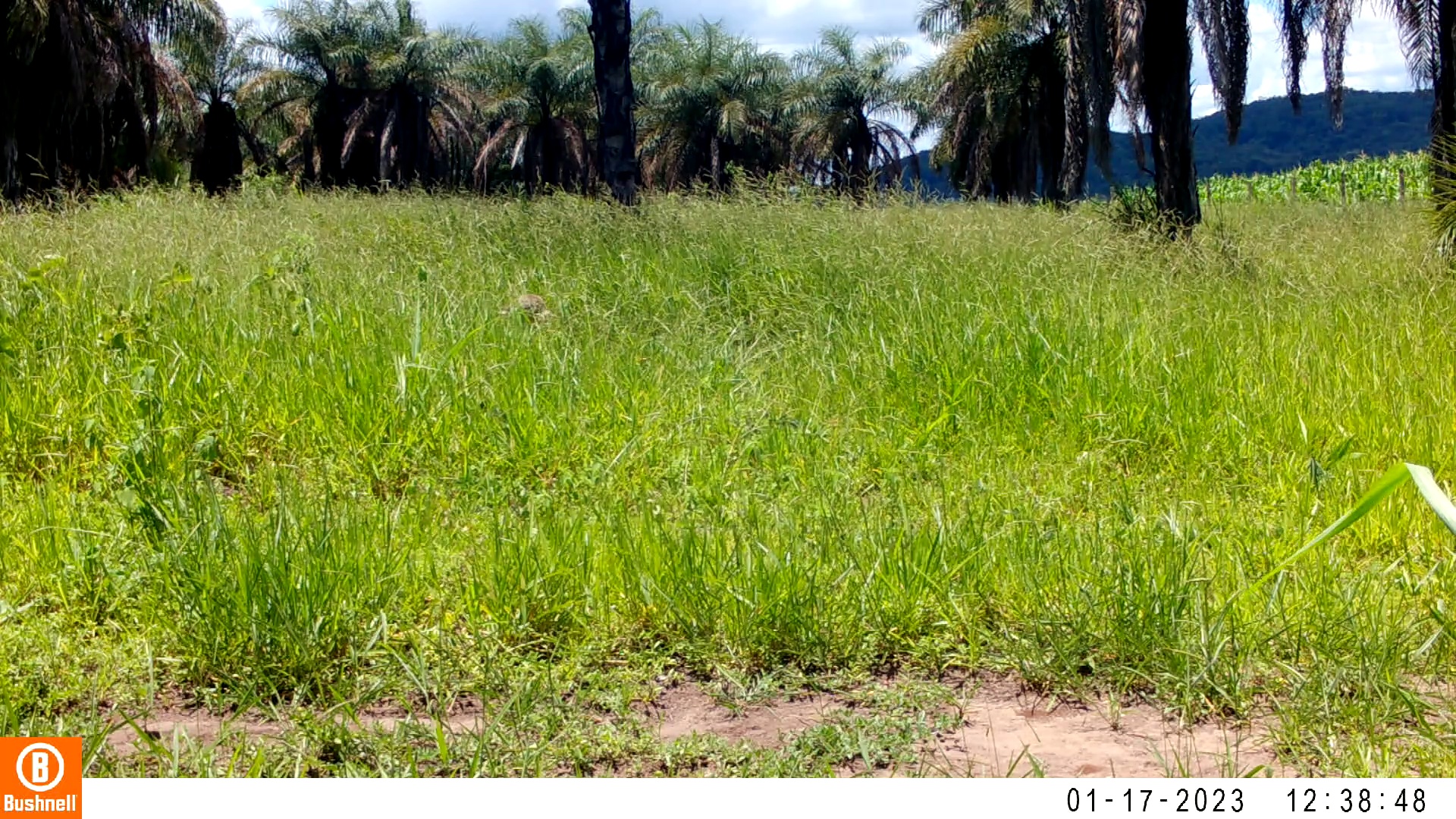

### 01170030-44304-104.jpg

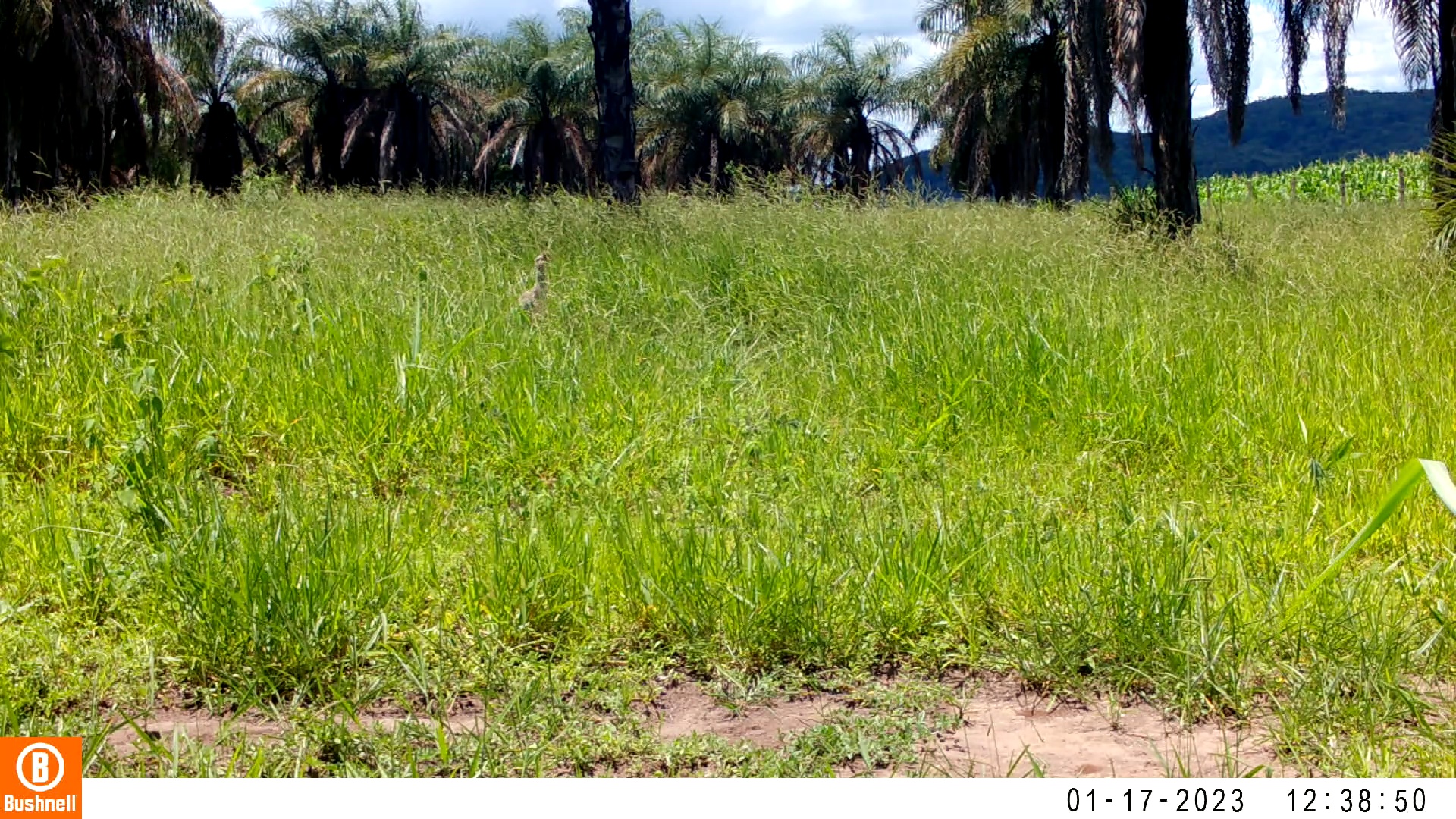

### 01170030-44304-156.jpg

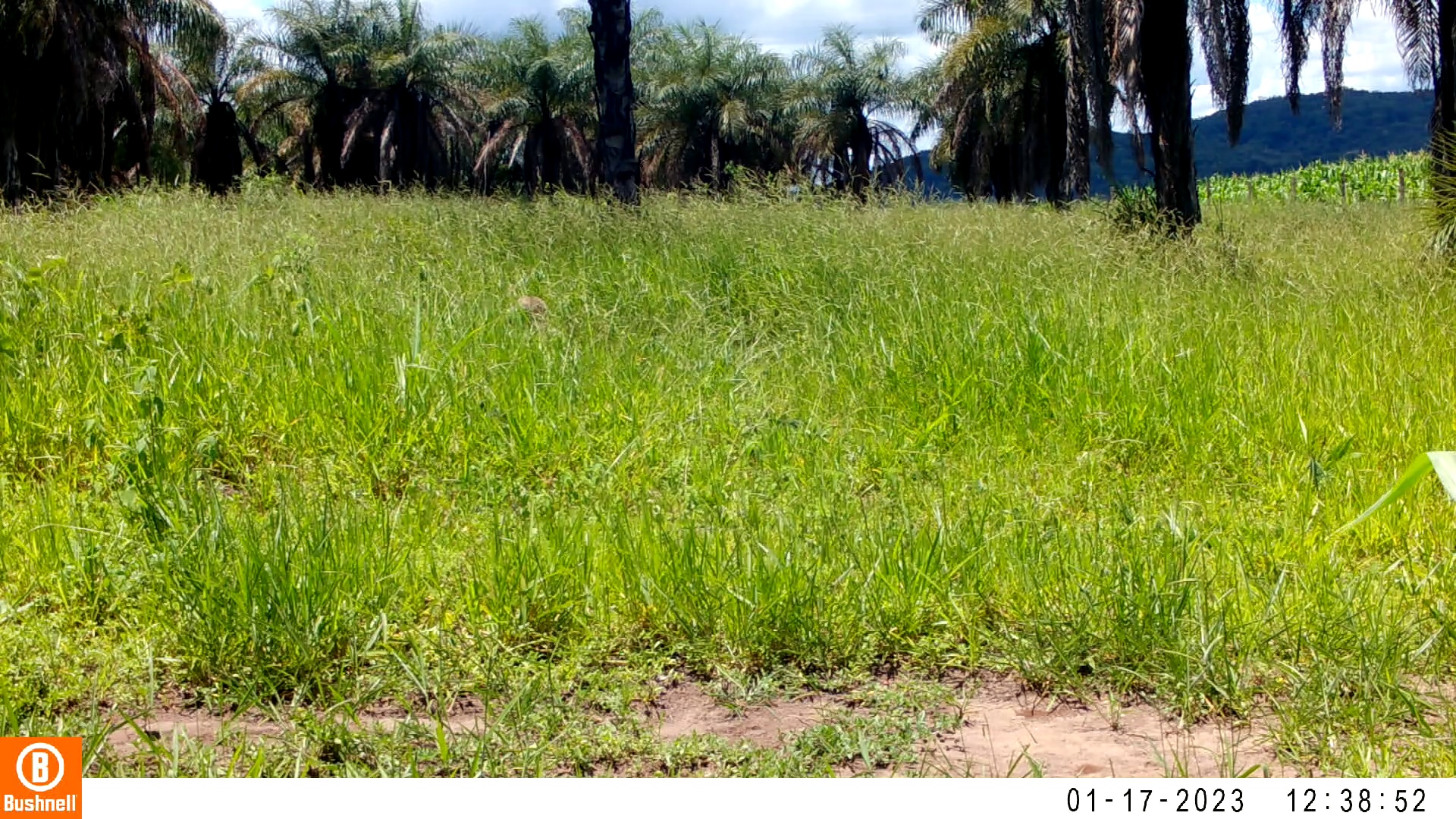

### 01170030-44304-208.jpg

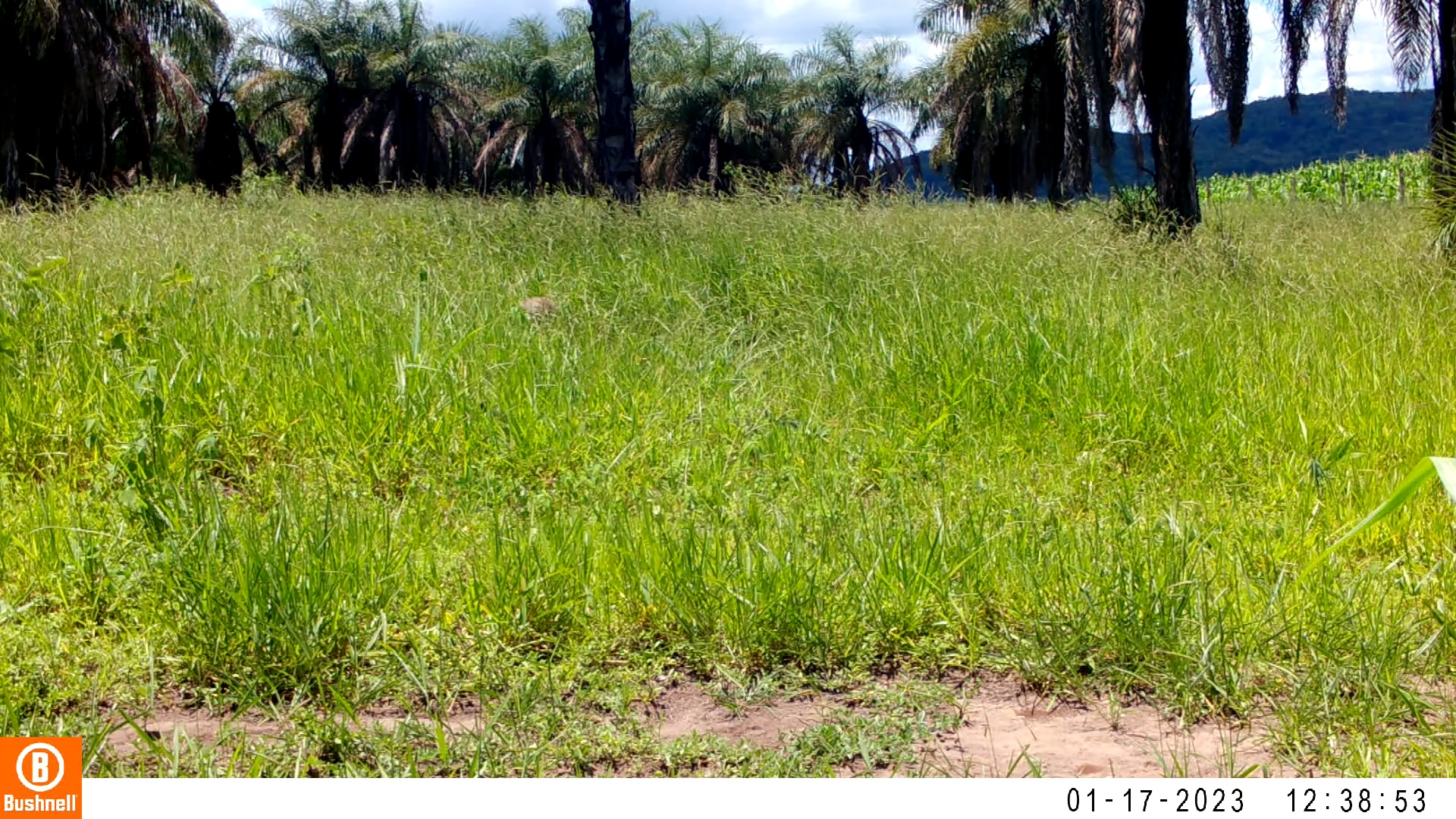

### 01170032-07403-52.jpg

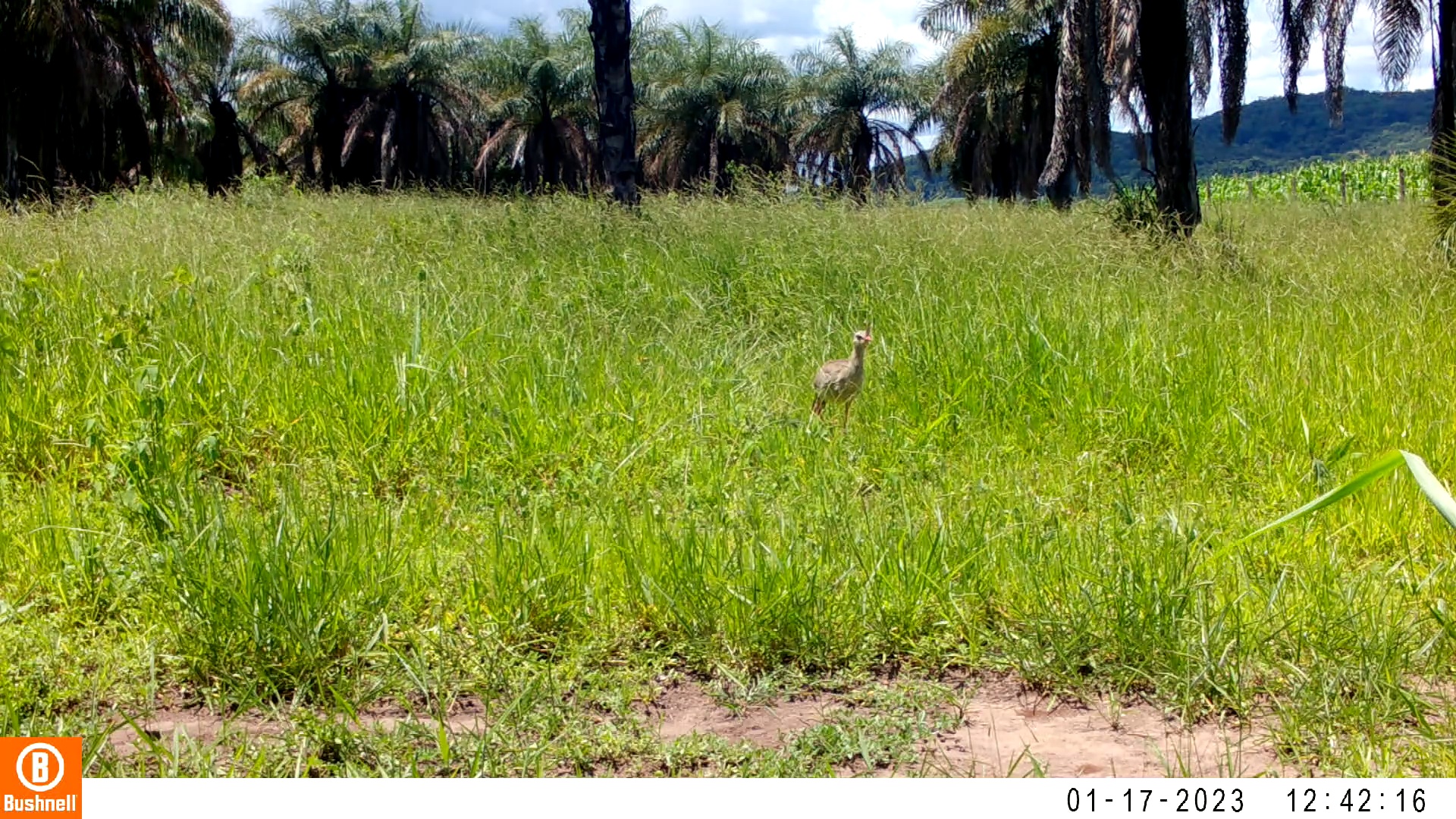

### 01170032-07403-104.jpg

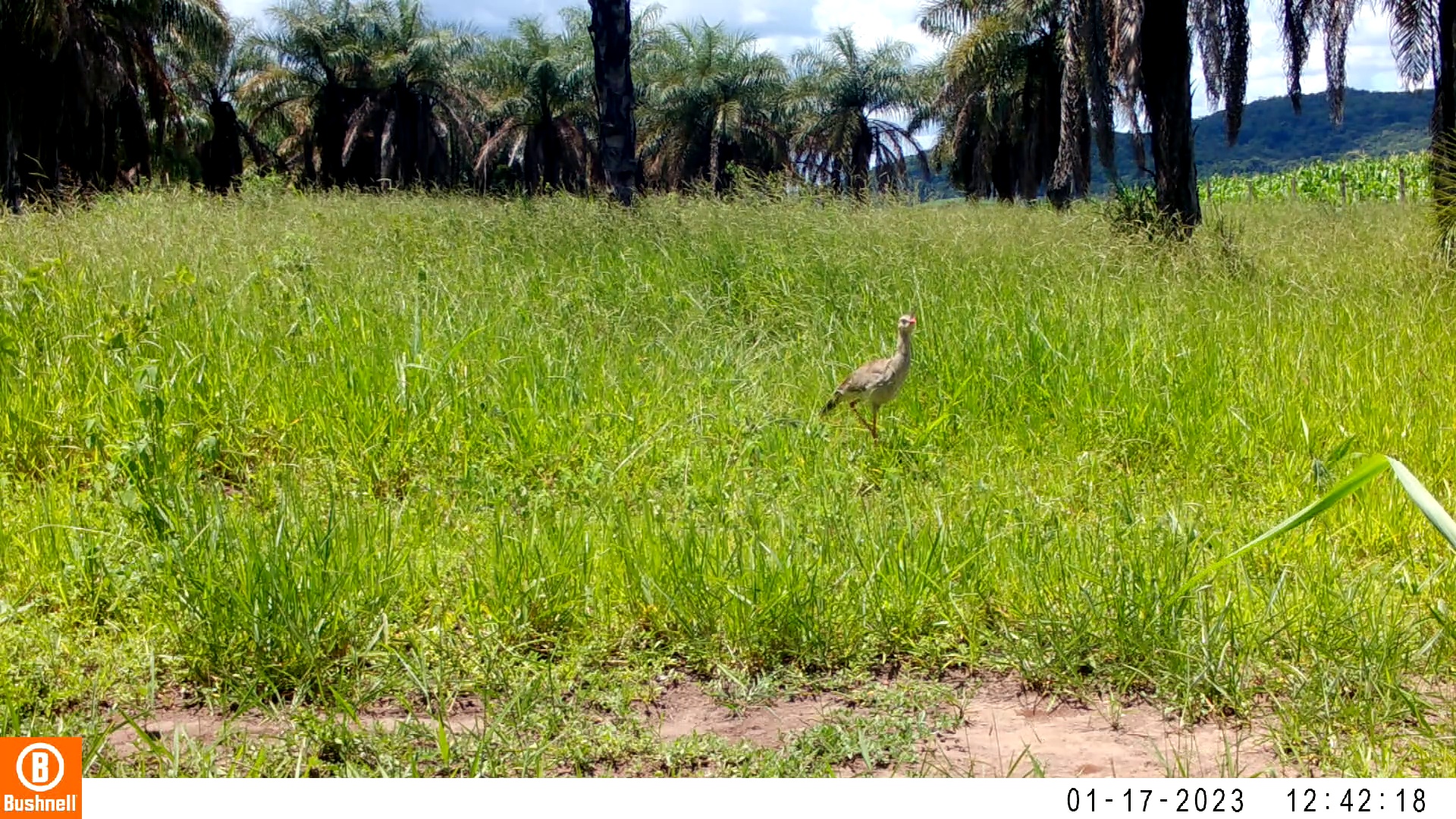

### 01170032-07403-156.jpg

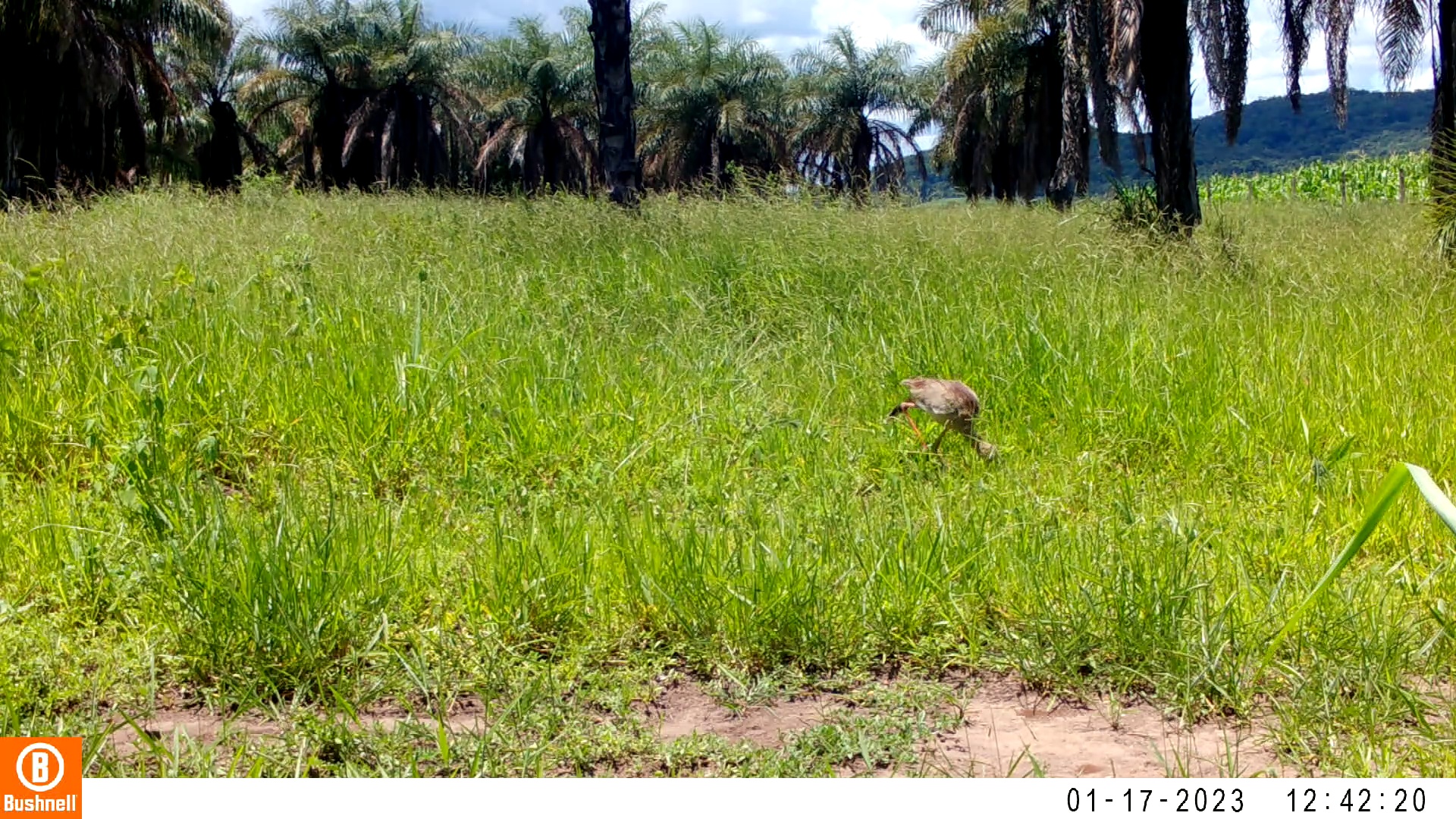

### 01170032-07403-208.jpg

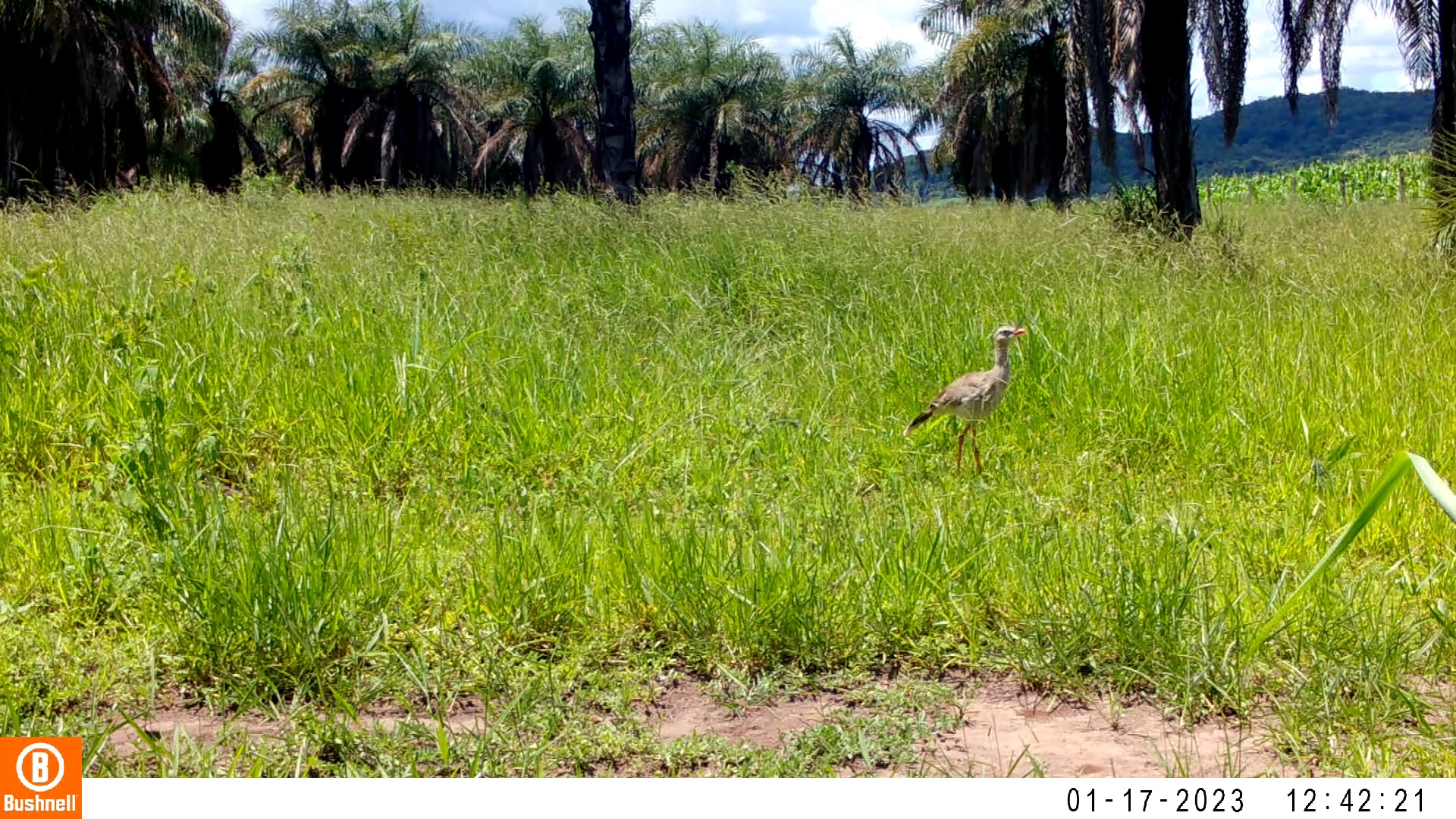

### 01170032-07403-260.jpg

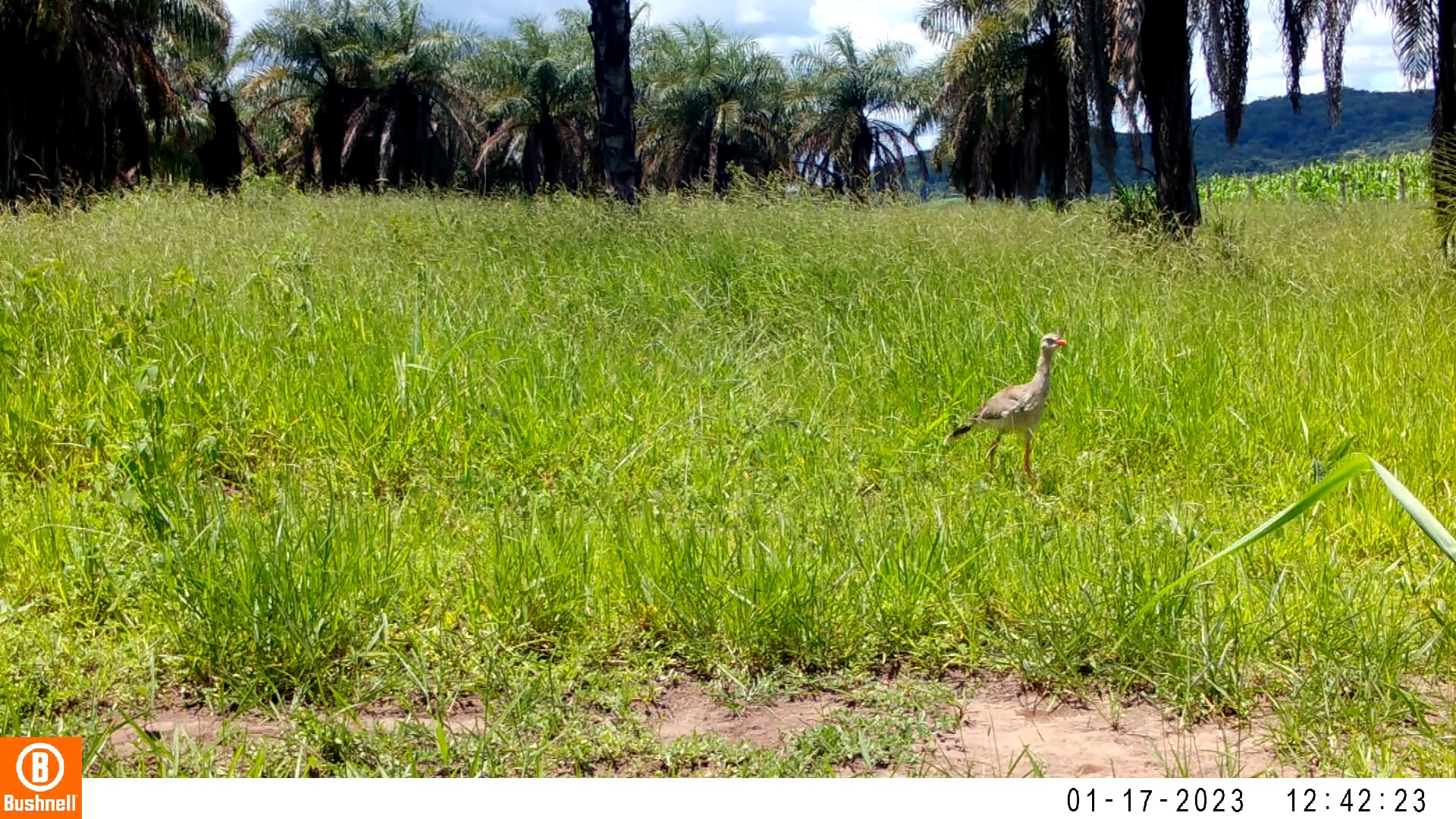

### 01170074-33303-0.jpg

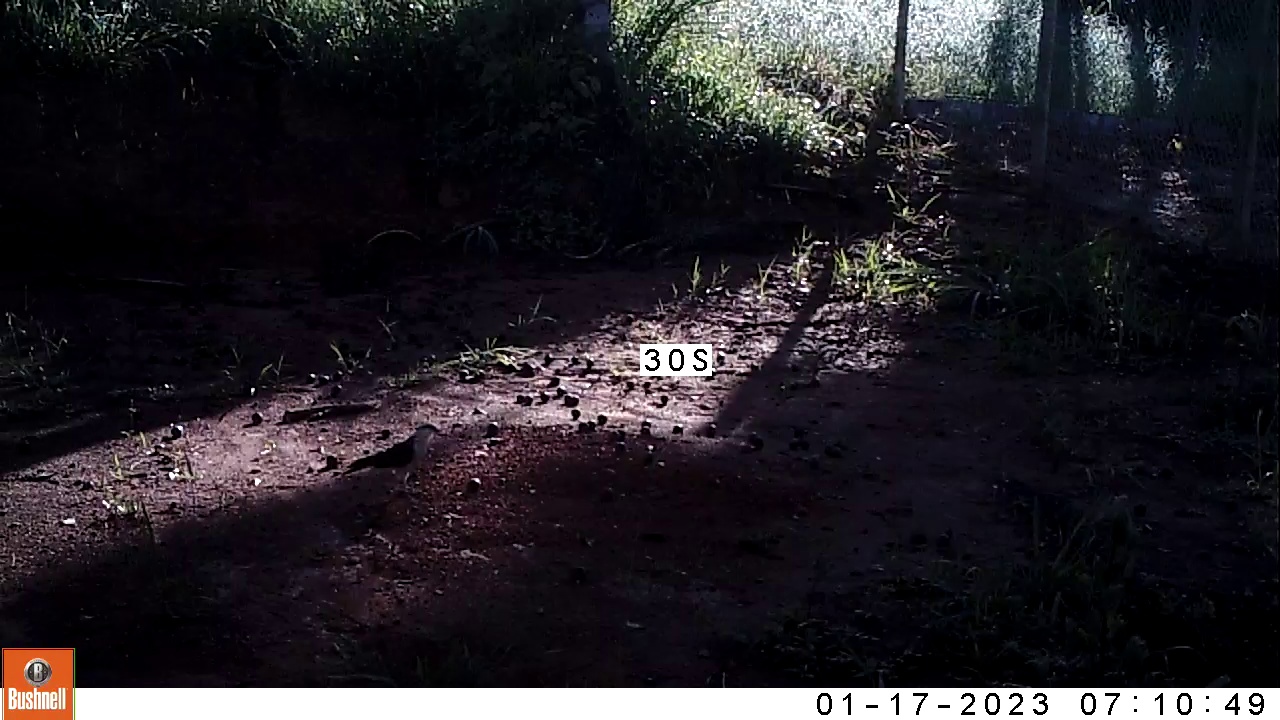

### 01170074-33303-300.jpg

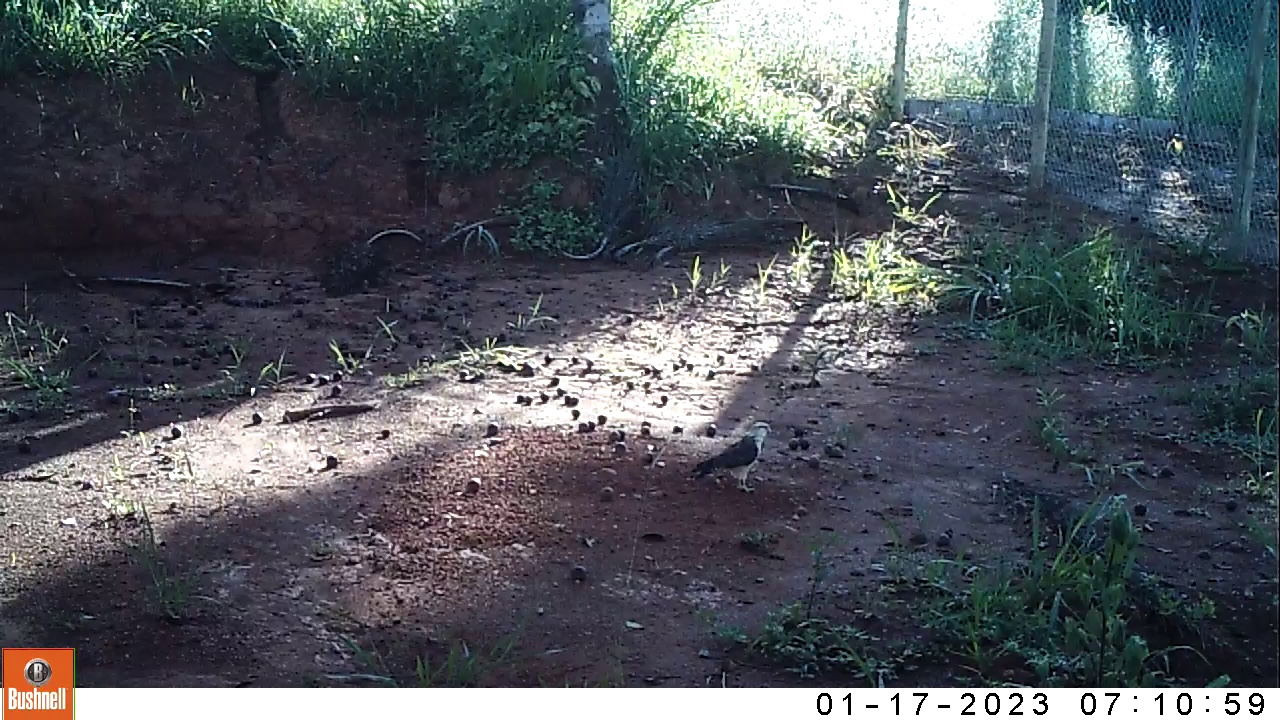

### 01170074-33303-450.jpg

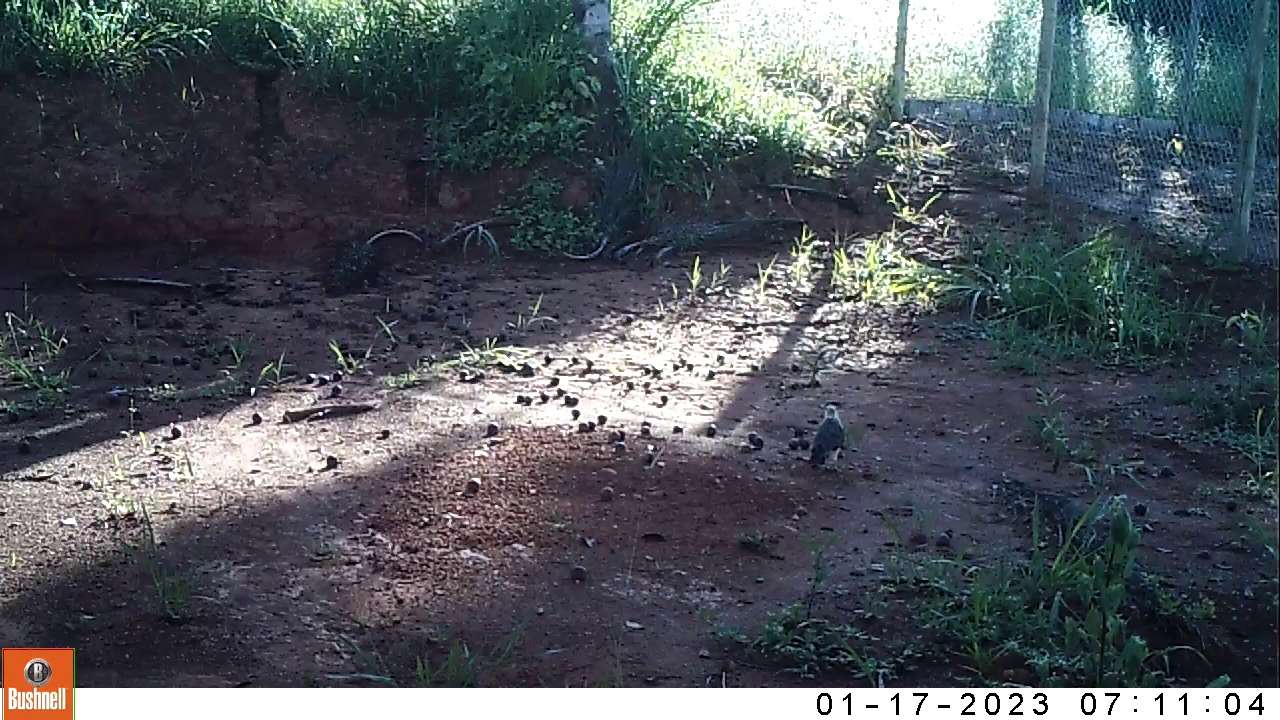

### 01170074-33303-600.jpg

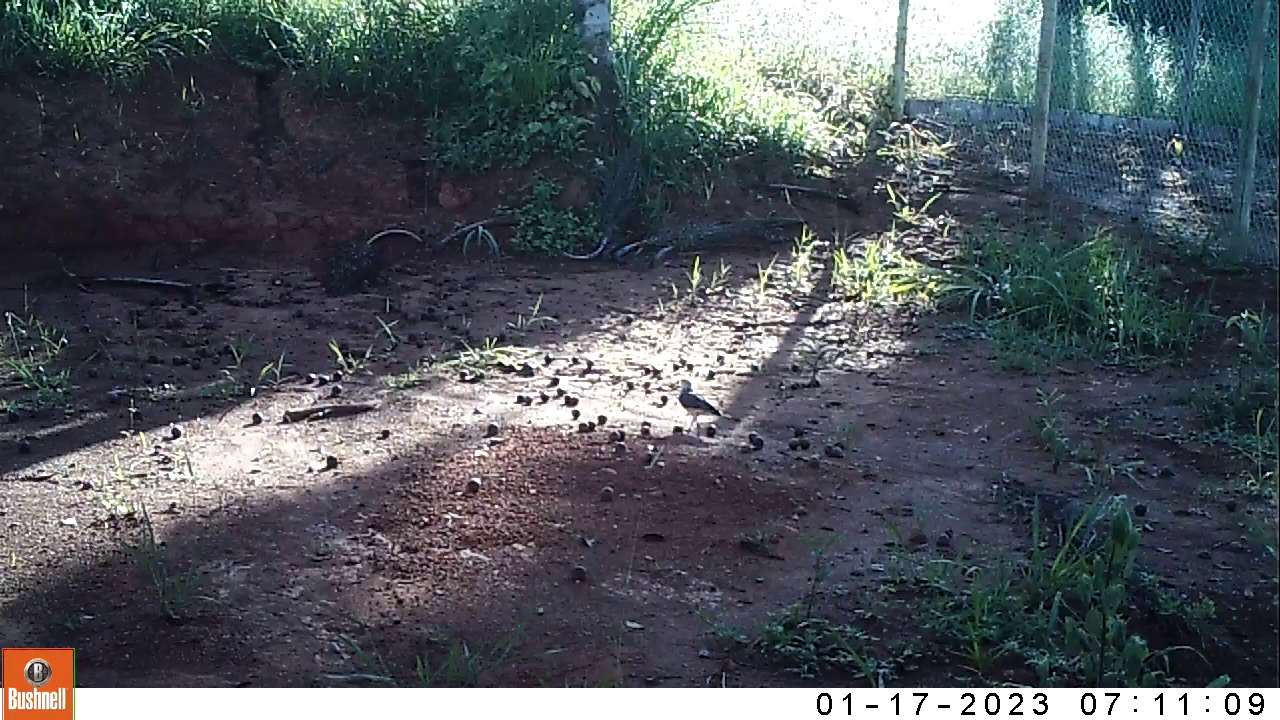

### 01170074-33303-750.jpg

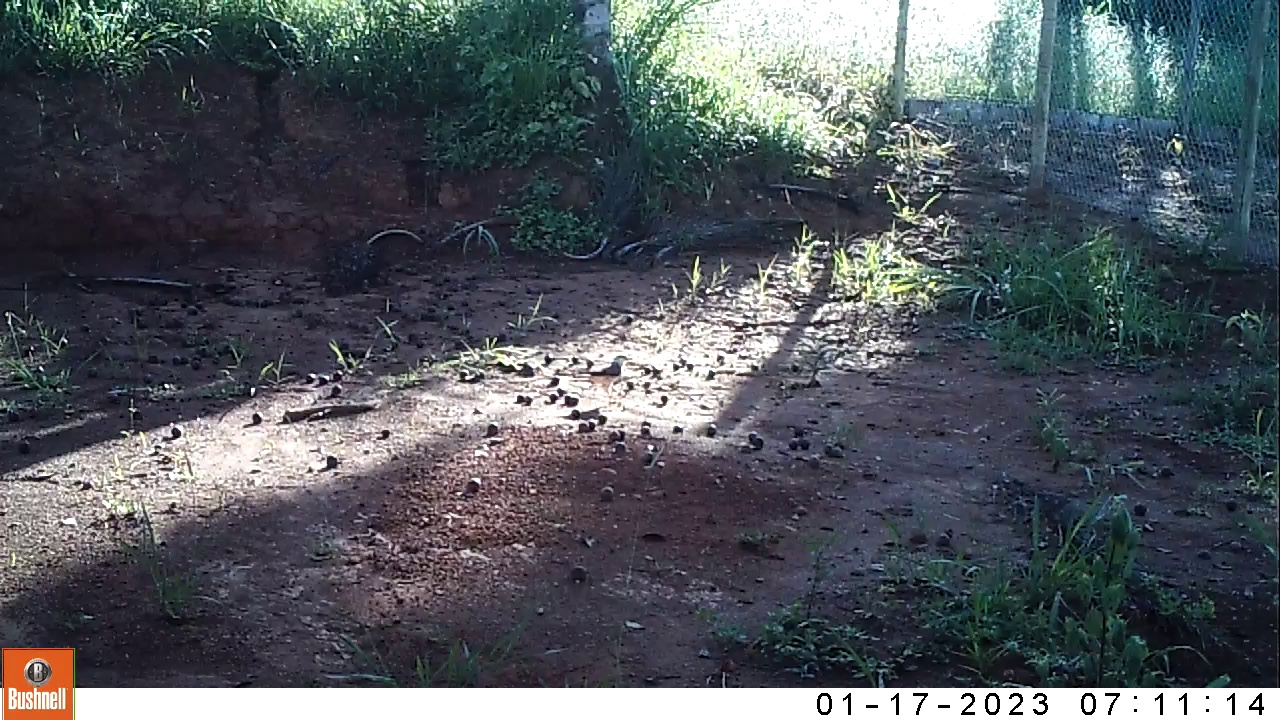

### 11060006-24362-0.jpg

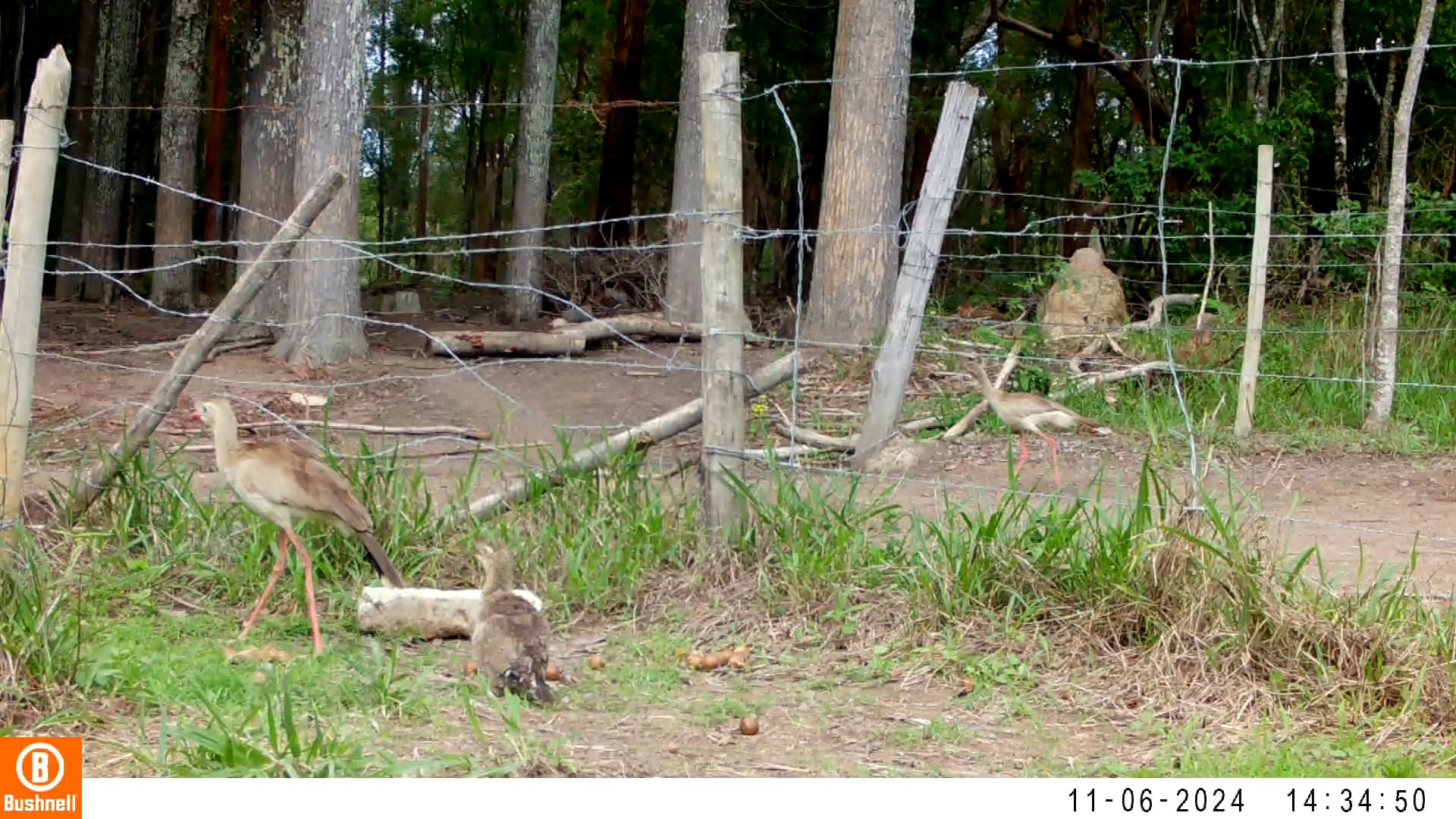

### 11060006-24362-302.jpg

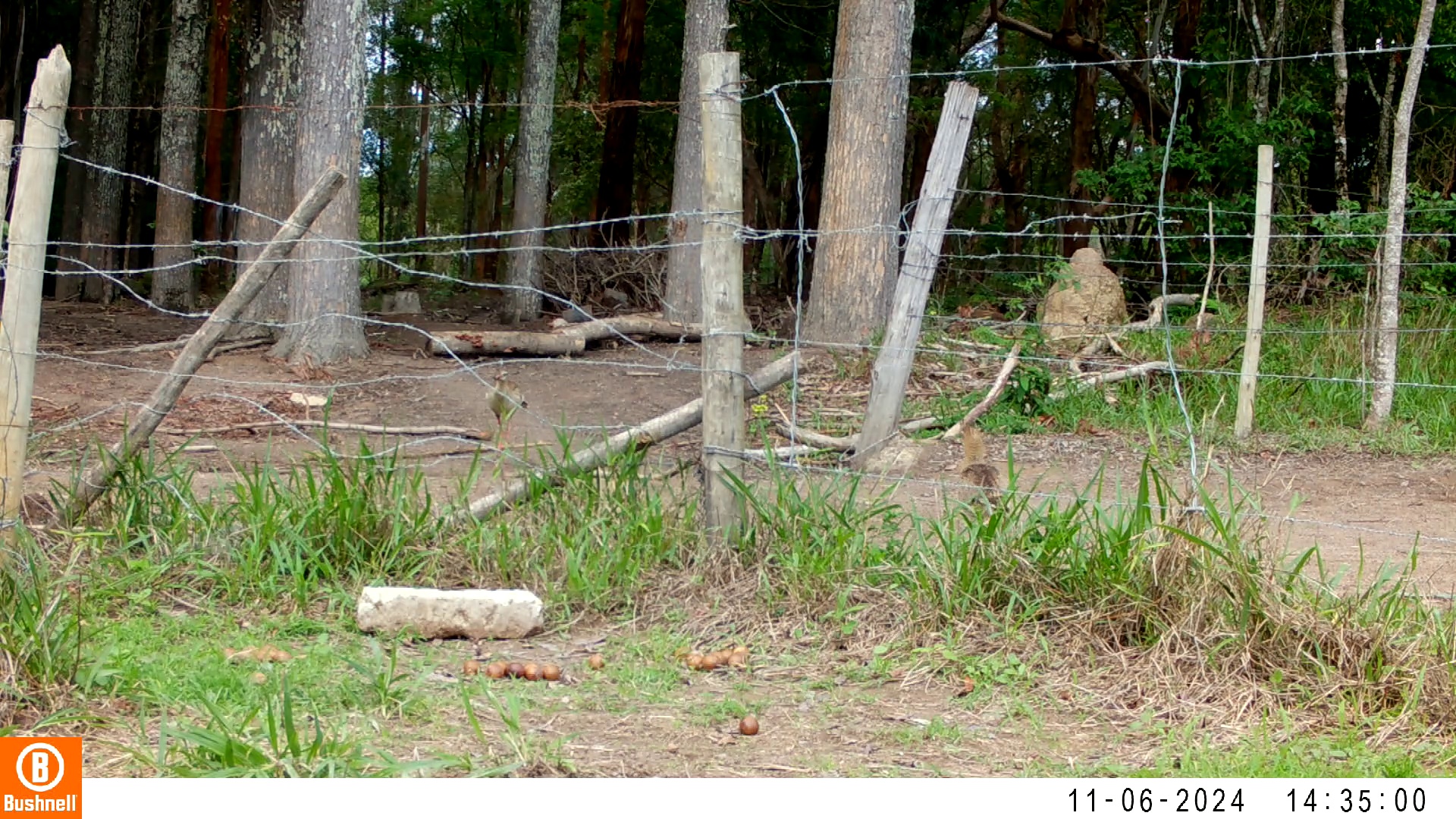

### 11060006-24362-604.jpg

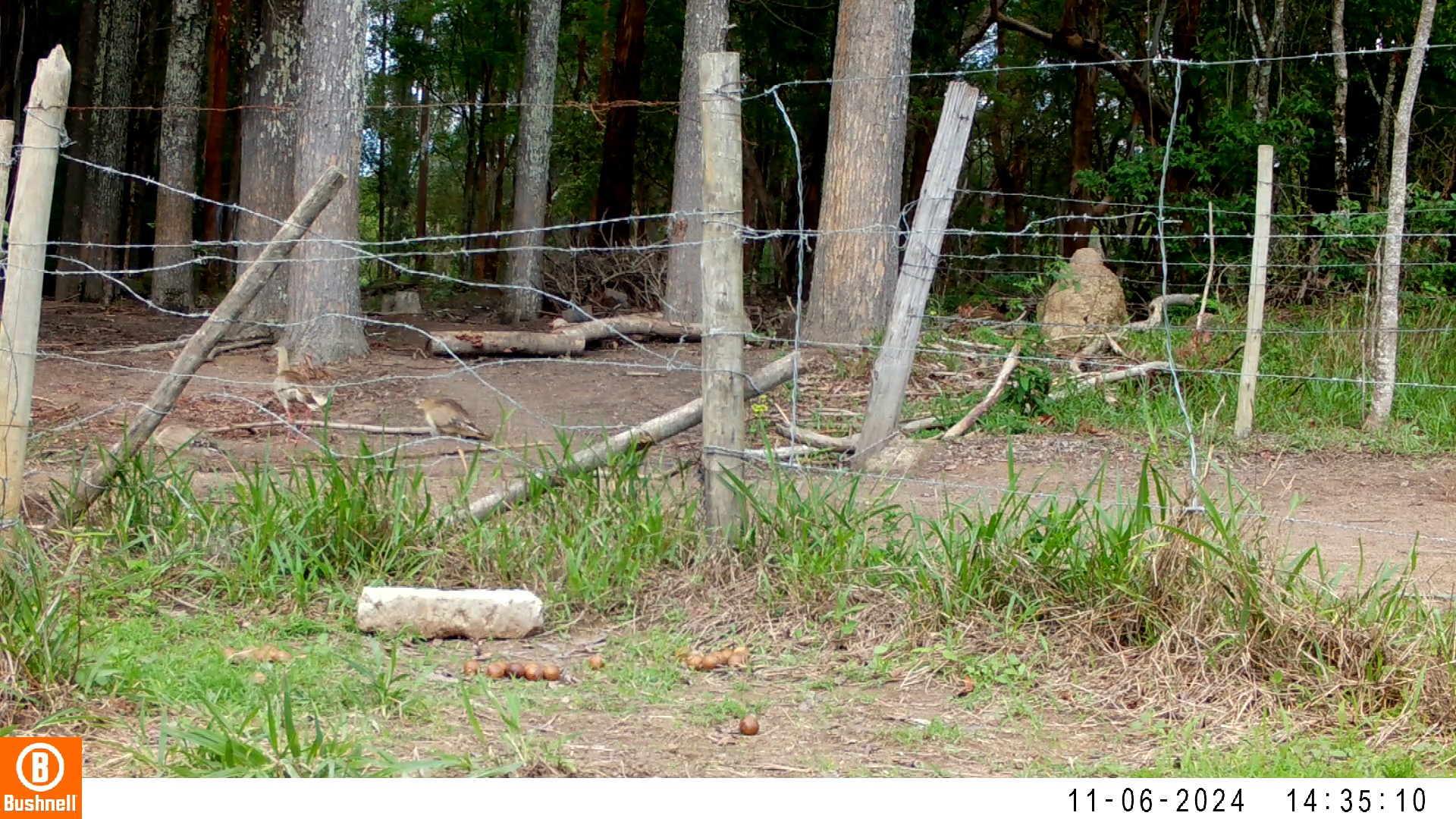

### 11060006-24362-906.jpg

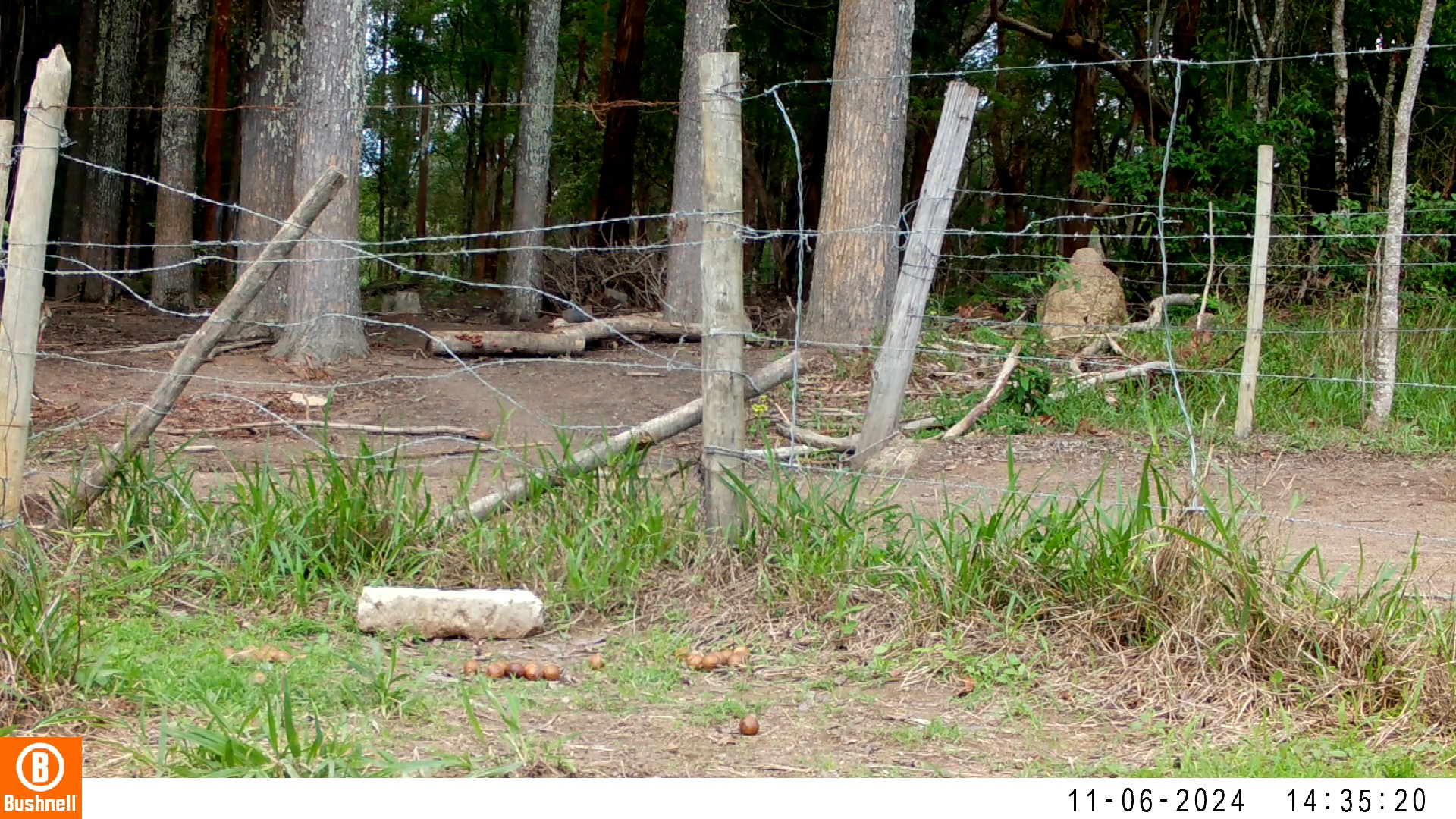

### 11060006-24362-1208.jpg

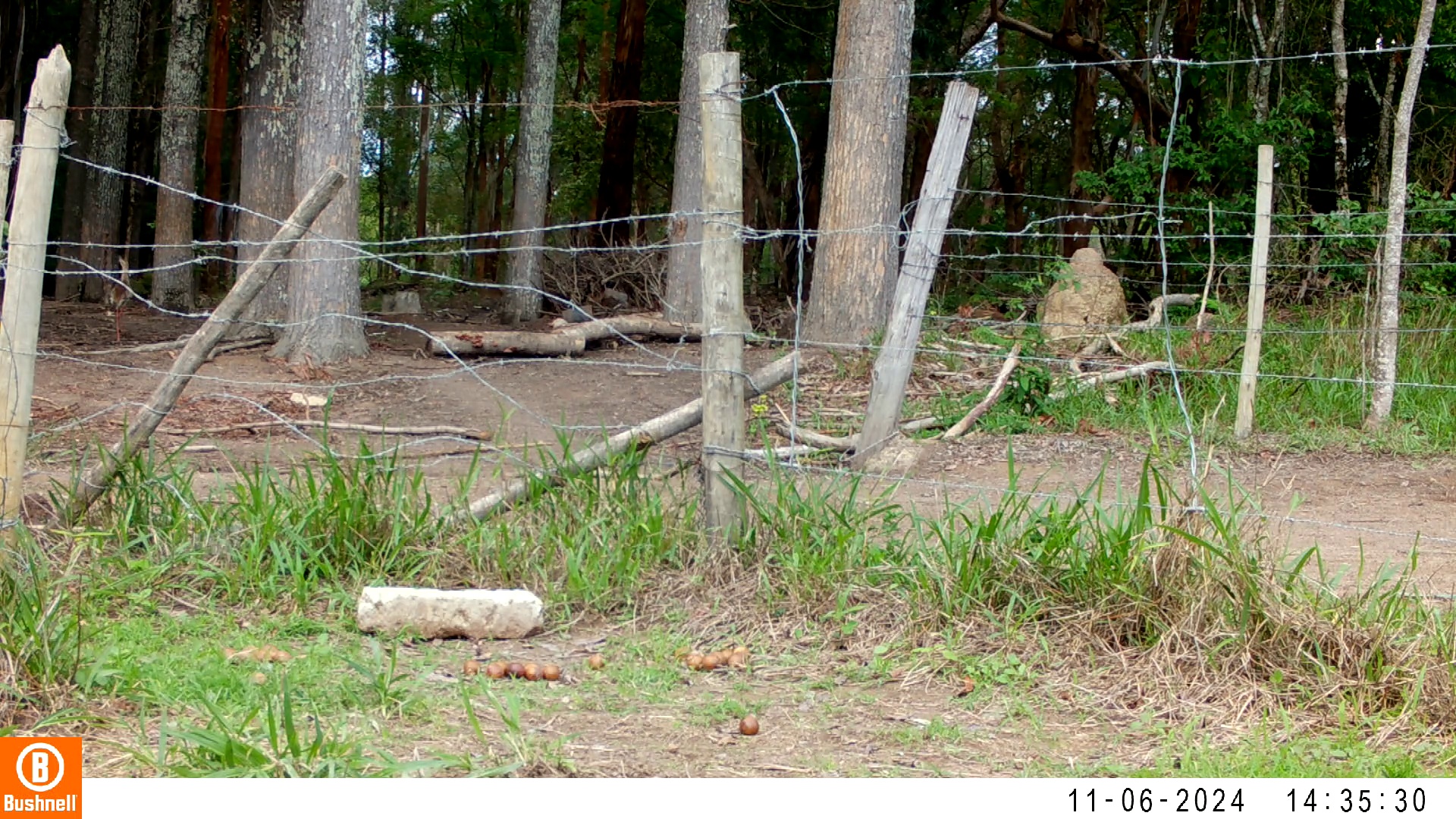

### 11060006-24362-1510.jpg

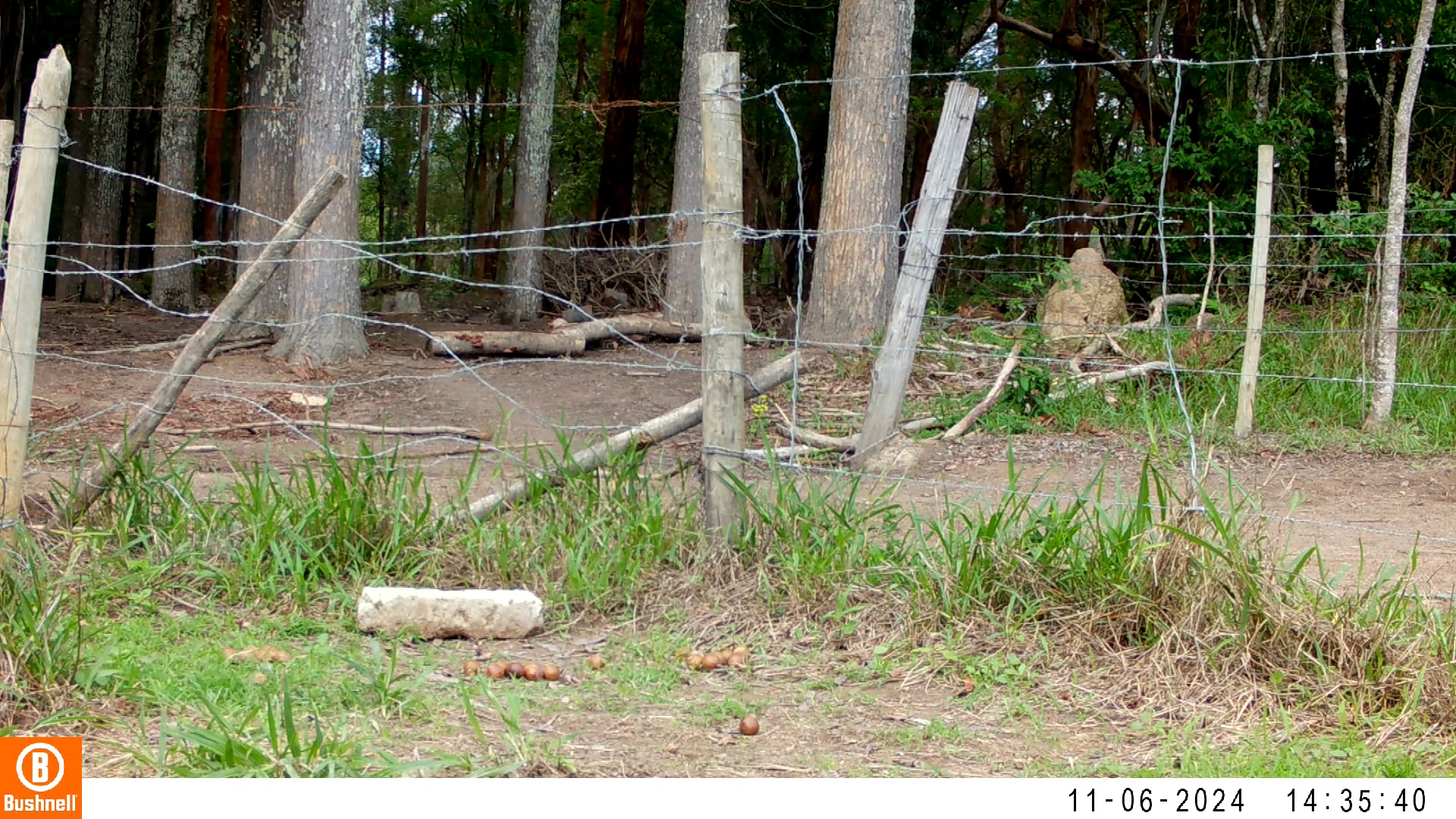

### 11080003-20734-0.jpg

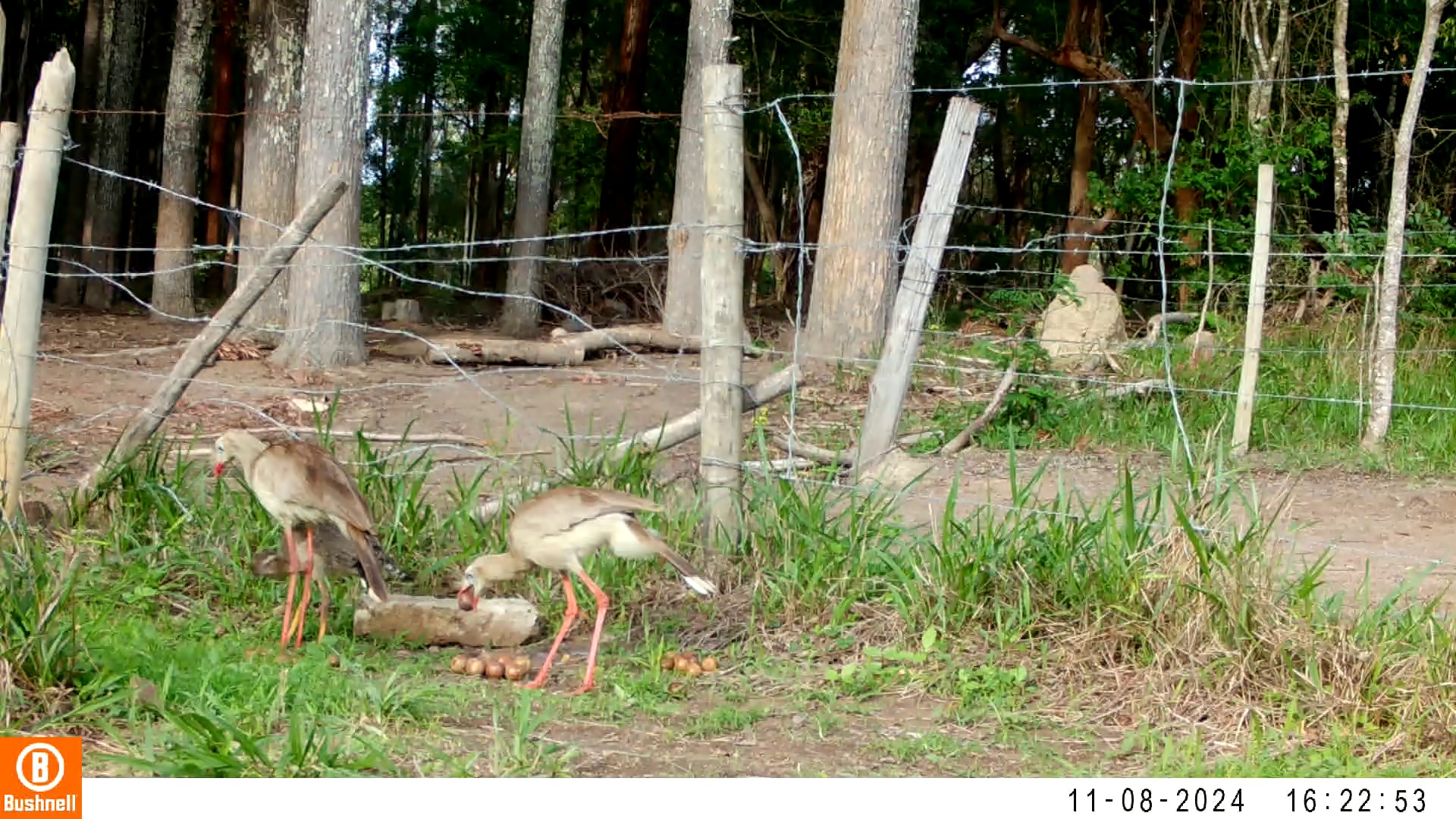

### 11080003-20734-906.jpg

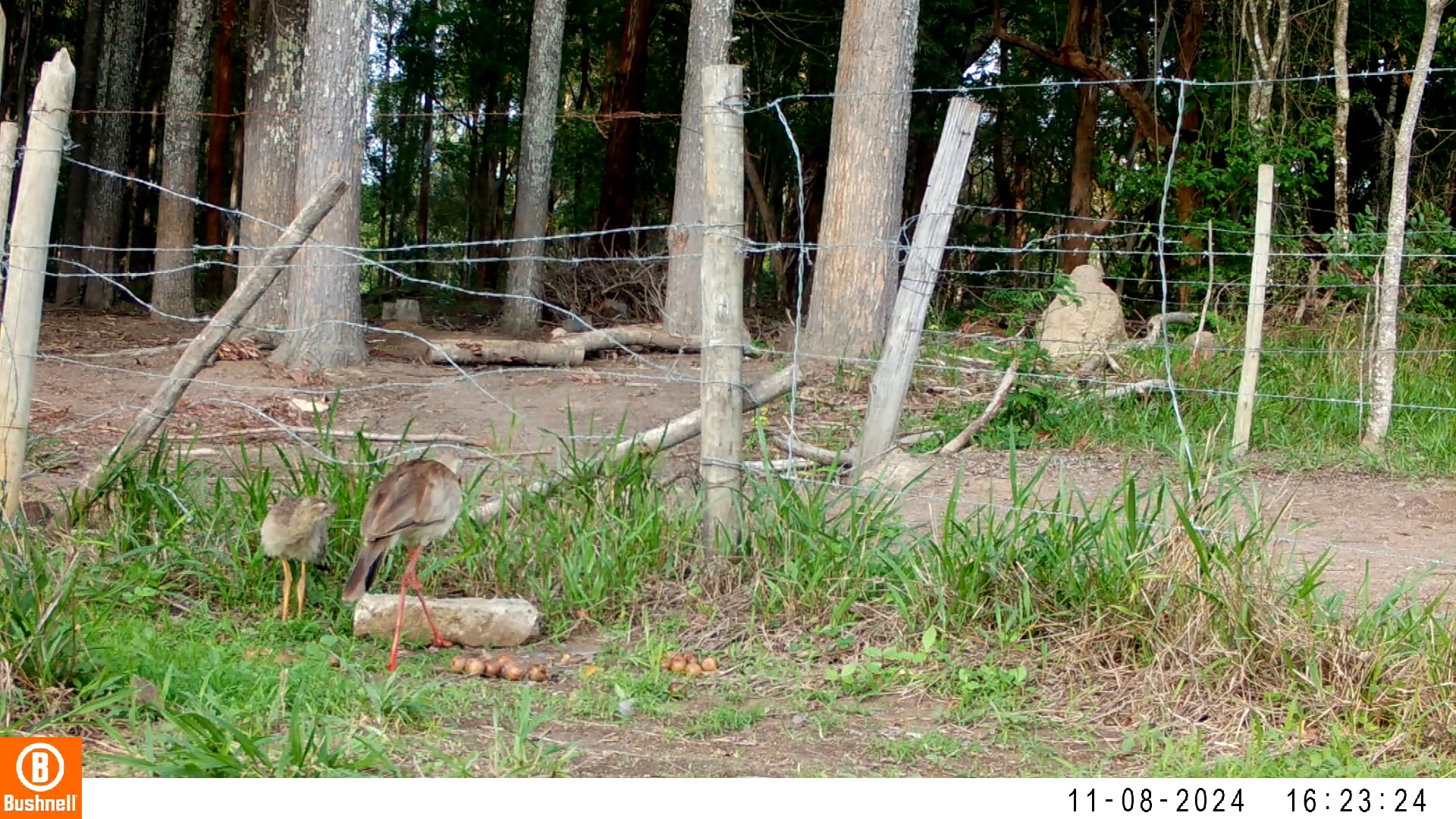

### 11080003-20734-1208.jpg

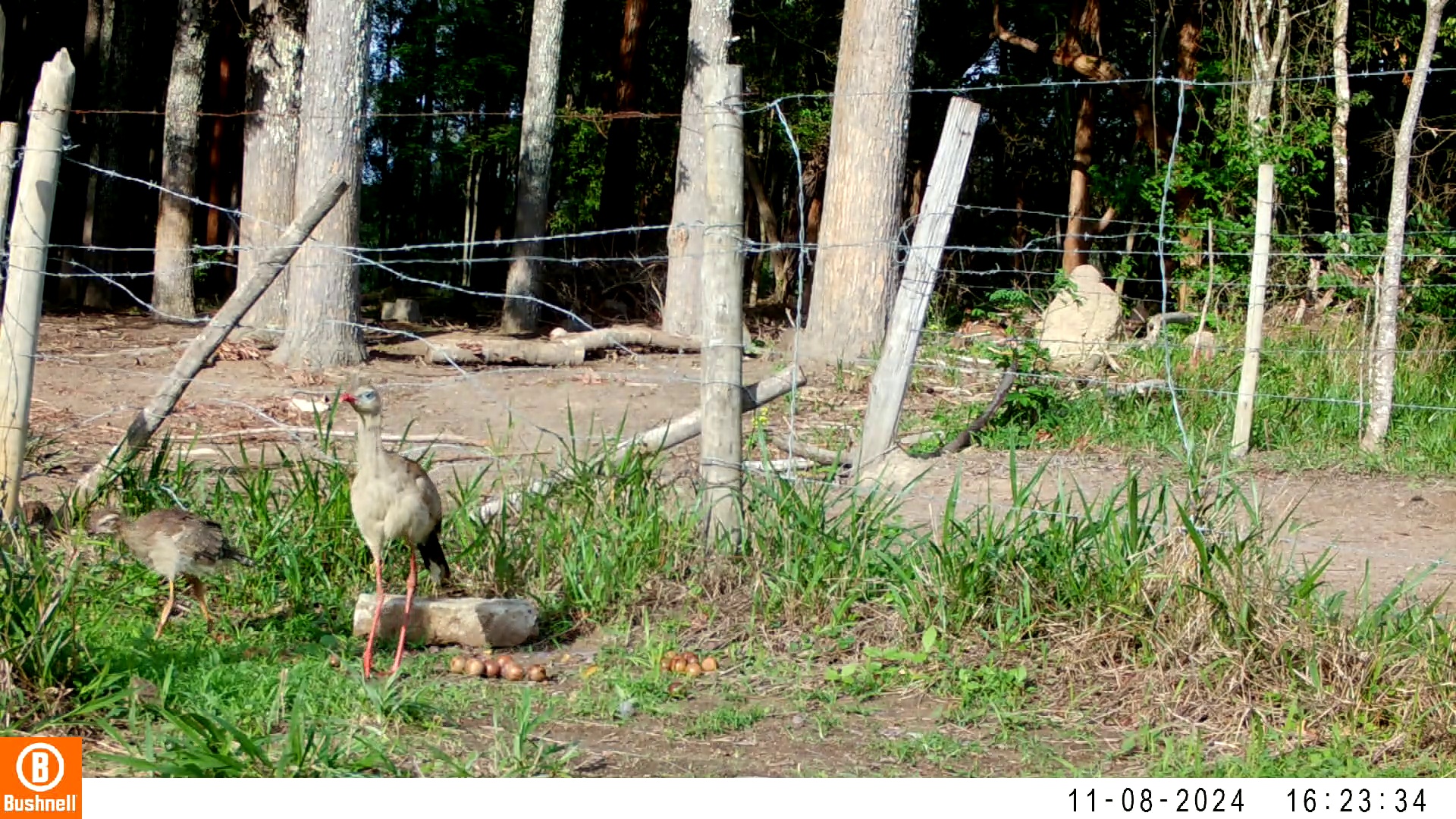

### 11080003-20734-1510.jpg

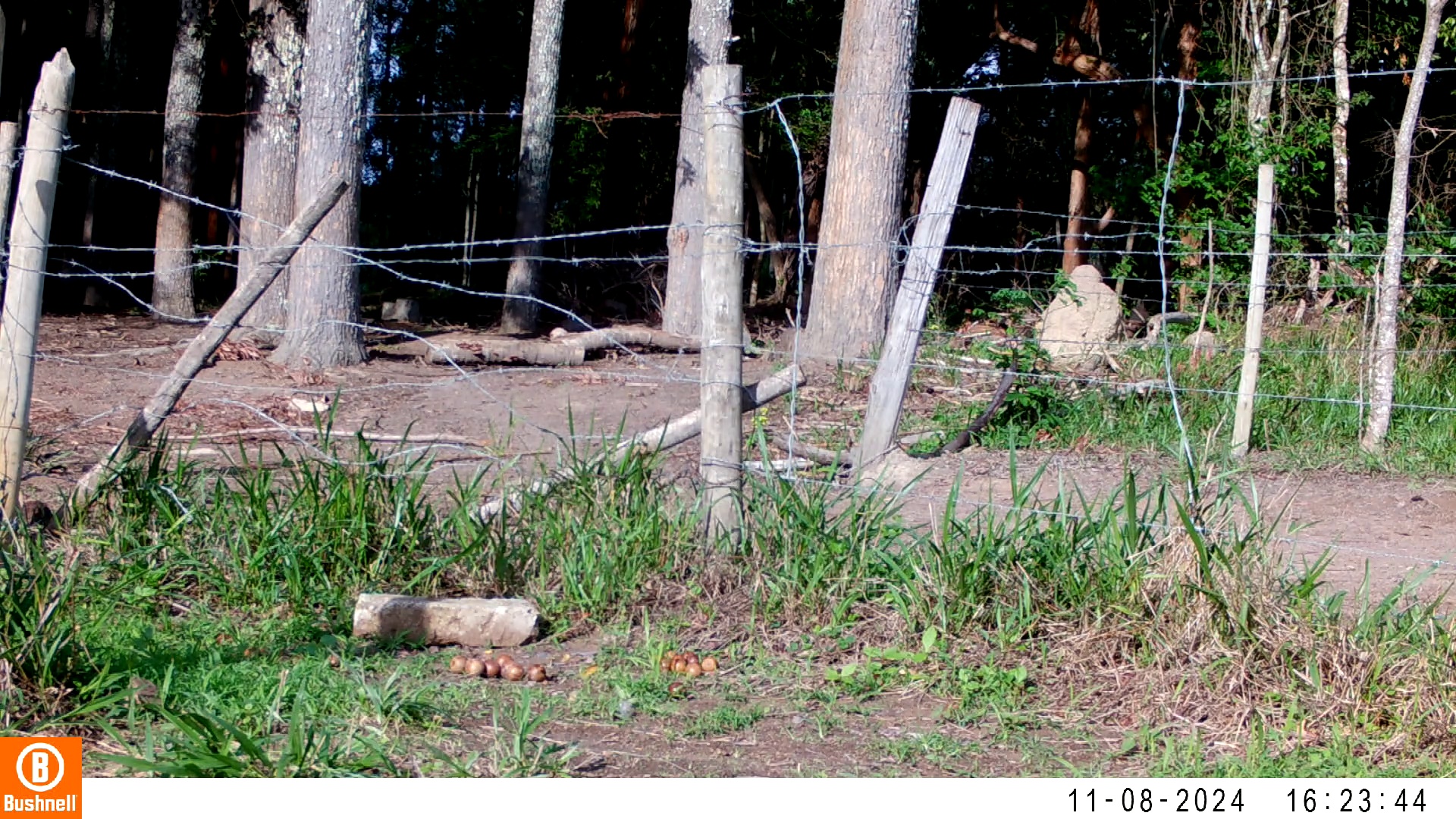

### 11160031-73776-906.jpg

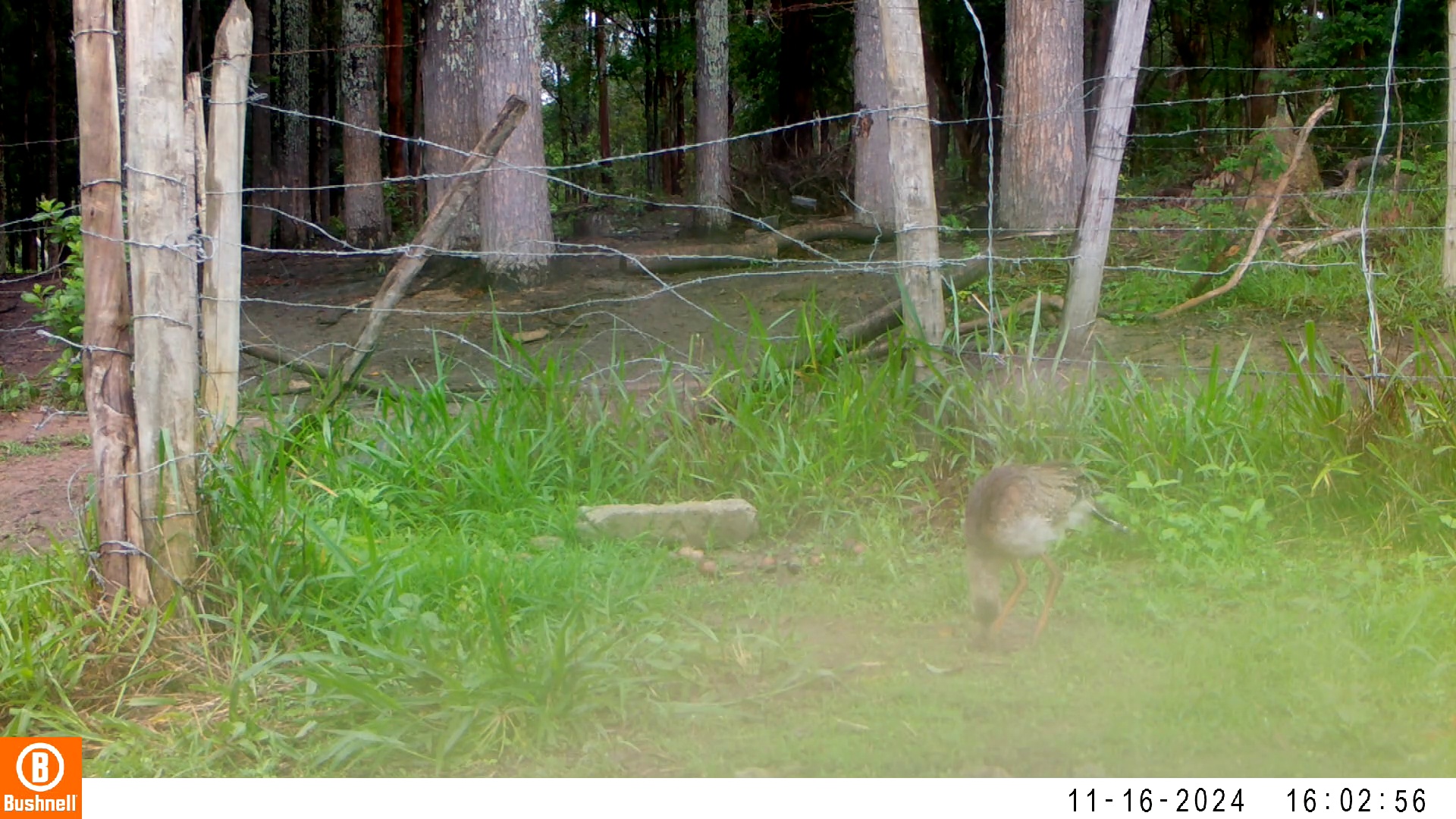

### 11170086-33174-906.jpg

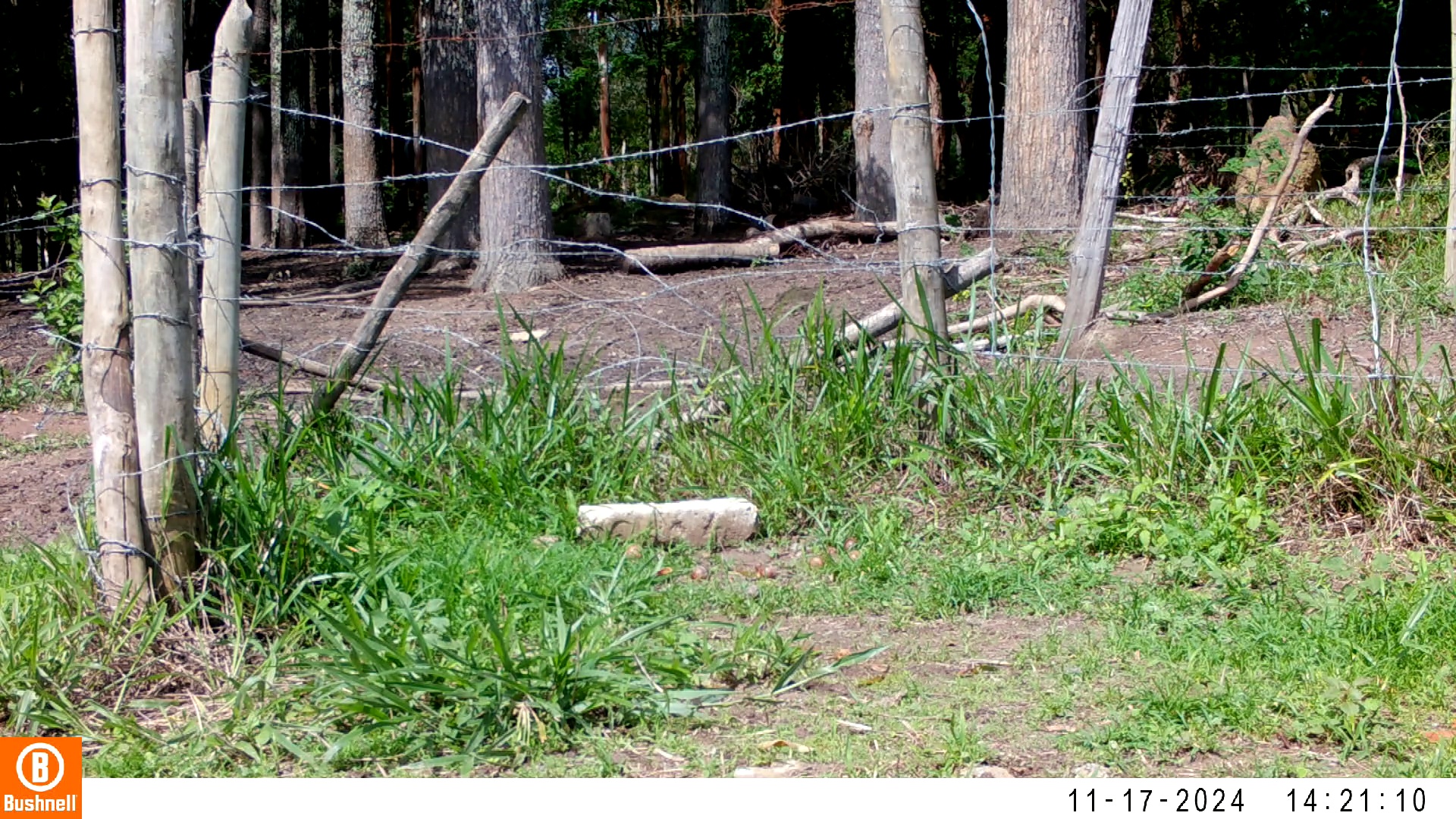

### 11170087-48404-1208.jpg

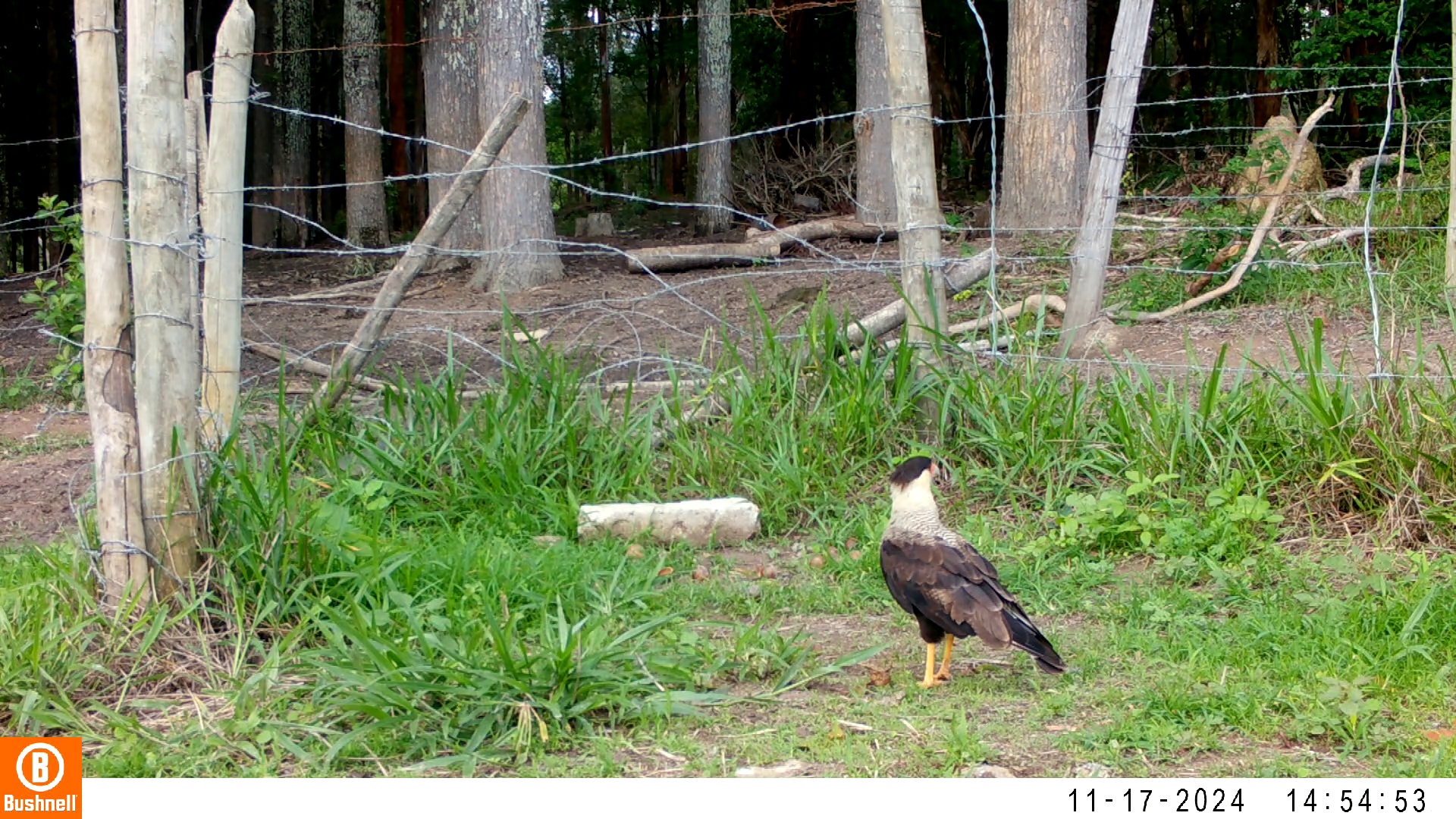

### 11180004-61866-906.jpg

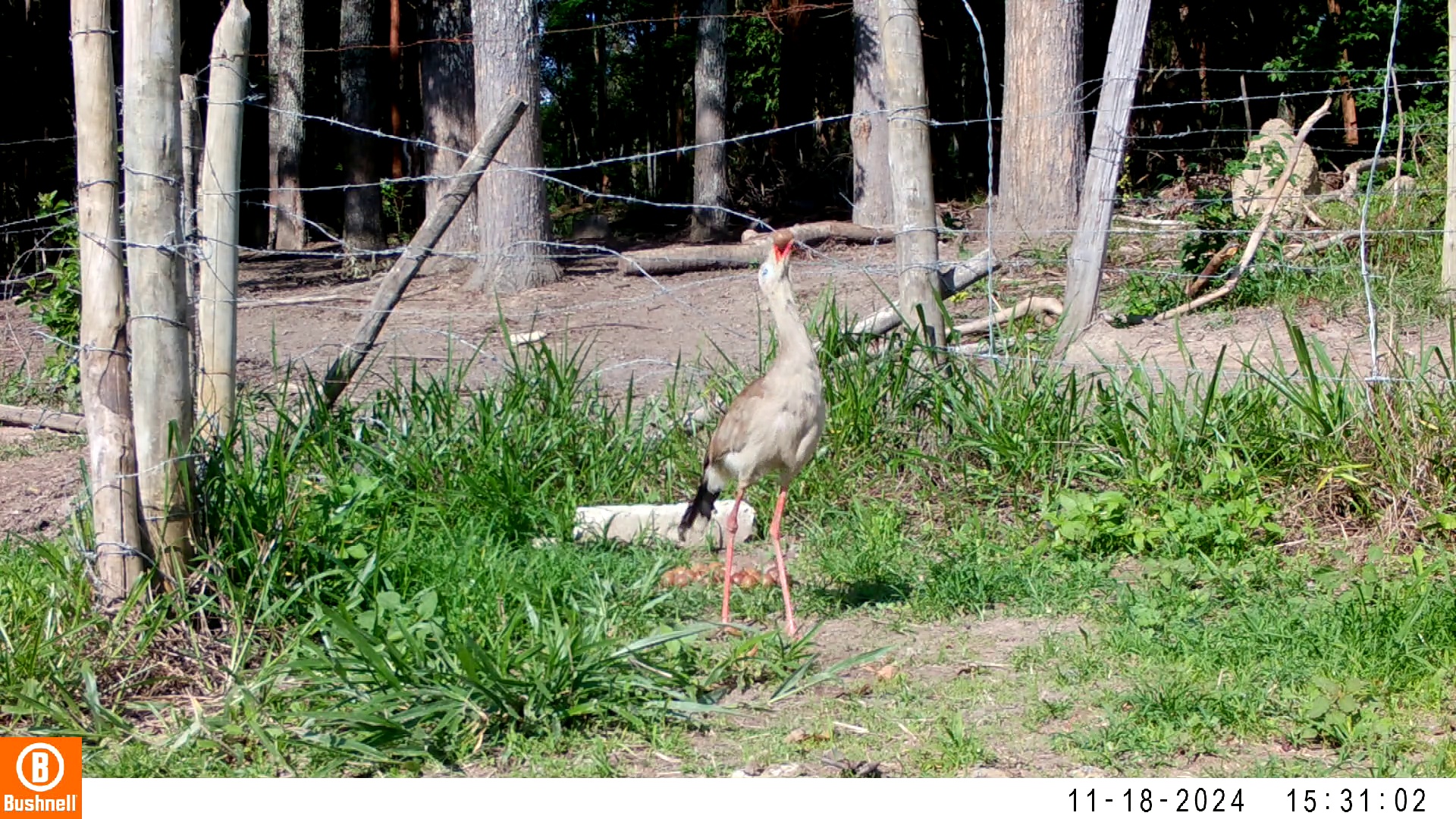

### 11190080-02410-0.jpg

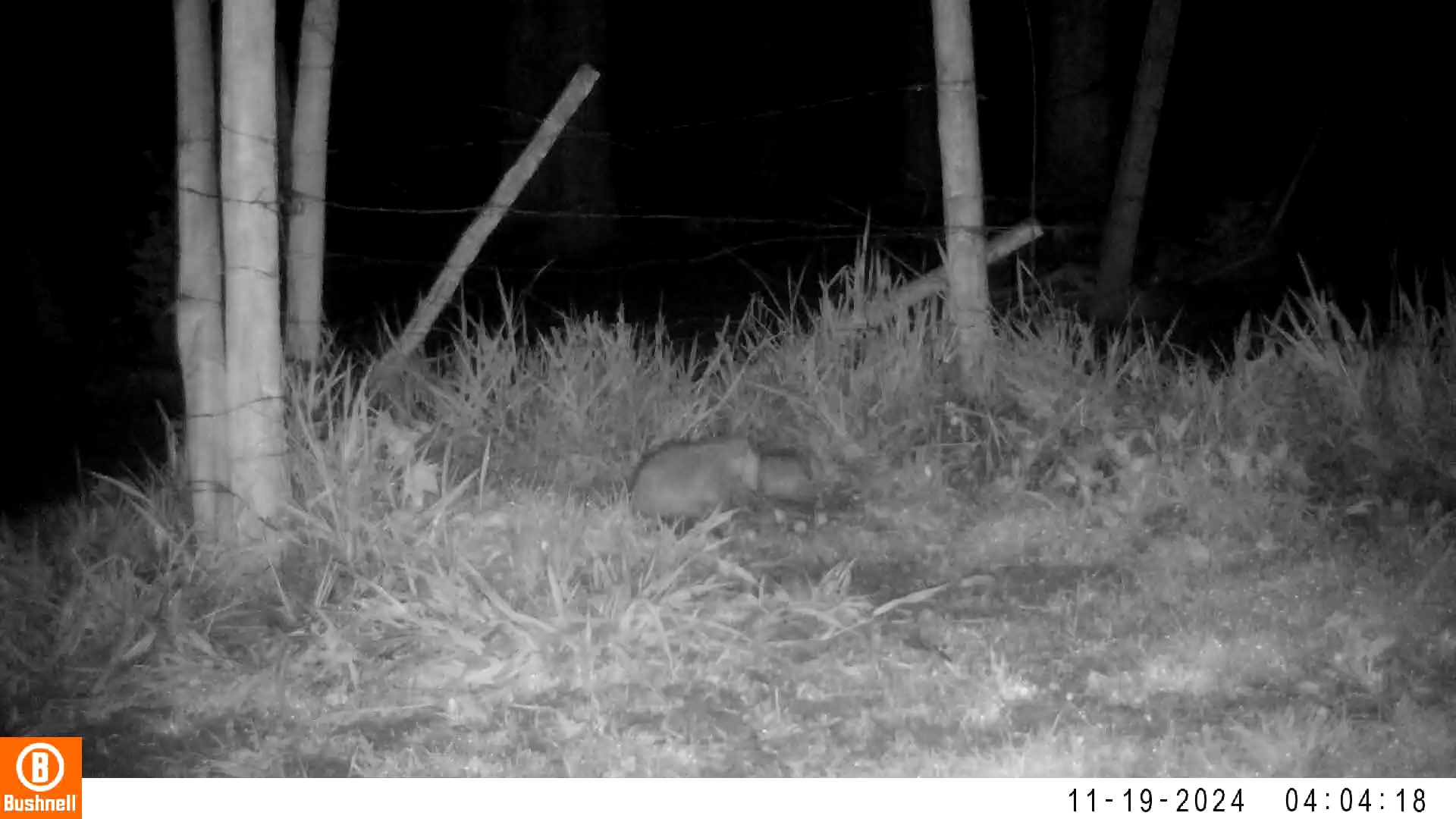

### 11190080-02410-302.jpg

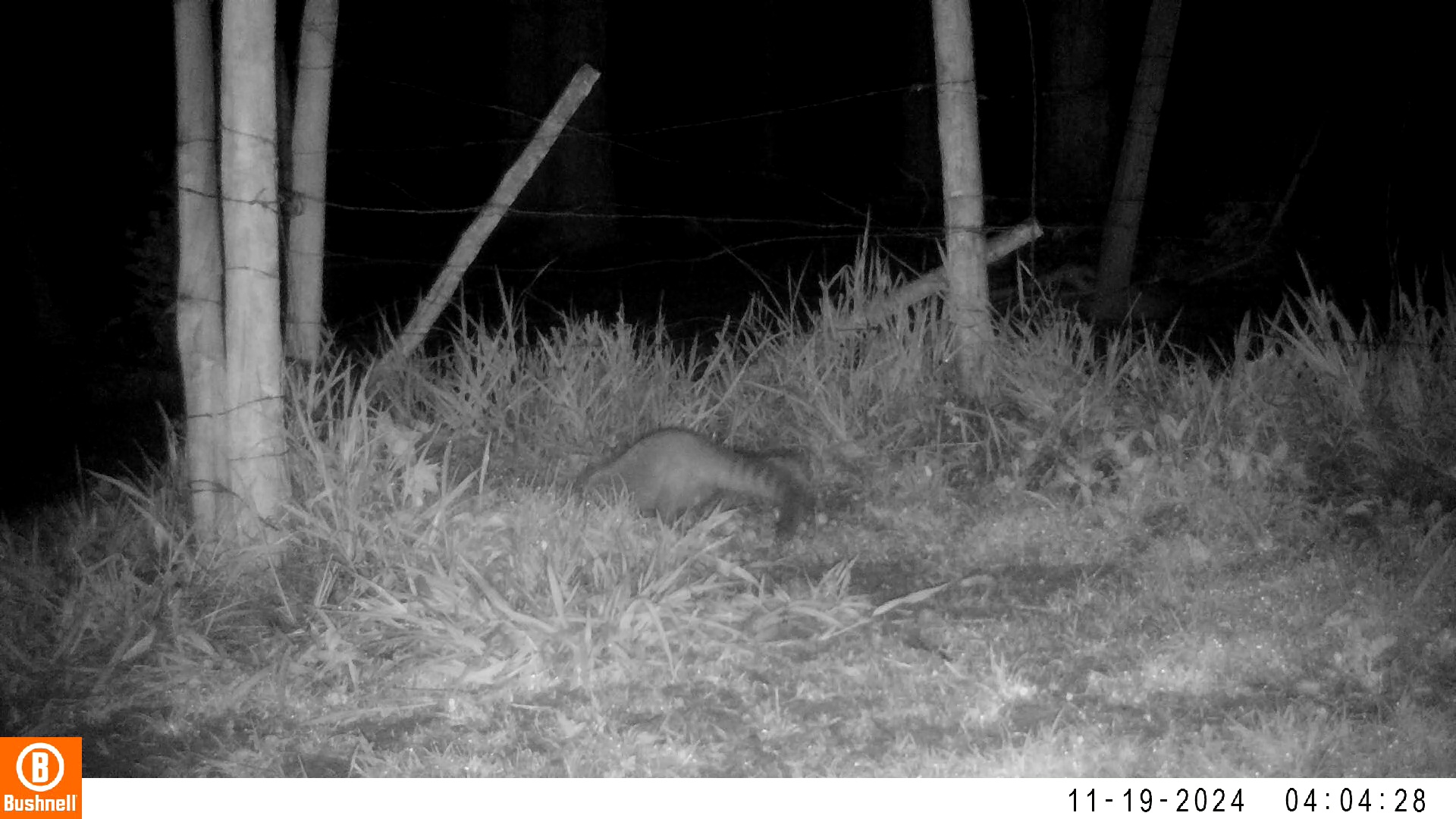

### 11190080-02410-1510.jpg

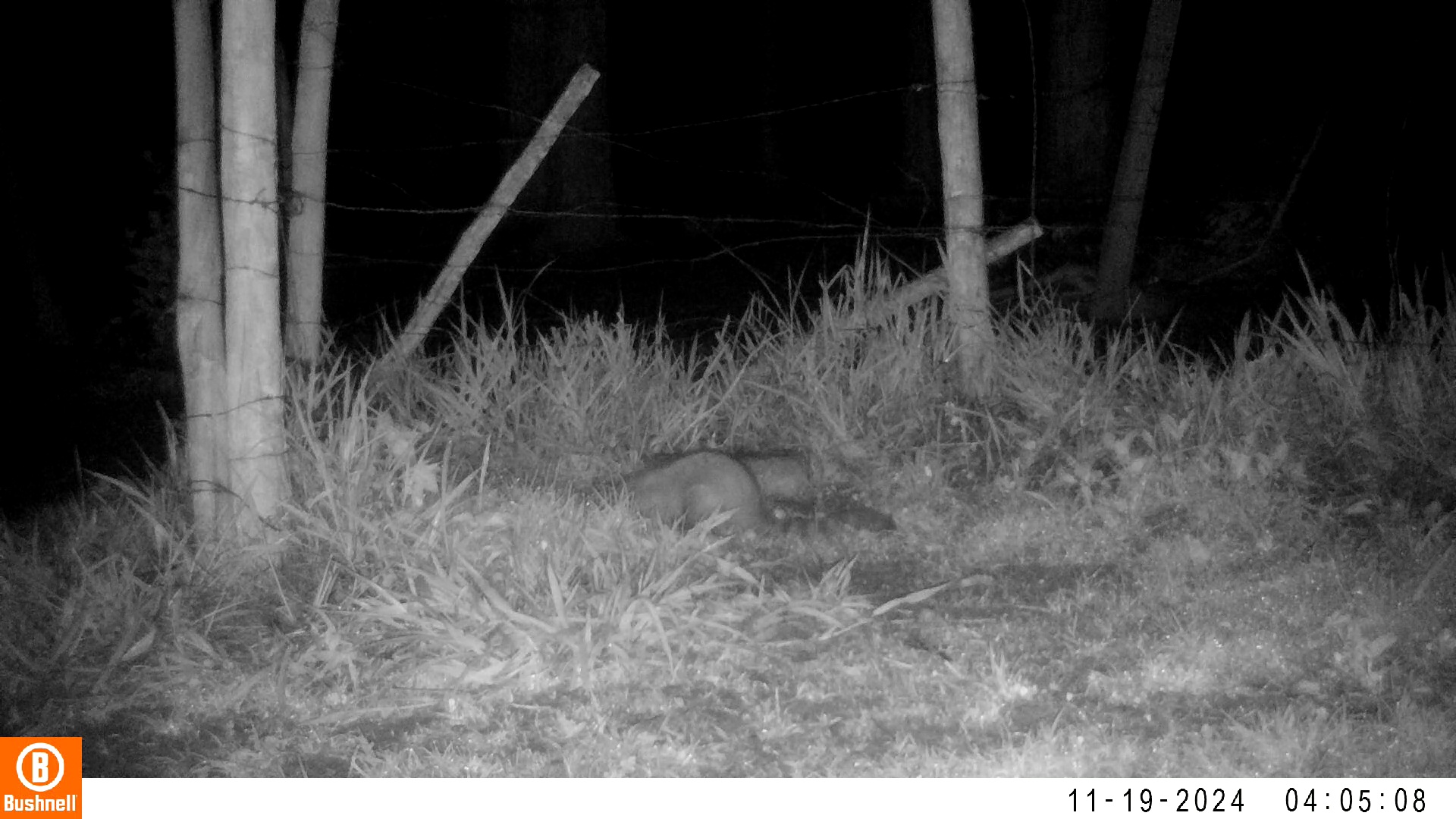

### 11190102-32683-1510.jpg

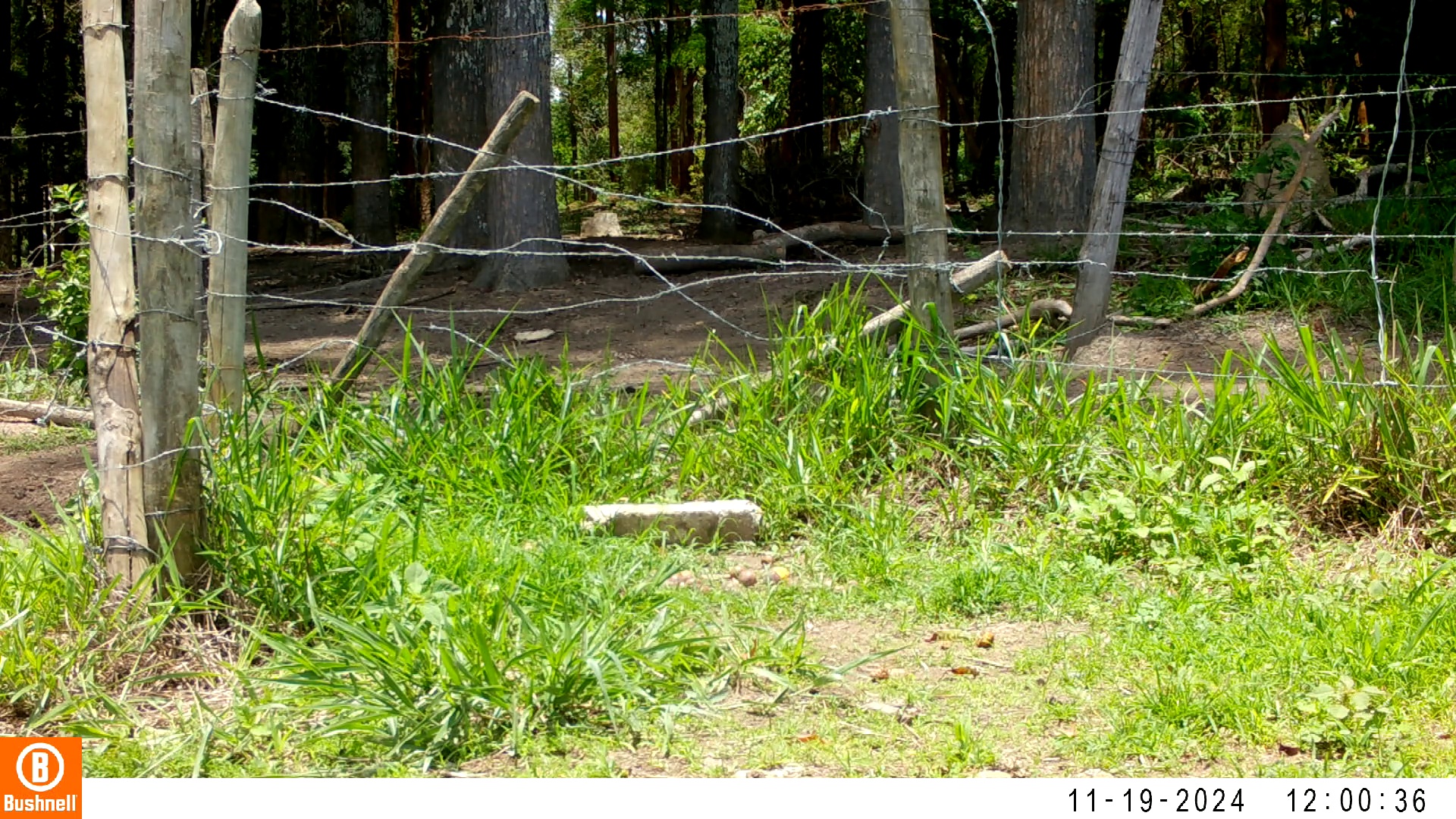

### 11210175-21978-1208.jpg

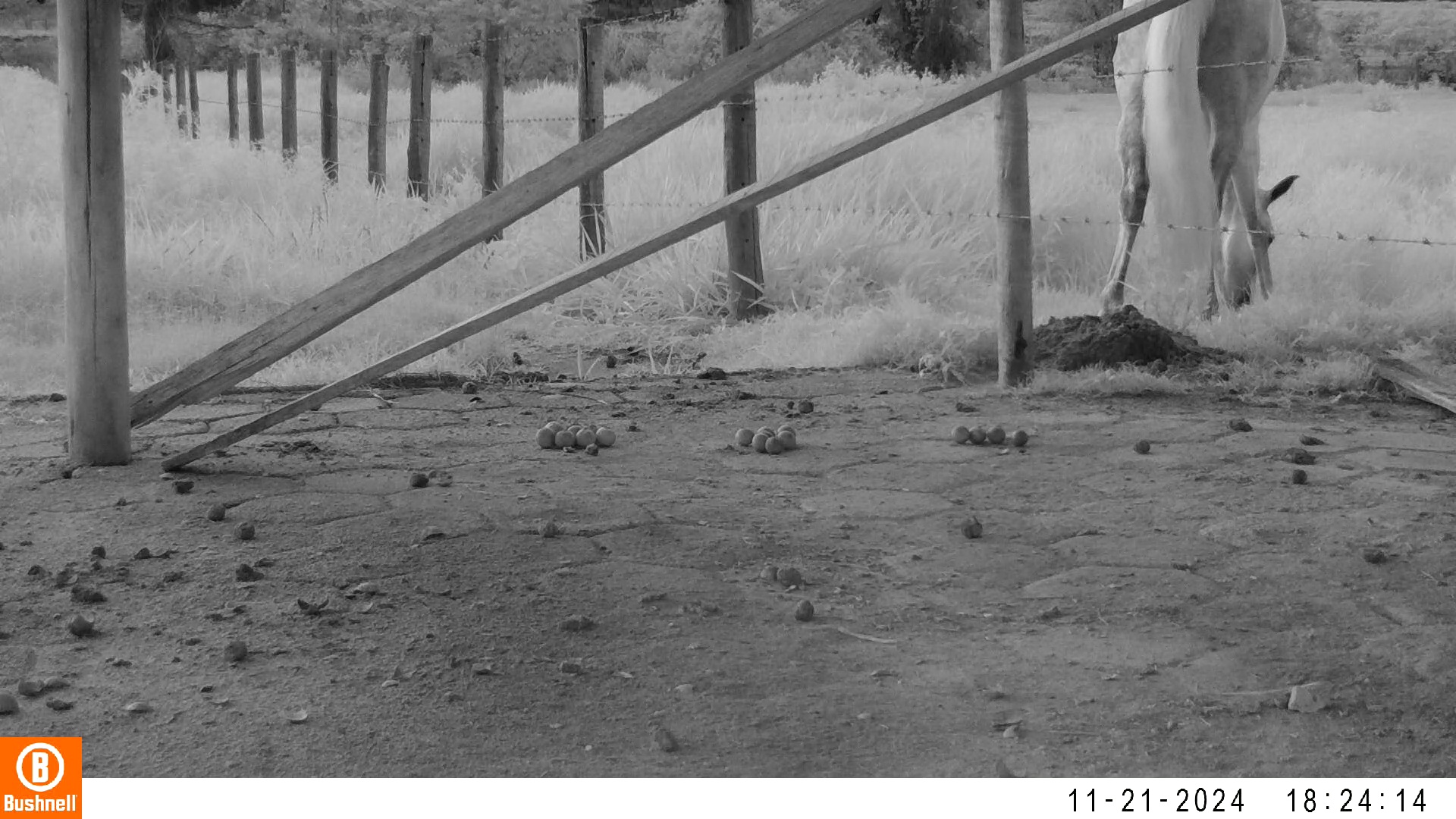

### 11220011-21818-1510.jpg

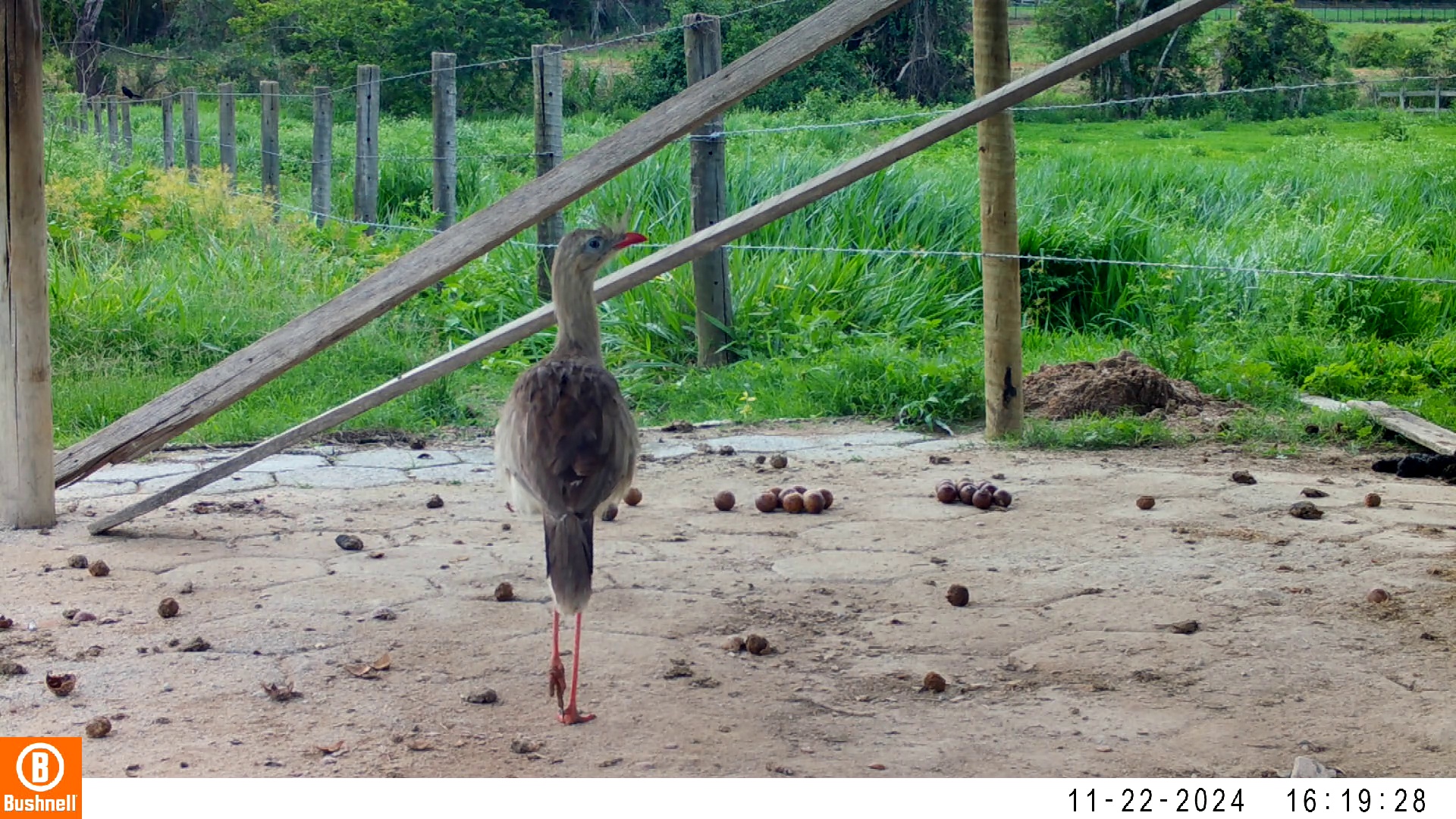

### 11230050-35931-1208.jpg

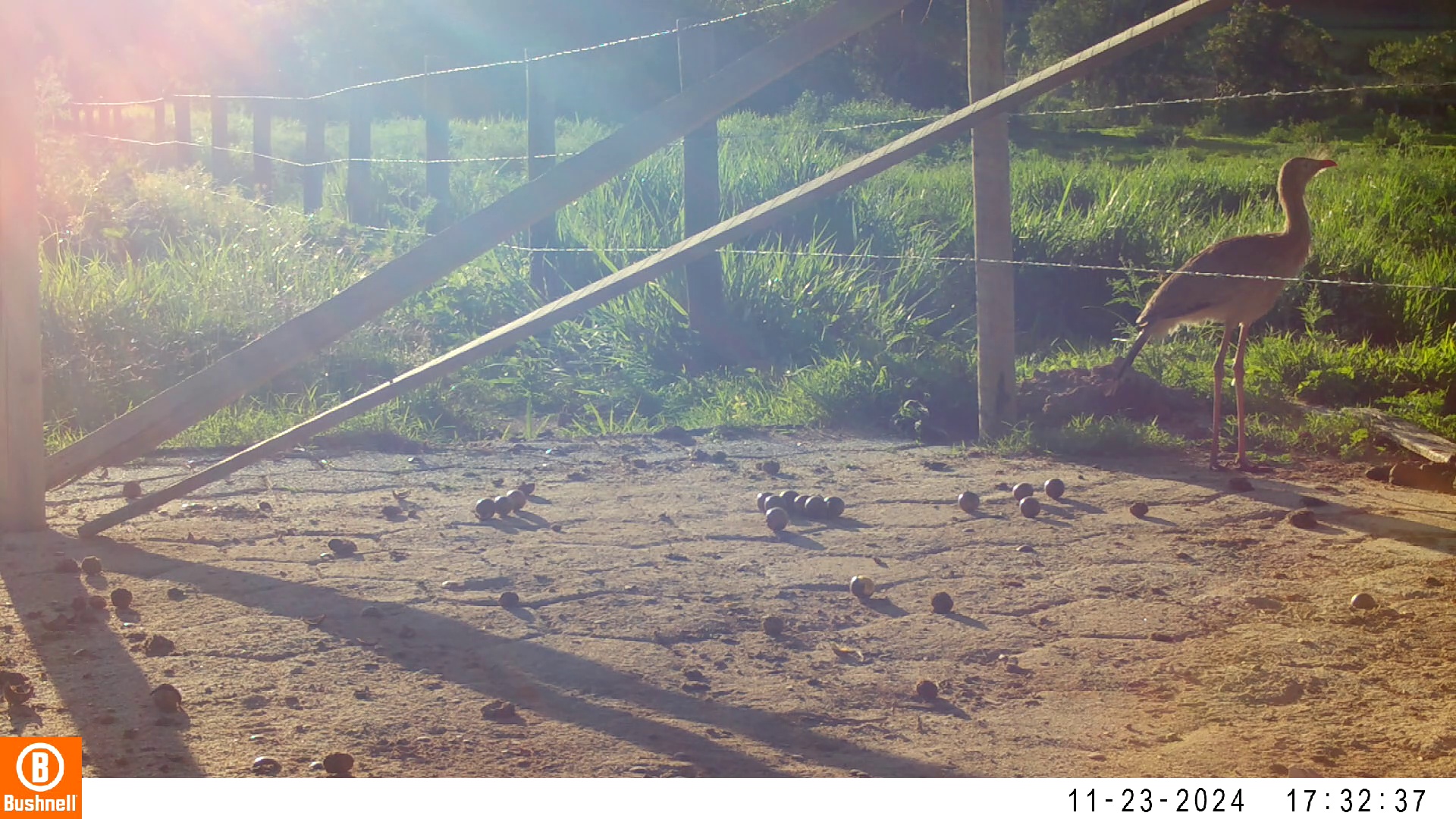

### 11240143-30572-906.jpg

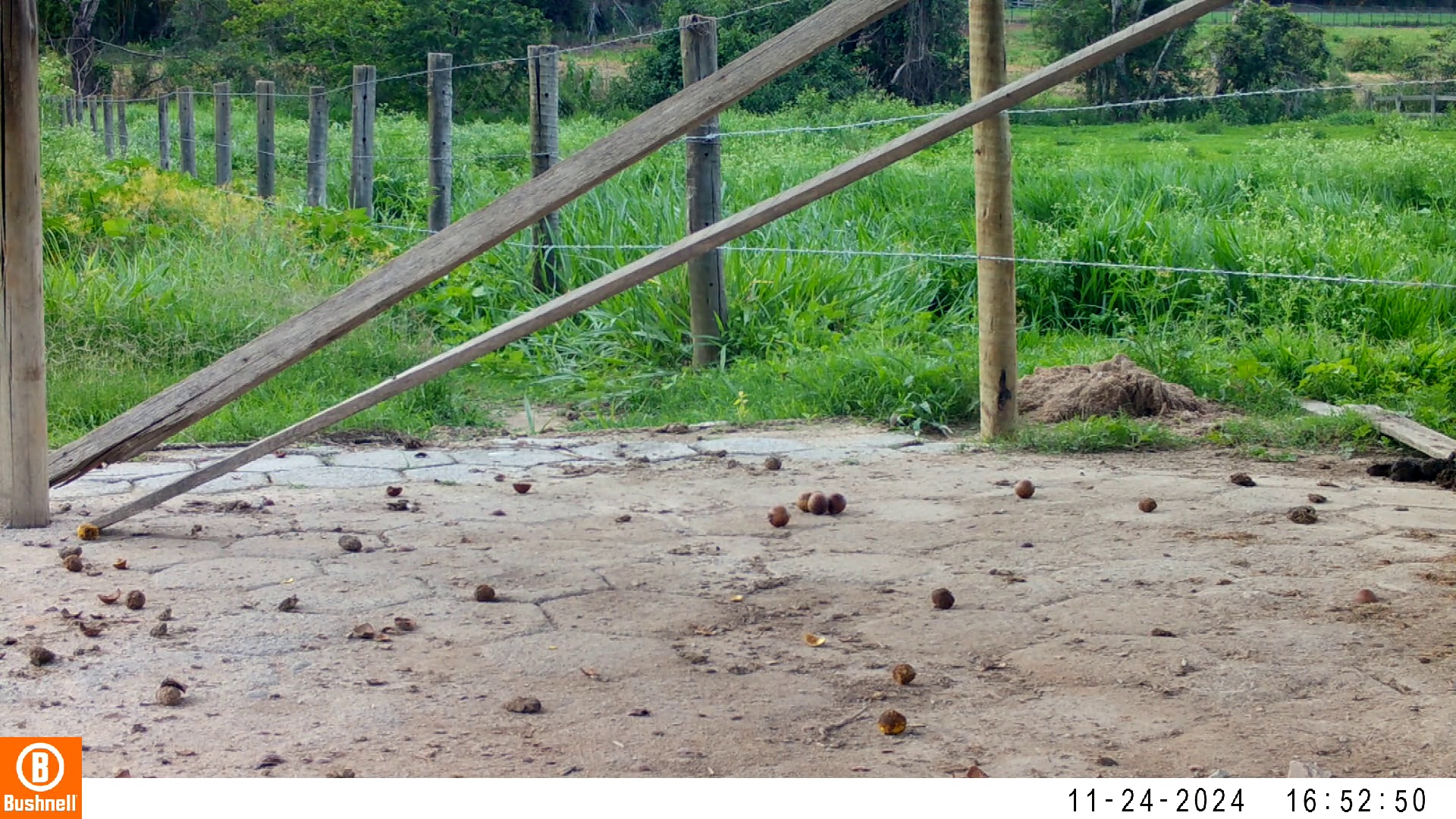

### 11240143-30572-1208.jpg

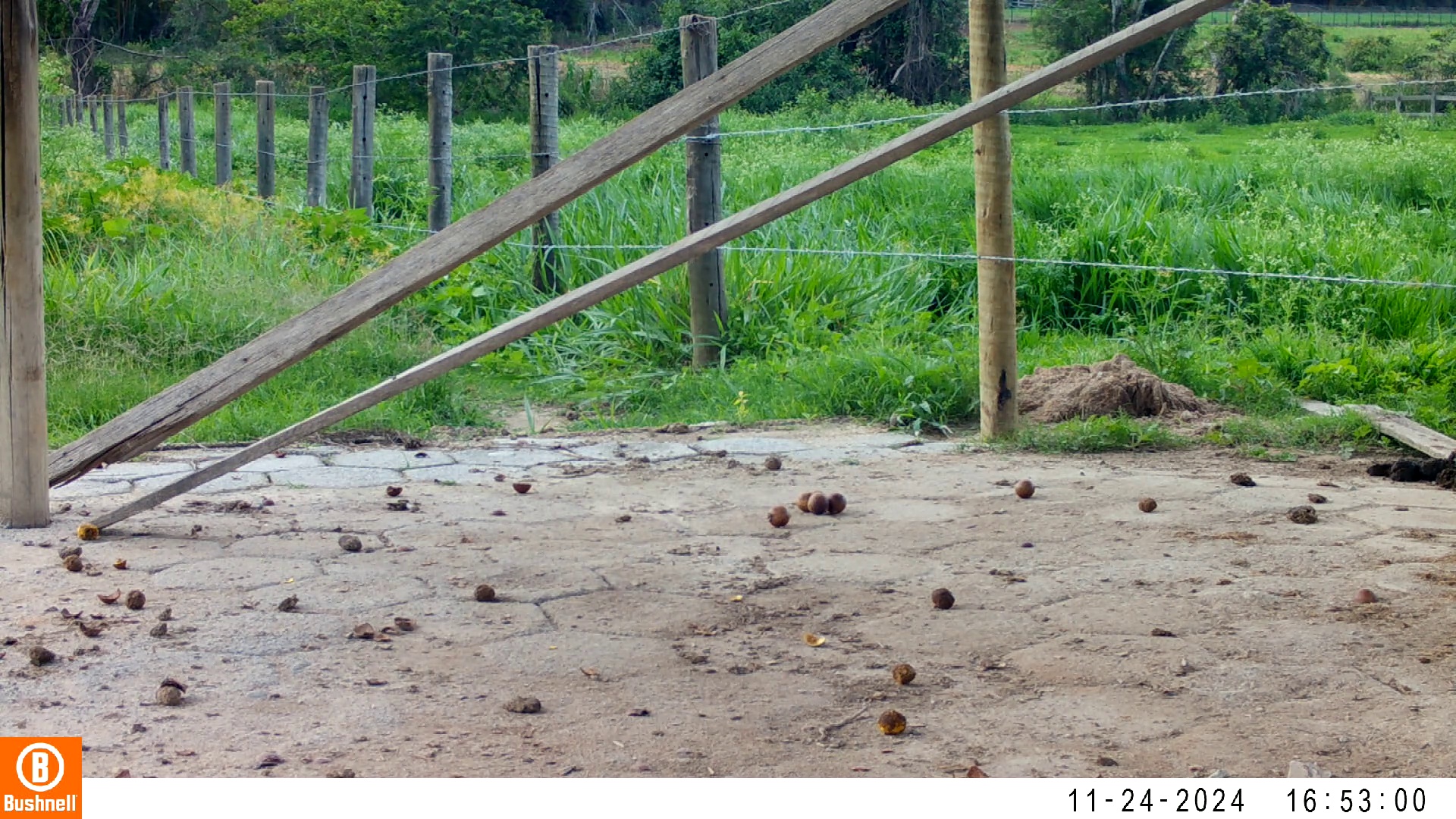

### 11250011-99191-1208.jpg

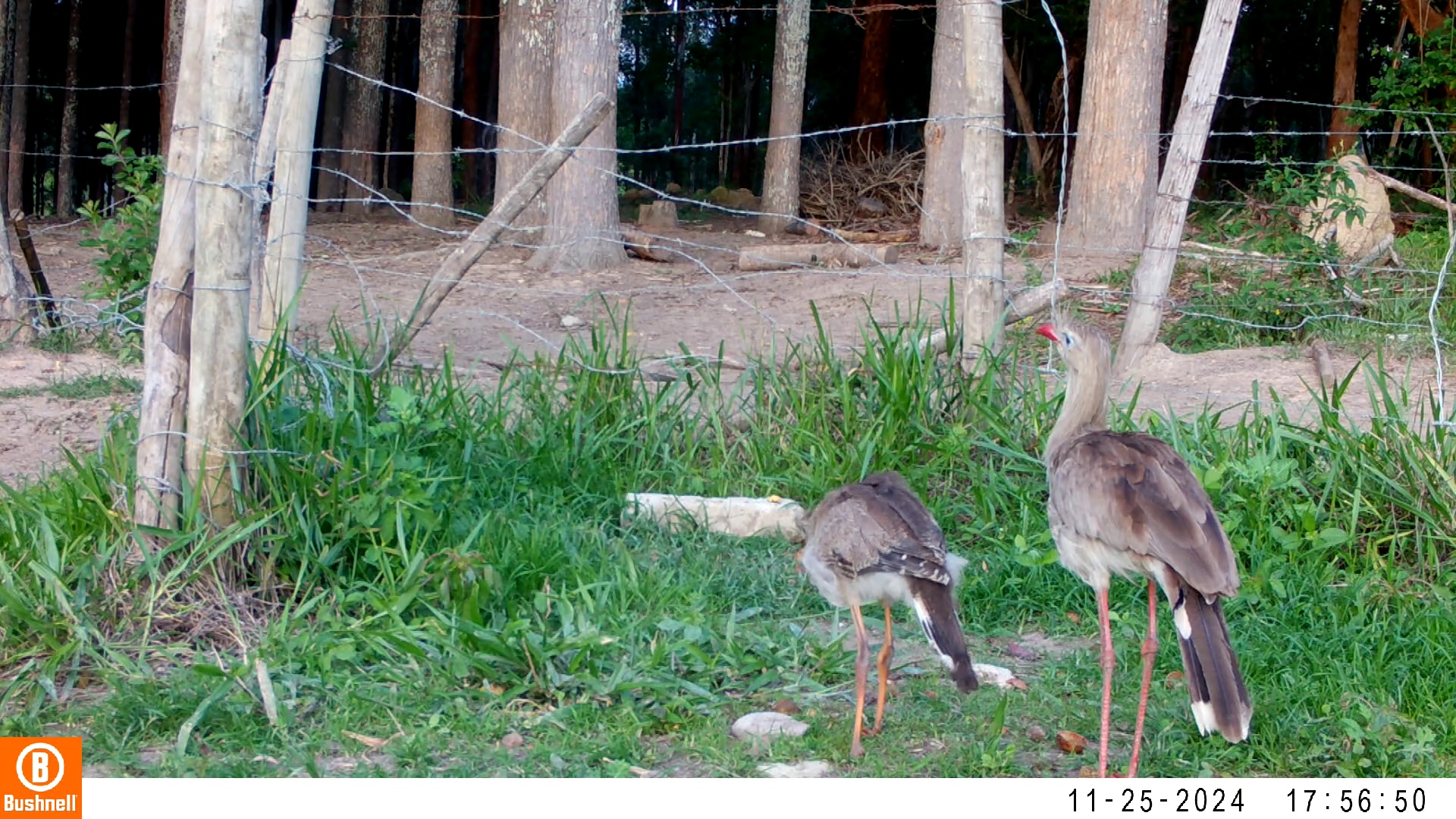

### 11250014-36058-302.jpg

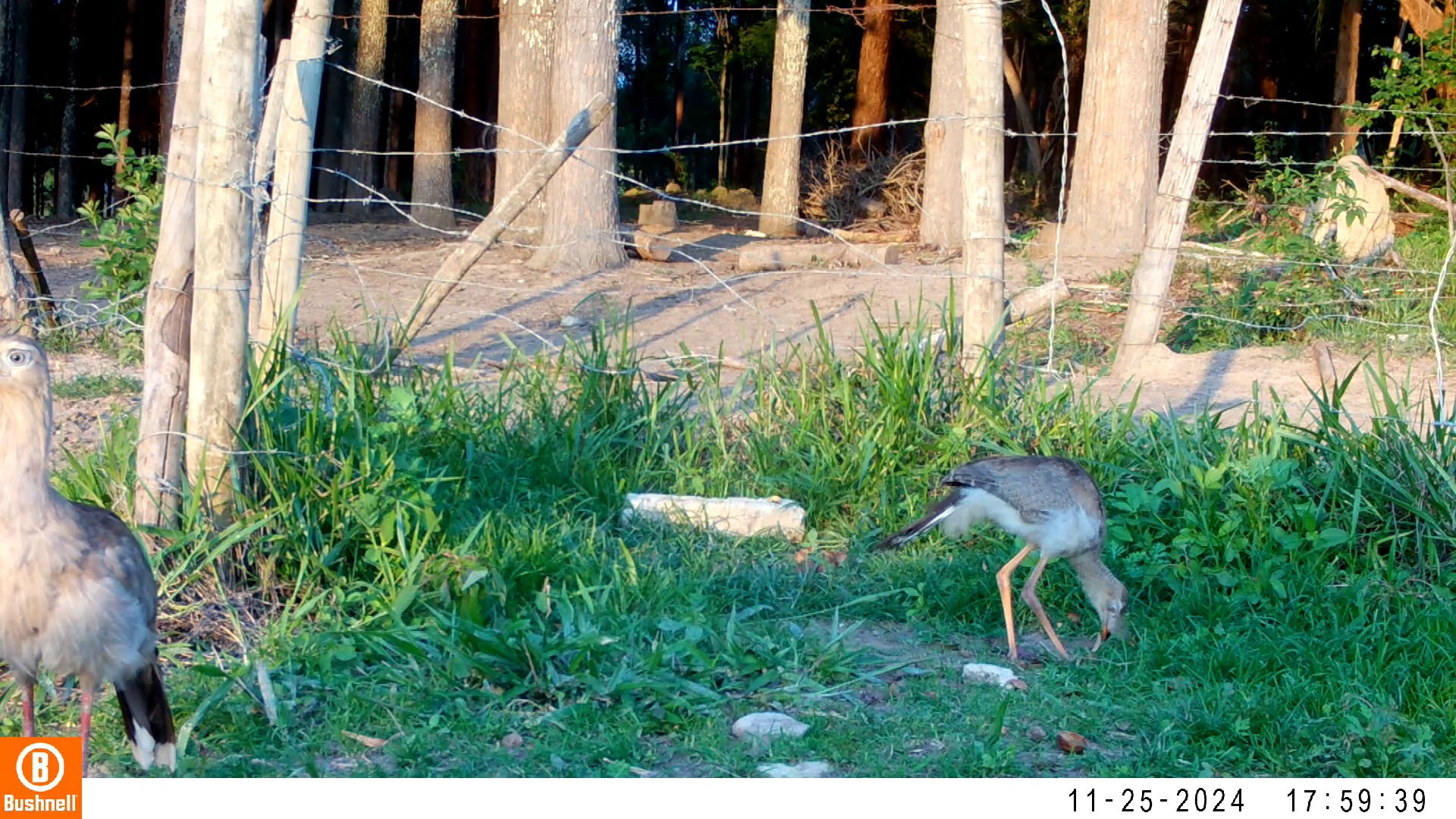

### 11250014-36058-604.jpg

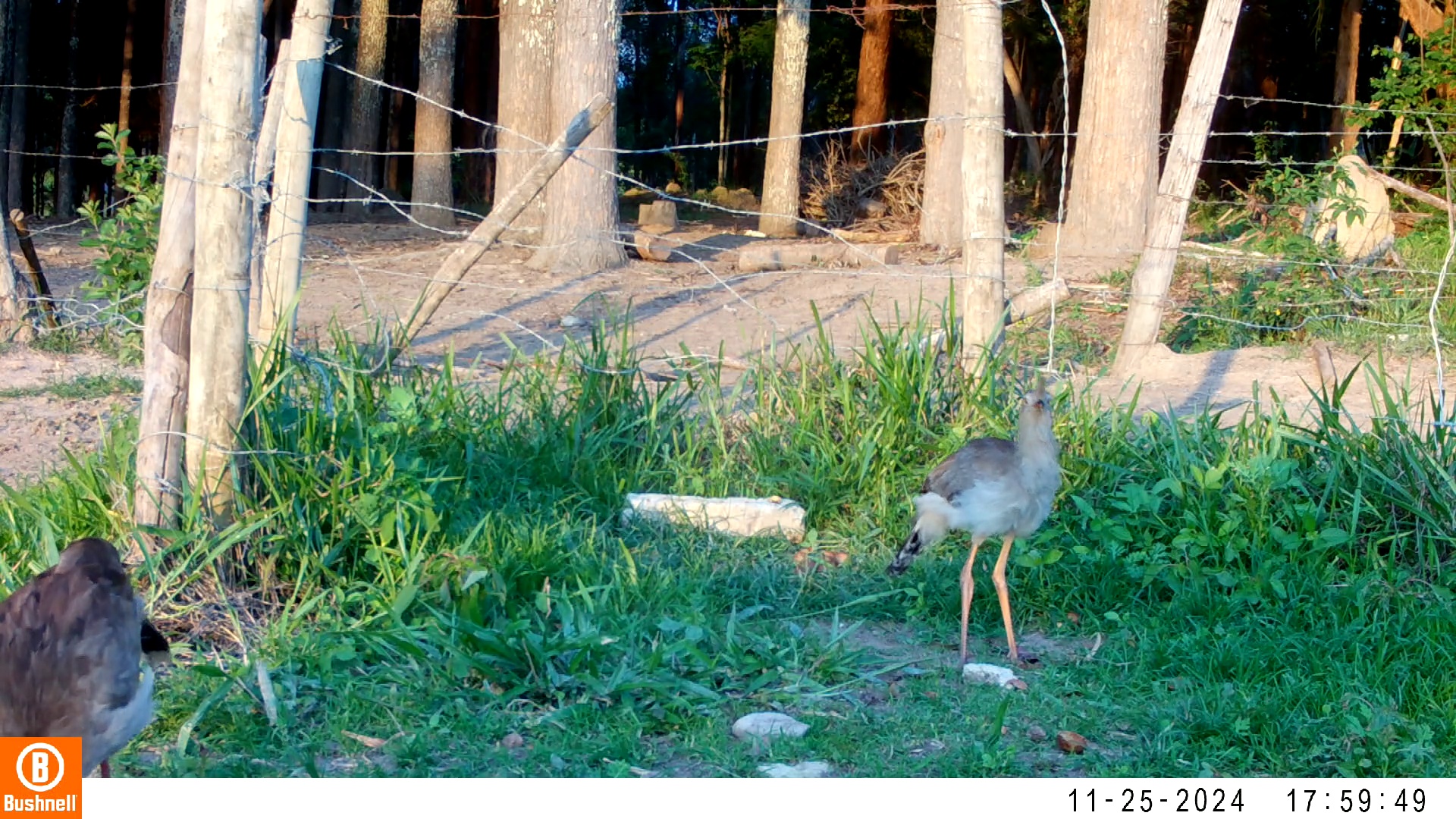
